# MicroRNA-29 Differentially Mediates Preeclampsia-Dysregulated Cellular Responses to Cytokines in Female and Male Fetal Endothelial Cells

**DOI:** 10.1101/2023.03.17.532827

**Authors:** Chi Zhou, Colman Freel, Olivia Mills, Xin-Ran Yang, Qin Yan, Jing Zheng

**Affiliations:** School of Animal and Comparative Biomedical Sciences, the University of Arizona, Tucson, AZ, United States; Department of Obstetrics and Gynecology, the University of Arizona, Tucson, AZ, United States; Department of Obstetrics and Gynecology, University of Wisconsin-Madison, Madison, WI, United States; University of Nebraska Medical Center, Omaha, NE, United States; Department of Obstetrics and Gynecology, Shanghai First Maternity and Infant Hospital, Tongji University School of Medicine, Shanghai, China

**Author notes:** **Corresponding Authors: Chi Zhou**, Ph.D., School of Animal and Comparative Biomedical Sciences, University of Arizona, 4101 N Campbell Ave., Tucson, AZ 85719, USA., **Jing Zheng**, Ph.D., Department of Obstetrics and Gynecology, University of Wisconsin-Madison, PAB1 UnityPoint Health-Meriter Hospital, 202 S. Park St., Madison, WI 53715, USA.

**Keywords:** Preeclampsia, MicroRNA-29, Cytokines, Fetal endothelial function, Sexual dimorphisms

## Abstract

**Introduction:** Preeclampsia (PE) differentially impairs female and male fetal endothelial cell function which is associated with the increased risks of adult-onset cardiovascular disorders in children born to mothers with PE. However, the underlying mechanisms are poorly defined. We **hypothesize** that dysregulation of microRNA-29a-3p and 29c-3p (miR-29a/c-3p) in PE disturbs gene expression and cellular responses to cytokines in fetal endothelial cells in a fetal sex-dependent manner.

**Methods:** RT-qPCR analysis of miR-29a/c-3p was performed on female and male unpassaged (P0) human umbilical vein endothelial cells (HUVECs) from normotensive (NT) and PE pregnancies. Bioinformatic analysis of an RNAseq dataset was performed to identify PE-dysregulated miR-29a/c-3p target genes in female and male P0-HUVECs. Gain- and loss-of-function assays were conducted to determine the effects of miR-29a/c-3p on endothelial monolayer integrity and proliferation in response to TGFβ1 and TNFα in NT and PE HUVECs at passage 1.

**Results:** PE downregulated miR-29a/c-3p in male, but not female P0-HUVECs. PE dysregulated significantly more miR-29a/c-3p target genes in female vs. male P0-HUVECs. Many of these PE-differentially dysregulated miR-29a/c-3p target genes are associated with critical cardiovascular diseases and endothelial functions. We further demonstrated that miR-29a/c-3p knockdown specifically recovered the PE-abolished TGFβ1-induced strengthening of endothelial monolayer integrity in female HUVECs, while miR-29a/c-3p overexpression specifically enhanced the TNFα-promoted cell proliferation in male PE HUVECs.

**Conclusions:** PE differentially dysregulates miR-29a/c-3p and their target genes associated with cardiovascular diseases- and endothelial function in female and male fetal endothelial cells, possibly contributing to the fetal sex-specific endothelial dysfunction observed in PE.

## Background

Preeclampsia (PE) is a hypertensive disorder that complicates 3∼8% of all human pregnancies(Anderson *et al*.) and costs billions of dollars to the U.S. healthcare system annually(Stevens *et al*.). PE is a leading cause of fetal and maternal morbidity and mortality during pregnancy(Askie *et al*.; Powe *et al*., 2011a), and adversely affects a wide range of fetal endothelial function such as cell proliferation, cell migration, monolayer integrity (or permeability), Ca^++^ response, and nitric oxide (NO) production(Wang *et al*.; Powe *et al*., 2011a; Boeldt *et al*.; Brodowski *et al*.; Zhou *et al*., 2019; Zhou *et al*., 2020). To date, the mechanisms underlying such fetal endothelial dysfunction in PE are poorly defined.

Pro-inflammatory cytokines are closely involved in the pathogenesis of PE(Raghupathy). For example, tumor necrosis factor-α (TNFα) can elevate blood pressure and induce proteinuria (two hallmarks of PE) in pregnant baboons(Sunderland *et al*.). Elevated maternal circulating TNFα is observed in several forms of pregnancy complications including PE(Wang & Walsh; Benyo *et al*.; Hung *et al*.; Raghupathy). TNFα expression is also significantly increased in human PE placentas(Wang & Walsh), contributing to impaired angiogenic activity in PE(Zhou *et al*.). Similarly, TNFα inhibits cell proliferation(Jiang *et al*.), migration, capillary tube formation(Hsu *et al*.), and downregulates endothelial nitric oxide synthase (eNOS)(Kim *et al*.), as well as affects cytokine-induced endothelial leukocyte adhesion molecule expression in human umbilical vein endothelial cells (HUVECs) in vitro(Collins *et al*.; Van Antwerp *et al*.; Mahboubi *et al*.). In addition, transforming growth factor-beta1 (TGFβ1), another growth factor and cytokine also regulates endothelial function, vascular development, and vascular barrier function(ten Dijke & Arthur; Walshe *et al*., 2009). In PE, TGFβ1 levels are significantly elevated in maternal circulation and are associated with PE-induced endothelial dysfunction(Muy-Rivera *et al*.; Peracoli *et al*.; Lau *et al*., 2013). MicroRNAs (miRNAs) are critical regulators of endothelial function (e.g., proliferation and migration)(Wu *et al*.; Zhou *et al*., 2017). MiR-29a-3p and miR-29c-3p (refer to as miR-29a/c-3p) are expressed in human endothelial cells and play important roles in maintaining endothelial function(Poliseno *et al*.; Wang *et al*.; Yang *et al*.). Dysregulation of miR-29a/c-3p is associated with cardiovascular diseases (e.g., stroke(Kajantie *et al*.) and heart failure(Thum *et al*.)), which may be partially due to their roles in endothelial cells. Overexpression of miR-29a-3p promotes angiogenesis, whereas knockdown of miR-29a-3p blocks TGFβ1-stimulated angiogenesis in a chick chorioallantoic membrane assay(Wang *et al*.). We have previously reported that PE downregulates miR-29a/c-3p in primary HUVECs and that knockdown of miR-29a/c-3p inhibits growth factor-stimulated fetal endothelial motility in HUVECs(Zhou *et al*.). However, it is unknown if PE differentially dysregulates the expression and function of miR-29a/c-3p in female (F) and male (M) HUVECs.

Increasing evidence has shown that sex is an important regulator of biological processes and cell function. Sexual dimorphisms of fetal endothelial function have been reported in HUVECs from normotensive (NT) pregnancies in cell proliferation, cell viability, tube formation capacity, migration, and endothelial eNOS expression(Addis *et al*.; Lorenz *et al*.). We have also demonstrated that PE differentially dysregulates F and M fetal endothelial transcriptomic profiles and endothelial cell responses (monolayer integrity and proliferation) to growth factors and cytokines (TNFα and TGFβ)(Zhou *et al*., 2019).

In this study, we tested the hypothesis that miR-29a/c-3p is differentially expressed and mediates PE-dysregulated cell responses to cytokines (TNFα and TGFβ) in fetal endothelial cells in a fetal sex-specific manner using HUVECs as a cell model. We determined the PE differentially dysregulated miR-29a/c-3p expression in F and M HUVECs and fetoplacental tissues from NT and PE pregnancies. Bioinformatics and gene ontology analyses were performed to identify PE-dysregulated miR-29a/c-3p target genes/pathways in F and M P0-HUVECs. We also examined the effect of miR-29a/c-3p knockdown and overexpression on endothelial monolayer integrity, cell proliferation, and migration in responses to cytokines in primary F and M HUVECs from NT and PE.

## Methods

The authors declare that all supporting data are available within the article and its online supplementary files.

### Ethical approval

All procedures were conducted in accordance with the Declaration of Helsinki. This research has been approved by the Institutional Review Board of the University of Arizona (Protocol# 2104723252) as well as the UnityPoint Health-Meriter Hospital (Madison, WI) and the University of Wisconsin-Madison (Protocol#2004-006). All subjects gave written, informed consent. PE was defined according to standard American College of Obstetricians and Gynecologists criteria(American College of *et al*., 2013). All PE patients included in this study were late-onset mild PE without major fetal morbidity. Due to the local demographics, all subjects included in this study are Caucasian.

### Isolation, purification, and characterization of primary HUVECs

F and M HUVECs were isolated immediately after deliveries of NT and PE pregnancies (Table.1). Cells were then purified using CD31 dynalbeads (Invitrogen, Carlsbad, CA) within ∼16h of culture as previously described(Zhou *et al*., 2017; Zhou *et al*., 2019). After purification, 80% of cells (referred to as P0-HUVECs) were immediately snap-frozen in liquid nitrogen until total RNA isolation, and the remaining 20% of cells were cultured to P1 (∼5 days of culture). The purity of each cell preparation was evaluated using the DiI-Ac-LDL uptake assay as previously described(Zhou *et al*., 2017; Zhou *et al*., 2019). Only cell preparations with ≥ 96% positive DiI-Ac-LDL uptake were used in this study.

**Table.1.** Patient Demographics*

| Diagnosis | Fetal sex | N | Maternal BMI | Maternal age (years) | Gestational age (weeks) | Systolic BP (mmHg) | Diastolic BP (mmHg) | Protein/Creatinine ratio (mg/ml) | Fetal weight (gram) | Apgar Score (5min) |
| --- | --- | --- | --- | --- | --- | --- | --- | --- | --- | --- |
| NT | F | 27 | 24.7 ± 0.6 | 32 ± 0.9 | 39.6 ± 0.2 | 109 ± 2.0 | 70 ± 1.8 | N/A | 3459 ± 73.6 | 9 ± 0.1 |
| PE | F | 15 | 24.9 ± 0.6 | 33 ± 1.5 | 37.3 ± 0.4 | 148 ± 4.3 | 90 ± 2.1 | 0.5 ± 0.5 | 3030 ± 143.4 | 9 ± 0.3 |
|  |  |  | P > 0.05 | P > 0.05 | P < 0.05 | P < 0.05 | P < 0.05 |  | P < 0.05 | P > 0.05 |
| NT | M | 20 | 24.1 ± 0.6 | 30.5 ± 1 | 39.3 ± 0.3 | 118 ± 3.3 | 73 ± 3.5 | N/A | 3645 ± 86.5 | 9 ± 0.1 |
| PE | M | 13 | 24.7 ± 0.7 | 30 ± 1.2 | 37.6 ± 0.5 | 143 ± 3.2 | 91 ± 2.7 | 0.6 ± 0.4 | 3150 ± 115.2 | 9 ± 0.2 |
|  |  |  | P > 0.05 | P > 0.05 | P < 0.05 | P < 0.05 | P < 0.05 |  | P < 0.05 | P > 0.05 |

\*BP: Blood Pressure; NT: Normotensive pregnancy; PE: Preeclampsia. All subjects are Caucasian.

### Human fetoplacental tissue collection

Human fetoplacental tissues were dissected from the placenta immediately after delivery. After dissection, fetoplacental tissue samples were snap-frozen in liquid nitrogen, stored at -80°C, and then ground to powder in liquid nitrogen followed by RNA isolation.

### RNA isolation and quality control

Small RNA-enriched total RNA samples were isolated from P0-HUVECs and placentas using the RNeasy Mini Kit (Qiagen, Valencia, CA). The concentration and quality of each RNA sample were assessed using a NanoDrop™ND-1000 spectrophotometer (NanoDrop Technologies, Wilmington, DE) and Agilent 2100-bioanalyzer (Agilent Technologies, Santa Clara, CA)(Zhou *et al*., 2017; Zhou *et al*., 2019). Only RNA samples with a high RNA integrity number (>8) were utilized in this study.

### RT-qPCR analysis of miR-29a/c-3p in human placentas and P0-HUVECs

To determine the PE-dysregulated miR-29a/c-3p expression in placentas and P0-HUVECs, RT-qPCR analysis was performed in F and M fetoplacental tissues and P0-HUVECs from NT and PE pregnancies (Table1; n=12-27cell preparations/group/sex) as previously described(Zhou *et al*., 2017). In brief, small RNA fragment enriched total RNA isolated from each sample was reverse transcribed into cDNA using a miScript II RT Kit (Qiagen). RT-qPCR was performed using miScriptSYBR Green PCR Kit (Qiagen) and commercially available miRNA Primer Assays (Table.S1) using a StepOne^Plus^ qPCR system (Life Technologies, Carlsbad, CA). Efficiencies of all target and control miRNA assays were between 90% and 110%. Data were first normalized to an external control (miRTC, Qiagen), followed by normalization to the geometric mean of endogenous control miRNAs (SNORD95, and SNORD96A). The normalized data were then analyzed using the 2^-ΔΔCT^ method(Yuan *et al*., 2006; Zhou *et al*., 2017).

### Bioinformatic analysis of RNAseq dataset on P0-HUVECs

We re-analyzed a previously published RNAseq dataset from P0-HUVECs (NCBI GEO accession: GSE116428(Zhou *et al*., 2019)) to identify PE dysregulated miR-29a/c-3p target genes in P0-HUVECs (Table.S2&S3). MiR-29a/c-3p target genes were determined using TarBase v.8(Karagkouni *et al*., 2018) (experiment supported miRNA targets database) and microT-CDS(Paraskevopoulou *et al*., 2013) databases. Functional genomics analysis of these PE-dysregulated miR-29a/c-3p target genes was performed to predict the enriched canonical pathways, diseases and biological functions, as well as gene networks using Ingenuity Pathway Analysis (IPA; www.qiagenbioinformatics.com)(Kramer *et al*., 2014; Zhou *et al*., 2019).

### Knockdown and overexpression of miR-29a/c-3p in F and M P1-HUVECs

Knockdown and overexpression of miR-29a/c-3p were performed using miScript miRNA mimics [Qiagen, refer as miRNA(+)] and miScript miRNA Inhibitor [Qiagen, refer as miRNA(i)] respectively, as described(Ukai *et al*., 2012; Zhou *et al*., 2017). Transfection dose and time were pre-determined as described(Zhou *et al*., 2017) (Fig.S1 and Supplemental Methods). MiRNA mimics and inhibitors were chemically synthesized and modified single-strand RNAs that specifically overexpress and inhibit target miRNA(Ukai *et al*., 2012; Zhou *et al*., 2017). In brief, individual primary P1-HUVECs at 50-60% confluence were transfected with miRNA mimic targeting human miR-29a/c-3p [miR-29a/c-3p(+); MSY0000086; Qiagen] and miRNA inhibitor targeting human miR-29a/c-3p [miR-29a/c-3p(i); MIN0000681; Qiagen] using the Qiagen HiPerFect Transfection Reagent for 24hr. Cells transfected with only the transfection reagent and miScript inhibitor negative control were used as the vehicle (Veh) and negative control (NC), respectively(Zhou *et al*., 2017). RT-qPCR was used to verify the efficiency of miRNA knockdown and overexpression.

### Cell functional assays

After successful knockdown and overexpression of miR-29a/c-3p, P1-HUVECs were treated with TNFα (10ng/ml), TGFβ (10ng/ml), or serum-free endothelial culture media (ECMb, control) followed by cell functional assays as previously described(Zhou *et al*., 2019) (n=5∼10 cell preparations/sex/group; Supplemental methods). The dose and duration of TNFα and TGFβ treatments for each cell functional assay were determined based on our previous reports(Zhou *et al*., 2017; Zhou *et al*., 2019). Endothelial monolayer integrity was determined using the ECIS Zθ+96-well array station (Applied BioPhysics, NY) using 96W10idf plates(Zhou *et al*., 2019). Cell proliferation was assessed using the CCK-8 kit (Dojindo Molecular Technologies, Rockville, MD)(Zhou *et al*., 2019).

### Statistical analyses

SigmaPlot software (Systat Software., San Jose, CA) was used for statistical analyses. Data are represented as the medians ± standard deviation (SD). Data analyses were performed using the Mann-Whitney Rank Sum Test or Kruskal-Wallis test as appropriate. Differences were considered significant when *P* <0.05. Benjamini and Hochberg False Discovery Rate (FDR)-adjustment(Zhou *et al*., 2017; Zhou *et al*., 2019) was used for multiple comparison correction as appropriate.

## Results

### PE differentially dysregulates miR-29a/c-3p expression in F and M placentas and HUVECs

Compared with NT, PE increased the levels of miR-29a-3p and miR-29c-3p by 72% and 106%, respectively, in F, but not M placentas (Fig.1A). Both miR-29a-3p and miR-29c-3p levels were significantly higher in F than in M PE placentas. In contrast, PE decreased the levels of miR-29a-3p and miR-29c-3p by 15% and 24%, respectively, in M but not F P0-HUVECs (Fig.1B). There were no significant differences in miR-29a-3p and miR-29c-3p levels between F and M placentas and P0-HUVECs from NT. However, miR-29a-3p and miR-29c-3p levels were significantly higher in F than in M placentas, while miR-29c-3p levels were higher in F vs. M PE P0-HUVECs.

**Figure 1.**
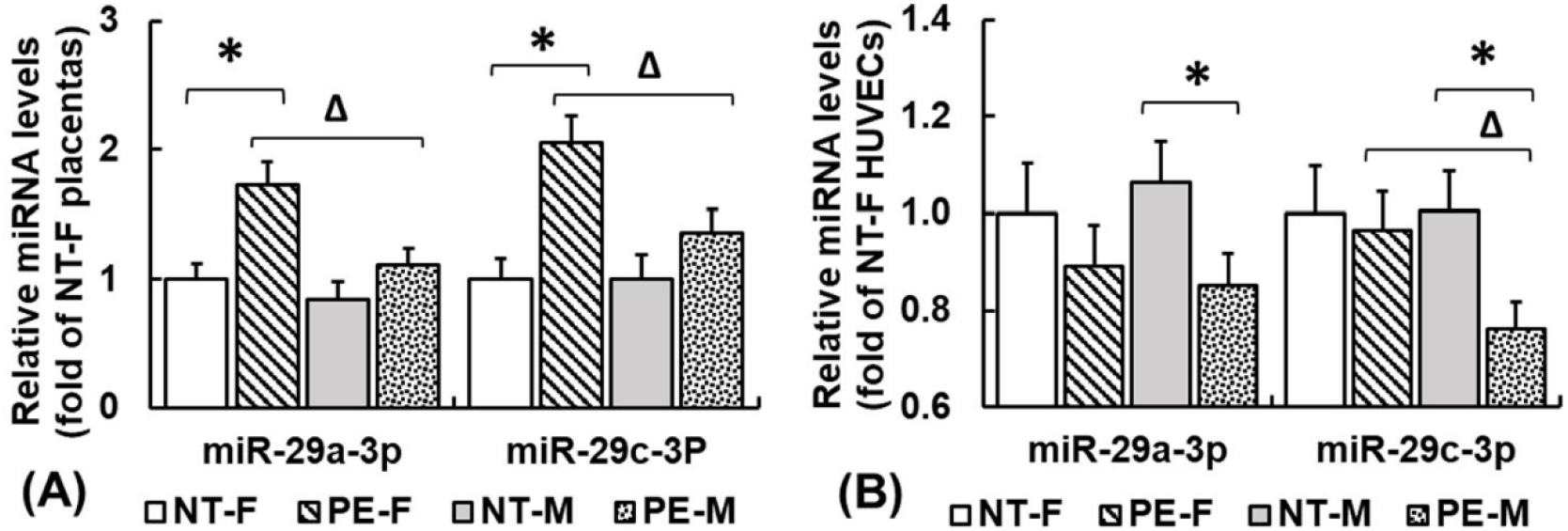
PE differentially dysregulates miR-29a/c-3p in F and M placentas (A) and P0-HUVECs (B). (A) Human placentas were collected immediately after delivery (NT-F, n=18; PE-F, n=12; NT-M, n=20; PE-M, n=14). (B) Individual primary P0-HUVECs preparations were isolated immediately after delivery and isolated within 16h (NT-F, n=27; PE-F, n=15; NT-M, n=20; PE-M, n=13). Data are expressed as fold of the corresponding NT-F group (*P* < 0.05, Mann-Whitney Rank Sum Test). **\***Differ between PE vs. NT in same fetal sex; ^**Δ**^Differ between F and M within the same diagnostic group (PE or NT).

### PE differentially dysregulates miR-29a/c-3p target genes in F and M HUVECs

Re-analysis of our previously published RNAseq dataset on P0-HUVECs (NCBI GEO accession: GSE116428(Zhou *et al*., 2019)) have identified 2883 (2357 experimentally supported and 526 theoretically predicted, Table.S2) miR-29a/c-3p target genes in P0-HUVECs (Fig. 2B). Among them, 1631 were common target genes of miR-29a/c-3p, while 477 and 775 were miR-29a-3p and miR-29c-3p specific target genes, respectively (Table.S2). Our analysis also showed that PE differentially dysregulated miR-29a/c-3p target genes in F and M P0-HUVECs (Fig.2A-C, Table.S3). A total of 125 miR-29a/c-3p target genes were differentially expressed (DE) between NT-F and NT-M P0-HUVECs (Fig.2B-C, Table.S3), among which two and one genes are located on X- and Y-chromosomes, respectively (Table.S3). In PE-F P0-HUVECs, 110 miR-29a/c-3p target genes were dysregulated with two DE genes located on X-chromosome (Fig.2B-C, Table.S3). In PE-M P0-HUVECs, 28 miR-29a/c-3p target genes were dysregulated with two DE-genes located on the X-chromosome; none of these DE genes located on the Y-chromosome (Fig.2B-C, Table.S3). Only five miR-29a/c-3p target genes (ARC, CILP2, NEFM, GRIN2B, and CHRNA7) were dysregulated in both PE-F and PE-M P0-HUVECs (Fig.2B-C, Table.S3). All these five genes were down-regulated in PE-F, but up-regulated in PE-M P0-HUVECs (Table.S3).

**Figure 2.**
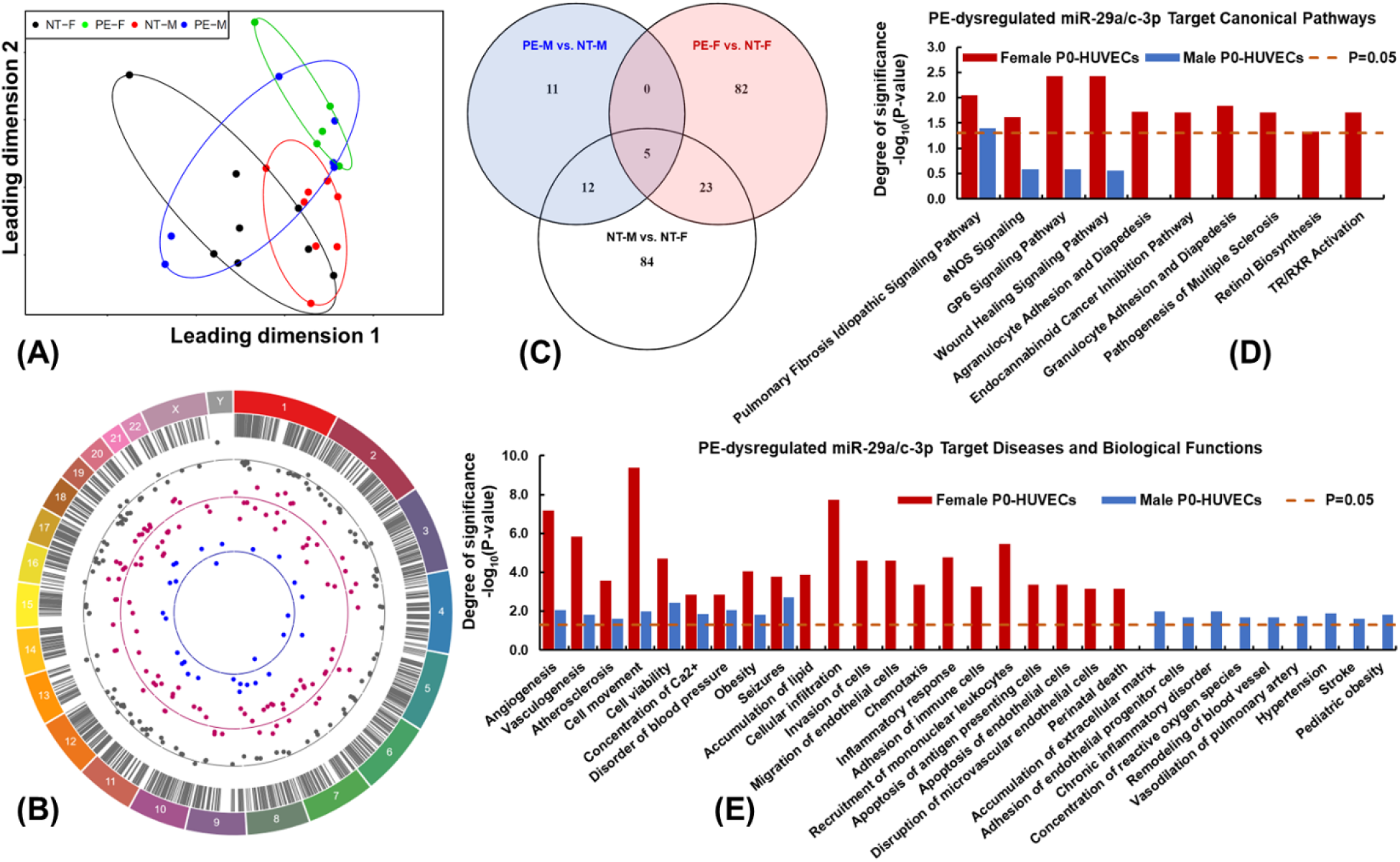
PE differentially dysregulates miR-29a/c-3p target genes. **(A)** Multi-Dimensional Scaling (MDS) plot representing the similarity and disparity among samples based on expression patterns of miR-29a/c-3p target genes. Each dot represents one biological sample. Distance between dots representing the differences of miR-29a/c-3p target genes expression profiles among samples. Eclipse shows the clustering of samples. **(B)** Circos plot illustrating the chromosomal position of DE miR-29a/c-3p target genes between NT-M vs. NT-F (grey dots, 125 DE-genes), PE-F vs. NT-F (pink dots, 110 DE-genes), and PE-M vs. NT-M (blue dots, 28 DE-genes). Each dot represents one gene. The numbers and letters in the outer ring indicate the chromosomal location. For each scatter plot track, dots outside and inside of the centerline are up- and down-regulated miR-29a/c-3p target genes by PE, respectively. **(C)** Overlap of DE miR-29a/c-3p target genes in NT-M vs NT-F, PE-F vs. NT-F, and PE-M vs. NT-M. Preeclampsia differentially dysregulated **(D)** canonical pathways-, as well as **(E)** diseases and biological functions-associated miR-29a/c-3p target genes in F and M P0-HUVECs. Significant enrichments were determined using IPA software (*P* < 0.05, Fisher’s exact test followed with BH-FDR multiple test correction).

### PE differentially dysregulates cardiovascular diseases- and endothelial function-associated miR-29a/c-3p target genes in F and M HUVECs

Canonical pathways enrichment analysis on PE-dysregulated genes in F and M P0-HUVECs (Fig.2D and S2) indicated that only one miR-29a/c-3p target canonical pathway (pulmonary fibrosis idiopathic signaling pathway) was significantly dysregulated in both PE-F and PE-M HUVECs. Nine canonical pathways were enriched only in PE-F HUVECs, and six of these pathways including wound healing signaling pathway and pathogenesis of multiple sclerosis were enriched only in PE-upregulated miR-29a/c-3p target genes in PE-F HUVECs.

Diseases and Bio-function enrichment analysis on PE-dysregulated genes in P0-HUVECs (Fig.2E and S3, Table.S4, S5 and S6) further showed that miR-29a/c-3p target genes associated with angiogenesis, atherosclerosis, cell viability, concentration of Ca^2+^, disorder of blood pressure, obesity, and seizures were dysregulated in PE-F and PE-M HUVECs. Specifically, atherosclerosis, concentration of Ca^2+^, disorder of blood pressure, and seizures-associated miR-29a/c-3p target genes were only enriched in PE-upregulated genes in M-HUVECs, while concentration of Ca^2+^, cell viability, and seizures-associated miR-29a/c-3p target genes were only enriched in PE-downregulated genes in F-HUVECs (Fig.S3, Table S5). Additionally, miR-29a/c-3p target genes associated with accumulation of lipid, cellular infiltration, invasion of cells, migration of endothelial cells, chemotaxis, inflammatory response, adhesion of immune cells, recruitment of mononuclear leukocytes, apoptosis of antigen presenting cells, and disruption of microvascular endothelial cells were only enriched in PE-upregulated genes in F-HUVECs. However, miR-29a/c-3p target genes associated with accumulation of extracellular matrix, adhesion of endothelial progenitor cells, concentration of reactive oxygen species, remodeling of blood vessel, vasodilation of pulmonary artery, hypertension, and stroke were only enriched in PE-upregulated genes in M-HUVECs.

Upstream regulator analysis on PE-dysregulated genes in P0-HUVECs (Table.2, Table.S7) revealed that TNF-, TGFB1-, IFNG-, IL1B-, MYC-, AGT-, FOXO1-, MAPK1-, PDGF BB-, F2-, and CSF2-regulated miR-29a/c-3p target genes were enriched in both PE-F and PE-M HUVECs, in which PE-F HUVECs has more PE-dysregulated miR-29a/c-3p target genes in all these gene networks than PE-M cells. In addition, NFκB-, IL33-, and IL1-regulated miR-29a/c-3p target genes were uniquely enriched in PE-F HUVECs.

**Table.2.** PE-dysregulated miR-29a/c-3p target gene network in female and male P0-HUVECs*

| Gene Network | PE vs. NT Female P0-HUVECs |  | PE vs. NT Male P0-HUVECs |  |
| --- | --- | --- | --- | --- |
|  | P-value | # Of miR-29a/c-3p Target in DE-Genes | P-value | # Of miR-29a/c-3p Target in DE-Genes |
| TNF-Regulated Genes | 1.35E-14 | 38 | 3.36E-04 | 9 |
| TGFB1-Regulated Genes | 1.81E-13 | 36 | 7.83E-07 | 12 |
| IFNG-Regulated Genes | 2.39E-10 | 28 | 1.65E-03 | 7 |
| IL1B-Regulated Genes | 4.06E-08 | 21 | 4.10E-02 | 4 |
| NFkB-Regulated Genes | 1.06E-11 | 20 | - | - |
| MYC-Regulated Genes | 2.63E-04 | 16 | 3.25E-03 | 6 |
| AGT-Regulated Genes | 2.71E-05 | 15 | 6.55E-04 | 6 |
| IL33-Regulated Genes | 5.62E-07 | 12 | - | - |
| FOXO1-Regulated Genes | 1.36E-05 | 11 | 2.24E-03 | 4 |
| MAPK1-Regulated Genes | 1.60E-06 | 11 | 9.93E-03 | 3 |
| PDGF BB-Regulated Genes | 6.67E-07 | 11 | 7.78E-03 | 3 |
| F2-Regulated Genes | 8.76E-07 | 10 | 4.43E-02 | 2 |
| IL1-Regulated Genes | 6.07E-06 | 10 | - | - |
| CSF2-Regulated Genes | 1.86E-04 | 10 | 2.42E-02 | 3 |

\*DE: Differentially expressed. NT: Normotensive pregnancy; PE: Preeclampsia; P0: Passage 0.

### MiR-29a/c-3p differentially regulate endothelial monolayer integrity in F and M HUVECs from NT and PE

Compared with NC, miR-29a/c-3p(+) at 10nM significantly increased (> 340%) the level of both miR-29a-3p and miR-29c-3p in HUVECs 24h after transfection and this overexpression maintained for up to 72h (Fig.S1). We previously reported that miR-29a/c-3p(i) at 50nM significantly decreased the levels of both miR-29a-3p and miR-29c-3p in HUVECs after 24-72h of transfection(Zhou *et al*., 2017). Hence miR-29a/c-3p(+) at 10nM and miR-29a/c-3p(i) at 50nM were utilized in all miR-29a/c-3p overexpression and knockdown experiments in this study. Veh and NC did not alter the endothelial monolayer integrity in all cell groups treated with ECMb, TGFβ1, and TNFα (Fig.3 and Fig.S4).

**Figure 3.**
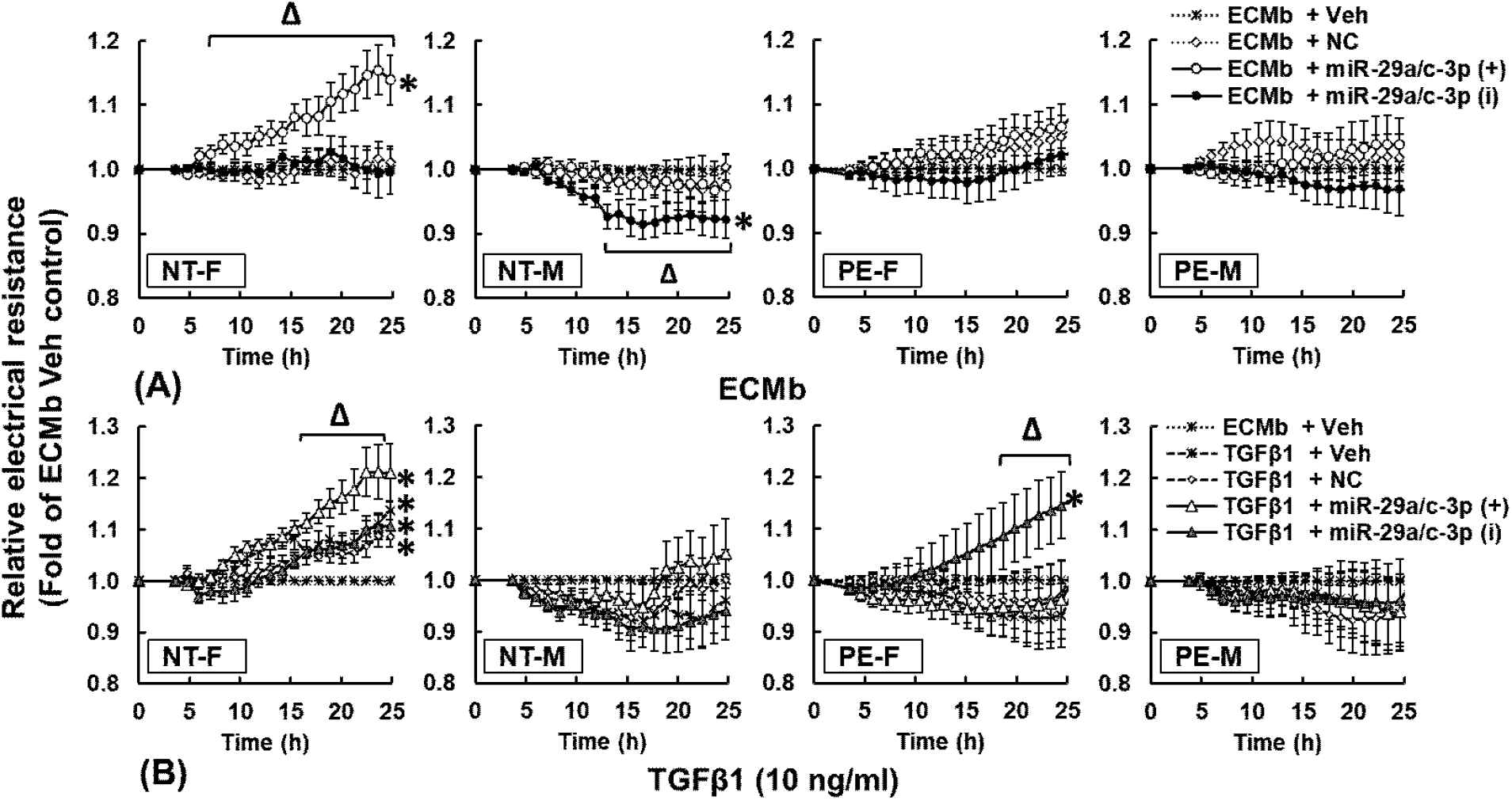
MiR-29a/c-3p differentially regulate basal endothelial monolayer integrity in F and M HUVECs from NT and PE. Cells were transfected with miR-29a/c-3p(+), miR-29a/c-3p(i), Negative Control (NC), or Vehicle (Veh), and then cultured until confluence (24-28h). After 6-8 hr of serum starvation, confluent cells were treated with ECMb (serum-free control), TGFβ1 (10 ng/ml), or TNFα (10 ng/ml) for 25h. Electrical resistance at 4000Hz was constantly recorded. Data are expressed as medians ± SEM fold of corresponding Vehicle control at the corresponding time. *Differ (*P*<0.05) from Veh control in ECMb; ^**Δ**^Differ from corresponding Veh and NC control groups. (*P*<0.05, Kruskal-Wallis test; n = 5-10 cell preparations/sex/group)

Compared with Veh and NC, miR-29a/c-3p(+) time-dependently increased electrical resistance (strengthening endothelial monolayer integrity) in NT-F HUVECs, but not in NT-M PE-F and PE-M HUVECs (Fig.3A). Specifically, in NT-F HUVECs, miR-29a/c-3p(+) strengthened endothelial monolayer integrity, beginning at 7h and reached a maximum increase of ∼15% at 24h. This miR-29a/c-3p(+)-enhanced monolayer integrity was lost in PE-F HUVECs. Compared with Veh and NC, miR-29a/c-3p(i) time-dependently decreased electrical resistance (weakening endothelial monolayer integrity) only in NT-M, but not in NT-F, PE-F, and PE-M HUVECs in ECMb (Fig.3A). Specifically, in NT-M HUVECs, miR-29a/c-3p(i) weakened endothelial monolayer integrity, beginning at 10h, reached a maximum of ∼9% decrease at 16h and maintained this level through 25h.

### MiR-29a/c-3p differentially regulate endothelial monolayer integrity in response to TGFβ1 in F and M HUVECs from NT and PE

Compared with ECMb, TGFβ1 time-dependently strengthened endothelial monolayer integrity in NT-F HUVECs, but not in NT-M, PE-F, and PE-M HUVECs. Specifically, TGFβ1 strengthened endothelial monolayer integrity in NT-F transfected with Veh and NC, starting at 15h and reaching 14% and 9% enhancement at 25h, respectively (Fig.3B). MiR-29a/c-3p(+) further enhanced the TGFβ1-strengthened endothelial monolayer integrity in NT-F HUVECs, starting at 16h and reaching 21% enhancement at 25h. The TGFβ1-strengthened endothelial monolayer integrity in F HUVECs was abolished in PE-F cells, while miR-29a/c-3p(i) recovered this TGFβ1-strengthened endothelial monolayer integrity in PE-F HUVECs (reaching 15% enhancement at 25h). MiR-29a/c-3p(i) did not alter endothelial monolayer integrity in NT-F, NT-M, and PE-M HUVECs.

Compared to ECMb, TNFα significantly decreased the endothelial monolayer integrity in NT-F, NT-M, PE-F, and PE-M HUVECs (Fig.S4). MiR-29a/c-3p(i) and miR-29a/c-3p(+) did not alter the TNFα-weakened endothelial monolayer integrity in NT-F, NT-M, PE-F, and PE-M HUVECs.

### MiR-29a/c-3p differentially regulate TNFα-induced cell proliferation in F and M HUVECs from PE

Veh and NC did not alter the cell proliferation in all cell groups treated with ECMb, TGFβ1, and TNFα. Compared with Veh and NC control, miR-29a/c-3p(+) and miR-29a/c-3p(i) did not alter the cell proliferation in response to ECMb and TGFβ1 in NT-F, NT-M, PE-F, and PE-M (Fig.4). Compared to ECMb control, TNFα promoted cell proliferation in PE-F (159%) and NT-M (153%) HUVECs (Fig.4C). Compared to Veh and NC, miR-29a/c-3p(+) further promoted the cell proliferation in PE-M HUVECs (181% of ECMb control), while MiR-29a/c-3p(i) did not affect the endothelial proliferation responses to TNFα in all HUVECs groups.

**Figure 4.**
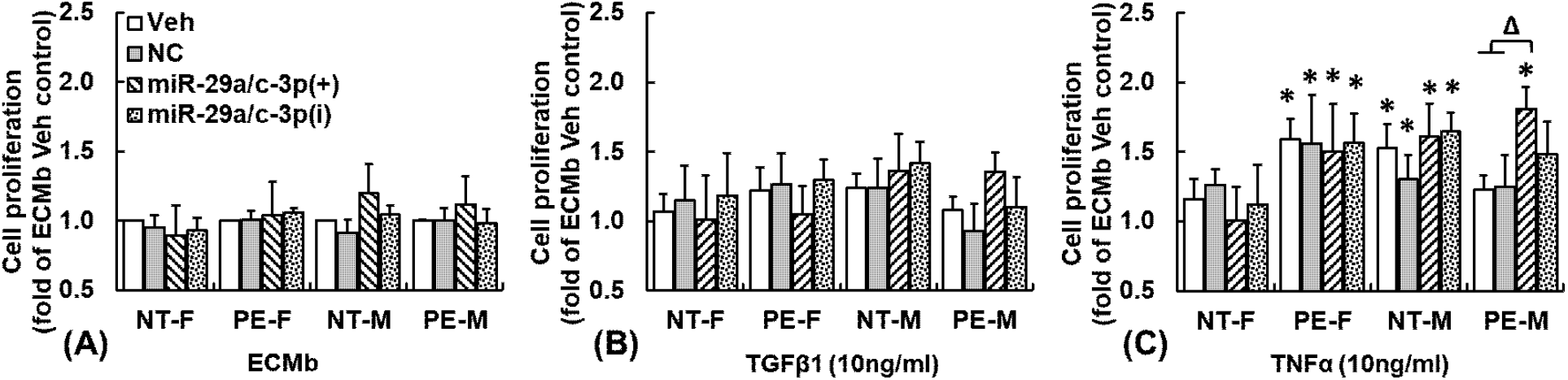
Effects of miR-29a/c-3p on cell proliferation in F and M P1-HUVECs from NT and PE. Sub-confluence cells were transfected with Veh, miR-29a/c-3p(+), or miR-29a/c-3p(i) for 24h. After 8 hr of serum starvation, sub-confluent cells were treated with ECMb [serum free control; **(A)**], TGFβ1 [10 ng/ml; **(B)**], or TNFα [10ng/ml; (**C)**] for 48h. Data are expressed as medians ± SEM fold of ECMb Vehicle control in NT-F. **\***Differ from corresponding Veh and NC controls in ECMb; ^**Δ**^Differ from corresponding NC and Veh control groups within the same treatment. (*P*<0.05, Kruskal-Wallis test; n=5-8 cell preparations/sex/group).

## Discussion

In this study, we have demonstrated for the first time that PE dysregulates miR-29a/c-3p in fetoplacental tissues and primary human fetal endothelial cells (HUVECs) in a fetal sex-specific manner. We have further shown fetal sex-specific dysregulation of cardiovascular diseases- and endothelial function-associated miR-29a/c-3p target genes in F and M fetal endothelial cells from PE. Moreover, miR-29a/c-3p overexpression and knockdown differentially affect PE-dysregulated fetal endothelial cell responses to cytokines in F and M HUVECs. These data indicate fetal sex-specific roles of miR-29a/c-3p in PE-dysregulated transcriptomic profiles and cell functions in fetal endothelial cells.

Expression of miR-29a/c-3p in human placentas are significantly higher in the third trimester than that in the first trimester(Gu *et al*., 2013), suggesting miR-29a/c-3p play important roles in placental development and function. Our current finding that PE upregulated miR-29a/c-3p levels in F, but not M fetoplacentas agrees with a previous report that miR-29a-3p is elevated in maternal plasma from mild PE patients(Li *et al*., 2013), indicating fetal sex-specific expression patterns of miR-29a/c-3p in PE placentas. To date, although we do not know the exact cell contribution to this PE-dysregulated miR-29a/c-3p expression, this fetal sex-specific upregulation of miR-29a/c-3p in PE placentas suggests that this dysregulation may contribute to the PE-impaired placental functions.

The current observation that PE only downregulates miR-29a/c-3p expression in M, but not in F HUVECs extends our previous report(Zhou *et al*., 2017; Zhou *et al*., 2019) and supports our hypothesis that PE differentially regulates miR-29a/c-3p expression in F and M HUVECs. However, this differential expression pattern in HUVECs is opposite to that in placentas, indicating different regulation of miR-29a/c-3p in placental tissues and HUVECs. We have previously reported that F HUVECs are transcriptionally more responsive to PE than their M counterparts(Zhou *et al*., 2019). In this study, we found that PE-dysregulated ∼4.5-fold of miR-29a/c-3p target genes in F HUVECs (125 genes) than in M HUVECs (28 genes). These data imply that while miR-29a/c-3p has a critical role in PE-dysregulated transcriptomes in F and M HUVECs, F HUVECs are more susceptible to miR-29a/c-3p regulation in PE. It is noteworthy that there are only five common PE-dysregulated miR-29a/c-3p target genes observed in F and M cells. However, these five DE genes had opposite regulation directions in F and M HUVECs: all were down-regulated in F but up-regulated in M HUVECs from PE. These DE genes include Activity Regulated Cytoskeleton Associated Protein (ARC) and Cartilage Intermediate Layer Protein 2 (CLIP2), both of which are important for intercellular microRNA transportation via extracellular exosome(Pastuzyn *et al*., 2018; Hu *et al*., 2020). Thus, fetal sex-specific dysregulation of these intercellular microRNA transportation-related genes may contribute to PE-dysregulated miR-29a/c-3p in HUVECs and fetoplacental tissues. This notion is consistent with our previous report that PE, in general, dysregulates more cardiovascular/endothelial function-associated pathways and biological functions in F than in M HUVECs(Zhou *et al*., 2019), as well as our observation in the current study that miR-29c-3p expression in PE-F is higher than in PE-M HUVECs.

Many of the PE-dysregulated miR-29a/c-3p target pathways and biological functions (e.g., cell movement, cell viability, angiogenesis, and concentration of Ca^2+^) in HUVECs are associated with PE-induced fetal endothelial dysfunction (e.g., reduced cell migration, and impaired calcium signaling) and vascular-related disorders (e.g., atherosclerosis, disorder of blood pressure, obesity, and seizures)(Wang *et al*.; Powe *et al*., 2011a; Boeldt *et al*.; Brodowski *et al*.; Zhou *et al*., 2019; Zhou *et al*., 2020). These data suggest that miR-29a/c-3p play important roles in vascular functions during pregnancy and PE-offspring, as PE-offspring are known to face increased risks of cardiovascular and metabolic disorders(Kajantie *et al*., 2009; Ryckman *et al*., 2013). Furthermore, PE uniquely dysregulated many endothelial function-associated miR-29a/c-3p target pathways and biological processes in F and M HUVECs. Notably, wound healing signaling, inflammatory response, and obesity were only enriched in PE-upregulated miR-29a/c-3p target genes in F HUVECs, while hypertension, stroke, and adhesion of endothelial progenitor cells were enriched only in PE-upregulated miR-29a/c-3p target genes in M HUVECs (Fig. 2D&E, Fig.S2&S3). In agreement with a previous report that dysregulated miRNA profiles in HUVECs are associated with dermal microvascular density neonates(Yu *et al*., 2018), These observations suggest that PE-dysregulated miR-29a/c-3p involve in the *in-utero* programming of fetal endothelial cells and prime PE offspring for higher risks of cardiovascular diseases later in their life.

The current finding that PE dysregulates much more inflammatory- and immune responses -associated genes (e.g., inflammatory response, adhesion of immune cells, apoptosis of antigen-presenting cells, NFkB, IL33, and IL1) in F than M HUVECs suggest a fetal sex-specific role of miR-29a/c-3p in the inflammatory response in PE. This is consistent with reports showing that the miR-29 family participates in the immunological responses after virus infections (HIV-1 and SARS-CoV-2)(Abel *et al*., 2021; Saulle *et al*., 2021).

Consistent with our bioinformatics analysis showing that PE-dysregulated miR-29a/c-3p target genes/pathways are highly associated with inflammatory responses in F and M HUVECs (Fig. 2), we observed that PE differentially regulated cell responses to inflammatory-related cytokines (TGFβ1 and TNFα) in F and M HUVECs. For instance, in agreement with our previous report(Zhou *et al*., 2019), TGFβ1 strengthens endothelial monolayer integrity only in NT F-HUVECs, which is abolished in PE, while TNFα decreases endothelial monolayer integrity in HUVECs. These data confirm the vital role of TGFβ1 and TNFα in fetal endothelial functions.

We observed that miR-29a/c-3p overexpression increased basal endothelial monolayer integrity only in NT-F, and miR-29a/c-3p knockdown decreased basal endothelial monolayer integrity only in NT-M cells, whereas this fetal sex-specific differential regulation was lost in PE-F and PE-M HUVECs. We also showed that miR-29a/c-3p overexpression further enhanced TGFβ1-strengthened endothelial monolayer integrity in F but not in M NT-HUVECs, while this fetal sex-specific regulation disappeared in PE HUVECs. Furthermore, miR-29a/c-3p knockdown recovered the TGFβ1-enhanced endothelial monolayer integrity in PE-F HUVECs but did not affect the monolayer integrity in PE-M cells. Together with our bioinformatic data that PE dysregulated more TGFβ1-regulated miR-29a/c-3p target genes in PE-F than PE-M HUVECs, these differential regulations implicate the fetal sex-specific importance of miR-29a/c-3p in maintaining basal and TGFβ1-induced endothelial monolayer integrity responses in HUVECs. Overall, PE-F HUVECs are more susceptible to miR-29a/c-3p regulated endothelial monolayer integrity responses than PE-M cells.

Overexpression and knockdown of miR-29a/c-3p do not alter the basal cell proliferation as well as cell proliferation in response to TGFβ1 and TNFα in NT HUVECs, indicating that miR-29a/c-3p is not critical to maintain the basal as well as TGFβ1- and TNFα-regulated cell proliferation responses in NT HUVECs. However, overexpression of miR-29a/c-3p recovers the TNFα-induced cell proliferation in PE-M and brings it to similar levels of NT-M cells. It appears that PE-M HUVECs are more responsive to the miR-29a/c-3p regulated cell proliferative responses than PE-F cells.

As the human umbilical vein carries oxygenated blood from the placenta to the growing fetus during pregnancy, HUVECs are a unique cell population that is directly exposed to altered humoral factors derived from placenta and maternal circulation and share many features of artery endothelial cells(Inoue *et al*., 1998; Lang *et al*., 2008; Jiang *et al*., 2013). Although a direct relationship between endothelial dysfunction observed in primary HUVECs and specific long-term cardiovascular risks in the offspring remain elusive, the emerging evidence has shown that dysregulated miRNA expression and dysfunction of HUVECs from hypertensive pregnancies including PE are associated with vascular dysfunction in offspring(Powe *et al*., 2011b; Staley *et al*., 2015; Yu *et al*., 2018).

## Conclusions

Our data have demonstrated that PE differentially dysregulates the expression of miR-29a/c-3p, miR-29a/c-3p target genes/pathways, and miR-29a/c-3p-associated endothelial cell function in response to TGFβ1 and TNFα in F and M HUVECs. These fetal sex-specific dysregulations may contribute to fetal sex-specific vascular dysfunction in PE and PE-associated adult-onset cardiovascular diseases in PE offspring.

## Perspectives

To date, there are limited therapeutic options for PE-induced fetal endothelial dysfunction due to our poor understanding of cellular and molecular mechanisms underlying PE. Here we reported fetal sexual dimorphic regulation of miR-29a/c-3p and miR-29a/c-3p target genes/pathways in HUVECs in PE. We also demonstrated fetal sex-specific dysregulation of miR-29a/c-3p-associated cellular responses to TGFβ1 and TNFα in PE HUVECs. These sexual dimorphisms of PE-dysregulated miR-29a/c-3p and their target genes/pathways may allow the discovery of novel fetal sex-specific therapeutic targets and risk predictors for adult-onset cardiovascular diseases in children born to PE mothers.

## Supporting information

Full Supplemental Materials

## Acknowledgments

We would like to thank Laura Hogan, Ph.D., a Science Editor at the University of Wisconsin Institute for Clinical and Translational Research (UW ICTR), for critically reading and editing this manuscript.

## Nonstandard abbreviations

(PE): Preeclampsia
(NT): Normotensive
(F): Female
(M): Male
(DE): Differentially expressed

