## Supplementary material for "MicroRNA-29 Differentially Mediates Preeclampsia-Dysregulated Cellular Responses to Cytokines in Female and Male Fetal Endothelial Cells": Full Supplemental Materials

[4]Current Institution: University of Nebraska Medical Center, Omaha, NE, United States.

**Short title:** MiR-29 Mediates Fetal Sex-specific Endothelial Dysfunction in PE

***Corresponding Authors:**

**Chi Zhou**, Ph.D., School of Animal and Comparative Biomedical Sciences, University of Arizona, 4101 N Campbell Ave., Tucson, AZ 85719, USA.

**Jing Zheng**, Ph.D., Department of Obstetrics and Gynecology, University of Wisconsin-Madison, PAB1 UnityPoint Health-Meriter Hospital, 202 S. Park St., Madison, WI 53715, USA

**Supplemental Materials**

Supplemental Methods

Tables S1-S7

Figures S1-S4

#### Supplemental Detailed Methods

##### Human fetoplacental tissue collection

Human fetoplacental tissues were dissected from the placenta immediately after delivery. After dissection, multiple tissue biopsies from each placenta were snap frozen in liquid nitrogen and stored at -80°C. Fetoplacental tissue biopsies from each placenta were ground to powder in liquid nitrogen followed by RNA isolation.

##### Isolation and characterization of HUVECs

HUVECs were freshly isolated from umbilical veins immediately after cesarean-section delivery from NT and PE pregnancies using an enzymatic method as described(Zhou *et al.*, 2017; Zhou *et al.*, 2019). To avoid the potential impact caused by the long-term *in vitro* culture, cells were purified using Dynabeads® CD31 Endothelial Cell (Life Technologies, Carlsbad, CA) after 16 h of culture under 37°C and 5% $CO_2$ in Endothelial Cell Medium (ECM; ScienCell Research Laboratories, Carlsbad, CA), which consisted of ECM basal medium (ECM-b) supplemented with 5% fetal bovine serum, 1% endothelial cell growth supplement, and 1% penicillin/streptomycin solution.

##### DiI-Ac-LDL uptake assay

The purity of HUVECs was examined as previously described(Zhou *et al.*, 2017; Zhou *et al.*, 2019). Cells were incubated with Acetylated Low-Density Lipoprotein labeled with 1,1'-dioctadecyl-3,3,3',3'-tetramethyl-indocarbocyanine perchlorate (DiI-Ac-LDL, 10μg/mL, Invitrogen, Waltham, MA) for 4 h at 37°C, fixed in 4% formaldehyde, and examined under a phase-contrast and fluorescent microscope. Images were taken utilizing a Biotek Cytation5 multimode imager with the Gen5 software. Cells exhibiting uptake of LDL were counted. Cells incubated without DiI-Ac-LDL were served as negative controls.

##### Real-time qPCR (RT-qPCR) analysis

To determine the expression of miRNAs of interest in human fetoplacental tissues and P0 HUVECs, 400ng small RNA fragment enriched total RNA isolated from each sample was reverse transcribed into cDNA using a miScript II RT Kit (Qiagen, CA). Diluted cDNA corresponding to 2 ng of original total RNA was utilized as the template in each RT-qPCR reaction. Each miRNA of interest was performed using commercially available miRNA miScript Primer Assays in Supplemental Table S1 (Qiagen), miScript SYBR Green PCR Kit (Qiagen), and StepOne$^{Plus}$

qPCR system (Life Technologies). The efficiencies of all target and control miRNA assays were between 90% and 110%. The RT-qPCR data were normalized to the external control (miRTC, Qiagen) along with internal control miRNAs (SNORD95, and SNORD96A). The normalized RT-qPCR data were then further analyzed using the $2^{-\Delta\Delta CT}$ method(Yuan *et al.*, 2006; Zhou *et al.*, 2017) to determine the relative abundance.

**Overexpression and Knockdown of miRNAs**

Due to the limited amount of P0 HUVECs (usually only $3\times10^5$-$1\times10^6$ per preparation), we used passages 1 (P1, ~5 days of culture) HUVECs preparations(Zhou *et al.*, 2019) in the miRNA overexpression and knockdown study. Overexpression and knockdown of each target miRNA were performed using miScript miRNA Mimic [Qiagen, refer as miRNA(+)] and miScript miRNA Inhibitor [Qiagen, refer as miRNA-(i)] which is chemically synthesized and modified single-strand RNA that specifically overexpress and inhibit target miRNA(Ukai *et al.*, 2012; Zhou *et al.*, 2017).

To overexpress miR-29a-3p and miR-29c-3p in HUVECs, P1 HUVECs at 50-60% confluence were transfected with miScript miRNA mimics (Qiagen) that specifically target human miR-29a-3p (Cat.# MSY0000086), miR-29c-3p (Cat.# MSY0000681), and miScript Inhibitor Negative Control (Cat.# 1027272) at 0, 5, or 10 nM using the HiPerFect Transfection Reagent (Qiagen) for 24h, 48h, and 72h. Cells transfected with the miScript Inhibitor Negative Control were used as negative controls (NC). Cells treated with only the HiPerFect Transfection Reagent were used as the vehicle (Veh) controls. RT-qPCR (see above) was used to verify the efficiency of miRNA knockdown. MiScript miR-29a-3p mimic (Qiagen, Cat.# MSY0000086) at 10nM significantly overexpressed (> 340%) overexpression of both miR-29a-3p and miR-29a-3p in HUVECs at 24h after transfection and this overexpression last through 72h after transfection. In this study, miScript miR-29a-3p mimics [Qiagen, Cat.# MSY0000086] at 10nM was used to overexpress both miR-29a-3p and miR-29a-3p in HUVECs [refer as miR-29a/c-3p(+)].

Knockdown of miR-29a-3p and miR-29c-3p in HUVECs were performed as we previously described(Zhou *et al.*, 2017). P1 HUVECs at 50-60% confluence were transfected with miScript miRNA Inhibitors (Qiagen) that specifically target human miR-29a-3p (Cat.# MIN0000086), miR-29c-3p (Cat.# MIN0000681), and miScript Inhibitor Negative Control (Cat.# 1027272) at 0, 50, or 100 nM using the HiPerFect Transfection Reagent (Qiagen) for 24h, 48h, and 72h. Cells transfected with the miScript Inhibitor Negative Control were used as negative controls (NC). Cells treated with only the HiPerFect Transfection Reagent were used as the vehicle (Veh) control. RT-qPCR (see above) was used to verify the efficiency of miRNA knockdown. As we previously reported, miScript miR-29c-3p inhibitor (Qiagen, Cat.# MIN0000681) at 50nM and 100nM significantly knockdown (> 90%) both miR-29a-3p and miR-29a-3p in HUVECs at 24h after transfection and this knockdown last through 72h after transfection(Zhou *et al.*, 2017). In this study, miScript miR-29c-3p inhibitor (Qiagen, Cat.# MIN0000681) at 50nM was used for knockdown both miR-29a-3p and miR-29c-3p in HUVECs [refer as miR-29a/c-3p(i)].

**Cell proliferation assay**

Cell proliferation was assessed using the CCK-8 kit (Dojindo Molecular Technologies, Rockville, MD) as described(Zhou *et al.*, 2019). Cells seeded in 96-well plates (5000 cells/well) were transfected with Veh, NC, miRNA-(+), and miRNA-(i) for 24h to overexpress or knockdown target miRNAs of interest. After miRNAs overexpression and knockdown, the cells were serum-starved in ECMb serum-free control media for 8h and then treated with ECM-b (serum-free control), TGFβ (10 ng/ml), and TNFα (10 ng/ml) for 48h (4 wells per treatment for each cell preparation). Cells were then incubated with the CCK-8 reagent for 1h, and absorbance at 450nm was measured using a Biotek Cytation5 multimode plate reader (Agilent, Winooski, VT).

**Cell monolayer integrity assay**

Cell monolayer integrity of HUVECs was determined using the ECIS Zθ+ 96-well array station (Applied BioPhysics, Troy, NY) using 96-well plates (96W20idf) as we previously described(Zhou *et al.*, 2019). Cells seeded at $2 \times 10^4$ per well were treated with were transfected with Veh, NC, miRNA-(+), and miRNA-(i) for 24h to overexpress or knockdown target miRNAs of interest. The cells were cultured until 100% confluence (~30h after seeding). The confluent cells were then treated with ECM-b (serum-free control), TGFβ (10 ng/ml), and TNFα (10 ng/ml) for 25h (4 wells per treatment for each cell preparation). ECIS system was used to real-time measure the electrical resistance over the monolayer of endothelial cells (indicating the monolayer integrity). Electrical resistance was constantly monitored and recorded up to 25 h.

**Table S1. List of miRNA Primer Assays used in miRNA RT-qPCR.**

| Mature miRNA ID | RT-qPCR Primer Assay type | Vendor | Catalog Number |
|---|---|---|---|
| Hsa-miR-29a-3p | Primer Assay for miRNA of interest | Qiagen | MS00003262 |
| Hsa-miR-29b-3p | Primer Assay for miRNA of interest | Qiagen | MS00006566 |
| Hsa-miR-29c-3p | Primer Assay for miRNA of interest | Qiagen | MS00003269 |
| MiRTC | Primer Assay for external control | Qiagen | MS00000001 |
| Hs_SNORD95 | Primer Assay for endogenous control (small nucleolar RNA) | Qiagen | MS00033726 |
| Hs_SNORD96A | Primer Assay for endogenous control (small nucleolar RNA) | Qiagen | MS00033733 |

Table S2. MiR-29a/c-3p target genes detected in P0-HUVECs.

| GeneID | gene | Human miR-29a/c-3p target | Database |
|---|---|---|---|
| ENSG00000002746 | HECW1 | miR-29a/c-3p common target gene | Theoretical prediction (microT-CDS algorithm) |
| ENSG00000002834 | LASP1 | miR-29a/c-3p common target gene | Experiment supported target gene (TarBase v.8) |
| ENSG00000002919 | SNX11 | miR-29a/c-3p common target gene | Experiment supported target gene (TarBase v.8) |
| ENSG00000003393 | ALS2 | miR-29a/c-3p common target gene | Theoretical prediction (microT-CDS algorithm) |
| ENSG00000003989 | SLC7A2 | miR-29a-3p target gene only | Experiment supported target gene (TarBase v.8) |
| ENSG00000004059 | ARF5 | miR-29a/c-3p common target gene | Theoretical prediction (microT-CDS algorithm) |
| ENSG00000004142 | POLDIP2 | miR-29a-3p target gene only | Experiment supported target gene (TarBase v.8) |
| ENSG00000004487 | KDM1A | miR-29a/c-3p common target gene | Experiment supported target gene (TarBase v.8) |
| ENSG00000004799 | PDK4 | miR-29a-3p target gene only | Experiment supported target gene (TarBase v.8) |
| ENSG00000004864 | SLC25A13 | miR-29a/c-3p common target gene | Experiment supported target gene (TarBase v.8) |
| ENSG00000004948 | CALCR | miR-29a/c-3p common target gene | Theoretical prediction (microT-CDS algorithm) |
| ENSG00000005007 | UPF1 | miR-29a-3p target gene only | Experiment supported target gene (TarBase v.8) |
| ENSG00000005022 | SLC25A5 | miR-29a/c-3p common target gene | Experiment supported target gene (TarBase v.8) |
| ENSG00000005100 | DHX33 | miR-29a-3p target gene only | Experiment supported target gene (TarBase v.8) |
| ENSG00000005302 | MSL3 | miR-29a/c-3p common target gene | Experiment supported target gene (TarBase v.8) |
| ENSG00000005700 | IBTK | miR-29a/c-3p common target gene | Experiment supported target gene (TarBase v.8) |
| ENSG00000005810 | MYCBP2 | miR-29a-3p target gene only | Experiment supported target gene (TarBase v.8) |
| ENSG00000005889 | ZFX | miR-29a/c-3p common target gene | Theoretical prediction (microT-CDS algorithm) |
| ENSG00000005893 | LAMP2 | miR-29a/c-3p common target gene | Experiment supported target gene (TarBase v.8) |
| ENSG00000006042 | TMEM98 | miR-29c-3p target gene only | Experiment supported target gene (TarBase v.8) |
| ENSG00000006047 | YBX2 | miR-29c-3p target gene only | Experiment supported target gene (TarBase v.8) |
| ENSG00000006125 | AP2B1 | miR-29a/c-3p common target gene | Experiment supported target gene (TarBase v.8) |
| ENSG00000006194 | ZNF263 | miR-29a/c-3p common target gene | Theoretical prediction (microT-CDS algorithm) |
| ENSG00000006210 | CX3CL1 | miR-29a/c-3p common target gene | Theoretical prediction (microT-CDS algorithm) |
| ENSG00000006451 | RALA | miR-29a/c-3p common target gene | Experiment supported target gene (TarBase v.8) |
| ENSG00000006607 | FARP2 | miR-29a/c-3p common target gene | Experiment supported target gene (TarBase v.8) |
| ENSG00000006625 | GGCT | miR-29c-3p target gene only | Experiment supported target gene (TarBase v.8) |
| ENSG00000006757 | PNPLA4 | miR-29c-3p target gene only | Experiment supported target gene (TarBase v.8) |
| ENSG00000006831 | ADIPOR2 | miR-29a-3p target gene only | Experiment supported target gene (TarBase v.8) |
| ENSG00000007168 | PAFAH1B1 | miR-29a-3p target gene only | Experiment supported target gene (TarBase v.8) |
| ENSG00000007237 | GAS7 | miR-29c-3p target gene only | Experiment supported target gene (TarBase v.8) |
| ENSG00000007384 | RHBDF1 | miR-29a-3p target gene only | Experiment supported target gene (TarBase v.8) |
| ENSG00000007923 | DNAJC11 | miR-29a-3p target gene only | Experiment supported target gene (TarBase v.8) |
| ENSG00000008083 | JARID2 | miR-29a/c-3p common target gene | Experiment supported target gene (TarBase v.8) |
| ENSG00000008086 | CDKL5 | miR-29c-3p target gene only | Experiment supported target gene (TarBase v.8) |
| ENSG00000008277 | ADAM22 | miR-29a/c-3p common target gene | Experiment supported target gene (TarBase v.8) |
| ENSG00000008294 | SPAG9 | miR-29a-3p target gene only | Experiment supported target gene (TarBase v.8) |
| ENSG00000008988 | RPS20 | miR-29c-3p target gene only | Experiment supported target gene (TarBase v.8) |
| ENSG00000009307 | CSDE1 | miR-29a/c-3p common target gene | Experiment supported target gene (TarBase v.8) |
| ENSG00000009413 | REV3L | miR-29a/c-3p common target gene | Experiment supported target gene (TarBase v.8) |
| ENSG00000009694 | TENM1 | miR-29a-3p target gene only | Experiment supported target gene (TarBase v.8) |
| ENSG00000009844 | VTA1 | miR-29a/c-3p common target gene | Experiment supported target gene (TarBase v.8) |
| ENSG00000010030 | ETV7 | miR-29c-3p target gene only | Experiment supported target gene (TarBase v.8) |
| ENSG00000010072 | SPRTN | miR-29a/c-3p common target gene | Experiment supported target gene (TarBase v.8) |
| ENSG00000010244 | ZNF207 | miR-29a/c-3p common target gene | Theoretical prediction (microT-CDS algorithm) |
| ENSG00000010256 | UQCRC1 | miR-29a-3p target gene only | Experiment supported target gene (TarBase v.8) |
| ENSG00000010278 | CD9 | miR-29a-3p target gene only | Experiment supported target gene (TarBase v.8) |
| ENSG00000010292 | NCAPD2 | miR-29a-3p target gene only | Experiment supported target gene (TarBase v.8) |
| ENSG00000010295 | IFFO1 | miR-29a/c-3p common target gene | Theoretical prediction (microT-CDS algorithm) |
| ENSG00000010404 | IDS | miR-29a/c-3p common target gene | Theoretical prediction (microT-CDS algorithm) |
| ENSG00000010438 | PRSS3 | miR-29c-3p target gene only | Experiment supported target gene (TarBase v.8) |
| ENSG00000010539 | ZNF200 | miR-29a-3p target gene only | Experiment supported target gene (TarBase v.8) |
| ENSG00000010810 | FYN | miR-29a/c-3p common target gene | Theoretical prediction (microT-CDS algorithm) |
| ENSG00000010932 | FMO1 | miR-29a/c-3p common target gene | Theoretical prediction (microT-CDS algorithm) |
| ENSG00000011105 | TSPAN9 | miR-29a/c-3p common target gene | Theoretical prediction (microT-CDS algorithm) |
| ENSG00000011114 | BTBD7 | miR-29a/c-3p common target gene | Experiment supported target gene (TarBase v.8) |
| ENSG00000011258 | MBTD1 | miR-29a/c-3p common target gene | Experiment supported target gene (TarBase v.8) |
| ENSG00000011347 | SYT7 | miR-29a/c-3p common target gene | Experiment supported target gene (TarBase v.8) |
| ENSG00000011422 | PLAUR | miR-29c-3p target gene only | Experiment supported target gene (TarBase v.8) |
| ENSG00000011426 | ANLN | miR-29a-3p target gene only | Experiment supported target gene (TarBase v.8) |
| ENSG00000011454 | RABGAP1 | miR-29a/c-3p common target gene | Experiment supported target gene (TarBase v.8) |
| ENSG00000011523 | CEP68 | miR-29a/c-3p common target gene | Theoretical prediction (microT-CDS algorithm) |
| ENSG00000011638 | TMEM159 | miR-29a/c-3p common target gene | Experiment supported target gene (TarBase v.8) |
| ENSG00000012174 | MBTPS2 | miR-29c-3p target gene only | Experiment supported target gene (TarBase v.8) |
| ENSG00000012211 | PRICKLE3 | miR-29c-3p target gene only | Experiment supported target gene (TarBase v.8) |
| ENSG00000012232 | EXTL3 | miR-29a-3p target gene only | Experiment supported target gene (TarBase v.8) |
| ENSG00000013016 | EHD3 | miR-29a/c-3p common target gene | Experiment supported target gene (TarBase v.8) |
| ENSG00000013297 | CLDN11 | miR-29a/c-3p common target gene | Experiment supported target gene (TarBase v.8) |
| ENSG00000013374 | NUB1 | miR-29a/c-3p common target gene | Theoretical prediction (microT-CDS algorithm) |
| ENSG00000013503 | POLR3B | miR-29a/c-3p common target gene | Experiment supported target gene (TarBase v.8) |
| ENSG00000013563 | DNASE1L1 | miR-29a/c-3p common target gene | Experiment supported target gene (TarBase v.8) |
| ENSG00000014919 | COX15 | miR-29a/c-3p common target gene | Experiment supported target gene (TarBase v.8) |
| ENSG00000015133 | CCDC88C | miR-29a/c-3p common target gene | Experiment supported target gene (TarBase v.8) |
| ENSG00000016082 | ISL1 | miR-29a/c-3p common target gene | Theoretical prediction (microT-CDS algorithm) |
| ENSG00000016391 | CHDH | miR-29a/c-3p common target gene | Experiment supported target gene (TarBase v.8) |
| ENSG00000016402 | IL20RA | miR-29c-3p target gene only | Experiment supported target gene (TarBase v.8) |
| ENSG00000017427 | IGF1 | miR-29a/c-3p common target gene | Experiment supported target gene (TarBase v.8) |
| ENSG00000017483 | SLC38A5 | miR-29a-3p target gene only | Experiment supported target gene (TarBase v.8) |
| ENSG00000018189 | RUFY3 | miR-29a-3p target gene only | Experiment supported target gene (TarBase v.8) |
| ENSG00000018236 | CNTN1 | miR-29a/c-3p common target gene | Experiment supported target gene (TarBase v.8) |
| ENSG00000018408 | WWTR1 | miR-29a/c-3p common target gene | Experiment supported target gene (TarBase v.8) |
| ENSG00000019144 | PHLDB1 | miR-29a/c-3p common target gene | Experiment supported target gene (TarBase v.8) |
| ENSG00000019485 | PRDM11 | miR-29a-3p target gene only | Experiment supported target gene (TarBase v.8) |
| ENSG00000019995 | ZRANB1 | miR-29a/c-3p common target gene | Experiment supported target gene (TarBase v.8) |
| ENSG00000021826 | CPS1 | miR-29a/c-3p common target gene | Experiment supported target gene (TarBase v.8) |
| ENSG00000023171 | GRAMD1B | miR-29a/c-3p common target gene | Theoretical prediction (microT-CDS algorithm) |
| ENSG00000023287 | RB1CC1 | miR-29a/c-3p common target gene | Experiment supported target gene (TarBase v.8) |
| ENSG00000023318 | ERP44 | miR-29a/c-3p common target gene | Theoretical prediction (microT-CDS algorithm) |
| ENSG00000023516 | AKAP11 | miR-29a/c-3p common target gene | Experiment supported target gene (TarBase v.8) |
| ENSG00000024422 | EHD2 | miR-29a/c-3p common target gene | Experiment supported target gene (TarBase v.8) |
| ENSG00000025423 | HSD17B6 | miR-29a-3p target gene only | Experiment supported target gene (TarBase v.8) |
| ENSG00000026103 | FAS | miR-29c-3p target gene only | Experiment supported target gene (TarBase v.8) |
| ENSG00000026652 | AGPAT4 | miR-29c-3p target gene only | Experiment supported target gene (TarBase v.8) |
| ENSG00000027001 | MIPEP | miR-29a/c-3p common target gene | Experiment supported target gene (TarBase v.8) |
| ENSG00000028277 | POU2F2 | miR-29a-3p target gene only | Experiment supported target gene (TarBase v.8) |
| ENSG00000028528 | SNX1 | miR-29c-3p target gene only | Experiment supported target gene (TarBase v.8) |
| ENSG00000029153 | ARNTL2 | miR-29a/c-3p common target gene | Theoretical prediction (microT-CDS algorithm) |
| ENSG00000029364 | SLC39A9 | miR-29a/c-3p common target gene | Experiment supported target gene (TarBase v.8) |
| ENSG00000029725 | RABEP1 | miR-29a/c-3p common target gene | Theoretical prediction (microT-CDS algorithm) |
| ENSG00000029993 | HMGB3 | miR-29c-3p target gene only | Experiment supported target gene (TarBase v.8) |
| ENSG00000030066 | NUP160 | miR-29a/c-3p common target gene | Experiment supported target gene (TarBase v.8) |
| ENSG00000030110 | BAK1 | miR-29a/c-3p common target gene | Experiment supported target gene (TarBase v.8) |
| ENSG00000031003 | FAM13B | miR-29a/c-3p common target gene | Experiment supported target gene (TarBase v.8) |
| ENSG00000031698 | SARS | miR-29a/c-3p common target gene | Experiment supported target gene (TarBase v.8) |
| ENSG00000033170 | FUT8 | miR-29a/c-3p common target gene | Experiment supported target gene (TarBase v.8) |
| ENSG00000033867 | SLC4A7 | miR-29a/c-3p common target gene | Experiment supported target gene (TarBase v.8) |
| ENSG00000034533 | ASTE1 | miR-29a/c-3p common target gene | Experiment supported target gene (TarBase v.8) |
| ENSG00000034677 | RNF19A | miR-29a/c-3p common target gene | Experiment supported target gene (TarBase v.8) |
| ENSG00000035141 | FAM136A | miR-29a/c-3p common target gene | Experiment supported target gene (TarBase v.8) |
| ENSG00000035403 | VCL | miR-29a/c-3p common target gene | Experiment supported target gene (TarBase v.8) |
| ENSG00000035499 | DEPDC1B | miR-29c-3p target gene only | Experiment supported target gene (TarBase v.8) |
| ENSG00000035687 | ADSS | miR-29a/c-3p common target gene | Experiment supported target gene (TarBase v.8) |
| ENSG00000035862 | TIMP2 | miR-29a/c-3p common target gene | Experiment supported target gene (TarBase v.8) |
| ENSG00000037749 | MFAP3 | miR-29a/c-3p common target gene | Experiment supported target gene (TarBase v.8) |
| ENSG00000038219 | BOD1L1 | miR-29a/c-3p common target gene | Experiment supported target gene (TarBase v.8) |
| ENSG00000038295 | TLL1 | miR-29a/c-3p common target gene | Theoretical prediction (microT-CDS algorithm) |
| ENSG00000038427 | VCAN | miR-29a/c-3p common target gene | Experiment supported target gene (TarBase v.8) |
| ENSG00000039068 | CDH1 | miR-29c-3p target gene only | Experiment supported target gene (TarBase v.8) |
| ENSG00000040199 | PHLPP2 | miR-29a/c-3p common target gene | Experiment supported target gene (TarBase v.8) |
| ENSG00000040275 | SPDL1 | miR-29a/c-3p common target gene | Experiment supported target gene (TarBase v.8) |
| ENSG00000041353 | RAB27B | miR-29a-3p target gene only | Experiment supported target gene (TarBase v.8) |
| ENSG00000041357 | PSMA4 | miR-29c-3p target gene only | Experiment supported target gene (TarBase v.8) |
| ENSG00000042062 | FAM65C | miR-29a-3p target gene only | Experiment supported target gene (TarBase v.8) |
| ENSG00000042286 | AIFM2 | miR-29a/c-3p common target gene | Theoretical prediction (microT-CDS algorithm) |
| ENSG00000042753 | AP2S1 | miR-29c-3p target gene only | Experiment supported target gene (TarBase v.8) |
| ENSG00000042781 | USH2A | miR-29c-3p target gene only | Experiment supported target gene (TarBase v.8) |
| ENSG00000044574 | HSPA5 | miR-29a-3p target gene only | Experiment supported target gene (TarBase v.8) |
| ENSG00000047346 | FAM214A | miR-29a/c-3p common target gene | Experiment supported target gene (TarBase v.8) |
| ENSG00000047597 | XK | miR-29c-3p target gene only | Experiment supported target gene (TarBase v.8) |
| ENSG00000047621 | C12orf4 | miR-29a/c-3p common target gene | Experiment supported target gene (TarBase v.8) |
| ENSG00000047634 | SCML1 | miR-29a/c-3p common target gene | Experiment supported target gene (TarBase v.8) |
| ENSG00000047849 | MAP4 | miR-29a/c-3p common target gene | Experiment supported target gene (TarBase v.8) |
| ENSG00000048028 | USP28 | miR-29a/c-3p common target gene | Theoretical prediction (microT-CDS algorithm) |
| ENSG00000048405 | ZNF800 | miR-29a-3p target gene only | Experiment supported target gene (TarBase v.8) |
| ENSG00000048707 | VPS13D | miR-29a/c-3p common target gene | Experiment supported target gene (TarBase v.8) |
| ENSG00000048740 | CELF2 | miR-29c-3p target gene only | Experiment supported target gene (TarBase v.8) |
| ENSG00000048828 | FAM120A | miR-29c-3p target gene only | Experiment supported target gene (TarBase v.8) |
| ENSG00000049192 | ADAMTS6 | miR-29a/c-3p common target gene | Theoretical prediction (microT-CDS algorithm) |
| ENSG00000049249 | TNFRSF9 | miR-29a/c-3p common target gene | Theoretical prediction (microT-CDS algorithm) |
| ENSG00000049540 | ELN | miR-29a/c-3p common target gene | Experiment supported target gene (TarBase v.8) |
| ENSG00000049618 | ARID1B | miR-29a/c-3p common target gene | Experiment supported target gene (TarBase v.8) |
| ENSG00000049656 | CLPTM1L | miR-29c-3p target gene only | Experiment supported target gene (TarBase v.8) |
| ENSG00000050030 | KIAA2022 | miR-29a/c-3p common target gene | Experiment supported target gene (TarBase v.8) |
| ENSG00000050405 | LIMA1 | miR-29a/c-3p common target gene | Experiment supported target gene (TarBase v.8) |
| ENSG00000051180 | RAD51 | miR-29c-3p target gene only | Experiment supported target gene (TarBase v.8) |
| ENSG00000051382 | PIK3CB | miR-29a/c-3p common target gene | Theoretical prediction (microT-CDS algorithm) |
| ENSG00000051825 | MPHOSPH9 | miR-29a-3p target gene only | Experiment supported target gene (TarBase v.8) |
| ENSG00000052723 | SIKE1 | miR-29a/c-3p common target gene | Experiment supported target gene (TarBase v.8) |
| ENSG00000053900 | ANAPC4 | miR-29a-3p target gene only | Experiment supported target gene (TarBase v.8) |
| ENSG00000054267 | ARID4B | miR-29a-3p target gene only | Experiment supported target gene (TarBase v.8) |
| ENSG00000055044 | NOP58 | miR-29a-3p target gene only | Experiment supported target gene (TarBase v.8) |
| ENSG00000055163 | CYFIP2 | miR-29c-3p target gene only | Experiment supported target gene (TarBase v.8) |
| ENSG00000057657 | PRDM1 | miR-29a-3p target gene only | Experiment supported target gene (TarBase v.8) |
| ENSG00000058063 | ATP11B | miR-29a/c-3p common target gene | Experiment supported target gene (TarBase v.8) |
| ENSG00000058085 | LAMC2 | miR-29a/c-3p common target gene | Experiment supported target gene (TarBase v.8) |
| ENSG00000058262 | SEC61A1 | miR-29a-3p target gene only | Experiment supported target gene (TarBase v.8) |
| ENSG00000058668 | ATP2B4 | miR-29a/c-3p common target gene | Experiment supported target gene (TarBase v.8) |
| ENSG00000058673 | ZC3H11A | miR-29a-3p target gene only | Experiment supported target gene (TarBase v.8) |
| ENSG00000058799 | YIPF1 | miR-29c-3p target gene only | Experiment supported target gene (TarBase v.8) |
| ENSG00000059378 | PARP12 | miR-29a/c-3p common target gene | Experiment supported target gene (TarBase v.8) |
| ENSG00000059728 | MXD1 | miR-29a/c-3p common target gene | Theoretical prediction (microT-CDS algorithm) |
| ENSG00000059804 | SLC2A3 | miR-29a/c-3p common target gene | Experiment supported target gene (TarBase v.8) |
| ENSG00000060138 | YBX3 | miR-29a/c-3p common target gene | Experiment supported target gene (TarBase v.8) |
| ENSG00000060237 | WNK1 | miR-29a-3p target gene only | Experiment supported target gene (TarBase v.8) |
| ENSG00000060642 | PIGV | miR-29a-3p target gene only | Experiment supported target gene (TarBase v.8) |
| ENSG00000060718 | COL11A1 | miR-29a/c-3p common target gene | Theoretical prediction (microT-CDS algorithm) |
| ENSG00000060749 | QSER1 | miR-29a/c-3p common target gene | Experiment supported target gene (TarBase v.8) |
| ENSG00000061273 | HDAC7 | miR-29a-3p target gene only | Experiment supported target gene (TarBase v.8) |
| ENSG00000062716 | VMP1 | miR-29a-3p target gene only | Experiment supported target gene (TarBase v.8) |
| ENSG00000063244 | U2AF2 | miR-29a-3p target gene only | Experiment supported target gene (TarBase v.8) |
| ENSG00000064012 | CASP8 | miR-29a/c-3p common target gene | Theoretical prediction (microT-CDS algorithm) |
| ENSG00000064199 | SPA17 | miR-29c-3p target gene only | Experiment supported target gene (TarBase v.8) |
| ENSG00000064225 | ST3GAL6 | miR-29a/c-3p common target gene | Theoretical prediction (microT-CDS algorithm) |
| ENSG00000064309 | CDON | miR-29a-3p target gene only | Experiment supported target gene (TarBase v.8) |
| ENSG00000064652 | SNX24 | miR-29a/c-3p common target gene | Theoretical prediction (microT-CDS algorithm) |
| ENSG00000064666 | CNN2 | miR-29a-3p target gene only | Experiment supported target gene (TarBase v.8) |
| ENSG00000064726 | BTBD1 | miR-29c-3p target gene only | Experiment supported target gene (TarBase v.8) |
| ENSG00000064995 | TAF11 | miR-29a/c-3p common target gene | Experiment supported target gene (TarBase v.8) |
| ENSG00000065268 | WDR18 | miR-29c-3p target gene only | Experiment supported target gene (TarBase v.8) |
| ENSG00000065308 | TRAM2 | miR-29a/c-3p common target gene | Experiment supported target gene (TarBase v.8) |
| ENSG00000065526 | SPEN | miR-29a/c-3p common target gene | Experiment supported target gene (TarBase v.8) |
| ENSG00000065559 | MAP2K4 | miR-29a/c-3p common target gene | Experiment supported target gene (TarBase v.8) |
| ENSG00000065609 | SNAP91 | miR-29a/c-3p common target gene | Experiment supported target gene (TarBase v.8) |
| ENSG00000065613 | SLK | miR-29a/c-3p common target gene | Theoretical prediction (microT-CDS algorithm) |
| ENSG00000065621 | GSTO2 | miR-29a/c-3p common target gene | Theoretical prediction (microT-CDS algorithm) |
| ENSG00000065809 | FAM107B | miR-29c-3p target gene only | Experiment supported target gene (TarBase v.8) |
| ENSG00000065883 | CDK13 | miR-29a-3p target gene only | Experiment supported target gene (TarBase v.8) |
| ENSG00000065911 | MTHFD2 | miR-29a/c-3p common target gene | Experiment supported target gene (TarBase v.8) |
| ENSG00000065970 | FOXJ2 | miR-29a/c-3p common target gene | Experiment supported target gene (TarBase v.8) |
| ENSG00000066084 | DIP2B | miR-29a/c-3p common target gene | Theoretical prediction (microT-CDS algorithm) |
| ENSG00000066135 | KDM4A | miR-29a-3p target gene only | Experiment supported target gene (TarBase v.8) |
| ENSG00000066427 | ATXN3 | miR-29a-3p target gene only | Experiment supported target gene (TarBase v.8) |
| ENSG00000066583 | ISOC1 | miR-29a/c-3p common target gene | Experiment supported target gene (TarBase v.8) |
| ENSG00000066697 | MSANTD3 | miR-29c-3p target gene only | Experiment supported target gene (TarBase v.8) |
| ENSG00000066735 | KIF26A | miR-29a/c-3p common target gene | Experiment supported target gene (TarBase v.8) |
| ENSG00000067048 | DDX3Y | miR-29a/c-3p common target gene | Experiment supported target gene (TarBase v.8) |
| ENSG00000067066 | SP100 | miR-29a/c-3p common target gene | Theoretical prediction (microT-CDS algorithm) |
| ENSG00000067167 | TRAM1 | miR-29a-3p target gene only | Experiment supported target gene (TarBase v.8) |
| ENSG00000067182 | TNFRSF1A | miR-29a/c-3p common target gene | Experiment supported target gene (TarBase v.8) |
| ENSG00000067369 | TP53BP1 | miR-29a/c-3p common target gene | Experiment supported target gene (TarBase v.8) |
| ENSG00000067445 | TRO | miR-29a/c-3p common target gene | Experiment supported target gene (TarBase v.8) |
| ENSG00000067596 | DHX8 | miR-29c-3p target gene only | Experiment supported target gene (TarBase v.8) |
| ENSG00000067704 | IARS2 | miR-29a/c-3p common target gene | Experiment supported target gene (TarBase v.8) |
| ENSG00000067798 | NAV3 | miR-29a/c-3p common target gene | Experiment supported target gene (TarBase v.8) |
| ENSG00000068024 | HDAC4 | miR-29a/c-3p common target gene | Experiment supported target gene (TarBase v.8) |
| ENSG00000068079 | IFI35 | miR-29c-3p target gene only | Experiment supported target gene (TarBase v.8) |
| ENSG00000068120 | COASY | miR-29a-3p target gene only | Experiment supported target gene (TarBase v.8) |
| ENSG00000068305 | MEF2A | miR-29a-3p target gene only | Experiment supported target gene (TarBase v.8) |
| ENSG00000068878 | PSME4 | miR-29a/c-3p common target gene | Experiment supported target gene (TarBase v.8) |
| ENSG00000069275 | NUCKS1 | miR-29a/c-3p common target gene | Experiment supported target gene (TarBase v.8) |
| ENSG00000069399 | BCL3 | miR-29c-3p target gene only | Experiment supported target gene (TarBase v.8) |
| ENSG00000069493 | CLEC2D | miR-29a/c-3p common target gene | Experiment supported target gene (TarBase v.8) |
| ENSG00000069667 | RORA | miR-29a/c-3p common target gene | Experiment supported target gene (TarBase v.8) |
| ENSG00000069702 | TGFBR3 | miR-29c-3p target gene only | Experiment supported target gene (TarBase v.8) |
| ENSG00000069849 | ATP1B3 | miR-29a/c-3p common target gene | Experiment supported target gene (TarBase v.8) |
| ENSG00000069956 | MAPK6 | miR-29a/c-3p common target gene | Experiment supported target gene (TarBase v.8) |
| ENSG00000070018 | LRP6 | miR-29a/c-3p common target gene | Experiment supported target gene (TarBase v.8) |
| ENSG00000070495 | JMJD6 | miR-29c-3p target gene only | Experiment supported target gene (TarBase v.8) |
| ENSG00000070614 | NDST1 | miR-29a/c-3p common target gene | Experiment supported target gene (TarBase v.8) |
| ENSG00000070669 | ASNS | miR-29a-3p target gene only | Experiment supported target gene (TarBase v.8) |
| ENSG00000070831 | CDC42 | miR-29a/c-3p common target gene | Experiment supported target gene (TarBase v.8) |
| ENSG00000070961 | ATP2B1 | miR-29a/c-3p common target gene | Experiment supported target gene (TarBase v.8) |
| ENSG00000071054 | MAP4K4 | miR-29a/c-3p common target gene | Experiment supported target gene (TarBase v.8) |
| ENSG00000071073 | MGAT4A | miR-29a/c-3p common target gene | Experiment supported target gene (TarBase v.8) |
| ENSG00000071246 | VASH1 | miR-29a/c-3p common target gene | Theoretical prediction (microT-CDS algorithm) |
| ENSG00000071575 | TRIB2 | miR-29a/c-3p common target gene | Experiment supported target gene (TarBase v.8) |
| ENSG00000072041 | SLC6A15 | miR-29a-3p target gene only | Experiment supported target gene (TarBase v.8) |
| ENSG00000072042 | RDH11 | miR-29c-3p target gene only | Experiment supported target gene (TarBase v.8) |
| ENSG00000072121 | ZFYVE26 | miR-29a/c-3p common target gene | Experiment supported target gene (TarBase v.8) |
| ENSG00000072134 | EPN2 | miR-29a-3p target gene only | Experiment supported target gene (TarBase v.8) |
| ENSG00000072210 | ALDH3A2 | miR-29a-3p target gene only | Experiment supported target gene (TarBase v.8) |
| ENSG00000072310 | SREBF1 | miR-29a/c-3p common target gene | Experiment supported target gene (TarBase v.8) |
| ENSG00000072364 | AFF4 | miR-29a/c-3p common target gene | Experiment supported target gene (TarBase v.8) |
| ENSG00000072422 | RHOBTB1 | miR-29a/c-3p common target gene | Theoretical prediction (microT-CDS algorithm) |
| ENSG00000072501 | SMC1A | miR-29a/c-3p common target gene | Experiment supported target gene (TarBase v.8) |
| ENSG00000072609 | CHFR | miR-29a/c-3p common target gene | Experiment supported target gene (TarBase v.8) |
| ENSG00000072736 | NFATC3 | miR-29c-3p target gene only | Experiment supported target gene (TarBase v.8) |
| ENSG00000072858 | SIDT1 | miR-29a/c-3p common target gene | Theoretical prediction (microT-CDS algorithm) |
| ENSG00000072864 | NDE1 | miR-29c-3p target gene only | Experiment supported target gene (TarBase v.8) |
| ENSG00000073331 | ALPK1 | miR-29c-3p target gene only | Experiment supported target gene (TarBase v.8) |
| ENSG00000073417 | PDE8A | miR-29a/c-3p common target gene | Theoretical prediction (microT-CDS algorithm) |
| ENSG00000073536 | NLE1 | miR-29a/c-3p common target gene | Experiment supported target gene (TarBase v.8) |
| ENSG00000073578 | SDHA | miR-29a-3p target gene only | Experiment supported target gene (TarBase v.8) |
| ENSG00000073584 | SMARCE1 | miR-29a/c-3p common target gene | Experiment supported target gene (TarBase v.8) |
| ENSG00000073614 | KDM5A | miR-29a/c-3p common target gene | Theoretical prediction (microT-CDS algorithm) |
| ENSG00000073712 | FERMT2 | miR-29a/c-3p common target gene | Experiment supported target gene (TarBase v.8) |

| | | | |
|---|---|---|---|
| ENSG00000073737 | DHRS9 | miR-29c-3p target gene only | Experiment supported target gene (TarBase v.8) |
| ENSG00000073756 | PTGS2 | miR-29a/c-3p common target gene | Experiment supported target gene (TarBase v.8) |
| ENSG00000073849 | ST6GAL1 | miR-29a-3p target gene only | Experiment supported target gene (TarBase v.8) |
| ENSG00000074410 | CA12 | miR-29a/c-3p common target gene | Experiment supported target gene (TarBase v.8) |
| ENSG00000074527 | NTN4 | miR-29a/c-3p common target gene | Experiment supported target gene (TarBase v.8) |
| ENSG00000075151 | EIF4G3 | miR-29a/c-3p common target gene | Experiment supported target gene (TarBase v.8) |
| ENSG00000075223 | SEMA3C | miR-29c-3p target gene only | Experiment supported target gene (TarBase v.8) |
| ENSG00000075407 | ZNF37A | miR-29a/c-3p common target gene | Theoretical prediction (microT-CDS algorithm) |
| ENSG00000075413 | MARK3 | miR-29a/c-3p common target gene | Experiment supported target gene (TarBase v.8) |
| ENSG00000075420 | FNDC3B | miR-29a/c-3p common target gene | Experiment supported target gene (TarBase v.8) |
| ENSG00000075618 | FSCN1 | miR-29a/c-3p common target gene | Experiment supported target gene (TarBase v.8) |
| ENSG00000075624 | ACTB | miR-29a-3p target gene only | Experiment supported target gene (TarBase v.8) |
| ENSG00000076513 | ANKRD13A | miR-29a/c-3p common target gene | Experiment supported target gene (TarBase v.8) |
| ENSG00000076604 | TRAF4 | miR-29a/c-3p common target gene | Experiment supported target gene (TarBase v.8) |
| ENSG00000076641 | PAG1 | miR-29a/c-3p common target gene | Experiment supported target gene (TarBase v.8) |
| ENSG00000076716 | GPC4 | miR-29a/c-3p common target gene | Theoretical prediction (microT-CDS algorithm) |
| ENSG00000076928 | ARHGEF1 | miR-29a-3p target gene only | Experiment supported target gene (TarBase v.8) |
| ENSG00000077044 | DGKD | miR-29a/c-3p common target gene | Experiment supported target gene (TarBase v.8) |
| ENSG00000077063 | CTTNBP2 | miR-29a-3p target gene only | Experiment supported target gene (TarBase v.8) |
| ENSG00000077147 | TM9SF3 | miR-29a/c-3p common target gene | Experiment supported target gene (TarBase v.8) |
| ENSG00000077238 | IL4R | miR-29a/c-3p common target gene | Experiment supported target gene (TarBase v.8) |
| ENSG00000077279 | DCX | miR-29a/c-3p common target gene | Theoretical prediction (microT-CDS algorithm) |
| ENSG00000077458 | FAM76B | miR-29a/c-3p common target gene | Experiment supported target gene (TarBase v.8) |
| ENSG00000077943 | ITGA8 | miR-29a-3p target gene only | Experiment supported target gene (TarBase v.8) |
| ENSG00000078043 | PIAS2 | miR-29a/c-3p common target gene | Experiment supported target gene (TarBase v.8) |
| ENSG00000078081 | LAMP3 | miR-29c-3p target gene only | Experiment supported target gene (TarBase v.8) |
| ENSG00000078098 | FAP | miR-29c-3p target gene only | Experiment supported target gene (TarBase v.8) |
| ENSG00000078124 | ACER3 | miR-29a/c-3p common target gene | Experiment supported target gene (TarBase v.8) |
| ENSG00000078140 | UBE2K | miR-29a/c-3p common target gene | Theoretical prediction (microT-CDS algorithm) |
| ENSG00000078142 | PIK3C3 | miR-29a/c-3p common target gene | Experiment supported target gene (TarBase v.8) |
| ENSG00000078177 | N4BP2 | miR-29a/c-3p common target gene | Experiment supported target gene (TarBase v.8) |
| ENSG00000078369 | GNB1 | miR-29a/c-3p common target gene | Experiment supported target gene (TarBase v.8) |
| ENSG00000078674 | PCM1 | miR-29a/c-3p common target gene | Experiment supported target gene (TarBase v.8) |
| ENSG00000078725 | BRINP1 | miR-29a/c-3p common target gene | Theoretical prediction (microT-CDS algorithm) |
| ENSG00000078804 | TP53INP2 | miR-29a/c-3p common target gene | Theoretical prediction (microT-CDS algorithm) |
| ENSG00000079102 | RUNX1T1 | miR-29a/c-3p common target gene | Theoretical prediction (microT-CDS algorithm) |
| ENSG00000079156 | OSBPL6 | miR-29a/c-3p common target gene | Theoretical prediction (microT-CDS algorithm) |
| ENSG00000079385 | CEACAM1 | miR-29a/c-3p common target gene | Theoretical prediction (microT-CDS algorithm) |
| ENSG00000079432 | CIC | miR-29a-3p target gene only | Experiment supported target gene (TarBase v.8) |
| ENSG00000079785 | DDX1 | miR-29a-3p target gene only | Experiment supported target gene (TarBase v.8) |
| ENSG00000080371 | RAB21 | miR-29a/c-3p common target gene | Experiment supported target gene (TarBase v.8) |
| ENSG00000080503 | SMARCA2 | miR-29a/c-3p common target gene | Theoretical prediction (microT-CDS algorithm) |
| ENSG00000080561 | MID2 | miR-29a-3p target gene only | Experiment supported target gene (TarBase v.8) |
| ENSG00000080573 | COL5A3 | miR-29a/c-3p common target gene | Experiment supported target gene (TarBase v.8) |
| ENSG00000080603 | SRCAP | miR-29a-3p target gene only | Experiment supported target gene (TarBase v.8) |
| ENSG00000081014 | AP4E1 | miR-29a/c-3p common target gene | Theoretical prediction (microT-CDS algorithm) |
| ENSG00000081052 | COL4A4 | miR-29a/c-3p common target gene | Experiment supported target gene (TarBase v.8) |
| ENSG00000081189 | MEF2C | miR-29c-3p target gene only | Experiment supported target gene (TarBase v.8) |
| ENSG00000081842 | PCDHA6 | miR-29a/c-3p common target gene | Theoretical prediction (microT-CDS algorithm) |
| ENSG00000081923 | ATP8B1 | miR-29a-3p target gene only | Experiment supported target gene (TarBase v.8) |
| ENSG00000082014 | SMARCD3 | miR-29c-3p target gene only | Experiment supported target gene (TarBase v.8) |
| ENSG00000082258 | CCNT2 | miR-29a/c-3p common target gene | Experiment supported target gene (TarBase v.8) |
| ENSG00000082269 | FAM135A | miR-29a-3p target gene only | Experiment supported target gene (TarBase v.8) |
| ENSG00000082293 | COL19A1 | miR-29a/c-3p common target gene | Experiment supported target gene (TarBase v.8) |
| ENSG00000082497 | SERTAD4 | miR-29a/c-3p common target gene | Theoretical prediction (microT-CDS algorithm) |
| ENSG00000082512 | TRAF5 | miR-29a-3p target gene only | Experiment supported target gene (TarBase v.8) |
| ENSG00000082701 | GSK3B | miR-29a/c-3p common target gene | Experiment supported target gene (TarBase v.8) |
| ENSG00000082781 | ITGB5 | miR-29a-3p target gene only | Experiment supported target gene (TarBase v.8) |
| ENSG00000082805 | ERC1 | miR-29a/c-3p common target gene | Experiment supported target gene (TarBase v.8) |
| ENSG00000083223 | ZCCHC6 | miR-29a/c-3p common target gene | Experiment supported target gene (TarBase v.8) |
| ENSG00000083544 | TDRD3 | miR-29a/c-3p common target gene | Theoretical prediction (microT-CDS algorithm) |
| ENSG00000083642 | PDS5B | miR-29a/c-3p common target gene | Experiment supported target gene (TarBase v.8) |

| | | | |
|---|---|---|---|
| ENSG00000083720 | OXCT1 | miR-29a/c-3p common target gene | Experiment supported target gene (TarBase v.8) |
| ENSG00000083857 | FAT1 | miR-29a-3p target gene only | Experiment supported target gene (TarBase v.8) |
| ENSG00000084092 | NOA1 | miR-29a-3p target gene only | Experiment supported target gene (TarBase v.8) |
| ENSG00000084093 | REST | miR-29a/c-3p common target gene | Experiment supported target gene (TarBase v.8) |
| ENSG00000084628 | NKAIN1 | miR-29a-3p target gene only | Theoretical prediction (microT-CDS algorithm) |
| ENSG00000084754 | HADHA | miR-29a-3p target gene only | Experiment supported target gene (TarBase v.8) |
| ENSG00000085185 | BCORL1 | miR-29a/c-3p common target gene | Theoretical prediction (microT-CDS algorithm) |
| ENSG00000085224 | ATRX | miR-29a/c-3p common target gene | Experiment supported target gene (TarBase v.8) |
| ENSG00000085449 | WDFY1 | miR-29a/c-3p common target gene | Experiment supported target gene (TarBase v.8) |
| ENSG00000085563 | ABCB1 | miR-29a-3p target gene only | Experiment supported target gene (TarBase v.8) |
| ENSG00000085662 | AKR1B1 | miR-29a-3p target gene only | Experiment supported target gene (TarBase v.8) |
| ENSG00000085741 | WNT11 | miR-29c-3p target gene only | Experiment supported target gene (TarBase v.8) |
| ENSG00000085832 | EPS15 | miR-29a/c-3p common target gene | Experiment supported target gene (TarBase v.8) |
| ENSG00000085978 | ATG16L1 | miR-29a-3p target gene only | Experiment supported target gene (TarBase v.8) |
| ENSG00000085998 | POMGNT1 | miR-29a/c-3p common target gene | Experiment supported target gene (TarBase v.8) |
| ENSG00000086061 | DNAJA1 | miR-29a-3p target gene only | Experiment supported target gene (TarBase v.8) |
| ENSG00000086062 | B4GALT1 | miR-29a/c-3p common target gene | Experiment supported target gene (TarBase v.8) |
| ENSG00000086475 | SEPHS1 | miR-29a/c-3p common target gene | Experiment supported target gene (TarBase v.8) |
| ENSG00000086548 | CEACAM6 | miR-29a/c-3p common target gene | Experiment supported target gene (TarBase v.8) |
| ENSG00000086598 | TMED2 | miR-29a/c-3p common target gene | Experiment supported target gene (TarBase v.8) |
| ENSG00000086696 | HSD17B2 | miR-29a-3p target gene only | Experiment supported target gene (TarBase v.8) |
| ENSG00000086758 | HUWE1 | miR-29a-3p target gene only | Experiment supported target gene (TarBase v.8) |
| ENSG00000087088 | BAX | miR-29a-3p target gene only | Experiment supported target gene (TarBase v.8) |
| ENSG00000087111 | PIGS | miR-29a/c-3p common target gene | Experiment supported target gene (TarBase v.8) |
| ENSG00000087116 | ADAMTS2 | miR-29a/c-3p common target gene | Experiment supported target gene (TarBase v.8) |
| ENSG00000087245 | MMP2 | miR-29a/c-3p common target gene | Experiment supported target gene (TarBase v.8) |
| ENSG00000087301 | TXNDC16 | miR-29c-3p target gene only | Experiment supported target gene (TarBase v.8) |
| ENSG00000087303 | NID2 | miR-29a/c-3p common target gene | Theoretical prediction (microT-CDS algorithm) |
| ENSG00000087448 | KLHL42 | miR-29c-3p target gene only | Experiment supported target gene (TarBase v.8) |
| ENSG00000087494 | PTHLH | miR-29c-3p target gene only | Experiment supported target gene (TarBase v.8) |
| ENSG00000087510 | TFAP2C | miR-29a/c-3p common target gene | Experiment supported target gene (TarBase v.8) |
| ENSG00000087586 | AURKA | miR-29a-3p target gene only | Experiment supported target gene (TarBase v.8) |
| ENSG00000087842 | PIR | miR-29c-3p target gene only | Experiment supported target gene (TarBase v.8) |
| ENSG00000088002 | SULT2B1 | miR-29c-3p target gene only | Experiment supported target gene (TarBase v.8) |
| ENSG00000088035 | ALG6 | miR-29c-3p target gene only | Experiment supported target gene (TarBase v.8) |
| ENSG00000088179 | PTPN4 | miR-29a-3p target gene only | Experiment supported target gene (TarBase v.8) |
| ENSG00000088305 | DNMT3B | miR-29a/c-3p common target gene | Experiment supported target gene (TarBase v.8) |
| ENSG00000088325 | TPX2 | miR-29a-3p target gene only | Experiment supported target gene (TarBase v.8) |
| ENSG00000088356 | PDRG1 | miR-29a/c-3p common target gene | Experiment supported target gene (TarBase v.8) |
| ENSG00000088367 | EPB41L1 | miR-29c-3p target gene only | Experiment supported target gene (TarBase v.8) |
| ENSG00000088387 | DOCK9 | miR-29a-3p target gene only | Experiment supported target gene (TarBase v.8) |
| ENSG00000088448 | ANKRD10 | miR-29a-3p target gene only | Experiment supported target gene (TarBase v.8) |
| ENSG00000088808 | PPP1R13B | miR-29a/c-3p common target gene | Experiment supported target gene (TarBase v.8) |
| ENSG00000088826 | SMOX | miR-29c-3p target gene only | Experiment supported target gene (TarBase v.8) |
| ENSG00000088888 | MAVS | miR-29a/c-3p common target gene | Experiment supported target gene (TarBase v.8) |
| ENSG00000088992 | TESC | miR-29c-3p target gene only | Experiment supported target gene (TarBase v.8) |
| ENSG00000089006 | SNX5 | miR-29a-3p target gene only | Experiment supported target gene (TarBase v.8) |
| ENSG00000089050 | RBBP9 | miR-29a-3p target gene only | Experiment supported target gene (TarBase v.8) |
| ENSG00000089127 | OAS1 | miR-29c-3p target gene only | Experiment supported target gene (TarBase v.8) |
| ENSG00000089157 | RPLP0 | miR-29a/c-3p common target gene | Experiment supported target gene (TarBase v.8) |
| ENSG00000089234 | BRAP | miR-29a/c-3p common target gene | Experiment supported target gene (TarBase v.8) |
| ENSG00000090060 | PAPOLA | miR-29a-3p target gene only | Experiment supported target gene (TarBase v.8) |
| ENSG00000090097 | PCBP4 | miR-29c-3p target gene only | Experiment supported target gene (TarBase v.8) |
| ENSG00000090104 | RGS1 | miR-29a/c-3p common target gene | Experiment supported target gene (TarBase v.8) |
| ENSG00000090372 | STRN4 | miR-29a/c-3p common target gene | Experiment supported target gene (TarBase v.8) |
| ENSG00000090520 | DNAJB11 | miR-29a/c-3p common target gene | Experiment supported target gene (TarBase v.8) |
| ENSG00000090863 | GLG1 | miR-29a-3p target gene only | Experiment supported target gene (TarBase v.8) |
| ENSG00000090905 | TNRC6A | miR-29a/c-3p common target gene | Experiment supported target gene (TarBase v.8) |
| ENSG00000090975 | PITPNM2 | miR-29a/c-3p common target gene | Theoretical prediction (microT-CDS algorithm) |
| ENSG00000091129 | NRCAM | miR-29a/c-3p common target gene | Theoretical prediction (microT-CDS algorithm) |
| ENSG00000091140 | DLD | miR-29a/c-3p common target gene | Experiment supported target gene (TarBase v.8) |
| ENSG00000091317 | CMTM6 | miR-29c-3p target gene only | Experiment supported target gene (TarBase v.8) |

| | | | |
|---|---|---|---|
| ENSG00000091409 | ITGA6 | miR-29a/c-3p common target gene | Experiment supported target gene (TarBase v.8) |
| ENSG00000091482 | SMPX | miR-29c-3p target gene only | Experiment supported target gene (TarBase v.8) |
| ENSG00000091490 | SEL1L3 | miR-29a-3p target gene only | Experiment supported target gene (TarBase v.8) |
| ENSG00000091527 | CDV3 | miR-29a-3p target gene only | Experiment supported target gene (TarBase v.8) |
| ENSG00000091656 | ZFHX4 | miR-29c-3p target gene only | Experiment supported target gene (TarBase v.8) |
| ENSG00000091831 | ESR1 | miR-29a-3p target gene only | Experiment supported target gene (TarBase v.8) |
| ENSG00000092108 | SCFD1 | miR-29a/c-3p common target gene | Experiment supported target gene (TarBase v.8) |
| ENSG00000092140 | G2E3 | miR-29a/c-3p common target gene | Experiment supported target gene (TarBase v.8) |
| ENSG00000092148 | HECTD1 | miR-29a/c-3p common target gene | Experiment supported target gene (TarBase v.8) |
| ENSG00000092208 | GEMIN2 | miR-29a/c-3p common target gene | Experiment supported target gene (TarBase v.8) |
| ENSG00000092345 | DAZL | miR-29a/c-3p common target gene | Theoretical prediction (microT-CDS algorithm) |
| ENSG00000092621 | PHGDH | miR-29a-3p target gene only | Experiment supported target gene (TarBase v.8) |
| ENSG00000092758 | COL9A3 | miR-29c-3p target gene only | Experiment supported target gene (TarBase v.8) |
| ENSG00000092847 | AGO1 | miR-29a/c-3p common target gene | Theoretical prediction (microT-CDS algorithm) |
| ENSG00000092850 | TEKT2 | miR-29c-3p target gene only | Experiment supported target gene (TarBase v.8) |
| ENSG00000092964 | DPYSL2 | miR-29a/c-3p common target gene | Experiment supported target gene (TarBase v.8) |
| ENSG00000092969 | TGFB2 | miR-29a/c-3p common target gene | Experiment supported target gene (TarBase v.8) |
| ENSG00000092978 | GPATCH2 | miR-29a/c-3p common target gene | Theoretical prediction (microT-CDS algorithm) |
| ENSG00000094755 | GABRP | miR-29a/c-3p common target gene | Theoretical prediction (microT-CDS algorithm) |
| ENSG00000094880 | CDC23 | miR-29a/c-3p common target gene | Experiment supported target gene (TarBase v.8) |
| ENSG00000094916 | CBX5 | miR-29a/c-3p common target gene | Experiment supported target gene (TarBase v.8) |
| ENSG00000095066 | HOOK2 | miR-29c-3p target gene only | Experiment supported target gene (TarBase v.8) |
| ENSG00000095139 | ARCN1 | miR-29a/c-3p common target gene | Experiment supported target gene (TarBase v.8) |
| ENSG00000095585 | BLNK | miR-29c-3p target gene only | Experiment supported target gene (TarBase v.8) |
| ENSG00000096433 | ITPR3 | miR-29a-3p target gene only | Experiment supported target gene (TarBase v.8) |
| ENSG00000096717 | SIRT1 | miR-29a/c-3p common target gene | Experiment supported target gene (TarBase v.8) |
| ENSG00000096746 | HNRNPH3 | miR-29a/c-3p common target gene | Experiment supported target gene (TarBase v.8) |
| ENSG00000097007 | ABL1 | miR-29a-3p target gene only | Experiment supported target gene (TarBase v.8) |
| ENSG00000097033 | SH3GLB1 | miR-29a/c-3p common target gene | Experiment supported target gene (TarBase v.8) |
| ENSG00000097046 | CDC7 | miR-29a/c-3p common target gene | Experiment supported target gene (TarBase v.8) |
| ENSG00000099139 | PCSK5 | miR-29a/c-3p common target gene | Experiment supported target gene (TarBase v.8) |
| ENSG00000099194 | SCD | miR-29a-3p target gene only | Experiment supported target gene (TarBase v.8) |
| ENSG00000099219 | ERMP1 | miR-29a-3p target gene only | Experiment supported target gene (TarBase v.8) |
| ENSG00000099800 | TIMM13 | miR-29c-3p target gene only | Experiment supported target gene (TarBase v.8) |
| ENSG00000099812 | MISP | miR-29c-3p target gene only | Experiment supported target gene (TarBase v.8) |
| ENSG00000099864 | PALM | miR-29a/c-3p common target gene | Theoretical prediction (microT-CDS algorithm) |
| ENSG00000099889 | ARVCF | miR-29a/c-3p common target gene | Theoretical prediction (microT-CDS algorithm) |
| ENSG00000099901 | RANBP1 | miR-29a-3p target gene only | Experiment supported target gene (TarBase v.8) |
| ENSG00000099942 | CRKL | miR-29a-3p target gene only | Experiment supported target gene (TarBase v.8) |
| ENSG00000099953 | MMP11 | miR-29a-3p target gene only | Experiment supported target gene (TarBase v.8) |
| ENSG00000100065 | CARD10 | miR-29c-3p target gene only | Experiment supported target gene (TarBase v.8) |
| ENSG00000100075 | SLC25A1 | miR-29c-3p target gene only | Experiment supported target gene (TarBase v.8) |
| ENSG00000100104 | SRRD | miR-29a-3p target gene only | Experiment supported target gene (TarBase v.8) |
| ENSG00000100105 | PATZ1 | miR-29a-3p target gene only | Experiment supported target gene (TarBase v.8) |
| ENSG00000100196 | KDELR3 | miR-29a-3p target gene only | Experiment supported target gene (TarBase v.8) |
| ENSG00000100201 | DDX17 | miR-29a/c-3p common target gene | Experiment supported target gene (TarBase v.8) |
| ENSG00000100211 | CBY1 | miR-29c-3p target gene only | Experiment supported target gene (TarBase v.8) |
| ENSG00000100219 | XBP1 | miR-29a-3p target gene only | Experiment supported target gene (TarBase v.8) |
| ENSG00000100221 | JOSD1 | miR-29a/c-3p common target gene | Experiment supported target gene (TarBase v.8) |
| ENSG00000100227 | POLDIP3 | miR-29c-3p target gene only | Experiment supported target gene (TarBase v.8) |
| ENSG00000100243 | CYB5R3 | miR-29a-3p target gene only | Experiment supported target gene (TarBase v.8) |
| ENSG00000100280 | AP1B1 | miR-29a/c-3p common target gene | Experiment supported target gene (TarBase v.8) |
| ENSG00000100292 | HMOX1 | miR-29c-3p target gene only | Experiment supported target gene (TarBase v.8) |
| ENSG00000100307 | CBX7 | miR-29c-3p target gene only | Experiment supported target gene (TarBase v.8) |
| ENSG00000100311 | PDGFB | miR-29a/c-3p common target gene | Theoretical prediction (microT-CDS algorithm) |
| ENSG00000100314 | CABP7 | miR-29c-3p target gene only | Experiment supported target gene (TarBase v.8) |
| ENSG00000100316 | RPL3 | miR-29a-3p target gene only | Experiment supported target gene (TarBase v.8) |
| ENSG00000100320 | RBFOX2 | miR-29a/c-3p common target gene | Experiment supported target gene (TarBase v.8) |
| ENSG00000100351 | GRAP2 | miR-29a/c-3p common target gene | Theoretical prediction (microT-CDS algorithm) |
| ENSG00000100354 | TNRC6B | miR-29a/c-3p common target gene | Experiment supported target gene (TarBase v.8) |
| ENSG00000100362 | PVALB | miR-29c-3p target gene only | Experiment supported target gene (TarBase v.8) |
| ENSG00000100372 | SLC25A17 | miR-29a/c-3p common target gene | Experiment supported target gene (TarBase v.8) |

| | | | |
|---|---|---|---|
| ENSG00000100393 | EP300 | miR-29a-3p target gene only | Experiment supported target gene (TarBase v.8) |
| ENSG00000100399 | CHADL | miR-29c-3p target gene only | Experiment supported target gene (TarBase v.8) |
| ENSG00000100401 | RANGAP1 | miR-29a/c-3p common target gene | Experiment supported target gene (TarBase v.8) |
| ENSG00000100429 | HDAC10 | miR-29a/c-3p common target gene | Theoretical prediction (microT-CDS algorithm) |
| ENSG00000100439 | ABHD4 | miR-29a/c-3p common target gene | Theoretical prediction (microT-CDS algorithm) |
| ENSG00000100453 | GZMB | miR-29c-3p target gene only | Experiment supported target gene (TarBase v.8) |
| ENSG00000100483 | VCPKMT | miR-29a/c-3p common target gene | Experiment supported target gene (TarBase v.8) |
| ENSG00000100505 | TRIM9 | miR-29a/c-3p common target gene | Theoretical prediction (microT-CDS algorithm) |
| ENSG00000100592 | DAAM1 | miR-29a/c-3p common target gene | Experiment supported target gene (TarBase v.8) |
| ENSG00000100596 | SPTLC2 | miR-29a/c-3p common target gene | Experiment supported target gene (TarBase v.8) |
| ENSG00000100600 | LGMN | miR-29a/c-3p common target gene | Theoretical prediction (microT-CDS algorithm) |
| ENSG00000100644 | HIF1A | miR-29a/c-3p common target gene | Experiment supported target gene (TarBase v.8) |
| ENSG00000100697 | DICER1 | miR-29a/c-3p common target gene | Experiment supported target gene (TarBase v.8) |
| ENSG00000100722 | ZC3H14 | miR-29c-3p target gene only | Experiment supported target gene (TarBase v.8) |
| ENSG00000100811 | YY1 | miR-29c-3p target gene only | Experiment supported target gene (TarBase v.8) |
| ENSG00000100814 | CCNB1IP1 | miR-29a/c-3p common target gene | Experiment supported target gene (TarBase v.8) |
| ENSG00000100911 | PSME2 | miR-29a/c-3p common target gene | Theoretical prediction (microT-CDS algorithm) |
| ENSG00000100918 | REC8 | miR-29c-3p target gene only | Experiment supported target gene (TarBase v.8) |
| ENSG00000100926 | TM9SF1 | miR-29a-3p target gene only | Experiment supported target gene (TarBase v.8) |
| ENSG00000100934 | SEC23A | miR-29a/c-3p common target gene | Experiment supported target gene (TarBase v.8) |
| ENSG00000100991 | TRPC4AP | miR-29a/c-3p common target gene | Experiment supported target gene (TarBase v.8) |
| ENSG00000101000 | PROCR | miR-29c-3p target gene only | Experiment supported target gene (TarBase v.8) |
| ENSG00000101057 | MYBL2 | miR-29a/c-3p common target gene | Experiment supported target gene (TarBase v.8) |
| ENSG00000101115 | SALL4 | miR-29c-3p target gene only | Experiment supported target gene (TarBase v.8) |
| ENSG00000101193 | GID8 | miR-29a/c-3p common target gene | Experiment supported target gene (TarBase v.8) |
| ENSG00000101199 | ARFGAP1 | miR-29a-3p target gene only | Experiment supported target gene (TarBase v.8) |
| ENSG00000101224 | CDC25B | miR-29c-3p target gene only | Experiment supported target gene (TarBase v.8) |
| ENSG00000101246 | ARFRP1 | miR-29c-3p target gene only | Experiment supported target gene (TarBase v.8) |
| ENSG00000101255 | TRIB3 | miR-29c-3p target gene only | Experiment supported target gene (TarBase v.8) |
| ENSG00000101265 | RASSF2 | miR-29c-3p target gene only | Experiment supported target gene (TarBase v.8) |
| ENSG00000101290 | CDS2 | miR-29a-3p target gene only | Experiment supported target gene (TarBase v.8) |
| ENSG00000101347 | SAMHD1 | miR-29a-3p target gene only | Experiment supported target gene (TarBase v.8) |
| ENSG00000101367 | MAPRE1 | miR-29a/c-3p common target gene | Theoretical prediction (microT-CDS algorithm) |
| ENSG00000101384 | JAG1 | miR-29a-3p target gene only | Experiment supported target gene (TarBase v.8) |
| ENSG00000101544 | ADNP2 | miR-29a/c-3p common target gene | Experiment supported target gene (TarBase v.8) |
| ENSG00000101624 | CEP76 | miR-29a/c-3p common target gene | Theoretical prediction (microT-CDS algorithm) |
| ENSG00000101680 | LAMA1 | miR-29a-3p target gene only | Experiment supported target gene (TarBase v.8) |
| ENSG00000101752 | MIB1 | miR-29a/c-3p common target gene | Experiment supported target gene (TarBase v.8) |
| ENSG00000101782 | RIOK3 | miR-29a/c-3p common target gene | Theoretical prediction (microT-CDS algorithm) |
| ENSG00000101901 | ALG13 | miR-29a/c-3p common target gene | Experiment supported target gene (TarBase v.8) |
| ENSG00000101955 | SRPX | miR-29a/c-3p common target gene | Experiment supported target gene (TarBase v.8) |
| ENSG00000101966 | XIAP | miR-29c-3p target gene only | Experiment supported target gene (TarBase v.8) |
| ENSG00000102038 | SMARCA1 | miR-29a/c-3p common target gene | Experiment supported target gene (TarBase v.8) |
| ENSG00000102078 | SLC25A14 | miR-29c-3p target gene only | Experiment supported target gene (TarBase v.8) |
| ENSG00000102096 | PIM2 | miR-29a/c-3p common target gene | Experiment supported target gene (TarBase v.8) |
| ENSG00000102098 | SCML2 | miR-29a/c-3p common target gene | Theoretical prediction (microT-CDS algorithm) |
| ENSG00000102100 | SLC35A2 | miR-29a/c-3p common target gene | Experiment supported target gene (TarBase v.8) |
| ENSG00000102172 | SMS | miR-29a/c-3p common target gene | Experiment supported target gene (TarBase v.8) |
| ENSG00000102174 | PHEX | miR-29a/c-3p common target gene | Theoretical prediction (microT-CDS algorithm) |
| ENSG00000102225 | CDK16 | miR-29a/c-3p common target gene | Experiment supported target gene (TarBase v.8) |
| ENSG00000102230 | PCYT1B | miR-29a/c-3p common target gene | Theoretical prediction (microT-CDS algorithm) |
| ENSG00000102243 | VGLL1 | miR-29a/c-3p common target gene | Theoretical prediction (microT-CDS algorithm) |
| ENSG00000102287 | GABRE | miR-29c-3p target gene only | Experiment supported target gene (TarBase v.8) |
| ENSG00000102384 | CENPI | miR-29a-3p target gene only | Experiment supported target gene (TarBase v.8) |
| ENSG00000102385 | DRP2 | miR-29a/c-3p common target gene | Theoretical prediction (microT-CDS algorithm) |
| ENSG00000102543 | CDADC1 | miR-29c-3p target gene only | Experiment supported target gene (TarBase v.8) |
| ENSG00000102547 | CAB39L | miR-29c-3p target gene only | Experiment supported target gene (TarBase v.8) |
| ENSG00000102780 | DGKH | miR-29a/c-3p common target gene | Theoretical prediction (microT-CDS algorithm) |
| ENSG00000102781 | KATNAL1 | miR-29a/c-3p common target gene | Experiment supported target gene (TarBase v.8) |
| ENSG00000102908 | NFAT5 | miR-29a/c-3p common target gene | Theoretical prediction (microT-CDS algorithm) |
| ENSG00000102934 | PLLP | miR-29c-3p target gene only | Experiment supported target gene (TarBase v.8) |
| ENSG00000102967 | DHODH | miR-29c-3p target gene only | Experiment supported target gene (TarBase v.8) |

| | | | |
|---|---|---|---|
| ENSG00000102978 | POLR2C | miR-29c-3p target gene only | Experiment supported target gene (TarBase v.8) |
| ENSG00000102996 | MMP15 | miR-29c-3p target gene only | Experiment supported target gene (TarBase v.8) |
| ENSG00000103034 | NDRG4 | miR-29a/c-3p common target gene | Theoretical prediction (microT-CDS algorithm) |
| ENSG00000103035 | PSMD7 | miR-29c-3p target gene only | Experiment supported target gene (TarBase v.8) |
| ENSG00000103044 | HAS3 | miR-29a/c-3p common target gene | Theoretical prediction (microT-CDS algorithm) |
| ENSG00000103051 | COG4 | miR-29a/c-3p common target gene | Theoretical prediction (microT-CDS algorithm) |
| ENSG00000103056 | SMPD3 | miR-29a/c-3p common target gene | Theoretical prediction (microT-CDS algorithm) |
| ENSG00000103067 | ESRP2 | miR-29a/c-3p common target gene | Theoretical prediction (microT-CDS algorithm) |
| ENSG00000103175 | WFDC1 | miR-29a-3p target gene only | Experiment supported target gene (TarBase v.8) |
| ENSG00000103194 | USP10 | miR-29a/c-3p common target gene | Experiment supported target gene (TarBase v.8) |
| ENSG00000103257 | SLC7A5 | miR-29a-3p target gene only | Experiment supported target gene (TarBase v.8) |
| ENSG00000103275 | UBE2I | miR-29a-3p target gene only | Experiment supported target gene (TarBase v.8) |
| ENSG00000103351 | CLUAP1 | miR-29c-3p target gene only | Experiment supported target gene (TarBase v.8) |
| ENSG00000103353 | UBFD1 | miR-29a/c-3p common target gene | Experiment supported target gene (TarBase v.8) |
| ENSG00000103404 | USP31 | miR-29a/c-3p common target gene | Experiment supported target gene (TarBase v.8) |
| ENSG00000103449 | SALL1 | miR-29c-3p target gene only | Experiment supported target gene (TarBase v.8) |
| ENSG00000103490 | PYCARD | miR-29c-3p target gene only | Experiment supported target gene (TarBase v.8) |
| ENSG00000103495 | MAZ | miR-29a/c-3p common target gene | Experiment supported target gene (TarBase v.8) |
| ENSG00000103550 | KNOP1 | miR-29a/c-3p common target gene | Experiment supported target gene (TarBase v.8) |
| ENSG00000103599 | IQCH | miR-29a/c-3p common target gene | Experiment supported target gene (TarBase v.8) |
| ENSG00000103657 | HERC1 | miR-29a/c-3p common target gene | Experiment supported target gene (TarBase v.8) |
| ENSG00000103811 | CTSH | miR-29c-3p target gene only | Experiment supported target gene (TarBase v.8) |
| ENSG00000103855 | CD276 | miR-29a/c-3p common target gene | Experiment supported target gene (TarBase v.8) |
| ENSG00000103888 | CEMIP | miR-29a/c-3p common target gene | Experiment supported target gene (TarBase v.8) |
| ENSG00000103942 | HOMER2 | miR-29c-3p target gene only | Experiment supported target gene (TarBase v.8) |
| ENSG00000103994 | ZNF106 | miR-29a-3p target gene only | Experiment supported target gene (TarBase v.8) |
| ENSG00000104064 | GABPB1 | miR-29a/c-3p common target gene | Experiment supported target gene (TarBase v.8) |
| ENSG00000104067 | TJP1 | miR-29a-3p target gene only | Experiment supported target gene (TarBase v.8) |
| ENSG00000104081 | BMF | miR-29a/c-3p common target gene | Experiment supported target gene (TarBase v.8) |
| ENSG00000104093 | DMXL2 | miR-29a/c-3p common target gene | Experiment supported target gene (TarBase v.8) |
| ENSG00000104131 | EIF3J | miR-29a/c-3p common target gene | Experiment supported target gene (TarBase v.8) |
| ENSG00000104142 | VPS18 | miR-29c-3p target gene only | Experiment supported target gene (TarBase v.8) |
| ENSG00000104147 | OIP5 | miR-29c-3p target gene only | Experiment supported target gene (TarBase v.8) |
| ENSG00000104213 | PDGFRL | miR-29a-3p target gene only | Experiment supported target gene (TarBase v.8) |
| ENSG00000104267 | CA2 | miR-29a/c-3p common target gene | Experiment supported target gene (TarBase v.8) |
| ENSG00000104332 | SFRP1 | miR-29a-3p target gene only | Experiment supported target gene (TarBase v.8) |
| ENSG00000104341 | LAPTM4B | miR-29a/c-3p common target gene | Experiment supported target gene (TarBase v.8) |
| ENSG00000104412 | EMC2 | miR-29a/c-3p common target gene | Experiment supported target gene (TarBase v.8) |
| ENSG00000104415 | WISP1 | miR-29a/c-3p common target gene | Theoretical prediction (microT-CDS algorithm) |
| ENSG00000104435 | STMN2 | miR-29a/c-3p common target gene | Experiment supported target gene (TarBase v.8) |
| ENSG00000104447 | TRPS1 | miR-29a/c-3p common target gene | Experiment supported target gene (TarBase v.8) |
| ENSG00000104497 | SNX16 | miR-29a/c-3p common target gene | Experiment supported target gene (TarBase v.8) |
| ENSG00000104549 | SQLE | miR-29a/c-3p common target gene | Experiment supported target gene (TarBase v.8) |
| ENSG00000104611 | SH2D4A | miR-29a-3p target gene only | Experiment supported target gene (TarBase v.8) |
| ENSG00000104643 | MTMR9 | miR-29a/c-3p common target gene | Experiment supported target gene (TarBase v.8) |
| ENSG00000104714 | ERICH1 | miR-29a/c-3p common target gene | Experiment supported target gene (TarBase v.8) |
| ENSG00000104722 | NEFM | miR-29a/c-3p common target gene | Experiment supported target gene (TarBase v.8) |
| ENSG00000104756 | KCTD9 | miR-29c-3p target gene only | Experiment supported target gene (TarBase v.8) |
| ENSG00000104783 | KCNN4 | miR-29c-3p target gene only | Experiment supported target gene (TarBase v.8) |
| ENSG00000104885 | DOT1L | miR-29a/c-3p common target gene | Experiment supported target gene (TarBase v.8) |
| ENSG00000104904 | OAZ1 | miR-29a-3p target gene only | Experiment supported target gene (TarBase v.8) |
| ENSG00000104967 | NOVA2 | miR-29a/c-3p common target gene | Experiment supported target gene (TarBase v.8) |
| ENSG00000104980 | TIMM44 | miR-29a/c-3p common target gene | Experiment supported target gene (TarBase v.8) |
| ENSG00000105011 | ASF1B | miR-29a-3p target gene only | Experiment supported target gene (TarBase v.8) |
| ENSG00000105085 | MED26 | miR-29a/c-3p common target gene | Theoretical prediction (microT-CDS algorithm) |
| ENSG00000105173 | CCNE1 | miR-29a/c-3p common target gene | Experiment supported target gene (TarBase v.8) |
| ENSG00000105220 | GPI | miR-29a/c-3p common target gene | Experiment supported target gene (TarBase v.8) |
| ENSG00000105221 | AKT2 | miR-29a/c-3p common target gene | Experiment supported target gene (TarBase v.8) |
| ENSG00000105323 | HNRNPUL1 | miR-29a/c-3p common target gene | Experiment supported target gene (TarBase v.8) |
| ENSG00000105559 | PLEKHA4 | miR-29c-3p target gene only | Experiment supported target gene (TarBase v.8) |
| ENSG00000105643 | ARRDC2 | miR-29c-3p target gene only | Experiment supported target gene (TarBase v.8) |
| ENSG00000105647 | PIK3R2 | miR-29a/c-3p common target gene | Theoretical prediction (microT-CDS algorithm) |
| ENSG00000105655 | ISYNA1 | miR-29a/c-3p common target gene | Experiment supported target gene (TarBase v.8) |
| ENSG00000105671 | DDX49 | miR-29c-3p target gene only | Experiment supported target gene (TarBase v.8) |
| ENSG00000105722 | ERF | miR-29a-3p target gene only | Experiment supported target gene (TarBase v.8) |
| ENSG00000105810 | CDK6 | miR-29a/c-3p common target gene | Experiment supported target gene (TarBase v.8) |
| ENSG00000105825 | TFPI2 | miR-29c-3p target gene only | Experiment supported target gene (TarBase v.8) |
| ENSG00000105856 | HBP1 | miR-29a/c-3p common target gene | Experiment supported target gene (TarBase v.8) |
| ENSG00000105879 | CBLL1 | miR-29a/c-3p common target gene | Experiment supported target gene (TarBase v.8) |
| ENSG00000105887 | MTPN | miR-29a/c-3p common target gene | Experiment supported target gene (TarBase v.8) |
| ENSG00000105967 | TFEC | miR-29a/c-3p common target gene | Theoretical prediction (microT-CDS algorithm) |
| ENSG00000105971 | CAV2 | miR-29a/c-3p common target gene | Experiment supported target gene (TarBase v.8) |
| ENSG00000105974 | CAV1 | miR-29c-3p target gene only | Experiment supported target gene (TarBase v.8) |
| ENSG00000106025 | TSPAN12 | miR-29a-3p target gene only | Experiment supported target gene (TarBase v.8) |
| ENSG00000106086 | PLEKHA8 | miR-29a/c-3p common target gene | Theoretical prediction (microT-CDS algorithm) |
| ENSG00000106144 | CASP2 | miR-29a/c-3p common target gene | Experiment supported target gene (TarBase v.8) |
| ENSG00000106261 | ZKSCAN1 | miR-29c-3p target gene only | Experiment supported target gene (TarBase v.8) |
| ENSG00000106305 | AIMP2 | miR-29a/c-3p common target gene | Experiment supported target gene (TarBase v.8) |
| ENSG00000106344 | RBM28 | miR-29a/c-3p common target gene | Experiment supported target gene (TarBase v.8) |
| ENSG00000106346 | USP42 | miR-29a/c-3p common target gene | Experiment supported target gene (TarBase v.8) |
| ENSG00000106348 | IMPDH1 | miR-29a/c-3p common target gene | Experiment supported target gene (TarBase v.8) |
| ENSG00000106462 | EZH2 | miR-29a/c-3p common target gene | Experiment supported target gene (TarBase v.8) |
| ENSG00000106477 | CEP41 | miR-29a/c-3p common target gene | Experiment supported target gene (TarBase v.8) |
| ENSG00000106479 | ZNF862 | miR-29a/c-3p common target gene | Theoretical prediction (microT-CDS algorithm) |
| ENSG00000106484 | MEST | miR-29a/c-3p common target gene | Experiment supported target gene (TarBase v.8) |
| ENSG00000106541 | AGR2 | miR-29a/c-3p common target gene | Theoretical prediction (microT-CDS algorithm) |
| ENSG00000106546 | AHR | miR-29a/c-3p common target gene | Experiment supported target gene (TarBase v.8) |
| ENSG00000106608 | URGCP | miR-29a/c-3p common target gene | Experiment supported target gene (TarBase v.8) |
| ENSG00000106682 | EIF4H | miR-29a/c-3p common target gene | Experiment supported target gene (TarBase v.8) |
| ENSG00000106683 | LIMK1 | miR-29c-3p target gene only | Experiment supported target gene (TarBase v.8) |
| ENSG00000106688 | SLC1A1 | miR-29a-3p target gene only | Experiment supported target gene (TarBase v.8) |
| ENSG00000106780 | MEGF9 | miR-29a/c-3p common target gene | Experiment supported target gene (TarBase v.8) |
| ENSG00000106993 | CDC37L1 | miR-29c-3p target gene only | Experiment supported target gene (TarBase v.8) |
| ENSG00000107201 | DDX58 | miR-29c-3p target gene only | Experiment supported target gene (TarBase v.8) |
| ENSG00000107443 | CCNJ | miR-29a/c-3p common target gene | Experiment supported target gene (TarBase v.8) |
| ENSG00000107518 | ATRNL1 | miR-29a-3p target gene only | Experiment supported target gene (TarBase v.8) |
| ENSG00000107521 | HPS1 | miR-29a-3p target gene only | Experiment supported target gene (TarBase v.8) |
| ENSG00000107581 | EIF3A | miR-29a-3p target gene only | Experiment supported target gene (TarBase v.8) |
| ENSG00000107643 | MAPK8 | miR-29a/c-3p common target gene | Experiment supported target gene (TarBase v.8) |
| ENSG00000107719 | PALD1 | miR-29a-3p target gene only | Experiment supported target gene (TarBase v.8) |
| ENSG00000107738 | C10orf54 | miR-29a/c-3p common target gene | Theoretical prediction (microT-CDS algorithm) |
| ENSG00000107771 | CCSER2 | miR-29a/c-3p common target gene | Experiment supported target gene (TarBase v.8) |
| ENSG00000107796 | ACTA2 | miR-29c-3p target gene only | Experiment supported target gene (TarBase v.8) |
| ENSG00000107833 | NPM3 | miR-29a/c-3p common target gene | Theoretical prediction (microT-CDS algorithm) |
| ENSG00000107864 | CPEB3 | miR-29a/c-3p common target gene | Experiment supported target gene (TarBase v.8) |
| ENSG00000107882 | SUFU | miR-29a/c-3p common target gene | Experiment supported target gene (TarBase v.8) |
| ENSG00000107929 | LARP4B | miR-29a/c-3p common target gene | Experiment supported target gene (TarBase v.8) |
| ENSG00000107957 | SH3PXD2A | miR-29a/c-3p common target gene | Experiment supported target gene (TarBase v.8) |
| ENSG00000107984 | DKK1 | miR-29c-3p target gene only | Experiment supported target gene (TarBase v.8) |
| ENSG00000108021 | FAM208B | miR-29a-3p target gene only | Experiment supported target gene (TarBase v.8) |
| ENSG00000108039 | XPNPEP1 | miR-29a/c-3p common target gene | Theoretical prediction (microT-CDS algorithm) |
| ENSG00000108175 | ZMIZ1 | miR-29a/c-3p common target gene | Experiment supported target gene (TarBase v.8) |
| ENSG00000108219 | TSPAN14 | miR-29a/c-3p common target gene | Theoretical prediction (microT-CDS algorithm) |
| ENSG00000108231 | LGI1 | miR-29c-3p target gene only | Experiment supported target gene (TarBase v.8) |
| ENSG00000108256 | NUFIP2 | miR-29a/c-3p common target gene | Experiment supported target gene (TarBase v.8) |
| ENSG00000108312 | UBTF | miR-29a/c-3p common target gene | Theoretical prediction (microT-CDS algorithm) |
| ENSG00000108352 | RAPGEFL1 | miR-29a/c-3p common target gene | Theoretical prediction (microT-CDS algorithm) |
| ENSG00000108379 | WNT3 | miR-29c-3p target gene only | Experiment supported target gene (TarBase v.8) |
| ENSG00000108384 | RAD51C | miR-29c-3p target gene only | Experiment supported target gene (TarBase v.8) |
| ENSG00000108387 | 4-Sep | miR-29c-3p target gene only | Experiment supported target gene (TarBase v.8) |
| ENSG00000108389 | MTMR4 | miR-29a/c-3p common target gene | Theoretical prediction (microT-CDS algorithm) |
| ENSG00000108395 | TRIM37 | miR-29a/c-3p common target gene | Experiment supported target gene (TarBase v.8) |
| ENSG00000108423 | TUBD1 | miR-29a/c-3p common target gene | Experiment supported target gene (TarBase v.8) |
| ENSG00000108448 | TRIM16L | miR-29a/c-3p common target gene | Theoretical prediction (microT-CDS algorithm) |
| ENSG00000108510 | MED13 | miR-29a/c-3p common target gene | Experiment supported target gene (TarBase v.8) |
| ENSG00000108523 | RNF167 | miR-29c-3p target gene only | Experiment supported target gene (TarBase v.8) |
| ENSG00000108578 | BLMH | miR-29a/c-3p common target gene | Experiment supported target gene (TarBase v.8) |
| ENSG00000108582 | CPD | miR-29a-3p target gene only | Experiment supported target gene (TarBase v.8) |
| ENSG00000108599 | AKAP10 | miR-29a/c-3p common target gene | Experiment supported target gene (TarBase v.8) |
| ENSG00000108654 | DDX5 | miR-29a/c-3p common target gene | Theoretical prediction (microT-CDS algorithm) |
| ENSG00000108788 | MLX | miR-29c-3p target gene only | Experiment supported target gene (TarBase v.8) |
| ENSG00000108821 | COL1A1 | miR-29a/c-3p common target gene | Experiment supported target gene (TarBase v.8) |
| ENSG00000108826 | MRPL27 | miR-29a-3p target gene only | Experiment supported target gene (TarBase v.8) |
| ENSG00000108829 | LRRC59 | miR-29a/c-3p common target gene | Theoretical prediction (microT-CDS algorithm) |
| ENSG00000108846 | ABCC3 | miR-29c-3p target gene only | Experiment supported target gene (TarBase v.8) |
| ENSG00000108848 | LUC7L3 | miR-29a/c-3p common target gene | Experiment supported target gene (TarBase v.8) |
| ENSG00000108854 | SMURF2 | miR-29a/c-3p common target gene | Theoretical prediction (microT-CDS algorithm) |
| ENSG00000108883 | EFTUD2 | miR-29a-3p target gene only | Experiment supported target gene (TarBase v.8) |
| ENSG00000108924 | HLF | miR-29c-3p target gene only | Experiment supported target gene (TarBase v.8) |
| ENSG00000108932 | SLC16A6 | miR-29a-3p target gene only | Experiment supported target gene (TarBase v.8) |
| ENSG00000108953 | YWHAE | miR-29a/c-3p common target gene | Experiment supported target gene (TarBase v.8) |
| ENSG00000108960 | MMD | miR-29c-3p target gene only | Experiment supported target gene (TarBase v.8) |
| ENSG00000108984 | MAP2K6 | miR-29a/c-3p common target gene | Experiment supported target gene (TarBase v.8) |
| ENSG00000109079 | TNFAIP1 | miR-29c-3p target gene only | Experiment supported target gene (TarBase v.8) |
| ENSG00000109099 | PMP22 | miR-29a/c-3p common target gene | Experiment supported target gene (TarBase v.8) |
| ENSG00000109107 | ALDOC | miR-29a-3p target gene only | Experiment supported target gene (TarBase v.8) |
| ENSG00000109133 | TMEM33 | miR-29c-3p target gene only | Experiment supported target gene (TarBase v.8) |
| ENSG00000109184 | DCUN1D4 | miR-29a/c-3p common target gene | Theoretical prediction (microT-CDS algorithm) |
| ENSG00000109220 | CHIC2 | miR-29a/c-3p common target gene | Experiment supported target gene (TarBase v.8) |
| ENSG00000109321 | AREG | miR-29a/c-3p common target gene | Experiment supported target gene (TarBase v.8) |
| ENSG00000109339 | MAPK10 | miR-29a/c-3p common target gene | Experiment supported target gene (TarBase v.8) |
| ENSG00000109381 | ELF2 | miR-29a/c-3p common target gene | Experiment supported target gene (TarBase v.8) |
| ENSG00000109424 | UCP1 | miR-29c-3p target gene only | Experiment supported target gene (TarBase v.8) |
| ENSG00000109618 | SEPSECS | miR-29a/c-3p common target gene | Theoretical prediction (microT-CDS algorithm) |
| ENSG00000109670 | FBXW7 | miR-29a/c-3p common target gene | Experiment supported target gene (TarBase v.8) |
| ENSG00000109685 | WHSC1 | miR-29a/c-3p common target gene | Theoretical prediction (microT-CDS algorithm) |
| ENSG00000109689 | STIM2 | miR-29a/c-3p common target gene | Experiment supported target gene (TarBase v.8) |
| ENSG00000109775 | UFSP2 | miR-29c-3p target gene only | Experiment supported target gene (TarBase v.8) |
| ENSG00000109854 | HTATIP2 | miR-29a/c-3p common target gene | Theoretical prediction (microT-CDS algorithm) |
| ENSG00000109929 | SC5D | miR-29a/c-3p common target gene | Experiment supported target gene (TarBase v.8) |
| ENSG00000110002 | VWA5A | miR-29a/c-3p common target gene | Experiment supported target gene (TarBase v.8) |
| ENSG00000110042 | DTX4 | miR-29a/c-3p common target gene | Experiment supported target gene (TarBase v.8) |
| ENSG00000110047 | EHD1 | miR-29a-3p target gene only | Experiment supported target gene (TarBase v.8) |
| ENSG00000110048 | OSBP | miR-29a/c-3p common target gene | Theoretical prediction (microT-CDS algorithm) |
| ENSG00000110075 | PPP6R3 | miR-29a-3p target gene only | Experiment supported target gene (TarBase v.8) |
| ENSG00000110104 | CCDC86 | miR-29c-3p target gene only | Experiment supported target gene (TarBase v.8) |
| ENSG00000110107 | PRPF19 | miR-29c-3p target gene only | Experiment supported target gene (TarBase v.8) |
| ENSG00000110203 | FOLR3 | miR-29c-3p target gene only | Experiment supported target gene (TarBase v.8) |
| ENSG00000110315 | RNF141 | miR-29a/c-3p common target gene | Experiment supported target gene (TarBase v.8) |
| ENSG00000110330 | BIRC2 | miR-29a/c-3p common target gene | Experiment supported target gene (TarBase v.8) |
| ENSG00000110344 | UBE4A | miR-29c-3p target gene only | Experiment supported target gene (TarBase v.8) |
| ENSG00000110367 | DDX6 | miR-29a/c-3p common target gene | Experiment supported target gene (TarBase v.8) |
| ENSG00000110427 | KIAA1549L | miR-29a/c-3p common target gene | Theoretical prediction (microT-CDS algorithm) |
| ENSG00000110429 | FBXO3 | miR-29c-3p target gene only | Experiment supported target gene (TarBase v.8) |
| ENSG00000110435 | PDHX | miR-29c-3p target gene only | Experiment supported target gene (TarBase v.8) |
| ENSG00000110436 | SLC1A2 | miR-29a/c-3p common target gene | Experiment supported target gene (TarBase v.8) |
| ENSG00000110446 | SLC15A3 | miR-29c-3p target gene only | Experiment supported target gene (TarBase v.8) |
| ENSG00000110514 | MADD | miR-29a-3p target gene only | Experiment supported target gene (TarBase v.8) |
| ENSG00000110693 | SOX6 | miR-29a/c-3p common target gene | Theoretical prediction (microT-CDS algorithm) |
| ENSG00000110777 | POU2AF1 | miR-29c-3p target gene only | Experiment supported target gene (TarBase v.8) |
| ENSG00000110799 | VWF | miR-29a-3p target gene only | Experiment supported target gene (TarBase v.8) |
| ENSG00000110841 | PPFIBP1 | miR-29a-3p target gene only | Experiment supported target gene (TarBase v.8) |
| ENSG00000110851 | PRDM4 | miR-29a-3p target gene only | Experiment supported target gene (TarBase v.8) |
| ENSG00000110871 | COQ5 | miR-29c-3p target gene only | Experiment supported target gene (TarBase v.8) |
| ENSG00000110876 | SELPLG | miR-29a-3p target gene only | Experiment supported target gene (TarBase v.8) |
| ENSG00000110880 | CORO1C | miR-29a-3p target gene only | Experiment supported target gene (TarBase v.8) |
| ENSG00000110881 | ASIC1 | miR-29a/c-3p common target gene | Theoretical prediction (microT-CDS algorithm) |
| ENSG00000110888 | CAPRIN2 | miR-29a/c-3p common target gene | Theoretical prediction (microT-CDS algorithm) |
| ENSG00000110911 | SLC11A2 | miR-29a-3p target gene only | Experiment supported target gene (TarBase v.8) |
| ENSG00000110925 | CSRNP2 | miR-29a/c-3p common target gene | Theoretical prediction (microT-CDS algorithm) |
| ENSG00000110931 | CAMKK2 | miR-29a/c-3p common target gene | Theoretical prediction (microT-CDS algorithm) |
| ENSG00000110987 | BCL7A | miR-29a-3p target gene only | Experiment supported target gene (TarBase v.8) |
| ENSG00000111011 | RSRC2 | miR-29c-3p target gene only | Experiment supported target gene (TarBase v.8) |
| ENSG00000111052 | LIN7A | miR-29a/c-3p common target gene | Theoretical prediction (microT-CDS algorithm) |
| ENSG00000111142 | METAP2 | miR-29a/c-3p common target gene | Experiment supported target gene (TarBase v.8) |
| ENSG00000111196 | MAGOHB | miR-29a-3p target gene only | Experiment supported target gene (TarBase v.8) |
| ENSG00000111224 | PARP11 | miR-29a/c-3p common target gene | Theoretical prediction (microT-CDS algorithm) |
| ENSG00000111276 | CDKN1B | miR-29c-3p target gene only | Experiment supported target gene (TarBase v.8) |
| ENSG00000111331 | OAS3 | miR-29c-3p target gene only | Experiment supported target gene (TarBase v.8) |
| ENSG00000111335 | OAS2 | miR-29a-3p target gene only | Experiment supported target gene (TarBase v.8) |
| ENSG00000111361 | EIF2B1 | miR-29c-3p target gene only | Experiment supported target gene (TarBase v.8) |
| ENSG00000111371 | SLC38A1 | miR-29a-3p target gene only | Experiment supported target gene (TarBase v.8) |
| ENSG00000111432 | FZD10 | miR-29c-3p target gene only | Experiment supported target gene (TarBase v.8) |
| ENSG00000111530 | CAND1 | miR-29a/c-3p common target gene | Experiment supported target gene (TarBase v.8) |
| ENSG00000111596 | CNOT2 | miR-29a/c-3p common target gene | Experiment supported target gene (TarBase v.8) |
| ENSG00000111615 | KRR1 | miR-29a-3p target gene only | Experiment supported target gene (TarBase v.8) |
| ENSG00000111640 | GAPDH | miR-29a/c-3p common target gene | Experiment supported target gene (TarBase v.8) |
| ENSG00000111725 | PRKAB1 | miR-29c-3p target gene only | Experiment supported target gene (TarBase v.8) |
| ENSG00000111726 | CMAS | miR-29c-3p target gene only | Experiment supported target gene (TarBase v.8) |
| ENSG00000111732 | AICDA | miR-29a/c-3p common target gene | Theoretical prediction (microT-CDS algorithm) |
| ENSG00000111737 | RAB35 | miR-29c-3p target gene only | Experiment supported target gene (TarBase v.8) |
| ENSG00000111816 | FRK | miR-29a/c-3p common target gene | Theoretical prediction (microT-CDS algorithm) |
| ENSG00000111832 | RWDD1 | miR-29a-3p target gene only | Experiment supported target gene (TarBase v.8) |
| ENSG00000111846 | GCNT2 | miR-29a-3p target gene only | Experiment supported target gene (TarBase v.8) |
| ENSG00000111859 | NEDD9 | miR-29a/c-3p common target gene | Theoretical prediction (microT-CDS algorithm) |
| ENSG00000111860 | CEP85L | miR-29a/c-3p common target gene | Theoretical prediction (microT-CDS algorithm) |
| ENSG00000111907 | TPD52L1 | miR-29a-3p target gene only | Experiment supported target gene (TarBase v.8) |
| ENSG00000111911 | HINT3 | miR-29a/c-3p common target gene | Experiment supported target gene (TarBase v.8) |
| ENSG00000111913 | FAM65B | miR-29a/c-3p common target gene | Theoretical prediction (microT-CDS algorithm) |
| ENSG00000111961 | SASH1 | miR-29a-3p target gene only | Experiment supported target gene (TarBase v.8) |
| ENSG00000112078 | KCTD20 | miR-29a/c-3p common target gene | Experiment supported target gene (TarBase v.8) |
| ENSG00000112079 | STK38 | miR-29a-3p target gene only | Experiment supported target gene (TarBase v.8) |
| ENSG00000112182 | BACH2 | miR-29a/c-3p common target gene | Experiment supported target gene (TarBase v.8) |
| ENSG00000112208 | BAG2 | miR-29a-3p target gene only | Experiment supported target gene (TarBase v.8) |
| ENSG00000112245 | PTP4A1 | miR-29a/c-3p common target gene | Experiment supported target gene (TarBase v.8) |
| ENSG00000112280 | COL9A1 | miR-29a/c-3p common target gene | Theoretical prediction (microT-CDS algorithm) |
| ENSG00000112290 | WASF1 | miR-29c-3p target gene only | Experiment supported target gene (TarBase v.8) |
| ENSG00000112297 | AIM1 | miR-29a/c-3p common target gene | Experiment supported target gene (TarBase v.8) |
| ENSG00000112305 | SMAP1 | miR-29c-3p target gene only | Experiment supported target gene (TarBase v.8) |
| ENSG00000112394 | SLC16A10 | miR-29a/c-3p common target gene | Theoretical prediction (microT-CDS algorithm) |
| ENSG00000112419 | PHACTR2 | miR-29a/c-3p common target gene | Theoretical prediction (microT-CDS algorithm) |
| ENSG00000112531 | QKI | miR-29a/c-3p common target gene | Experiment supported target gene (TarBase v.8) |
| ENSG00000112561 | TFEB | miR-29a/c-3p common target gene | Theoretical prediction (microT-CDS algorithm) |
| ENSG00000112562 | SMOC2 | miR-29c-3p target gene only | Experiment supported target gene (TarBase v.8) |
| ENSG00000112659 | CUL9 | miR-29c-3p target gene only | Experiment supported target gene (TarBase v.8) |
| ENSG00000112695 | COX7A2 | miR-29a-3p target gene only | Experiment supported target gene (TarBase v.8) |
| ENSG00000112697 | TMEM30A | miR-29a-3p target gene only | Experiment supported target gene (TarBase v.8) |
| ENSG00000112715 | VEGFA | miR-29a/c-3p common target gene | Experiment supported target gene (TarBase v.8) |
| ENSG00000112796 | ENPP5 | miR-29a/c-3p common target gene | Experiment supported target gene (TarBase v.8) |
| ENSG00000112837 | TBX18 | miR-29a-3p target gene only | Experiment supported target gene (TarBase v.8) |
| ENSG00000112972 | HMGCS1 | miR-29a/c-3p common target gene | Experiment supported target gene (TarBase v.8) |
| ENSG00000112977 | DAP | miR-29a-3p target gene only | Experiment supported target gene (TarBase v.8) |
| ENSG00000112984 | KIF20A | miR-29a/c-3p common target gene | Experiment supported target gene (TarBase v.8) |
| ENSG00000112992 | NNT | miR-29a-3p target gene only | Experiment supported target gene (TarBase v.8) |
| ENSG00000113070 | HBEGF | miR-29a/c-3p common target gene | Experiment supported target gene (TarBase v.8) |
| ENSG00000113083 | LOX | miR-29a/c-3p common target gene | Experiment supported target gene (TarBase v.8) |
| ENSG00000113140 | SPARC | miR-29a/c-3p common target gene | Experiment supported target gene (TarBase v.8) |
| ENSG00000113161 | HMGCR | miR-29a/c-3p common target gene | Experiment supported target gene (TarBase v.8) |

| | | | |
|---|---|---|---|
| ENSG00000113273 | ARSB | miR-29a/c-3p common target gene | Experiment supported target gene (TarBase v.8) |
| ENSG00000113300 | CNOT6 | miR-29a/c-3p common target gene | Theoretical prediction (microT-CDS algorithm) |
| ENSG00000113327 | GABRG2 | miR-29a/c-3p common target gene | Experiment supported target gene (TarBase v.8) |
| ENSG00000113356 | POLR3G | miR-29a-3p target gene only | Experiment supported target gene (TarBase v.8) |
| ENSG00000113368 | LMNB1 | miR-29a/c-3p common target gene | Experiment supported target gene (TarBase v.8) |
| ENSG00000113369 | ARRDC3 | miR-29a/c-3p common target gene | Experiment supported target gene (TarBase v.8) |
| ENSG00000113387 | SUB1 | miR-29a/c-3p common target gene | Experiment supported target gene (TarBase v.8) |
| ENSG00000113448 | PDE4D | miR-29a/c-3p common target gene | Experiment supported target gene (TarBase v.8) |
| ENSG00000113522 | RAD50 | miR-29c-3p target gene only | Experiment supported target gene (TarBase v.8) |
| ENSG00000113575 | PPP2CA | miR-29a-3p target gene only | Experiment supported target gene (TarBase v.8) |
| ENSG00000113580 | NR3C1 | miR-29a/c-3p common target gene | Experiment supported target gene (TarBase v.8) |
| ENSG00000113583 | C5orf15 | miR-29a/c-3p common target gene | Experiment supported target gene (TarBase v.8) |
| ENSG00000113594 | LIFR | miR-29a/c-3p common target gene | Experiment supported target gene (TarBase v.8) |
| ENSG00000113621 | TXNDC15 | miR-29c-3p target gene only | Experiment supported target gene (TarBase v.8) |
| ENSG00000113638 | TTC33 | miR-29c-3p target gene only | Experiment supported target gene (TarBase v.8) |
| ENSG00000113643 | RARS | miR-29a-3p target gene only | Experiment supported target gene (TarBase v.8) |
| ENSG00000113648 | H2AFY | miR-29a/c-3p common target gene | Experiment supported target gene (TarBase v.8) |
| ENSG00000113712 | CSNK1A1 | miR-29a-3p target gene only | Experiment supported target gene (TarBase v.8) |
| ENSG00000113721 | PDGFRB | miR-29a/c-3p common target gene | Experiment supported target gene (TarBase v.8) |
| ENSG00000113761 | ZNF346 | miR-29a-3p target gene only | Experiment supported target gene (TarBase v.8) |
| ENSG00000113838 | TBCCD1 | miR-29a/c-3p common target gene | Experiment supported target gene (TarBase v.8) |
| ENSG00000114013 | CD86 | miR-29c-3p target gene only | Experiment supported target gene (TarBase v.8) |
| ENSG00000114021 | NIT2 | miR-29a/c-3p common target gene | Theoretical prediction (microT-CDS algorithm) |
| ENSG00000114030 | KPNA1 | miR-29a/c-3p common target gene | Experiment supported target gene (TarBase v.8) |
| ENSG00000114098 | ARMC8 | miR-29a/c-3p common target gene | Theoretical prediction (microT-CDS algorithm) |
| ENSG00000114120 | SLC25A36 | miR-29a/c-3p common target gene | Experiment supported target gene (TarBase v.8) |
| ENSG00000114125 | RNF7 | miR-29a-3p target gene only | Experiment supported target gene (TarBase v.8) |
| ENSG00000114270 | COL7A1 | miR-29a/c-3p common target gene | Experiment supported target gene (TarBase v.8) |
| ENSG00000114353 | GNAI2 | miR-29a/c-3p common target gene | Theoretical prediction (microT-CDS algorithm) |
| ENSG00000114439 | BBX | miR-29a/c-3p common target gene | Experiment supported target gene (TarBase v.8) |
| ENSG00000114520 | SNX4 | miR-29a/c-3p common target gene | Experiment supported target gene (TarBase v.8) |
| ENSG00000114529 | C3orf52 | miR-29c-3p target gene only | Experiment supported target gene (TarBase v.8) |
| ENSG00000114544 | SLC41A3 | miR-29a-3p target gene only | Experiment supported target gene (TarBase v.8) |
| ENSG00000114554 | PLXNA1 | miR-29a/c-3p common target gene | Experiment supported target gene (TarBase v.8) |
| ENSG00000114573 | ATP6V1A | miR-29a/c-3p common target gene | Experiment supported target gene (TarBase v.8) |
| ENSG00000114638 | UPK1B | miR-29c-3p target gene only | Experiment supported target gene (TarBase v.8) |
| ENSG00000114650 | SCAP | miR-29a/c-3p common target gene | Experiment supported target gene (TarBase v.8) |
| ENSG00000114698 | PLSCR4 | miR-29a-3p target gene only | Experiment supported target gene (TarBase v.8) |
| ENSG00000114739 | ACVR2B | miR-29a/c-3p common target gene | Theoretical prediction (microT-CDS algorithm) |
| ENSG00000114742 | WDR48 | miR-29a/c-3p common target gene | Theoretical prediction (microT-CDS algorithm) |
| ENSG00000114744 | COMMD2 | miR-29a/c-3p common target gene | Experiment supported target gene (TarBase v.8) |
| ENSG00000114767 | RRP9 | miR-29c-3p target gene only | Experiment supported target gene (TarBase v.8) |
| ENSG00000114771 | AADAC | miR-29c-3p target gene only | Experiment supported target gene (TarBase v.8) |
| ENSG00000114784 | EIF1B | miR-29c-3p target gene only | Experiment supported target gene (TarBase v.8) |
| ENSG00000114796 | KLHL24 | miR-29a/c-3p common target gene | Theoretical prediction (microT-CDS algorithm) |
| ENSG00000114853 | ZBTB47 | miR-29a/c-3p common target gene | Experiment supported target gene (TarBase v.8) |
| ENSG00000114857 | NKTR | miR-29a-3p target gene only | Experiment supported target gene (TarBase v.8) |
| ENSG00000114861 | FOXP1 | miR-29a/c-3p common target gene | Experiment supported target gene (TarBase v.8) |
| ENSG00000114867 | EIF4G1 | miR-29a-3p target gene only | Experiment supported target gene (TarBase v.8) |
| ENSG00000114978 | MOB1A | miR-29a/c-3p common target gene | Experiment supported target gene (TarBase v.8) |
| ENSG00000115020 | PIKFYVE | miR-29a/c-3p common target gene | Experiment supported target gene (TarBase v.8) |
| ENSG00000115112 | TFCP2L1 | miR-29c-3p target gene only | Experiment supported target gene (TarBase v.8) |
| ENSG00000115129 | TP53I3 | miR-29c-3p target gene only | Experiment supported target gene (TarBase v.8) |
| ENSG00000115183 | TANC1 | miR-29a/c-3p common target gene | Theoretical prediction (microT-CDS algorithm) |
| ENSG00000115194 | SLC30A3 | miR-29a/c-3p common target gene | Experiment supported target gene (TarBase v.8) |
| ENSG00000115207 | GTF3C2 | miR-29a-3p target gene only | Experiment supported target gene (TarBase v.8) |
| ENSG00000115216 | NRBP1 | miR-29a/c-3p common target gene | Experiment supported target gene (TarBase v.8) |
| ENSG00000115234 | SNX17 | miR-29a-3p target gene only | Experiment supported target gene (TarBase v.8) |
| ENSG00000115267 | IFIH1 | miR-29c-3p target gene only | Experiment supported target gene (TarBase v.8) |
| ENSG00000115271 | GCA | miR-29c-3p target gene only | Experiment supported target gene (TarBase v.8) |
| ENSG00000115282 | TTC31 | miR-29a/c-3p common target gene | Theoretical prediction (microT-CDS algorithm) |
| ENSG00000115297 | TLX2 | miR-29a/c-3p common target gene | Theoretical prediction (microT-CDS algorithm) |
| ENSG00000115306 | SPTBN1 | miR-29a/c-3p common target gene | Experiment supported target gene (TarBase v.8) |
| ENSG00000115355 | CCDC88A | miR-29a/c-3p common target gene | Experiment supported target gene (TarBase v.8) |
| ENSG00000115363 | EVA1A | miR-29a/c-3p common target gene | Theoretical prediction (microT-CDS algorithm) |
| ENSG00000115365 | LANCL1 | miR-29c-3p target gene only | Experiment supported target gene (TarBase v.8) |
| ENSG00000115368 | WDR75 | miR-29a/c-3p common target gene | Theoretical prediction (microT-CDS algorithm) |
| ENSG00000115380 | EFEMP1 | miR-29c-3p target gene only | Experiment supported target gene (TarBase v.8) |
| ENSG00000115415 | STAT1 | miR-29c-3p target gene only | Experiment supported target gene (TarBase v.8) |
| ENSG00000115421 | PAPOLG | miR-29a/c-3p common target gene | Theoretical prediction (microT-CDS algorithm) |
| ENSG00000115464 | USP34 | miR-29a-3p target gene only | Experiment supported target gene (TarBase v.8) |
| ENSG00000115484 | CCT4 | miR-29a-3p target gene only | Experiment supported target gene (TarBase v.8) |
| ENSG00000115526 | CHST10 | miR-29a-3p target gene only | Experiment supported target gene (TarBase v.8) |
| ENSG00000115590 | IL1R2 | miR-29c-3p target gene only | Experiment supported target gene (TarBase v.8) |
| ENSG00000115594 | IL1R1 | miR-29a-3p target gene only | Experiment supported target gene (TarBase v.8) |
| ENSG00000115602 | IL1RL1 | miR-29a/c-3p common target gene | Theoretical prediction (microT-CDS algorithm) |
| ENSG00000115657 | ABCB6 | miR-29a/c-3p common target gene | Experiment supported target gene (TarBase v.8) |
| ENSG00000115808 | STRN | miR-29a/c-3p common target gene | Experiment supported target gene (TarBase v.8) |
| ENSG00000115884 | SDC1 | miR-29c-3p target gene only | Experiment supported target gene (TarBase v.8) |
| ENSG00000115904 | SOS1 | miR-29a/c-3p common target gene | Experiment supported target gene (TarBase v.8) |
| ENSG00000115944 | COX7A2L | miR-29a/c-3p common target gene | Experiment supported target gene (TarBase v.8) |
| ENSG00000115970 | THADA | miR-29a-3p target gene only | Experiment supported target gene (TarBase v.8) |
| ENSG00000115993 | TRAK2 | miR-29a/c-3p common target gene | Experiment supported target gene (TarBase v.8) |
| ENSG00000116044 | NFE2L2 | miR-29a-3p target gene only | Experiment supported target gene (TarBase v.8) |
| ENSG00000116062 | MSH6 | miR-29a/c-3p common target gene | Experiment supported target gene (TarBase v.8) |
| ENSG00000116095 | PLEKHA3 | miR-29a/c-3p common target gene | Theoretical prediction (microT-CDS algorithm) |
| ENSG00000116128 | BCL9 | miR-29a-3p target gene only | Experiment supported target gene (TarBase v.8) |
| ENSG00000116133 | DHCR24 | miR-29a-3p target gene only | Experiment supported target gene (TarBase v.8) |
| ENSG00000116157 | GPX7 | miR-29a/c-3p common target gene | Experiment supported target gene (TarBase v.8) |
| ENSG00000116285 | ERRFI1 | miR-29a-3p target gene only | Experiment supported target gene (TarBase v.8) |
| ENSG00000116406 | EDEM3 | miR-29c-3p target gene only | Experiment supported target gene (TarBase v.8) |
| ENSG00000116473 | RAP1A | miR-29a/c-3p common target gene | Theoretical prediction (microT-CDS algorithm) |
| ENSG00000116539 | ASH1L | miR-29a-3p target gene only | Experiment supported target gene (TarBase v.8) |
| ENSG00000116560 | SFPQ | miR-29a/c-3p common target gene | Experiment supported target gene (TarBase v.8) |
| ENSG00000116663 | FBXO6 | miR-29c-3p target gene only | Experiment supported target gene (TarBase v.8) |
| ENSG00000116667 | C1orf21 | miR-29a/c-3p common target gene | Theoretical prediction (microT-CDS algorithm) |
| ENSG00000116685 | KIAA2013 | miR-29a-3p target gene only | Experiment supported target gene (TarBase v.8) |
| ENSG00000116704 | SLC35D1 | miR-29a-3p target gene only | Experiment supported target gene (TarBase v.8) |
| ENSG00000116711 | PLA2G4A | miR-29a-3p target gene only | Experiment supported target gene (TarBase v.8) |
| ENSG00000116741 | RGS2 | miR-29c-3p target gene only | Experiment supported target gene (TarBase v.8) |
| ENSG00000116885 | OSCP1 | miR-29c-3p target gene only | Experiment supported target gene (TarBase v.8) |
| ENSG00000116954 | RRAGC | miR-29a/c-3p common target gene | Experiment supported target gene (TarBase v.8) |
| ENSG00000116962 | NID1 | miR-29a/c-3p common target gene | Theoretical prediction (microT-CDS algorithm) |
| ENSG00000116977 | LGALS8 | miR-29a/c-3p common target gene | Experiment supported target gene (TarBase v.8) |
| ENSG00000116983 | HPCAL4 | miR-29a/c-3p common target gene | Theoretical prediction (microT-CDS algorithm) |
| ENSG00000116984 | MTR | miR-29c-3p target gene only | Experiment supported target gene (TarBase v.8) |
| ENSG00000117000 | RLF | miR-29a/c-3p common target gene | Experiment supported target gene (TarBase v.8) |
| ENSG00000117016 | RIMS3 | miR-29c-3p target gene only | Experiment supported target gene (TarBase v.8) |
| ENSG00000117020 | AKT3 | miR-29a/c-3p common target gene | Experiment supported target gene (TarBase v.8) |
| ENSG00000117090 | SLAMF1 | miR-29a-3p target gene only | Theoretical prediction (microT-CDS algorithm) |
| ENSG00000117139 | KDM5B | miR-29a/c-3p common target gene | Experiment supported target gene (TarBase v.8) |
| ENSG00000117148 | ACTL8 | miR-29c-3p target gene only | Experiment supported target gene (TarBase v.8) |
| ENSG00000117152 | RGS4 | miR-29a/c-3p common target gene | Experiment supported target gene (TarBase v.8) |
| ENSG00000117154 | IGSF21 | miR-29c-3p target gene only | Experiment supported target gene (TarBase v.8) |
| ENSG00000117226 | GBP3 | miR-29c-3p target gene only | Experiment supported target gene (TarBase v.8) |
| ENSG00000117262 | GPR89A | miR-29a/c-3p common target gene | Theoretical prediction (microT-CDS algorithm) |
| ENSG00000117318 | ID3 | miR-29a/c-3p common target gene | Experiment supported target gene (TarBase v.8) |
| ENSG00000117335 | CD46 | miR-29a/c-3p common target gene | Experiment supported target gene (TarBase v.8) |
| ENSG00000117360 | PRPF3 | miR-29c-3p target gene only | Experiment supported target gene (TarBase v.8) |
| ENSG00000117362 | APH1A | miR-29a-3p target gene only | Experiment supported target gene (TarBase v.8) |
| ENSG00000117408 | IPO13 | miR-29c-3p target gene only | Experiment supported target gene (TarBase v.8) |
| ENSG00000117411 | B4GALT2 | miR-29c-3p target gene only | Experiment supported target gene (TarBase v.8) |
| ENSG00000117450 | PRDX1 | miR-29c-3p target gene only | Experiment supported target gene (TarBase v.8) |
| ENSG00000117461 | PIK3R3 | miR-29a-3p target gene only | Experiment supported target gene (TarBase v.8) |
| ENSG00000117472 | TSPAN1 | miR-29a/c-3p common target gene | Experiment supported target gene (TarBase v.8) |
| ENSG00000117479 | SLC19A2 | miR-29c-3p target gene only | Experiment supported target gene (TarBase v.8) |
| ENSG00000117505 | DR1 | miR-29a-3p target gene only | Experiment supported target gene (TarBase v.8) |
| ENSG00000117523 | PRRC2C | miR-29a/c-3p common target gene | Experiment supported target gene (TarBase v.8) |
| ENSG00000117525 | F3 | miR-29c-3p target gene only | Experiment supported target gene (TarBase v.8) |
| ENSG00000117543 | DPH5 | miR-29c-3p target gene only | Experiment supported target gene (TarBase v.8) |
| ENSG00000117569 | PTBP2 | miR-29a/c-3p common target gene | Theoretical prediction (microT-CDS algorithm) |
| ENSG00000117586 | TNFSF4 | miR-29a-3p target gene only | Experiment supported target gene (TarBase v.8) |
| ENSG00000117602 | RCAN3 | miR-29a-3p target gene only | Experiment supported target gene (TarBase v.8) |
| ENSG00000117758 | STX12 | miR-29a-3p target gene only | Experiment supported target gene (TarBase v.8) |
| ENSG00000118113 | MMP8 | miR-29a/c-3p common target gene | Theoretical prediction (microT-CDS algorithm) |
| ENSG00000118181 | RPS25 | miR-29c-3p target gene only | Experiment supported target gene (TarBase v.8) |
| ENSG00000118200 | CAMSAP2 | miR-29a/c-3p common target gene | Experiment supported target gene (TarBase v.8) |
| ENSG00000118298 | CA14 | miR-29c-3p target gene only | Experiment supported target gene (TarBase v.8) |
| ENSG00000118402 | ELOVL4 | miR-29a/c-3p common target gene | Experiment supported target gene (TarBase v.8) |
| ENSG00000118418 | HMGN3 | miR-29a/c-3p common target gene | Experiment supported target gene (TarBase v.8) |
| ENSG00000118432 | CNR1 | miR-29a-3p target gene only | Experiment supported target gene (TarBase v.8) |
| ENSG00000118454 | ANKRD13C | miR-29a/c-3p common target gene | Experiment supported target gene (TarBase v.8) |
| ENSG00000118503 | TNFAIP3 | miR-29a/c-3p common target gene | Experiment supported target gene (TarBase v.8) |
| ENSG00000118513 | MYB | miR-29c-3p target gene only | Experiment supported target gene (TarBase v.8) |
| ENSG00000118515 | SGK1 | miR-29a/c-3p common target gene | Experiment supported target gene (TarBase v.8) |
| ENSG00000118579 | MED28 | miR-29a/c-3p common target gene | Experiment supported target gene (TarBase v.8) |
| ENSG00000118689 | FOXO3 | miR-29a/c-3p common target gene | Theoretical prediction (microT-CDS algorithm) |
| ENSG00000118705 | RPN2 | miR-29a-3p target gene only | Experiment supported target gene (TarBase v.8) |
| ENSG00000118922 | KLF12 | miR-29a/c-3p common target gene | Experiment supported target gene (TarBase v.8) |
| ENSG00000118971 | CCND2 | miR-29a/c-3p common target gene | Experiment supported target gene (TarBase v.8) |
| ENSG00000118985 | ELL2 | miR-29c-3p target gene only | Experiment supported target gene (TarBase v.8) |
| ENSG00000119041 | GTF3C3 | miR-29a/c-3p common target gene | Theoretical prediction (microT-CDS algorithm) |
| ENSG00000119125 | GDA | miR-29a-3p target gene only | Experiment supported target gene (TarBase v.8) |
| ENSG00000119203 | CPSF3 | miR-29c-3p target gene only | Experiment supported target gene (TarBase v.8) |
| ENSG00000119280 | C1orf198 | miR-29a-3p target gene only | Experiment supported target gene (TarBase v.8) |
| ENSG00000119326 | CTNNAL1 | miR-29c-3p target gene only | Experiment supported target gene (TarBase v.8) |
| ENSG00000119335 | SET | miR-29a/c-3p common target gene | Theoretical prediction (microT-CDS algorithm) |
| ENSG00000119396 | RAB14 | miR-29a/c-3p common target gene | Theoretical prediction (microT-CDS algorithm) |
| ENSG00000119402 | FBXW2 | miR-29a/c-3p common target gene | Experiment supported target gene (TarBase v.8) |
| ENSG00000119408 | NEK6 | miR-29c-3p target gene only | Experiment supported target gene (TarBase v.8) |
| ENSG00000119414 | PPP6C | miR-29a/c-3p common target gene | Theoretical prediction (microT-CDS algorithm) |
| ENSG00000119638 | NEK9 | miR-29a/c-3p common target gene | Experiment supported target gene (TarBase v.8) |
| ENSG00000119686 | FLVCR2 | miR-29c-3p target gene only | Experiment supported target gene (TarBase v.8) |
| ENSG00000119772 | DNMT3A | miR-29a/c-3p common target gene | Experiment supported target gene (TarBase v.8) |
| ENSG00000119778 | ATAD2B | miR-29a/c-3p common target gene | Experiment supported target gene (TarBase v.8) |
| ENSG00000119866 | BCL11A | miR-29a/c-3p common target gene | Experiment supported target gene (TarBase v.8) |
| ENSG00000119900 | OGFRL1 | miR-29a/c-3p common target gene | Experiment supported target gene (TarBase v.8) |
| ENSG00000119917 | IFIT3 | miR-29c-3p target gene only | Experiment supported target gene (TarBase v.8) |
| ENSG00000119922 | IFIT2 | miR-29c-3p target gene only | Experiment supported target gene (TarBase v.8) |
| ENSG00000119929 | CUTC | miR-29c-3p target gene only | Experiment supported target gene (TarBase v.8) |
| ENSG00000119938 | PPP1R3C | miR-29c-3p target gene only | Experiment supported target gene (TarBase v.8) |
| ENSG00000120063 | GNA13 | miR-29a/c-3p common target gene | Experiment supported target gene (TarBase v.8) |
| ENSG00000120137 | PANK3 | miR-29a/c-3p common target gene | Experiment supported target gene (TarBase v.8) |
| ENSG00000120211 | INSL4 | miR-29c-3p target gene only | Experiment supported target gene (TarBase v.8) |
| ENSG00000120251 | GRIA2 | miR-29c-3p target gene only | Experiment supported target gene (TarBase v.8) |
| ENSG00000120256 | LRP11 | miR-29a/c-3p common target gene | Experiment supported target gene (TarBase v.8) |
| ENSG00000120318 | ARAP3 | miR-29c-3p target gene only | Experiment supported target gene (TarBase v.8) |
| ENSG00000120437 | ACAT2 | miR-29a-3p target gene only | Experiment supported target gene (TarBase v.8) |
| ENSG00000120438 | TCP1 | miR-29a-3p target gene only | Experiment supported target gene (TarBase v.8) |
| ENSG00000120526 | NUDCD1 | miR-29a/c-3p common target gene | Theoretical prediction (microT-CDS algorithm) |
| ENSG00000120594 | PLXDC2 | miR-29a/c-3p common target gene | Experiment supported target gene (TarBase v.8) |
| ENSG00000120616 | EPC1 | miR-29a/c-3p common target gene | Theoretical prediction (microT-CDS algorithm) |
| ENSG00000120656 | TAF12 | miR-29c-3p target gene only | Experiment supported target gene (TarBase v.8) |
| ENSG00000120685 | PROSER1 | miR-29a-3p target gene only | Theoretical prediction (microT-CDS algorithm) |
| ENSG00000120708 | TGFBI | miR-29c-3p target gene only | Experiment supported target gene (TarBase v.8) |
| ENSG00000120727 | PAIP2 | miR-29a-3p target gene only | Theoretical prediction (microT-CDS algorithm) |
| ENSG00000120742 | SERP1 | miR-29c-3p target gene only | Experiment supported target gene (TarBase v.8) |
| ENSG00000120802 | TMPO | miR-29a-3p target gene only | Experiment supported target gene (TarBase v.8) |
| ENSG00000120875 | DUSP4 | miR-29c-3p target gene only | Experiment supported target gene (TarBase v.8) |
| ENSG00000120889 | TNFRSF10B | miR-29a/c-3p common target gene | Experiment supported target gene (TarBase v.8) |
| ENSG00000120910 | PPP3CC | miR-29c-3p target gene only | Experiment supported target gene (TarBase v.8) |
| ENSG00000120992 | LYPLA1 | miR-29a/c-3p common target gene | Theoretical prediction (microT-CDS algorithm) |
| ENSG00000121005 | CRISPLD1 | miR-29a/c-3p common target gene | Experiment supported target gene (TarBase v.8) |
| ENSG00000121022 | COPS5 | miR-29c-3p target gene only | Experiment supported target gene (TarBase v.8) |
| ENSG00000121594 | CD80 | miR-29a/c-3p common target gene | Theoretical prediction (microT-CDS algorithm) |
| ENSG00000121716 | PILRB | miR-29a/c-3p common target gene | Experiment supported target gene (TarBase v.8) |
| ENSG00000121741 | ZMYM2 | miR-29a/c-3p common target gene | Experiment supported target gene (TarBase v.8) |
| ENSG00000121858 | TNFSF10 | miR-29c-3p target gene only | Experiment supported target gene (TarBase v.8) |
| ENSG00000121864 | ZNF639 | miR-29a/c-3p common target gene | Experiment supported target gene (TarBase v.8) |
| ENSG00000121989 | ACVR2A | miR-29a/c-3p common target gene | Theoretical prediction (microT-CDS algorithm) |
| ENSG00000122068 | FYTTD1 | miR-29c-3p target gene only | Experiment supported target gene (TarBase v.8) |
| ENSG00000122176 | FMOD | miR-29a-3p target gene only | Experiment supported target gene (TarBase v.8) |
| ENSG00000122203 | KIAA1191 | miR-29a-3p target gene only | Experiment supported target gene (TarBase v.8) |
| ENSG00000122218 | COPA | miR-29a-3p target gene only | Experiment supported target gene (TarBase v.8) |
| ENSG00000122482 | ZNF644 | miR-29a/c-3p common target gene | Theoretical prediction (microT-CDS algorithm) |
| ENSG00000122490 | PQLC1 | miR-29a/c-3p common target gene | Theoretical prediction (microT-CDS algorithm) |
| ENSG00000122591 | FAM126A | miR-29a/c-3p common target gene | Experiment supported target gene (TarBase v.8) |
| ENSG00000122641 | INHBA | miR-29c-3p target gene only | Experiment supported target gene (TarBase v.8) |
| ENSG00000122729 | ACO1 | miR-29a/c-3p common target gene | Experiment supported target gene (TarBase v.8) |
| ENSG00000122778 | KIAA1549 | miR-29a/c-3p common target gene | Experiment supported target gene (TarBase v.8) |
| ENSG00000122779 | TRIM24 | miR-29c-3p target gene only | Experiment supported target gene (TarBase v.8) |
| ENSG00000122873 | CISD1 | miR-29a-3p target gene only | Experiment supported target gene (TarBase v.8) |
| ENSG00000123066 | MED13L | miR-29a/c-3p common target gene | Experiment supported target gene (TarBase v.8) |
| ENSG00000123091 | RNF11 | miR-29a/c-3p common target gene | Experiment supported target gene (TarBase v.8) |
| ENSG00000123094 | RASSF8 | miR-29a/c-3p common target gene | Experiment supported target gene (TarBase v.8) |
| ENSG00000123095 | BHLHE41 | miR-29c-3p target gene only | Experiment supported target gene (TarBase v.8) |
| ENSG00000123219 | CENPK | miR-29a/c-3p common target gene | Experiment supported target gene (TarBase v.8) |
| ENSG00000123243 | ITIH5 | miR-29a-3p target gene only | Experiment supported target gene (TarBase v.8) |
| ENSG00000123364 | HOXC13 | miR-29c-3p target gene only | Experiment supported target gene (TarBase v.8) |
| ENSG00000123405 | NFE2 | miR-29c-3p target gene only | Experiment supported target gene (TarBase v.8) |
| ENSG00000123500 | COL10A1 | miR-29a/c-3p common target gene | Experiment supported target gene (TarBase v.8) |
| ENSG00000123545 | NDUFAF4 | miR-29c-3p target gene only | Experiment supported target gene (TarBase v.8) |
| ENSG00000123560 | PLP1 | miR-29a/c-3p common target gene | Theoretical prediction (microT-CDS algorithm) |
| ENSG00000123562 | MORF4L2 | miR-29a/c-3p common target gene | Experiment supported target gene (TarBase v.8) |
| ENSG00000123609 | NMI | miR-29a/c-3p common target gene | Experiment supported target gene (TarBase v.8) |
| ENSG00000123610 | TNFAIP6 | miR-29a-3p target gene only | Experiment supported target gene (TarBase v.8) |
| ENSG00000123612 | ACVR1C | miR-29c-3p target gene only | Experiment supported target gene (TarBase v.8) |
| ENSG00000123643 | SLC36A1 | miR-29a/c-3p common target gene | Theoretical prediction (microT-CDS algorithm) |
| ENSG00000123684 | LPGAT1 | miR-29c-3p target gene only | Experiment supported target gene (TarBase v.8) |
| ENSG00000123685 | BATF3 | miR-29c-3p target gene only | Experiment supported target gene (TarBase v.8) |
| ENSG00000123689 | G0S2 | miR-29c-3p target gene only | Experiment supported target gene (TarBase v.8) |
| ENSG00000123700 | KCNJ2 | miR-29a/c-3p common target gene | Experiment supported target gene (TarBase v.8) |
| ENSG00000123810 | B9D2 | miR-29c-3p target gene only | Experiment supported target gene (TarBase v.8) |
| ENSG00000123838 | C4BPA | miR-29c-3p target gene only | Experiment supported target gene (TarBase v.8) |
| ENSG00000123843 | C4BPB | miR-29c-3p target gene only | Experiment supported target gene (TarBase v.8) |
| ENSG00000123989 | CHPF | miR-29c-3p target gene only | Experiment supported target gene (TarBase v.8) |
| ENSG00000123992 | DNPEP | miR-29a-3p target gene only | Experiment supported target gene (TarBase v.8) |
| ENSG00000124006 | OBSL1 | miR-29a/c-3p common target gene | Experiment supported target gene (TarBase v.8) |
| ENSG00000124116 | WFDC3 | miR-29c-3p target gene only | Experiment supported target gene (TarBase v.8) |
| ENSG00000124120 | TTPAL | miR-29c-3p target gene only | Experiment supported target gene (TarBase v.8) |
| ENSG00000124151 | NCOA3 | miR-29a/c-3p common target gene | Experiment supported target gene (TarBase v.8) |
| ENSG00000124193 | SRSF6 | miR-29a-3p target gene only | Experiment supported target gene (TarBase v.8) |
| ENSG00000124198 | ARFGEF2 | miR-29c-3p target gene only | Experiment supported target gene (TarBase v.8) |
| ENSG00000124201 | ZNFX1 | miR-29c-3p target gene only | Experiment supported target gene (TarBase v.8) |
| ENSG00000124222 | STX16 | miR-29a/c-3p common target gene | Experiment supported target gene (TarBase v.8) |
| ENSG00000124299 | PEPD | miR-29a/c-3p common target gene | Experiment supported target gene (TarBase v.8) |
| ENSG00000124333 | VAMP7 | miR-29c-3p target gene only | Experiment supported target gene (TarBase v.8) |
| ENSG00000124440 | HIF3A | miR-29a/c-3p common target gene | Theoretical prediction (microT-CDS algorithm) |

| | | | |
|---|---|---|---|
| ENSG00000124486 | USP9X | miR-29a-3p target gene only | Experiment supported target gene (TarBase v.8) |
| ENSG00000124570 | SERPINB6 | miR-29a/c-3p common target gene | Experiment supported target gene (TarBase v.8) |
| ENSG00000124571 | XPO5 | miR-29a/c-3p common target gene | Experiment supported target gene (TarBase v.8) |
| ENSG00000124588 | NQO2 | miR-29c-3p target gene only | Experiment supported target gene (TarBase v.8) |
| ENSG00000124641 | MED20 | miR-29c-3p target gene only | Experiment supported target gene (TarBase v.8) |
| ENSG00000124664 | SPDEF | miR-29c-3p target gene only | Experiment supported target gene (TarBase v.8) |
| ENSG00000124702 | KLHDC3 | miR-29a/c-3p common target gene | Experiment supported target gene (TarBase v.8) |
| ENSG00000124749 | COL21A1 | miR-29a/c-3p common target gene | Experiment supported target gene (TarBase v.8) |
| ENSG00000124762 | CDKN1A | miR-29a/c-3p common target gene | Experiment supported target gene (TarBase v.8) |
| ENSG00000124766 | SOX4 | miR-29a/c-3p common target gene | Experiment supported target gene (TarBase v.8) |
| ENSG00000124767 | GLO1 | miR-29a/c-3p common target gene | Experiment supported target gene (TarBase v.8) |
| ENSG00000124782 | RREB1 | miR-29a/c-3p common target gene | Experiment supported target gene (TarBase v.8) |
| ENSG00000124788 | ATXN1 | miR-29a/c-3p common target gene | Experiment supported target gene (TarBase v.8) |
| ENSG00000124942 | AHNAK | miR-29a/c-3p common target gene | Experiment supported target gene (TarBase v.8) |
| ENSG00000125037 | EMC3 | miR-29a-3p target gene only | Theoretical prediction (microT-CDS algorithm) |
| ENSG00000125089 | SH3TC1 | miR-29a-3p target gene only | Experiment supported target gene (TarBase v.8) |
| ENSG00000125107 | CNOT1 | miR-29a/c-3p common target gene | Experiment supported target gene (TarBase v.8) |
| ENSG00000125249 | RAP2A | miR-29c-3p target gene only | Experiment supported target gene (TarBase v.8) |
| ENSG00000125266 | EFNB2 | miR-29a-3p target gene only | Experiment supported target gene (TarBase v.8) |
| ENSG00000125384 | PTGER2 | miR-29a/c-3p common target gene | Experiment supported target gene (TarBase v.8) |
| ENSG00000125430 | HS3ST3B1 | miR-29a/c-3p common target gene | Theoretical prediction (microT-CDS algorithm) |
| ENSG00000125484 | GTF3C4 | miR-29a/c-3p common target gene | Theoretical prediction (microT-CDS algorithm) |
| ENSG00000125772 | GPCPD1 | miR-29a/c-3p common target gene | Experiment supported target gene (TarBase v.8) |
| ENSG00000125779 | PANK2 | miR-29a-3p target gene only | Experiment supported target gene (TarBase v.8) |
| ENSG00000125810 | CD93 | miR-29a/c-3p common target gene | Experiment supported target gene (TarBase v.8) |
| ENSG00000125814 | NAPB | miR-29a/c-3p common target gene | Theoretical prediction (microT-CDS algorithm) |
| ENSG00000125817 | CENPB | miR-29a/c-3p common target gene | Theoretical prediction (microT-CDS algorithm) |
| ENSG00000125818 | PSMF1 | miR-29a/c-3p common target gene | Experiment supported target gene (TarBase v.8) |
| ENSG00000125826 | RBCK1 | miR-29a/c-3p common target gene | Experiment supported target gene (TarBase v.8) |
| ENSG00000125843 | AP5S1 | miR-29a/c-3p common target gene | Experiment supported target gene (TarBase v.8) |
| ENSG00000125845 | BMP2 | miR-29a-3p target gene only | Experiment supported target gene (TarBase v.8) |
| ENSG00000125848 | FLRT3 | miR-29c-3p target gene only | Experiment supported target gene (TarBase v.8) |
| ENSG00000125870 | SNRPB2 | miR-29c-3p target gene only | Experiment supported target gene (TarBase v.8) |
| ENSG00000125966 | MMP24 | miR-29a/c-3p common target gene | Experiment supported target gene (TarBase v.8) |
| ENSG00000125977 | EIF2S2 | miR-29a/c-3p common target gene | Experiment supported target gene (TarBase v.8) |
| ENSG00000126003 | PLAGL2 | miR-29c-3p target gene only | Experiment supported target gene (TarBase v.8) |
| ENSG00000126016 | AMOT | miR-29a/c-3p common target gene | Experiment supported target gene (TarBase v.8) |
| ENSG00000126070 | AGO3 | miR-29a/c-3p common target gene | Experiment supported target gene (TarBase v.8) |
| ENSG00000126216 | TUBGCP3 | miR-29a/c-3p common target gene | Experiment supported target gene (TarBase v.8) |
| ENSG00000126218 | F10 | miR-29a-3p target gene only | Experiment supported target gene (TarBase v.8) |
| ENSG00000126351 | THRA | miR-29a/c-3p common target gene | Theoretical prediction (microT-CDS algorithm) |
| ENSG00000126457 | PRMT1 | miR-29a/c-3p common target gene | Experiment supported target gene (TarBase v.8) |
| ENSG00000126464 | PRR12 | miR-29a/c-3p common target gene | Experiment supported target gene (TarBase v.8) |
| ENSG00000126549 | STATH | miR-29c-3p target gene only | Experiment supported target gene (TarBase v.8) |
| ENSG00000126603 | GLIS2 | miR-29a/c-3p common target gene | Theoretical prediction (microT-CDS algorithm) |
| ENSG00000126778 | SIX1 | miR-29c-3p target gene only | Experiment supported target gene (TarBase v.8) |
| ENSG00000126858 | RHOT1 | miR-29a/c-3p common target gene | Theoretical prediction (microT-CDS algorithm) |
| ENSG00000126882 | FAM78A | miR-29a-3p target gene only | Experiment supported target gene (TarBase v.8) |
| ENSG00000126970 | ZC4H2 | miR-29a/c-3p common target gene | Theoretical prediction (microT-CDS algorithm) |
| ENSG00000127022 | CANX | miR-29a/c-3p common target gene | Experiment supported target gene (TarBase v.8) |
| ENSG00000127124 | HIVEP3 | miR-29a/c-3p common target gene | Experiment supported target gene (TarBase v.8) |
| ENSG00000127129 | EDN2 | miR-29c-3p target gene only | Experiment supported target gene (TarBase v.8) |
| ENSG00000127152 | BCL11B | miR-29a/c-3p common target gene | Theoretical prediction (microT-CDS algorithm) |
| ENSG00000127220 | ABHD8 | miR-29c-3p target gene only | Experiment supported target gene (TarBase v.8) |
| ENSG00000127324 | TSPAN8 | miR-29a/c-3p common target gene | Experiment supported target gene (TarBase v.8) |
| ENSG00000127334 | DYRK2 | miR-29a/c-3p common target gene | Experiment supported target gene (TarBase v.8) |
| ENSG00000127423 | AUNIP | miR-29a/c-3p common target gene | Experiment supported target gene (TarBase v.8) |
| ENSG00000127481 | UBR4 | miR-29a-3p target gene only | Experiment supported target gene (TarBase v.8) |
| ENSG00000127483 | HP1BP3 | miR-29a/c-3p common target gene | Experiment supported target gene (TarBase v.8) |
| ENSG00000127527 | EPS15L1 | miR-29a/c-3p common target gene | Experiment supported target gene (TarBase v.8) |
| ENSG00000127528 | KLF2 | miR-29a/c-3p common target gene | Experiment supported target gene (TarBase v.8) |
| ENSG00000127529 | OR7C2 | miR-29c-3p target gene only | Experiment supported target gene (TarBase v.8) |
| ENSG00000127603 | MACF1 | miR-29a/c-3p common target gene | Experiment supported target gene (TarBase v.8) |
| ENSG00000127663 | KDM4B | miR-29a/c-3p common target gene | Theoretical prediction (microT-CDS algorithm) |
| ENSG00000127946 | HIP1 | miR-29a-3p target gene only | Experiment supported target gene (TarBase v.8) |
| ENSG00000127947 | PTPN12 | miR-29a-3p target gene only | Experiment supported target gene (TarBase v.8) |
| ENSG00000127990 | SGCE | miR-29a-3p target gene only | Experiment supported target gene (TarBase v.8) |
| ENSG00000128016 | ZFP36 | miR-29a/c-3p common target gene | Experiment supported target gene (TarBase v.8) |
| ENSG00000128228 | SDF2L1 | miR-29a-3p target gene only | Experiment supported target gene (TarBase v.8) |
| ENSG00000128242 | GAL3ST1 | miR-29c-3p target gene only | Experiment supported target gene (TarBase v.8) |
| ENSG00000128245 | YWHAH | miR-29a/c-3p common target gene | Experiment supported target gene (TarBase v.8) |
| ENSG00000128274 | A4GALT | miR-29c-3p target gene only | Experiment supported target gene (TarBase v.8) |
| ENSG00000128342 | LIF | miR-29a/c-3p common target gene | Experiment supported target gene (TarBase v.8) |
| ENSG00000128585 | MKLN1 | miR-29a/c-3p common target gene | Experiment supported target gene (TarBase v.8) |
| ENSG00000128590 | DNAJB9 | miR-29c-3p target gene only | Experiment supported target gene (TarBase v.8) |
| ENSG00000128595 | CALU | miR-29a/c-3p common target gene | Experiment supported target gene (TarBase v.8) |
| ENSG00000128607 | KLHDC10 | miR-29a/c-3p common target gene | Experiment supported target gene (TarBase v.8) |
| ENSG00000128645 | HOXD1 | miR-29a/c-3p common target gene | Experiment supported target gene (TarBase v.8) |
| ENSG00000128708 | HAT1 | miR-29a/c-3p common target gene | Experiment supported target gene (TarBase v.8) |
| ENSG00000128805 | ARHGAP22 | miR-29c-3p target gene only | Experiment supported target gene (TarBase v.8) |
| ENSG00000128886 | ELL3 | miR-29c-3p target gene only | Experiment supported target gene (TarBase v.8) |
| ENSG00000129003 | VPS13C | miR-29a/c-3p common target gene | Experiment supported target gene (TarBase v.8) |
| ENSG00000129009 | ISLR | miR-29a/c-3p common target gene | Experiment supported target gene (TarBase v.8) |
| ENSG00000129038 | LOXL1 | miR-29c-3p target gene only | Experiment supported target gene (TarBase v.8) |
| ENSG00000129055 | ANAPC13 | miR-29c-3p target gene only | Experiment supported target gene (TarBase v.8) |
| ENSG00000129071 | MBD4 | miR-29a/c-3p common target gene | Experiment supported target gene (TarBase v.8) |
| ENSG00000129195 | FAM64A | miR-29c-3p target gene only | Experiment supported target gene (TarBase v.8) |
| ENSG00000129451 | KLK10 | miR-29c-3p target gene only | Experiment supported target gene (TarBase v.8) |
| ENSG00000129472 | RAB2B | miR-29c-3p target gene only | Experiment supported target gene (TarBase v.8) |
| ENSG00000129473 | BCL2L2 | miR-29a/c-3p common target gene | Experiment supported target gene (TarBase v.8) |
| ENSG00000129757 | CDKN1C | miR-29c-3p target gene only | Experiment supported target gene (TarBase v.8) |
| ENSG00000129925 | TMEM8A | miR-29a-3p target gene only | Experiment supported target gene (TarBase v.8) |
| ENSG00000129991 | TNNI3 | miR-29c-3p target gene only | Experiment supported target gene (TarBase v.8) |
| ENSG00000130005 | GAMT | miR-29c-3p target gene only | Experiment supported target gene (TarBase v.8) |
| ENSG00000130023 | ERMARD | miR-29c-3p target gene only | Experiment supported target gene (TarBase v.8) |
| ENSG00000130024 | PHF10 | miR-29c-3p target gene only | Experiment supported target gene (TarBase v.8) |
| ENSG00000130052 | STARD8 | miR-29a/c-3p common target gene | Theoretical prediction (microT-CDS algorithm) |
| ENSG00000130066 | SAT1 | miR-29a-3p target gene only | Experiment supported target gene (TarBase v.8) |
| ENSG00000130150 | MOSPD2 | miR-29a/c-3p common target gene | Theoretical prediction (microT-CDS algorithm) |
| ENSG00000130158 | DOCK6 | miR-29a-3p target gene only | Experiment supported target gene (TarBase v.8) |
| ENSG00000130165 | ELOF1 | miR-29a/c-3p common target gene | Theoretical prediction (microT-CDS algorithm) |
| ENSG00000130176 | CNN1 | miR-29a-3p target gene only | Experiment supported target gene (TarBase v.8) |
| ENSG00000130255 | RPL36 | miR-29a/c-3p common target gene | Experiment supported target gene (TarBase v.8) |
| ENSG00000130303 | BST2 | miR-29a/c-3p common target gene | Experiment supported target gene (TarBase v.8) |
| ENSG00000130309 | COLGALT1 | miR-29c-3p target gene only | Experiment supported target gene (TarBase v.8) |
| ENSG00000130311 | DDA1 | miR-29a-3p target gene only | Experiment supported target gene (TarBase v.8) |
| ENSG00000130338 | TULP4 | miR-29a/c-3p common target gene | Experiment supported target gene (TarBase v.8) |
| ENSG00000130475 | FCHO1 | miR-29c-3p target gene only | Experiment supported target gene (TarBase v.8) |
| ENSG00000130477 | UNC13A | miR-29a/c-3p common target gene | Experiment supported target gene (TarBase v.8) |
| ENSG00000130487 | KLHDC7B | miR-29c-3p target gene only | Experiment supported target gene (TarBase v.8) |
| ENSG00000130508 | PXDN | miR-29a/c-3p common target gene | Experiment supported target gene (TarBase v.8) |
| ENSG00000130540 | SULT4A1 | miR-29c-3p target gene only | Experiment supported target gene (TarBase v.8) |
| ENSG00000130544 | ZNF557 | miR-29c-3p target gene only | Experiment supported target gene (TarBase v.8) |
| ENSG00000130584 | ZBTB46 | miR-29a/c-3p common target gene | Theoretical prediction (microT-CDS algorithm) |
| ENSG00000130635 | COL5A1 | miR-29a/c-3p common target gene | Experiment supported target gene (TarBase v.8) |
| ENSG00000130702 | LAMA5 | miR-29a-3p target gene only | Experiment supported target gene (TarBase v.8) |
| ENSG00000130766 | SESN2 | miR-29c-3p target gene only | Experiment supported target gene (TarBase v.8) |
| ENSG00000130775 | THEMIS2 | miR-29a/c-3p common target gene | Experiment supported target gene (TarBase v.8) |
| ENSG00000130813 | C19orf66 | miR-29c-3p target gene only | Experiment supported target gene (TarBase v.8) |
| ENSG00000130881 | LRP3 | miR-29c-3p target gene only | Experiment supported target gene (TarBase v.8) |
| ENSG00000130935 | NOL11 | miR-29a/c-3p common target gene | Experiment supported target gene (TarBase v.8) |
| ENSG00000130985 | UBA1 | miR-29a/c-3p common target gene | Experiment supported target gene (TarBase v.8) |
| ENSG00000131015 | ULBP2 | miR-29a/c-3p common target gene | Experiment supported target gene (TarBase v.8) |
| ENSG00000131023 | LATS1 | miR-29a/c-3p common target gene | Experiment supported target gene (TarBase v.8) |
| ENSG00000131043 | AAR2 | miR-29a/c-3p common target gene | Experiment supported target gene (TarBase v.8) |
| ENSG00000131127 | ZNF141 | miR-29a/c-3p common target gene | Theoretical prediction (microT-CDS algorithm) |
| ENSG00000131148 | EMC8 | miR-29a/c-3p common target gene | Experiment supported target gene (TarBase v.8) |
| ENSG00000131153 | GINS2 | miR-29c-3p target gene only | Experiment supported target gene (TarBase v.8) |
| ENSG00000131187 | F12 | miR-29c-3p target gene only | Experiment supported target gene (TarBase v.8) |
| ENSG00000131269 | ABCB7 | miR-29a/c-3p common target gene | Experiment supported target gene (TarBase v.8) |
| ENSG00000131323 | TRAF3 | miR-29a/c-3p common target gene | Theoretical prediction (microT-CDS algorithm) |
| ENSG00000131351 | HAUS8 | miR-29a/c-3p common target gene | Experiment supported target gene (TarBase v.8) |
| ENSG00000131370 | SH3BP5 | miR-29a-3p target gene only | Experiment supported target gene (TarBase v.8) |
| ENSG00000131373 | HACL1 | miR-29a/c-3p common target gene | Experiment supported target gene (TarBase v.8) |
| ENSG00000131398 | KCNC3 | miR-29a/c-3p common target gene | Theoretical prediction (microT-CDS algorithm) |
| ENSG00000131459 | GFPT2 | miR-29c-3p target gene only | Experiment supported target gene (TarBase v.8) |
| ENSG00000131504 | DIAPH1 | miR-29a-3p target gene only | Experiment supported target gene (TarBase v.8) |
| ENSG00000131620 | ANO1 | miR-29a-3p target gene only | Experiment supported target gene (TarBase v.8) |
| ENSG00000131650 | KREMEN2 | miR-29a/c-3p common target gene | Experiment supported target gene (TarBase v.8) |
| ENSG00000131697 | NPHP4 | miR-29a/c-3p common target gene | Experiment supported target gene (TarBase v.8) |
| ENSG00000131711 | MAP1B | miR-29a/c-3p common target gene | Experiment supported target gene (TarBase v.8) |
| ENSG00000131747 | TOP2A | miR-29a/c-3p common target gene | Experiment supported target gene (TarBase v.8) |
| ENSG00000131748 | STARD3 | miR-29a/c-3p common target gene | Theoretical prediction (microT-CDS algorithm) |
| ENSG00000131771 | PPP1R1B | miR-29c-3p target gene only | Experiment supported target gene (TarBase v.8) |
| ENSG00000131779 | PEX11B | miR-29c-3p target gene only | Experiment supported target gene (TarBase v.8) |
| ENSG00000131791 | PRKAB2 | miR-29a/c-3p common target gene | Experiment supported target gene (TarBase v.8) |
| ENSG00000131844 | MCCC2 | miR-29a/c-3p common target gene | Experiment supported target gene (TarBase v.8) |
| ENSG00000131864 | USP29 | miR-29a/c-3p common target gene | Theoretical prediction (microT-CDS algorithm) |
| ENSG00000131873 | CHSY1 | miR-29a/c-3p common target gene | Experiment supported target gene (TarBase v.8) |
| ENSG00000131941 | RHPN2 | miR-29c-3p target gene only | Experiment supported target gene (TarBase v.8) |
| ENSG00000131966 | ACTR10 | miR-29a/c-3p common target gene | Experiment supported target gene (TarBase v.8) |
| ENSG00000132004 | FBXW9 | miR-29c-3p target gene only | Experiment supported target gene (TarBase v.8) |
| ENSG00000132016 | C19orf57 | miR-29c-3p target gene only | Experiment supported target gene (TarBase v.8) |
| ENSG00000132109 | TRIM21 | miR-29c-3p target gene only | Experiment supported target gene (TarBase v.8) |
| ENSG00000132356 | PRKAA1 | miR-29a/c-3p common target gene | Experiment supported target gene (TarBase v.8) |
| ENSG00000132383 | RPA1 | miR-29c-3p target gene only | Experiment supported target gene (TarBase v.8) |
| ENSG00000132394 | EEFSEC | miR-29a/c-3p common target gene | Experiment supported target gene (TarBase v.8) |
| ENSG00000132432 | SEC61G | miR-29a-3p target gene only | Experiment supported target gene (TarBase v.8) |
| ENSG00000132466 | ANKRD17 | miR-29a/c-3p common target gene | Experiment supported target gene (TarBase v.8) |
| ENSG00000132475 | H3F3B | miR-29a-3p target gene only | Experiment supported target gene (TarBase v.8) |
| ENSG00000132507 | EIF5A | miR-29a/c-3p common target gene | Experiment supported target gene (TarBase v.8) |
| ENSG00000132510 | KDM6B | miR-29a/c-3p common target gene | Experiment supported target gene (TarBase v.8) |
| ENSG00000132530 | XAF1 | miR-29c-3p target gene only | Experiment supported target gene (TarBase v.8) |
| ENSG00000132549 | VPS13B | miR-29a-3p target gene only | Experiment supported target gene (TarBase v.8) |
| ENSG00000132613 | MTSS1L | miR-29a/c-3p common target gene | Experiment supported target gene (TarBase v.8) |
| ENSG00000132622 | HSPA12B | miR-29a-3p target gene only | Experiment supported target gene (TarBase v.8) |
| ENSG00000132623 | ANKEF1 | miR-29a/c-3p common target gene | Experiment supported target gene (TarBase v.8) |
| ENSG00000132640 | BTBD3 | miR-29a/c-3p common target gene | Experiment supported target gene (TarBase v.8) |
| ENSG00000132718 | SYT11 | miR-29a-3p target gene only | Experiment supported target gene (TarBase v.8) |
| ENSG00000132780 | NASP | miR-29a/c-3p common target gene | Experiment supported target gene (TarBase v.8) |
| ENSG00000132824 | SERINC3 | miR-29a-3p target gene only | Experiment supported target gene (TarBase v.8) |
| ENSG00000132906 | CASP9 | miR-29a-3p target gene only | Experiment supported target gene (TarBase v.8) |
| ENSG00000132953 | XPO4 | miR-29a/c-3p common target gene | Theoretical prediction (microT-CDS algorithm) |
| ENSG00000132970 | WASF3 | miR-29c-3p target gene only | Experiment supported target gene (TarBase v.8) |
| ENSG00000133030 | MPRIP | miR-29a-3p target gene only | Experiment supported target gene (TarBase v.8) |
| ENSG00000133110 | POSTN | miR-29a/c-3p common target gene | Experiment supported target gene (TarBase v.8) |
| ENSG00000133111 | RFXAP | miR-29a/c-3p common target gene | Experiment supported target gene (TarBase v.8) |
| ENSG00000133135 | RNF128 | miR-29a/c-3p common target gene | Experiment supported target gene (TarBase v.8) |
| ENSG00000133138 | TBC1D8B | miR-29a-3p target gene only | Experiment supported target gene (TarBase v.8) |
| ENSG00000133142 | TCEAL4 | miR-29c-3p target gene only | Experiment supported target gene (TarBase v.8) |
| ENSG00000133466 | C1QTNF6 | miR-29a/c-3p common target gene | Experiment supported target gene (TarBase v.8) |
| ENSG00000133657 | ATP13A3 | miR-29a/c-3p common target gene | Experiment supported target gene (TarBase v.8) |
| ENSG00000133678 | TMEM254 | miR-29c-3p target gene only | Experiment supported target gene (TarBase v.8) |
| ENSG00000133706 | LARS | miR-29a/c-3p common target gene | Experiment supported target gene (TarBase v.8) |
| ENSG00000133789 | SWAP70 | miR-29a/c-3p common target gene | Experiment supported target gene (TarBase v.8) |
| ENSG00000133805 | AMPD3 | miR-29a/c-3p common target gene | Experiment supported target gene (TarBase v.8) |
| ENSG00000133818 | RRAS2 | miR-29a/c-3p common target gene | Theoretical prediction (microT-CDS algorithm) |
| ENSG00000133874 | RNF122 | miR-29a/c-3p common target gene | Experiment supported target gene (TarBase v.8) |
| ENSG00000133943 | C14orf159 | miR-29a/c-3p common target gene | Experiment supported target gene (TarBase v.8) |
| ENSG00000133985 | TTC9 | miR-29a/c-3p common target gene | Theoretical prediction (microT-CDS algorithm) |
| ENSG00000134001 | EIF2S1 | miR-29a/c-3p common target gene | Experiment supported target gene (TarBase v.8) |
| ENSG00000134013 | LOXL2 | miR-29a/c-3p common target gene | Experiment supported target gene (TarBase v.8) |
| ENSG00000134028 | ADAMDEC1 | miR-29a/c-3p common target gene | Theoretical prediction (microT-CDS algorithm) |
| ENSG00000134077 | THUMPD3 | miR-29a-3p target gene only | Experiment supported target gene (TarBase v.8) |
| ENSG00000134109 | EDEM1 | miR-29a-3p target gene only | Experiment supported target gene (TarBase v.8) |
| ENSG00000134121 | CHL1 | miR-29a-3p target gene only | Experiment supported target gene (TarBase v.8) |
| ENSG00000134240 | HMGCS2 | miR-29a/c-3p common target gene | Theoretical prediction (microT-CDS algorithm) |
| ENSG00000134250 | NOTCH2 | miR-29a/c-3p common target gene | Experiment supported target gene (TarBase v.8) |
| ENSG00000134294 | SLC38A2 | miR-29a-3p target gene only | Experiment supported target gene (TarBase v.8) |
| ENSG00000134313 | KIDINS220 | miR-29a/c-3p common target gene | Theoretical prediction (microT-CDS algorithm) |
| ENSG00000134321 | RSAD2 | miR-29c-3p target gene only | Experiment supported target gene (TarBase v.8) |
| ENSG00000134323 | MYCN | miR-29a/c-3p common target gene | Experiment supported target gene (TarBase v.8) |
| ENSG00000134326 | CMPK2 | miR-29c-3p target gene only | Experiment supported target gene (TarBase v.8) |
| ENSG00000134369 | NAV1 | miR-29a/c-3p common target gene | Experiment supported target gene (TarBase v.8) |
| ENSG00000134419 | RPS15A | miR-29a/c-3p common target gene | Experiment supported target gene (TarBase v.8) |
| ENSG00000134440 | NARS | miR-29a/c-3p common target gene | Experiment supported target gene (TarBase v.8) |
| ENSG00000134460 | IL2RA | miR-29c-3p target gene only | Theoretical prediction (microT-CDS algorithm) |
| ENSG00000134531 | EMP1 | miR-29a/c-3p common target gene | Experiment supported target gene (TarBase v.8) |
| ENSG00000134574 | DDB2 | miR-29c-3p target gene only | Experiment supported target gene (TarBase v.8) |
| ENSG00000134684 | YARS | miR-29a/c-3p common target gene | Experiment supported target gene (TarBase v.8) |
| ENSG00000134716 | CYP2J2 | miR-29c-3p target gene only | Experiment supported target gene (TarBase v.8) |
| ENSG00000134744 | ZCCHC11 | miR-29a/c-3p common target gene | Experiment supported target gene (TarBase v.8) |
| ENSG00000134748 | PRPF38A | miR-29c-3p target gene only | Experiment supported target gene (TarBase v.8) |
| ENSG00000134755 | DSC2 | miR-29c-3p target gene only | Experiment supported target gene (TarBase v.8) |
| ENSG00000134758 | RNF138 | miR-29a/c-3p common target gene | Experiment supported target gene (TarBase v.8) |
| ENSG00000134775 | FHOD3 | miR-29c-3p target gene only | Experiment supported target gene (TarBase v.8) |
| ENSG00000134825 | TMEM258 | miR-29a-3p target gene only | Experiment supported target gene (TarBase v.8) |
| ENSG00000134851 | TMEM165 | miR-29a-3p target gene only | Experiment supported target gene (TarBase v.8) |
| ENSG00000134852 | CLOCK | miR-29a/c-3p common target gene | Theoretical prediction (microT-CDS algorithm) |
| ENSG00000134864 | GGACT | miR-29c-3p target gene only | Experiment supported target gene (TarBase v.8) |
| ENSG00000134871 | COL4A2 | miR-29a/c-3p common target gene | Experiment supported target gene (TarBase v.8) |
| ENSG00000134874 | DZIP1 | miR-29c-3p target gene only | Experiment supported target gene (TarBase v.8) |
| ENSG00000134882 | UBAC2 | miR-29a-3p target gene only | Experiment supported target gene (TarBase v.8) |
| ENSG00000134901 | KDELC1 | miR-29a/c-3p common target gene | Experiment supported target gene (TarBase v.8) |
| ENSG00000134982 | APC | miR-29a/c-3p common target gene | Experiment supported target gene (TarBase v.8) |
| ENSG00000134986 | NREP | miR-29a/c-3p common target gene | Experiment supported target gene (TarBase v.8) |
| ENSG00000135048 | TMEM2 | miR-29a/c-3p common target gene | Experiment supported target gene (TarBase v.8) |
| ENSG00000135069 | PSAT1 | miR-29a-3p target gene only | Experiment supported target gene (TarBase v.8) |
| ENSG00000135074 | ADAM19 | miR-29a/c-3p common target gene | Experiment supported target gene (TarBase v.8) |
| ENSG00000135114 | OASL | miR-29c-3p target gene only | Experiment supported target gene (TarBase v.8) |
| ENSG00000135116 | HRK | miR-29a/c-3p common target gene | Experiment supported target gene (TarBase v.8) |
| ENSG00000135119 | RNFT2 | miR-29c-3p target gene only | Experiment supported target gene (TarBase v.8) |
| ENSG00000135205 | CCDC146 | miR-29a/c-3p common target gene | Theoretical prediction (microT-CDS algorithm) |
| ENSG00000135218 | CD36 | miR-29a-3p target gene only | Experiment supported target gene (TarBase v.8) |
| ENSG00000135241 | PNPLA8 | miR-29a/c-3p common target gene | Experiment supported target gene (TarBase v.8) |
| ENSG00000135250 | SRPK2 | miR-29c-3p target gene only | Experiment supported target gene (TarBase v.8) |
| ENSG00000135316 | SYNCRIP | miR-29a/c-3p common target gene | Theoretical prediction (microT-CDS algorithm) |
| ENSG00000135387 | CAPRIN1 | miR-29a/c-3p common target gene | Experiment supported target gene (TarBase v.8) |
| ENSG00000135390 | ATP5G2 | miR-29a/c-3p common target gene | Experiment supported target gene (TarBase v.8) |
| ENSG00000135423 | GLS2 | miR-29c-3p target gene only | Experiment supported target gene (TarBase v.8) |
| ENSG00000135426 | TESPA1 | miR-29a/c-3p common target gene | Theoretical prediction (microT-CDS algorithm) |
| ENSG00000135457 | TFCP2 | miR-29a/c-3p common target gene | Experiment supported target gene (TarBase v.8) |
| ENSG00000135473 | PAN2 | miR-29a/c-3p common target gene | Experiment supported target gene (TarBase v.8) |
| ENSG00000135486 | HNRNPA1 | miR-29a/c-3p common target gene | Experiment supported target gene (TarBase v.8) |
| ENSG00000135547 | HEY2 | miR-29c-3p target gene only | Theoretical prediction (microT-CDS algorithm) |
| ENSG00000135636 | DYSF | miR-29a-3p target gene only | Experiment supported target gene (TarBase v.8) |
| ENSG00000135677 | GNS | miR-29a/c-3p common target gene | Theoretical prediction (microT-CDS algorithm) |
| ENSG00000135679 | MDM2 | miR-29a/c-3p common target gene | Experiment supported target gene (TarBase v.8) |

| | | | |
|---|---|---|---|
| ENSG00000135709 | KIAA0513 | miR-29c-3p target gene only | Experiment supported target gene (TarBase v.8) |
| ENSG00000135740 | SLC9A5 | miR-29a/c-3p common target gene | Theoretical prediction (microT-CDS algorithm) |
| ENSG00000135750 | KCNK1 | miR-29c-3p target gene only | Experiment supported target gene (TarBase v.8) |
| ENSG00000135773 | CAPN9 | miR-29c-3p target gene only | Experiment supported target gene (TarBase v.8) |
| ENSG00000135828 | RNASEL | miR-29a/c-3p common target gene | Experiment supported target gene (TarBase v.8) |
| ENSG00000135829 | DHX9 | miR-29a/c-3p common target gene | Experiment supported target gene (TarBase v.8) |
| ENSG00000135835 | KIAA1614 | miR-29a/c-3p common target gene | Experiment supported target gene (TarBase v.8) |
| ENSG00000135862 | LAMC1 | miR-29a/c-3p common target gene | Experiment supported target gene (TarBase v.8) |
| ENSG00000135870 | RC3H1 | miR-29a-3p target gene only | Experiment supported target gene (TarBase v.8) |
| ENSG00000135899 | SP110 | miR-29c-3p target gene only | Experiment supported target gene (TarBase v.8) |
| ENSG00000135902 | CHRND | miR-29a/c-3p common target gene | Theoretical prediction (microT-CDS algorithm) |
| ENSG00000135913 | USP37 | miR-29a/c-3p common target gene | Experiment supported target gene (TarBase v.8) |
| ENSG00000135930 | EIF4E2 | miR-29a/c-3p common target gene | Experiment supported target gene (TarBase v.8) |
| ENSG00000135931 | ARMC9 | miR-29c-3p target gene only | Experiment supported target gene (TarBase v.8) |
| ENSG00000135945 | REV1 | miR-29c-3p target gene only | Experiment supported target gene (TarBase v.8) |
| ENSG00000135956 | TMEM127 | miR-29a/c-3p common target gene | Experiment supported target gene (TarBase v.8) |
| ENSG00000136014 | USP44 | miR-29a/c-3p common target gene | Experiment supported target gene (TarBase v.8) |
| ENSG00000136021 | SCYL2 | miR-29a/c-3p common target gene | Experiment supported target gene (TarBase v.8) |
| ENSG00000136100 | VPS36 | miR-29c-3p target gene only | Experiment supported target gene (TarBase v.8) |
| ENSG00000136114 | THSD1 | miR-29a-3p target gene only | Experiment supported target gene (TarBase v.8) |
| ENSG00000136143 | SUCLA2 | miR-29c-3p target gene only | Experiment supported target gene (TarBase v.8) |
| ENSG00000136160 | EDNRB | miR-29a/c-3p common target gene | Theoretical prediction (microT-CDS algorithm) |
| ENSG00000136169 | SETDB2 | miR-29a/c-3p common target gene | Theoretical prediction (microT-CDS algorithm) |
| ENSG00000136205 | TNS3 | miR-29a-3p target gene only | Experiment supported target gene (TarBase v.8) |
| ENSG00000136235 | GPNMB | miR-29a-3p target gene only | Experiment supported target gene (TarBase v.8) |
| ENSG00000136240 | KDELR2 | miR-29a-3p target gene only | Experiment supported target gene (TarBase v.8) |
| ENSG00000136295 | TTYH3 | miR-29a/c-3p common target gene | Experiment supported target gene (TarBase v.8) |
| ENSG00000136378 | ADAMTS7 | miR-29a/c-3p common target gene | Theoretical prediction (microT-CDS algorithm) |
| ENSG00000136381 | IREB2 | miR-29a/c-3p common target gene | Experiment supported target gene (TarBase v.8) |
| ENSG00000136450 | SRSF1 | miR-29a-3p target gene only | Experiment supported target gene (TarBase v.8) |
| ENSG00000136485 | DCAF7 | miR-29c-3p target gene only | Experiment supported target gene (TarBase v.8) |
| ENSG00000136514 | RTP4 | miR-29c-3p target gene only | Experiment supported target gene (TarBase v.8) |
| ENSG00000136527 | TRA2B | miR-29a/c-3p common target gene | Experiment supported target gene (TarBase v.8) |
| ENSG00000136531 | SCN2A | miR-29a-3p target gene only | Experiment supported target gene (TarBase v.8) |
| ENSG00000136542 | GALNT5 | miR-29a/c-3p common target gene | Experiment supported target gene (TarBase v.8) |
| ENSG00000136628 | EPRS | miR-29a/c-3p common target gene | Experiment supported target gene (TarBase v.8) |
| ENSG00000136630 | HLX | miR-29c-3p target gene only | Experiment supported target gene (TarBase v.8) |
| ENSG00000136636 | KCTD3 | miR-29a/c-3p common target gene | Experiment supported target gene (TarBase v.8) |
| ENSG00000136695 | IL36RN | miR-29c-3p target gene only | Experiment supported target gene (TarBase v.8) |
| ENSG00000136699 | SMPD4 | miR-29a/c-3p common target gene | Experiment supported target gene (TarBase v.8) |
| ENSG00000136783 | NIPSNAP3A | miR-29a/c-3p common target gene | Experiment supported target gene (TarBase v.8) |
| ENSG00000136807 | CDK9 | miR-29a-3p target gene only | Experiment supported target gene (TarBase v.8) |
| ENSG00000136826 | KLF4 | miR-29a/c-3p common target gene | Experiment supported target gene (TarBase v.8) |
| ENSG00000136827 | TOR1A | miR-29c-3p target gene only | Experiment supported target gene (TarBase v.8) |
| ENSG00000136828 | RALGPS1 | miR-29a/c-3p common target gene | Experiment supported target gene (TarBase v.8) |
| ENSG00000136830 | FAM129B | miR-29c-3p target gene only | Experiment supported target gene (TarBase v.8) |
| ENSG00000136856 | SLC2A8 | miR-29c-3p target gene only | Experiment supported target gene (TarBase v.8) |
| ENSG00000136859 | ANGPTL2 | miR-29a-3p target gene only | Experiment supported target gene (TarBase v.8) |
| ENSG00000136868 | SLC31A1 | miR-29a/c-3p common target gene | Experiment supported target gene (TarBase v.8) |
| ENSG00000136874 | STX17 | miR-29a/c-3p common target gene | Theoretical prediction (microT-CDS algorithm) |
| ENSG00000136897 | MRPL50 | miR-29a/c-3p common target gene | Experiment supported target gene (TarBase v.8) |
| ENSG00000136940 | PDCL | miR-29a/c-3p common target gene | Experiment supported target gene (TarBase v.8) |
| ENSG00000136960 | ENPP2 | miR-29a/c-3p common target gene | Experiment supported target gene (TarBase v.8) |
| ENSG00000136982 | DSCC1 | miR-29c-3p target gene only | Experiment supported target gene (TarBase v.8) |
| ENSG00000136986 | DERL1 | miR-29c-3p target gene only | Experiment supported target gene (TarBase v.8) |
| ENSG00000137040 | RANBP6 | miR-29a-3p target gene only | Experiment supported target gene (TarBase v.8) |
| ENSG00000137055 | PLAA | miR-29a/c-3p common target gene | Experiment supported target gene (TarBase v.8) |
| ENSG00000137193 | PIM1 | miR-29c-3p target gene only | Experiment supported target gene (TarBase v.8) |
| ENSG00000137204 | SLC22A7 | miR-29a/c-3p common target gene | Theoretical prediction (microT-CDS algorithm) |
| ENSG00000137267 | TUBB2A | miR-29a/c-3p common target gene | Experiment supported target gene (TarBase v.8) |
| ENSG00000137440 | FGFBP1 | miR-29c-3p target gene only | Experiment supported target gene (TarBase v.8) |
| ENSG00000137502 | RAB30 | miR-29a/c-3p common target gene | Experiment supported target gene (TarBase v.8) |
| ENSG00000137509 | PRCP | miR-29a/c-3p common target gene | Experiment supported target gene (TarBase v.8) |
| ENSG00000137558 | PI15 | miR-29a/c-3p common target gene | Experiment supported target gene (TarBase v.8) |
| ENSG00000137571 | SLCO5A1 | miR-29a/c-3p common target gene | Theoretical prediction (microT-CDS algorithm) |
| ENSG00000137628 | DDX60 | miR-29a-3p target gene only | Experiment supported target gene (TarBase v.8) |
| ENSG00000137692 | DCUN1D5 | miR-29a/c-3p common target gene | Experiment supported target gene (TarBase v.8) |
| ENSG00000137699 | TRIM29 | miR-29c-3p target gene only | Experiment supported target gene (TarBase v.8) |
| ENSG00000137713 | PPP2R1B | miR-29c-3p target gene only | Experiment supported target gene (TarBase v.8) |
| ENSG00000137720 | C11orf1 | miR-29c-3p target gene only | Experiment supported target gene (TarBase v.8) |
| ENSG00000137752 | CASP1 | miR-29a-3p target gene only | Experiment supported target gene (TarBase v.8) |
| ENSG00000137760 | ALKBH8 | miR-29a/c-3p common target gene | Theoretical prediction (microT-CDS algorithm) |
| ENSG00000137770 | CTDSPL2 | miR-29a/c-3p common target gene | Experiment supported target gene (TarBase v.8) |
| ENSG00000137801 | THBS1 | miR-29a/c-3p common target gene | Experiment supported target gene (TarBase v.8) |
| ENSG00000137809 | ITGA11 | miR-29a/c-3p common target gene | Experiment supported target gene (TarBase v.8) |
| ENSG00000137824 | RMDN3 | miR-29a/c-3p common target gene | Theoretical prediction (microT-CDS algorithm) |
| ENSG00000137831 | UACA | miR-29a/c-3p common target gene | Experiment supported target gene (TarBase v.8) |
| ENSG00000137880 | GCHFR | miR-29c-3p target gene only | Experiment supported target gene (TarBase v.8) |
| ENSG00000137941 | TTLL7 | miR-29c-3p target gene only | Experiment supported target gene (TarBase v.8) |
| ENSG00000137959 | IFI44L | miR-29c-3p target gene only | Experiment supported target gene (TarBase v.8) |
| ENSG00000137965 | IFI44 | miR-29c-3p target gene only | Experiment supported target gene (TarBase v.8) |
| ENSG00000137968 | SLC44A5 | miR-29a/c-3p common target gene | Theoretical prediction (microT-CDS algorithm) |
| ENSG00000137992 | DBT | miR-29a/c-3p common target gene | Experiment supported target gene (TarBase v.8) |
| ENSG00000138002 | IFT172 | miR-29c-3p target gene only | Experiment supported target gene (TarBase v.8) |
| ENSG00000138028 | CGREF1 | miR-29c-3p target gene only | Experiment supported target gene (TarBase v.8) |
| ENSG00000138030 | KHK | miR-29c-3p target gene only | Experiment supported target gene (TarBase v.8) |
| ENSG00000138032 | PPM1B | miR-29a-3p target gene only | Experiment supported target gene (TarBase v.8) |
| ENSG00000138035 | PNPT1 | miR-29a/c-3p common target gene | Theoretical prediction (microT-CDS algorithm) |
| ENSG00000138061 | CYP1B1 | miR-29c-3p target gene only | Experiment supported target gene (TarBase v.8) |
| ENSG00000138069 | RAB1A | miR-29a/c-3p common target gene | Experiment supported target gene (TarBase v.8) |
| ENSG00000138078 | PREPL | miR-29a/c-3p common target gene | Experiment supported target gene (TarBase v.8) |
| ENSG00000138160 | KIF11 | miR-29a-3p target gene only | Experiment supported target gene (TarBase v.8) |
| ENSG00000138166 | DUSP5 | miR-29a-3p target gene only | Experiment supported target gene (TarBase v.8) |
| ENSG00000138185 | ENTPD1 | miR-29a-3p target gene only | Experiment supported target gene (TarBase v.8) |
| ENSG00000138303 | ASCC1 | miR-29c-3p target gene only | Experiment supported target gene (TarBase v.8) |
| ENSG00000138316 | ADAMTS14 | miR-29a/c-3p common target gene | Theoretical prediction (microT-CDS algorithm) |
| ENSG00000138326 | RPS24 | miR-29c-3p target gene only | Experiment supported target gene (TarBase v.8) |
| ENSG00000138336 | TET1 | miR-29a/c-3p common target gene | Experiment supported target gene (TarBase v.8) |
| ENSG00000138347 | MYPN | miR-29a-3p target gene only | Experiment supported target gene (TarBase v.8) |
| ENSG00000138398 | PPIG | miR-29a/c-3p common target gene | Experiment supported target gene (TarBase v.8) |
| ENSG00000138439 | FAM117B | miR-29a-3p target gene only | Experiment supported target gene (TarBase v.8) |
| ENSG00000138449 | SLC40A1 | miR-29a-3p target gene only | Experiment supported target gene (TarBase v.8) |
| ENSG00000138496 | PARP9 | miR-29c-3p target gene only | Experiment supported target gene (TarBase v.8) |
| ENSG00000138594 | TMOD3 | miR-29a/c-3p common target gene | Experiment supported target gene (TarBase v.8) |
| ENSG00000138614 | VWA9 | miR-29c-3p target gene only | Experiment supported target gene (TarBase v.8) |
| ENSG00000138642 | HERC6 | miR-29c-3p target gene only | Experiment supported target gene (TarBase v.8) |
| ENSG00000138674 | SEC31A | miR-29a/c-3p common target gene | Experiment supported target gene (TarBase v.8) |
| ENSG00000138758 | 11-Sep | miR-29a/c-3p common target gene | Experiment supported target gene (TarBase v.8) |
| ENSG00000138759 | FRAS1 | miR-29a/c-3p common target gene | Theoretical prediction (microT-CDS algorithm) |
| ENSG00000138768 | USO1 | miR-29a/c-3p common target gene | Experiment supported target gene (TarBase v.8) |
| ENSG00000138769 | CDKL2 | miR-29a/c-3p common target gene | Theoretical prediction (microT-CDS algorithm) |
| ENSG00000138778 | CENPE | miR-29c-3p target gene only | Experiment supported target gene (TarBase v.8) |
| ENSG00000138792 | ENPEP | miR-29a-3p target gene only | Experiment supported target gene (TarBase v.8) |
| ENSG00000138796 | HADH | miR-29c-3p target gene only | Experiment supported target gene (TarBase v.8) |
| ENSG00000138801 | PAPSS1 | miR-29a-3p target gene only | Experiment supported target gene (TarBase v.8) |
| ENSG00000138829 | FBN2 | miR-29a-3p target gene only | Experiment supported target gene (TarBase v.8) |
| ENSG00000139083 | ETV6 | miR-29a/c-3p common target gene | Theoretical prediction (microT-CDS algorithm) |
| ENSG00000139132 | FGD4 | miR-29a/c-3p common target gene | Theoretical prediction (microT-CDS algorithm) |
| ENSG00000139163 | ETNK1 | miR-29a-3p target gene only | Experiment supported target gene (TarBase v.8) |
| ENSG00000139174 | PRICKLE1 | miR-29a-3p target gene only | Experiment supported target gene (TarBase v.8) |
| ENSG00000139211 | AMIGO2 | miR-29a/c-3p common target gene | Experiment supported target gene (TarBase v.8) |
| ENSG00000139218 | SCAF11 | miR-29a/c-3p common target gene | Experiment supported target gene (TarBase v.8) |
| ENSG00000139219 | COL2A1 | miR-29a/c-3p common target gene | Theoretical prediction (microT-CDS algorithm) |
| ENSG00000139263 | LRIG3 | miR-29c-3p target gene only | Experiment supported target gene (TarBase v.8) |
| ENSG00000139324 | TMTC3 | miR-29a/c-3p common target gene | Experiment supported target gene (TarBase v.8) |
| ENSG00000139372 | TDG | miR-29a/c-3p common target gene | Experiment supported target gene (TarBase v.8) |
| ENSG00000139436 | GIT2 | miR-29a-3p target gene only | Experiment supported target gene (TarBase v.8) |
| ENSG00000139567 | ACVRL1 | miR-29a-3p target gene only | Experiment supported target gene (TarBase v.8) |
| ENSG00000139597 | N4BP2L1 | miR-29a/c-3p common target gene | Experiment supported target gene (TarBase v.8) |
| ENSG00000139636 | LMBR1L | miR-29a/c-3p common target gene | Theoretical prediction (microT-CDS algorithm) |
| ENSG00000139641 | ESYT1 | miR-29a-3p target gene only | Experiment supported target gene (TarBase v.8) |
| ENSG00000139645 | ANKRD52 | miR-29a/c-3p common target gene | Experiment supported target gene (TarBase v.8) |
| ENSG00000139697 | SBNO1 | miR-29a/c-3p common target gene | Experiment supported target gene (TarBase v.8) |
| ENSG00000139734 | DIAPH3 | miR-29a-3p target gene only | Experiment supported target gene (TarBase v.8) |
| ENSG00000139746 | RBM26 | miR-29a/c-3p common target gene | Theoretical prediction (microT-CDS algorithm) |
| ENSG00000139800 | ZIC5 | miR-29a/c-3p common target gene | Theoretical prediction (microT-CDS algorithm) |
| ENSG00000139826 | ABHD13 | miR-29a/c-3p common target gene | Experiment supported target gene (TarBase v.8) |
| ENSG00000139910 | NOVA1 | miR-29a/c-3p common target gene | Theoretical prediction (microT-CDS algorithm) |
| ENSG00000139915 | MDGA2 | miR-29a/c-3p common target gene | Theoretical prediction (microT-CDS algorithm) |
| ENSG00000139973 | SYT16 | miR-29a-3p target gene only | Experiment supported target gene (TarBase v.8) |
| ENSG00000139990 | DCAF5 | miR-29a-3p target gene only | Theoretical prediction (microT-CDS algorithm) |
| ENSG00000139998 | RAB15 | miR-29a/c-3p common target gene | Theoretical prediction (microT-CDS algorithm) |
| ENSG00000140044 | JDP2 | miR-29c-3p target gene only | Experiment supported target gene (TarBase v.8) |
| ENSG00000140263 | SORD | miR-29c-3p target gene only | Experiment supported target gene (TarBase v.8) |
| ENSG00000140299 | BNIP2 | miR-29a/c-3p common target gene | Experiment supported target gene (TarBase v.8) |
| ENSG00000140323 | DISP2 | miR-29a/c-3p common target gene | Experiment supported target gene (TarBase v.8) |
| ENSG00000140398 | NEIL1 | miR-29c-3p target gene only | Experiment supported target gene (TarBase v.8) |
| ENSG00000140416 | TPM1 | miR-29a/c-3p common target gene | Experiment supported target gene (TarBase v.8) |
| ENSG00000140450 | ARRDC4 | miR-29a/c-3p common target gene | Experiment supported target gene (TarBase v.8) |
| ENSG00000140455 | USP3 | miR-29a-3p target gene only | Experiment supported target gene (TarBase v.8) |
| ENSG00000140464 | PML | miR-29c-3p target gene only | Experiment supported target gene (TarBase v.8) |
| ENSG00000140465 | CYP1A1 | miR-29c-3p target gene only | Experiment supported target gene (TarBase v.8) |
| ENSG00000140470 | ADAMTS17 | miR-29a/c-3p common target gene | Theoretical prediction (microT-CDS algorithm) |
| ENSG00000140511 | HAPLN3 | miR-29a/c-3p common target gene | Experiment supported target gene (TarBase v.8) |
| ENSG00000140548 | ZNF710 | miR-29a-3p target gene only | Experiment supported target gene (TarBase v.8) |
| ENSG00000140553 | UNC45A | miR-29a-3p target gene only | Experiment supported target gene (TarBase v.8) |
| ENSG00000140836 | ZFHX3 | miR-29a-3p target gene only | Experiment supported target gene (TarBase v.8) |
| ENSG00000140873 | ADAMTS18 | miR-29a/c-3p common target gene | Theoretical prediction (microT-CDS algorithm) |
| ENSG00000140905 | GCSH | miR-29a/c-3p common target gene | Experiment supported target gene (TarBase v.8) |
| ENSG00000140939 | NOL3 | miR-29c-3p target gene only | Experiment supported target gene (TarBase v.8) |
| ENSG00000140948 | ZCCHC14 | miR-29a/c-3p common target gene | Experiment supported target gene (TarBase v.8) |
| ENSG00000140983 | RHOT2 | miR-29c-3p target gene only | Experiment supported target gene (TarBase v.8) |
| ENSG00000140995 | DEF8 | miR-29a-3p target gene only | Experiment supported target gene (TarBase v.8) |
| ENSG00000141179 | PCTP | miR-29c-3p target gene only | Experiment supported target gene (TarBase v.8) |
| ENSG00000141258 | SGSM2 | miR-29a-3p target gene only | Experiment supported target gene (TarBase v.8) |
| ENSG00000141293 | SKAP1 | miR-29c-3p target gene only | Experiment supported target gene (TarBase v.8) |
| ENSG00000141338 | ABCA8 | miR-29a-3p target gene only | Experiment supported target gene (TarBase v.8) |
| ENSG00000141367 | CLTC | miR-29a-3p target gene only | Experiment supported target gene (TarBase v.8) |
| ENSG00000141429 | GALNT1 | miR-29c-3p target gene only | Experiment supported target gene (TarBase v.8) |
| ENSG00000141431 | ASXL3 | miR-29a/c-3p common target gene | Theoretical prediction (microT-CDS algorithm) |
| ENSG00000141446 | ESCO1 | miR-29a/c-3p common target gene | Theoretical prediction (microT-CDS algorithm) |
| ENSG00000141449 | GREB1L | miR-29a/c-3p common target gene | Theoretical prediction (microT-CDS algorithm) |
| ENSG00000141458 | NPC1 | miR-29a/c-3p common target gene | Experiment supported target gene (TarBase v.8) |
| ENSG00000141480 | ARRB2 | miR-29a-3p target gene only | Experiment supported target gene (TarBase v.8) |
| ENSG00000141505 | ASGR1 | miR-29a-3p target gene only | Experiment supported target gene (TarBase v.8) |
| ENSG00000141510 | TP53 | miR-29a/c-3p common target gene | Experiment supported target gene (TarBase v.8) |
| ENSG00000141526 | SLC16A3 | miR-29a/c-3p common target gene | Experiment supported target gene (TarBase v.8) |
| ENSG00000141562 | NARF | miR-29a/c-3p common target gene | Theoretical prediction (microT-CDS algorithm) |
| ENSG00000141582 | CBX4 | miR-29a/c-3p common target gene | Experiment supported target gene (TarBase v.8) |
| ENSG00000141622 | RNF165 | miR-29a/c-3p common target gene | Theoretical prediction (microT-CDS algorithm) |
| ENSG00000141627 | DYM | miR-29c-3p target gene only | Experiment supported target gene (TarBase v.8) |
| ENSG00000141665 | FBXO15 | miR-29c-3p target gene only | Experiment supported target gene (TarBase v.8) |
| ENSG00000141682 | PMAIP1 | miR-29c-3p target gene only | Experiment supported target gene (TarBase v.8) |
| ENSG00000141867 | BRD4 | miR-29a/c-3p common target gene | Experiment supported target gene (TarBase v.8) |
| ENSG00000141905 | NFIC | miR-29a/c-3p common target gene | Theoretical prediction (microT-CDS algorithm) |
| ENSG00000141985 | SH3GL1 | miR-29a/c-3p common target gene | Experiment supported target gene (TarBase v.8) |
| ENSG00000142089 | IFITM3 | miR-29c-3p target gene only | Experiment supported target gene (TarBase v.8) |
| ENSG00000142156 | COL6A1 | miR-29a/c-3p common target gene | Experiment supported target gene (TarBase v.8) |
| ENSG00000142173 | COL6A2 | miR-29a/c-3p common target gene | Experiment supported target gene (TarBase v.8) |
| ENSG00000142227 | EMP3 | miR-29a/c-3p common target gene | Experiment supported target gene (TarBase v.8) |
| ENSG00000142303 | ADAMTS10 | miR-29a/c-3p common target gene | Theoretical prediction (microT-CDS algorithm) |
| ENSG00000142327 | RNPEPL1 | miR-29a-3p target gene only | Experiment supported target gene (TarBase v.8) |
| ENSG00000142556 | ZNF614 | miR-29a/c-3p common target gene | Experiment supported target gene (TarBase v.8) |
| ENSG00000142599 | RERE | miR-29a/c-3p common target gene | Experiment supported target gene (TarBase v.8) |
| ENSG00000142627 | EPHA2 | miR-29a-3p target gene only | Experiment supported target gene (TarBase v.8) |
| ENSG00000142798 | HSPG2 | miR-29a-3p target gene only | Experiment supported target gene (TarBase v.8) |
| ENSG00000142864 | SERBP1 | miR-29a/c-3p common target gene | Experiment supported target gene (TarBase v.8) |
| ENSG00000142871 | CYR61 | miR-29c-3p target gene only | Experiment supported target gene (TarBase v.8) |
| ENSG00000142949 | PTPRF | miR-29a/c-3p common target gene | Experiment supported target gene (TarBase v.8) |
| ENSG00000143006 | DMRTB1 | miR-29a/c-3p common target gene | Theoretical prediction (microT-CDS algorithm) |
| ENSG00000143028 | SYPL2 | miR-29a/c-3p common target gene | Theoretical prediction (microT-CDS algorithm) |
| ENSG00000143061 | IGSF3 | miR-29c-3p target gene only | Experiment supported target gene (TarBase v.8) |
| ENSG00000143153 | ATP1B1 | miR-29a/c-3p common target gene | Experiment supported target gene (TarBase v.8) |
| ENSG00000143156 | NME7 | miR-29a/c-3p common target gene | Experiment supported target gene (TarBase v.8) |
| ENSG00000143162 | CREG1 | miR-29a/c-3p common target gene | Experiment supported target gene (TarBase v.8) |
| ENSG00000143179 | UCK2 | miR-29c-3p target gene only | Experiment supported target gene (TarBase v.8) |
| ENSG00000143183 | TMCO1 | miR-29c-3p target gene only | Experiment supported target gene (TarBase v.8) |
| ENSG00000143207 | RFWD2 | miR-29c-3p target gene only | Experiment supported target gene (TarBase v.8) |
| ENSG00000143248 | RGS5 | miR-29a/c-3p common target gene | Experiment supported target gene (TarBase v.8) |
| ENSG00000143319 | ISG20L2 | miR-29a/c-3p common target gene | Experiment supported target gene (TarBase v.8) |
| ENSG00000143320 | CRABP2 | miR-29a/c-3p common target gene | Experiment supported target gene (TarBase v.8) |
| ENSG00000143321 | HDGF | miR-29a/c-3p common target gene | Experiment supported target gene (TarBase v.8) |
| ENSG00000143341 | HMCN1 | miR-29a/c-3p common target gene | Theoretical prediction (microT-CDS algorithm) |
| ENSG00000143374 | TARS2 | miR-29c-3p target gene only | Experiment supported target gene (TarBase v.8) |
| ENSG00000143379 | SETDB1 | miR-29a/c-3p common target gene | Theoretical prediction (microT-CDS algorithm) |
| ENSG00000143384 | MCL1 | miR-29a/c-3p common target gene | Experiment supported target gene (TarBase v.8) |
| ENSG00000143387 | CTSK | miR-29a-3p target gene only | Experiment supported target gene (TarBase v.8) |
| ENSG00000143412 | ANXA9 | miR-29c-3p target gene only | Experiment supported target gene (TarBase v.8) |
| ENSG00000143437 | ARNT | miR-29a/c-3p common target gene | Experiment supported target gene (TarBase v.8) |
| ENSG00000143442 | POGZ | miR-29a-3p target gene only | Experiment supported target gene (TarBase v.8) |
| ENSG00000143457 | GOLPH3L | miR-29a-3p target gene only | Experiment supported target gene (TarBase v.8) |
| ENSG00000143473 | KCNH1 | miR-29c-3p target gene only | Experiment supported target gene (TarBase v.8) |
| ENSG00000143612 | C1orf43 | miR-29a/c-3p common target gene | Experiment supported target gene (TarBase v.8) |
| ENSG00000143614 | GATAD2B | miR-29a/c-3p common target gene | Experiment supported target gene (TarBase v.8) |
| ENSG00000143622 | RIT1 | miR-29a/c-3p common target gene | Experiment supported target gene (TarBase v.8) |
| ENSG00000143624 | INTS3 | miR-29a-3p target gene only | Theoretical prediction (microT-CDS algorithm) |
| ENSG00000143633 | C1orf131 | miR-29a/c-3p common target gene | Experiment supported target gene (TarBase v.8) |
| ENSG00000143740 | SNAP47 | miR-29a/c-3p common target gene | Theoretical prediction (microT-CDS algorithm) |
| ENSG00000143776 | CDC42BPA | miR-29a/c-3p common target gene | Theoretical prediction (microT-CDS algorithm) |
| ENSG00000143799 | PARP1 | miR-29a/c-3p common target gene | Experiment supported target gene (TarBase v.8) |
| ENSG00000143801 | PSEN2 | miR-29c-3p target gene only | Experiment supported target gene (TarBase v.8) |
| ENSG00000143847 | PPFIA4 | miR-29c-3p target gene only | Experiment supported target gene (TarBase v.8) |
| ENSG00000143858 | SYT2 | miR-29a-3p target gene only | Experiment supported target gene (TarBase v.8) |
| ENSG00000143870 | PDIA6 | miR-29a-3p target gene only | Experiment supported target gene (TarBase v.8) |
| ENSG00000143924 | EML4 | miR-29c-3p target gene only | Experiment supported target gene (TarBase v.8) |
| ENSG00000143947 | RPS27A | miR-29a/c-3p common target gene | Experiment supported target gene (TarBase v.8) |
| ENSG00000143970 | ASXL2 | miR-29a/c-3p common target gene | Experiment supported target gene (TarBase v.8) |
| ENSG00000144063 | MALL | miR-29c-3p target gene only | Experiment supported target gene (TarBase v.8) |
| ENSG00000144115 | THNSL2 | miR-29c-3p target gene only | Experiment supported target gene (TarBase v.8) |
| ENSG00000144233 | AMMECR1L | miR-29a/c-3p common target gene | Experiment supported target gene (TarBase v.8) |
| ENSG00000144339 | TMEFF2 | miR-29a/c-3p common target gene | Experiment supported target gene (TarBase v.8) |
| ENSG00000144381 | HSPD1 | miR-29a/c-3p common target gene | Experiment supported target gene (TarBase v.8) |
| ENSG00000144401 | METTL21A | miR-29c-3p target gene only | Experiment supported target gene (TarBase v.8) |
| ENSG00000144455 | SUMF1 | miR-29c-3p target gene only | Experiment supported target gene (TarBase v.8) |
| ENSG00000144460 | NYAP2 | miR-29a/c-3p common target gene | Experiment supported target gene (TarBase v.8) |
| ENSG00000144619 | CNTN4 | miR-29a-3p target gene only | Experiment supported target gene (TarBase v.8) |
| ENSG00000144677 | CTDSPL | miR-29c-3p target gene only | Experiment supported target gene (TarBase v.8) |
| ENSG00000144724 | PTPRG | miR-29a-3p target gene only | Theoretical prediction (microT-CDS algorithm) |

| | | | |
|---|---|---|---|
| ENSG00000144747 | TMF1 | miR-29a/c-3p common target gene | Experiment supported target gene (TarBase v.8) |
| ENSG00000144791 | LIMD1 | miR-29a-3p target gene only | Experiment supported target gene (TarBase v.8) |
| ENSG00000144810 | COL8A1 | miR-29a/c-3p common target gene | Theoretical prediction (microT-CDS algorithm) |
| ENSG00000144824 | PHLDB2 | miR-29a/c-3p common target gene | Theoretical prediction (microT-CDS algorithm) |
| ENSG00000144867 | SRPRB | miR-29c-3p target gene only | Experiment supported target gene (TarBase v.8) |
| ENSG00000144893 | MED12L | miR-29a/c-3p common target gene | Theoretical prediction (microT-CDS algorithm) |
| ENSG00000144909 | OSBPL11 | miR-29a/c-3p common target gene | Experiment supported target gene (TarBase v.8) |
| ENSG00000145246 | ATP10D | miR-29a/c-3p common target gene | Theoretical prediction (microT-CDS algorithm) |
| ENSG00000145287 | PLAC8 | miR-29c-3p target gene only | Experiment supported target gene (TarBase v.8) |
| ENSG00000145293 | ENOPH1 | miR-29a-3p target gene only | Experiment supported target gene (TarBase v.8) |
| ENSG00000145331 | TRMT10A | miR-29c-3p target gene only | Experiment supported target gene (TarBase v.8) |
| ENSG00000145332 | KLHL8 | miR-29a/c-3p common target gene | Theoretical prediction (microT-CDS algorithm) |
| ENSG00000145335 | SNCA | miR-29a-3p target gene only | Experiment supported target gene (TarBase v.8) |
| ENSG00000145386 | CCNA2 | miR-29a/c-3p common target gene | Experiment supported target gene (TarBase v.8) |
| ENSG00000145391 | SETD7 | miR-29a/c-3p common target gene | Experiment supported target gene (TarBase v.8) |
| ENSG00000145416 | 1-Mar | miR-29a/c-3p common target gene | Experiment supported target gene (TarBase v.8) |
| ENSG00000145431 | PDGFC | miR-29a/c-3p common target gene | Theoretical prediction (microT-CDS algorithm) |
| ENSG00000145545 | SRD5A1 | miR-29c-3p target gene only | Experiment supported target gene (TarBase v.8) |
| ENSG00000145604 | SKP2 | miR-29a/c-3p common target gene | Experiment supported target gene (TarBase v.8) |
| ENSG00000145632 | PLK2 | miR-29a-3p target gene only | Experiment supported target gene (TarBase v.8) |
| ENSG00000145675 | PIK3R1 | miR-29a/c-3p common target gene | Experiment supported target gene (TarBase v.8) |
| ENSG00000145725 | PPIP5K2 | miR-29c-3p target gene only | Experiment supported target gene (TarBase v.8) |
| ENSG00000145780 | FEM1C | miR-29a/c-3p common target gene | Experiment supported target gene (TarBase v.8) |
| ENSG00000145808 | ADAMTS19 | miR-29a/c-3p common target gene | Theoretical prediction (microT-CDS algorithm) |
| ENSG00000145824 | CXCL14 | miR-29a-3p target gene only | Experiment supported target gene (TarBase v.8) |
| ENSG00000145882 | PCYOX1L | miR-29c-3p target gene only | Experiment supported target gene (TarBase v.8) |
| ENSG00000145901 | TNIP1 | miR-29a-3p target gene only | Experiment supported target gene (TarBase v.8) |
| ENSG00000145907 | G3BP1 | miR-29a/c-3p common target gene | Experiment supported target gene (TarBase v.8) |
| ENSG00000145945 | FAM50B | miR-29a-3p target gene only | Experiment supported target gene (TarBase v.8) |
| ENSG00000145979 | TBC1D7 | miR-29a/c-3p common target gene | Experiment supported target gene (TarBase v.8) |
| ENSG00000145990 | GFOD1 | miR-29a/c-3p common target gene | Experiment supported target gene (TarBase v.8) |
| ENSG00000146067 | FAM193B | miR-29a/c-3p common target gene | Experiment supported target gene (TarBase v.8) |
| ENSG00000146072 | TNFRSF21 | miR-29a/c-3p common target gene | Experiment supported target gene (TarBase v.8) |
| ENSG00000146151 | HMGCLL1 | miR-29a/c-3p common target gene | Experiment supported target gene (TarBase v.8) |
| ENSG00000146267 | FAXC | miR-29a/c-3p common target gene | Theoretical prediction (microT-CDS algorithm) |
| ENSG00000146281 | PM20D2 | miR-29a/c-3p common target gene | Experiment supported target gene (TarBase v.8) |
| ENSG00000146373 | RNF217 | miR-29a/c-3p common target gene | Experiment supported target gene (TarBase v.8) |
| ENSG00000146409 | SLC18B1 | miR-29c-3p target gene only | Experiment supported target gene (TarBase v.8) |
| ENSG00000146416 | AIG1 | miR-29c-3p target gene only | Experiment supported target gene (TarBase v.8) |
| ENSG00000146425 | DYNLT1 | miR-29a/c-3p common target gene | Experiment supported target gene (TarBase v.8) |
| ENSG00000146433 | TMEM181 | miR-29a/c-3p common target gene | Experiment supported target gene (TarBase v.8) |
| ENSG00000146555 | SDK1 | miR-29a/c-3p common target gene | Theoretical prediction (microT-CDS algorithm) |
| ENSG00000146587 | RBAK | miR-29a/c-3p common target gene | Experiment supported target gene (TarBase v.8) |
| ENSG00000146592 | CREB5 | miR-29a/c-3p common target gene | Experiment supported target gene (TarBase v.8) |
| ENSG00000146674 | IGFBP3 | miR-29a/c-3p common target gene | Experiment supported target gene (TarBase v.8) |
| ENSG00000146757 | ZNF92 | miR-29c-3p target gene only | Experiment supported target gene (TarBase v.8) |
| ENSG00000146904 | EPHA1 | miR-29c-3p target gene only | Experiment supported target gene (TarBase v.8) |
| ENSG00000146950 | SHROOM2 | miR-29a/c-3p common target gene | Theoretical prediction (microT-CDS algorithm) |
| ENSG00000147202 | DIAPH2 | miR-29a/c-3p common target gene | Experiment supported target gene (TarBase v.8) |
| ENSG00000147255 | IGSF1 | miR-29a/c-3p common target gene | Theoretical prediction (microT-CDS algorithm) |
| ENSG00000147256 | ARHGAP36 | miR-29a/c-3p common target gene | Theoretical prediction (microT-CDS algorithm) |
| ENSG00000147257 | GPC3 | miR-29a-3p target gene only | Experiment supported target gene (TarBase v.8) |
| ENSG00000147408 | CSGALNACT1 | miR-29a-3p target gene only | Experiment supported target gene (TarBase v.8) |
| ENSG00000147475 | ERLIN2 | miR-29a/c-3p common target gene | Theoretical prediction (microT-CDS algorithm) |
| ENSG00000147526 | TACC1 | miR-29a/c-3p common target gene | Experiment supported target gene (TarBase v.8) |
| ENSG00000147533 | GOLGA7 | miR-29a/c-3p common target gene | Experiment supported target gene (TarBase v.8) |
| ENSG00000147576 | ADHFE1 | miR-29a/c-3p common target gene | Theoretical prediction (microT-CDS algorithm) |
| ENSG00000147592 | LACTB2 | miR-29c-3p target gene only | Experiment supported target gene (TarBase v.8) |
| ENSG00000147601 | TERF1 | miR-29a/c-3p common target gene | Theoretical prediction (microT-CDS algorithm) |
| ENSG00000147679 | UTP23 | miR-29c-3p target gene only | Experiment supported target gene (TarBase v.8) |
| ENSG00000147862 | NFIB | miR-29a/c-3p common target gene | Experiment supported target gene (TarBase v.8) |
| ENSG00000147894 | C9orf72 | miR-29a/c-3p common target gene | Theoretical prediction (microT-CDS algorithm) |

| | | | |
|---|---|---|---|
| ENSG00000148053 | NTRK2 | miR-29a/c-3p common target gene | Experiment supported target gene (TarBase v.8) |
| ENSG00000148057 | IDNK | miR-29c-3p target gene only | Experiment supported target gene (TarBase v.8) |
| ENSG00000148120 | C9orf3 | miR-29c-3p target gene only | Experiment supported target gene (TarBase v.8) |
| ENSG00000148154 | UGCG | miR-29a-3p target gene only | Experiment supported target gene (TarBase v.8) |
| ENSG00000148229 | POLE3 | miR-29a/c-3p common target gene | Experiment supported target gene (TarBase v.8) |
| ENSG00000148335 | NTMT1 | miR-29a/c-3p common target gene | Experiment supported target gene (TarBase v.8) |
| ENSG00000148358 | GPR107 | miR-29c-3p target gene only | Experiment supported target gene (TarBase v.8) |
| ENSG00000148396 | SEC16A | miR-29a/c-3p common target gene | Experiment supported target gene (TarBase v.8) |
| ENSG00000148429 | USP6NL | miR-29a/c-3p common target gene | Theoretical prediction (microT-CDS algorithm) |
| ENSG00000148450 | MSRB2 | miR-29c-3p target gene only | Experiment supported target gene (TarBase v.8) |
| ENSG00000148483 | TMEM236 | miR-29a/c-3p common target gene | Theoretical prediction (microT-CDS algorithm) |
| ENSG00000148672 | GLUD1 | miR-29a/c-3p common target gene | Theoretical prediction (microT-CDS algorithm) |
| ENSG00000148677 | ANKRD1 | miR-29a-3p target gene only | Experiment supported target gene (TarBase v.8) |
| ENSG00000148700 | ADD3 | miR-29a/c-3p common target gene | Experiment supported target gene (TarBase v.8) |
| ENSG00000148773 | MKI67 | miR-29a/c-3p common target gene | Experiment supported target gene (TarBase v.8) |
| ENSG00000148798 | INA | miR-29a/c-3p common target gene | Theoretical prediction (microT-CDS algorithm) |
| ENSG00000148835 | TAF5 | miR-29a/c-3p common target gene | Experiment supported target gene (TarBase v.8) |
| ENSG00000148848 | ADAM12 | miR-29a/c-3p common target gene | Experiment supported target gene (TarBase v.8) |
| ENSG00000148925 | BTBD10 | miR-29a/c-3p common target gene | Experiment supported target gene (TarBase v.8) |
| ENSG00000148985 | PGAP2 | miR-29a/c-3p common target gene | Experiment supported target gene (TarBase v.8) |
| ENSG00000149115 | TNKS1BP1 | miR-29a/c-3p common target gene | Experiment supported target gene (TarBase v.8) |
| ENSG00000149136 | SSRP1 | miR-29a-3p target gene only | Experiment supported target gene (TarBase v.8) |
| ENSG00000149150 | SLC43A1 | miR-29c-3p target gene only | Experiment supported target gene (TarBase v.8) |
| ENSG00000149177 | PTPRJ | miR-29a/c-3p common target gene | Theoretical prediction (microT-CDS algorithm) |
| ENSG00000149182 | ARFGAP2 | miR-29c-3p target gene only | Experiment supported target gene (TarBase v.8) |
| ENSG00000149187 | CELF1 | miR-29a/c-3p common target gene | Experiment supported target gene (TarBase v.8) |
| ENSG00000149201 | CCDC81 | miR-29a-3p target gene only | Theoretical prediction (microT-CDS algorithm) |
| ENSG00000149257 | SERPINH1 | miR-29a/c-3p common target gene | Experiment supported target gene (TarBase v.8) |
| ENSG00000149311 | ATM | miR-29a/c-3p common target gene | Experiment supported target gene (TarBase v.8) |
| ENSG00000149313 | AASDHPPT | miR-29a/c-3p common target gene | Experiment supported target gene (TarBase v.8) |
| ENSG00000149346 | SLX4IP | miR-29c-3p target gene only | Experiment supported target gene (TarBase v.8) |
| ENSG00000149357 | LAMTOR1 | miR-29a/c-3p common target gene | Experiment supported target gene (TarBase v.8) |
| ENSG00000149499 | EML3 | miR-29a-3p target gene only | Experiment supported target gene (TarBase v.8) |
| ENSG00000149532 | CPSF7 | miR-29a/c-3p common target gene | Experiment supported target gene (TarBase v.8) |
| ENSG00000149541 | B3GAT3 | miR-29a-3p target gene only | Experiment supported target gene (TarBase v.8) |
| ENSG00000149547 | EI24 | miR-29c-3p target gene only | Experiment supported target gene (TarBase v.8) |
| ENSG00000149554 | CHEK1 | miR-29a-3p target gene only | Experiment supported target gene (TarBase v.8) |
| ENSG00000149564 | ESAM | miR-29a-3p target gene only | Experiment supported target gene (TarBase v.8) |
| ENSG00000149577 | SIDT2 | miR-29a/c-3p common target gene | Experiment supported target gene (TarBase v.8) |
| ENSG00000149600 | COMMD7 | miR-29a/c-3p common target gene | Experiment supported target gene (TarBase v.8) |
| ENSG00000149639 | SOGA1 | miR-29a/c-3p common target gene | Experiment supported target gene (TarBase v.8) |
| ENSG00000149658 | YTHDF1 | miR-29a/c-3p common target gene | Experiment supported target gene (TarBase v.8) |
| ENSG00000149679 | CABLES2 | miR-29a-3p target gene only | Experiment supported target gene (TarBase v.8) |
| ENSG00000149743 | TRPT1 | miR-29c-3p target gene only | Experiment supported target gene (TarBase v.8) |
| ENSG00000149798 | CDC42EP2 | miR-29c-3p target gene only | Experiment supported target gene (TarBase v.8) |
| ENSG00000149926 | FAM57B | miR-29a/c-3p common target gene | Experiment supported target gene (TarBase v.8) |
| ENSG00000149948 | HMGA2 | miR-29a/c-3p common target gene | Experiment supported target gene (TarBase v.8) |
| ENSG00000149972 | CNTN5 | miR-29a/c-3p common target gene | Theoretical prediction (microT-CDS algorithm) |
| ENSG00000150093 | ITGB1 | miR-29a/c-3p common target gene | Experiment supported target gene (TarBase v.8) |
| ENSG00000150275 | PCDH15 | miR-29a/c-3p common target gene | Theoretical prediction (microT-CDS algorithm) |
| ENSG00000150394 | CDH8 | miR-29a/c-3p common target gene | Theoretical prediction (microT-CDS algorithm) |
| ENSG00000150457 | LATS2 | miR-29c-3p target gene only | Experiment supported target gene (TarBase v.8) |
| ENSG00000150593 | PDCD4 | miR-29a/c-3p common target gene | Experiment supported target gene (TarBase v.8) |
| ENSG00000150656 | CNDP1 | miR-29a/c-3p common target gene | Experiment supported target gene (TarBase v.8) |
| ENSG00000150672 | DLG2 | miR-29a/c-3p common target gene | Experiment supported target gene (TarBase v.8) |
| ENSG00000150764 | DIXDC1 | miR-29a/c-3p common target gene | Theoretical prediction (microT-CDS algorithm) |
| ENSG00000150776 | C11orf57 | miR-29a/c-3p common target gene | Theoretical prediction (microT-CDS algorithm) |
| ENSG00000150779 | TIMM8B | miR-29a/c-3p common target gene | Experiment supported target gene (TarBase v.8) |
| ENSG00000150867 | PIP4K2A | miR-29c-3p target gene only | Experiment supported target gene (TarBase v.8) |
| ENSG00000150893 | FREM2 | miR-29a/c-3p common target gene | Theoretical prediction (microT-CDS algorithm) |
| ENSG00000150977 | RILPL2 | miR-29c-3p target gene only | Experiment supported target gene (TarBase v.8) |
| ENSG00000150991 | UBC | miR-29a/c-3p common target gene | Experiment supported target gene (TarBase v.8) |

| | | | |
|---|---|---|---|
| ENSG00000151025 | GPR158 | miR-29c-3p target gene only | Experiment supported target gene (TarBase v.8) |
| ENSG00000151065 | DCP1B | miR-29c-3p target gene only | Experiment supported target gene (TarBase v.8) |
| ENSG00000151117 | TMEM86A | miR-29a/c-3p common target gene | Theoretical prediction (microT-CDS algorithm) |
| ENSG00000151131 | C12orf45 | miR-29a/c-3p common target gene | Theoretical prediction (microT-CDS algorithm) |
| ENSG00000151148 | UBE3B | miR-29a/c-3p common target gene | Experiment supported target gene (TarBase v.8) |
| ENSG00000151150 | ANK3 | miR-29a/c-3p common target gene | Experiment supported target gene (TarBase v.8) |
| ENSG00000151233 | GXYLT1 | miR-29a-3p target gene only | Experiment supported target gene (TarBase v.8) |
| ENSG00000151240 | DIP2C | miR-29a/c-3p common target gene | Experiment supported target gene (TarBase v.8) |
| ENSG00000151292 | CSNK1G3 | miR-29a/c-3p common target gene | Experiment supported target gene (TarBase v.8) |
| ENSG00000151322 | NPAS3 | miR-29a/c-3p common target gene | Theoretical prediction (microT-CDS algorithm) |
| ENSG00000151376 | ME3 | miR-29a/c-3p common target gene | Experiment supported target gene (TarBase v.8) |
| ENSG00000151388 | ADAMTS12 | miR-29a/c-3p common target gene | Theoretical prediction (microT-CDS algorithm) |
| ENSG00000151422 | FER | miR-29a/c-3p common target gene | Theoretical prediction (microT-CDS algorithm) |
| ENSG00000151445 | VIPAS39 | miR-29a/c-3p common target gene | Experiment supported target gene (TarBase v.8) |
| ENSG00000151458 | ANKRD50 | miR-29c-3p target gene only | Experiment supported target gene (TarBase v.8) |
| ENSG00000151466 | SCLT1 | miR-29a/c-3p common target gene | Theoretical prediction (microT-CDS algorithm) |
| ENSG00000151474 | FRMD4A | miR-29a/c-3p common target gene | Theoretical prediction (microT-CDS algorithm) |
| ENSG00000151502 | VPS26B | miR-29a/c-3p common target gene | Experiment supported target gene (TarBase v.8) |
| ENSG00000151532 | VTI1A | miR-29c-3p target gene only | Experiment supported target gene (TarBase v.8) |
| ENSG00000151553 | FAM160B1 | miR-29a-3p target gene only | Experiment supported target gene (TarBase v.8) |
| ENSG00000151572 | ANO4 | miR-29a/c-3p common target gene | Experiment supported target gene (TarBase v.8) |
| ENSG00000151611 | MMAA | miR-29a/c-3p common target gene | Theoretical prediction (microT-CDS algorithm) |
| ENSG00000151612 | ZNF827 | miR-29a/c-3p common target gene | Experiment supported target gene (TarBase v.8) |
| ENSG00000151623 | NR3C2 | miR-29a/c-3p common target gene | Experiment supported target gene (TarBase v.8) |
| ENSG00000151693 | ASAP2 | miR-29a/c-3p common target gene | Experiment supported target gene (TarBase v.8) |
| ENSG00000151729 | SLC25A4 | miR-29c-3p target gene only | Experiment supported target gene (TarBase v.8) |
| ENSG00000151743 | AMN1 | miR-29c-3p target gene only | Experiment supported target gene (TarBase v.8) |
| ENSG00000151834 | GABRA2 | miR-29a-3p target gene only | Experiment supported target gene (TarBase v.8) |
| ENSG00000151835 | SACS | miR-29a/c-3p common target gene | Experiment supported target gene (TarBase v.8) |
| ENSG00000151883 | PARP8 | miR-29c-3p target gene only | Experiment supported target gene (TarBase v.8) |
| ENSG00000151893 | CACUL1 | miR-29a/c-3p common target gene | Experiment supported target gene (TarBase v.8) |
| ENSG00000151923 | TIAL1 | miR-29a/c-3p common target gene | Experiment supported target gene (TarBase v.8) |
| ENSG00000151967 | SCHIP1 | miR-29a/c-3p common target gene | Experiment supported target gene (TarBase v.8) |
| ENSG00000152076 | CCDC74B | miR-29c-3p target gene only | Experiment supported target gene (TarBase v.8) |
| ENSG00000152102 | FAM168B | miR-29a/c-3p common target gene | Experiment supported target gene (TarBase v.8) |
| ENSG00000152133 | GPATCH11 | miR-29c-3p target gene only | Experiment supported target gene (TarBase v.8) |
| ENSG00000152193 | RNF219 | miR-29c-3p target gene only | Experiment supported target gene (TarBase v.8) |
| ENSG00000152332 | UHMK1 | miR-29a/c-3p common target gene | Experiment supported target gene (TarBase v.8) |
| ENSG00000152409 | JMY | miR-29a/c-3p common target gene | Theoretical prediction (microT-CDS algorithm) |
| ENSG00000152439 | ZNF773 | miR-29a/c-3p common target gene | Theoretical prediction (microT-CDS algorithm) |
| ENSG00000152495 | CAMK4 | miR-29a/c-3p common target gene | Theoretical prediction (microT-CDS algorithm) |
| ENSG00000152556 | PFKM | miR-29a/c-3p common target gene | Experiment supported target gene (TarBase v.8) |
| ENSG00000152558 | TMEM123 | miR-29a/c-3p common target gene | Experiment supported target gene (TarBase v.8) |
| ENSG00000152620 | NADK2 | miR-29a/c-3p common target gene | Experiment supported target gene (TarBase v.8) |
| ENSG00000152689 | RASGRP3 | miR-29c-3p target gene only | Experiment supported target gene (TarBase v.8) |
| ENSG00000152778 | IFIT5 | miR-29c-3p target gene only | Experiment supported target gene (TarBase v.8) |
| ENSG00000152894 | PTPRK | miR-29c-3p target gene only | Experiment supported target gene (TarBase v.8) |
| ENSG00000152904 | GGPS1 | miR-29a/c-3p common target gene | Experiment supported target gene (TarBase v.8) |
| ENSG00000152932 | RAB3C | miR-29a/c-3p common target gene | Experiment supported target gene (TarBase v.8) |
| ENSG00000152939 | MARVELD2 | miR-29c-3p target gene only | Experiment supported target gene (TarBase v.8) |
| ENSG00000153064 | BANK1 | miR-29c-3p target gene only | Experiment supported target gene (TarBase v.8) |
| ENSG00000153147 | SMARCA5 | miR-29a-3p target gene only | Experiment supported target gene (TarBase v.8) |
| ENSG00000153187 | HNRNPU | miR-29a/c-3p common target gene | Experiment supported target gene (TarBase v.8) |
| ENSG00000153246 | PLA2R1 | miR-29a-3p target gene only | Experiment supported target gene (TarBase v.8) |
| ENSG00000153253 | SCN3A | miR-29a-3p target gene only | Experiment supported target gene (TarBase v.8) |
| ENSG00000153395 | LPCAT1 | miR-29c-3p target gene only | Experiment supported target gene (TarBase v.8) |
| ENSG00000153561 | RMND5A | miR-29a/c-3p common target gene | Experiment supported target gene (TarBase v.8) |
| ENSG00000153707 | PTPRD | miR-29a/c-3p common target gene | Experiment supported target gene (TarBase v.8) |
| ENSG00000153714 | LURAP1L | miR-29a-3p target gene only | Experiment supported target gene (TarBase v.8) |
| ENSG00000153721 | CNKSR3 | miR-29a/c-3p common target gene | Theoretical prediction (microT-CDS algorithm) |
| ENSG00000153790 | C7orf31 | miR-29c-3p target gene only | Experiment supported target gene (TarBase v.8) |
| ENSG00000153814 | JAZF1 | miR-29a/c-3p common target gene | Experiment supported target gene (TarBase v.8) |

| | | | |
|---|---|---|---|
| ENSG00000153827 | TRIP12 | miR-29a-3p target gene only | Experiment supported target gene (TarBase v.8) |
| ENSG00000153885 | KCTD15 | miR-29c-3p target gene only | Experiment supported target gene (TarBase v.8) |
| ENSG00000153898 | MCOLN2 | miR-29c-3p target gene only | Experiment supported target gene (TarBase v.8) |
| ENSG00000153904 | DDAH1 | miR-29c-3p target gene only | Experiment supported target gene (TarBase v.8) |
| ENSG00000153922 | CHD1 | miR-29a-3p target gene only | Experiment supported target gene (TarBase v.8) |
| ENSG00000154114 | TBCEL | miR-29a/c-3p common target gene | Experiment supported target gene (TarBase v.8) |
| ENSG00000154124 | OTULIN | miR-29a/c-3p common target gene | Experiment supported target gene (TarBase v.8) |
| ENSG00000154237 | LRRK1 | miR-29a/c-3p common target gene | Theoretical prediction (microT-CDS algorithm) |
| ENSG00000154310 | TNIK | miR-29a/c-3p common target gene | Theoretical prediction (microT-CDS algorithm) |
| ENSG00000154319 | FAM167A | miR-29a/c-3p common target gene | Experiment supported target gene (TarBase v.8) |
| ENSG00000154380 | ENAH | miR-29a/c-3p common target gene | Experiment supported target gene (TarBase v.8) |
| ENSG00000154429 | CCSAP | miR-29a/c-3p common target gene | Experiment supported target gene (TarBase v.8) |
| ENSG00000154447 | SH3RF1 | miR-29c-3p target gene only | Experiment supported target gene (TarBase v.8) |
| ENSG00000154451 | GBP5 | miR-29c-3p target gene only | Experiment supported target gene (TarBase v.8) |
| ENSG00000154639 | CXADR | miR-29c-3p target gene only | Experiment supported target gene (TarBase v.8) |
| ENSG00000154642 | C21orf91 | miR-29c-3p target gene only | Experiment supported target gene (TarBase v.8) |
| ENSG00000154845 | PPP4R1 | miR-29c-3p target gene only | Experiment supported target gene (TarBase v.8) |
| ENSG00000154864 | PIEZO2 | miR-29a-3p target gene only | Experiment supported target gene (TarBase v.8) |
| ENSG00000155008 | APOOL | miR-29a/c-3p common target gene | Experiment supported target gene (TarBase v.8) |
| ENSG00000155096 | AZIN1 | miR-29a-3p target gene only | Experiment supported target gene (TarBase v.8) |
| ENSG00000155099 | TMEM55A | miR-29c-3p target gene only | Experiment supported target gene (TarBase v.8) |
| ENSG00000155115 | GTF3C6 | miR-29c-3p target gene only | Experiment supported target gene (TarBase v.8) |
| ENSG00000155189 | AGPAT5 | miR-29a/c-3p common target gene | Experiment supported target gene (TarBase v.8) |
| ENSG00000155254 | MARVELD1 | miR-29c-3p target gene only | Experiment supported target gene (TarBase v.8) |
| ENSG00000155256 | ZFYVE27 | miR-29a/c-3p common target gene | Experiment supported target gene (TarBase v.8) |
| ENSG00000155330 | C16orf87 | miR-29a/c-3p common target gene | Experiment supported target gene (TarBase v.8) |
| ENSG00000155380 | SLC16A1 | miR-29a/c-3p common target gene | Experiment supported target gene (TarBase v.8) |
| ENSG00000155465 | SLC7A7 | miR-29c-3p target gene only | Experiment supported target gene (TarBase v.8) |
| ENSG00000155508 | CNOT8 | miR-29a/c-3p common target gene | Experiment supported target gene (TarBase v.8) |
| ENSG00000155744 | FAM126B | miR-29a/c-3p common target gene | Experiment supported target gene (TarBase v.8) |
| ENSG00000155755 | TMEM237 | miR-29a-3p target gene only | Experiment supported target gene (TarBase v.8) |
| ENSG00000155792 | DEPTOR | miR-29c-3p target gene only | Experiment supported target gene (TarBase v.8) |
| ENSG00000155858 | LSM11 | miR-29a/c-3p common target gene | Theoretical prediction (microT-CDS algorithm) |
| ENSG00000155876 | RRAGA | miR-29a/c-3p common target gene | Experiment supported target gene (TarBase v.8) |
| ENSG00000155893 | PXYLP1 | miR-29a/c-3p common target gene | Experiment supported target gene (TarBase v.8) |
| ENSG00000155918 | RAET1L | miR-29a/c-3p common target gene | Theoretical prediction (microT-CDS algorithm) |
| ENSG00000155974 | GRIP1 | miR-29a/c-3p common target gene | Theoretical prediction (microT-CDS algorithm) |
| ENSG00000155980 | KIF5A | miR-29a/c-3p common target gene | Theoretical prediction (microT-CDS algorithm) |
| ENSG00000156026 | MCU | miR-29a/c-3p common target gene | Experiment supported target gene (TarBase v.8) |
| ENSG00000156030 | ELMSAN1 | miR-29a/c-3p common target gene | Experiment supported target gene (TarBase v.8) |
| ENSG00000156103 | MMP16 | miR-29a/c-3p common target gene | Experiment supported target gene (TarBase v.8) |
| ENSG00000156113 | KCNMA1 | miR-29a/c-3p common target gene | Theoretical prediction (microT-CDS algorithm) |
| ENSG00000156140 | ADAMTS3 | miR-29a/c-3p common target gene | Theoretical prediction (microT-CDS algorithm) |
| ENSG00000156239 | N6AMT1 | miR-29c-3p target gene only | Experiment supported target gene (TarBase v.8) |
| ENSG00000156299 | TIAM1 | miR-29a/c-3p common target gene | Theoretical prediction (microT-CDS algorithm) |
| ENSG00000156345 | CDK20 | miR-29c-3p target gene only | Experiment supported target gene (TarBase v.8) |
| ENSG00000156374 | PCGF6 | miR-29c-3p target gene only | Experiment supported target gene (TarBase v.8) |
| ENSG00000156384 | SFR1 | miR-29c-3p target gene only | Experiment supported target gene (TarBase v.8) |
| ENSG00000156427 | FGF18 | miR-29c-3p target gene only | Experiment supported target gene (TarBase v.8) |
| ENSG00000156504 | FAM122B | miR-29a/c-3p common target gene | Experiment supported target gene (TarBase v.8) |
| ENSG00000156587 | UBE2L6 | miR-29a-3p target gene only | Theoretical prediction (microT-CDS algorithm) |
| ENSG00000156599 | ZDHHC5 | miR-29a/c-3p common target gene | Experiment supported target gene (TarBase v.8) |
| ENSG00000156650 | KAT6B | miR-29a/c-3p common target gene | Experiment supported target gene (TarBase v.8) |
| ENSG00000156802 | ATAD2 | miR-29a/c-3p common target gene | Experiment supported target gene (TarBase v.8) |
| ENSG00000156804 | FBXO32 | miR-29a-3p target gene only | Experiment supported target gene (TarBase v.8) |
| ENSG00000156853 | ZNF689 | miR-29c-3p target gene only | Experiment supported target gene (TarBase v.8) |
| ENSG00000156876 | SASS6 | miR-29c-3p target gene only | Experiment supported target gene (TarBase v.8) |
| ENSG00000156976 | EIF4A2 | miR-29a/c-3p common target gene | Experiment supported target gene (TarBase v.8) |
| ENSG00000157111 | TMEM171 | miR-29c-3p target gene only | Experiment supported target gene (TarBase v.8) |
| ENSG00000157168 | NRG1 | miR-29a-3p target gene only | Experiment supported target gene (TarBase v.8) |
| ENSG00000157184 | CPT2 | miR-29a/c-3p common target gene | Experiment supported target gene (TarBase v.8) |
| ENSG00000157193 | LRP8 | miR-29a/c-3p common target gene | Experiment supported target gene (TarBase v.8) |

| | | | |
|---|---|---|---|
| ENSG00000157211 | CDCP2 | miR-29a/c-3p common target gene | Theoretical prediction (microT-CDS algorithm) |
| ENSG00000157227 | MMP14 | miR-29a/c-3p common target gene | Experiment supported target gene (TarBase v.8) |
| ENSG00000157303 | SUSD3 | miR-29c-3p target gene only | Experiment supported target gene (TarBase v.8) |
| ENSG00000157456 | CCNB2 | miR-29a-3p target gene only | Experiment supported target gene (TarBase v.8) |
| ENSG00000157483 | MYO1E | miR-29a/c-3p common target gene | Theoretical prediction (microT-CDS algorithm) |
| ENSG00000157510 | AFAP1L1 | miR-29a/c-3p common target gene | Theoretical prediction (microT-CDS algorithm) |
| ENSG00000157514 | TSC22D3 | miR-29a/c-3p common target gene | Experiment supported target gene (TarBase v.8) |
| ENSG00000157542 | KCNJ6 | miR-29a/c-3p common target gene | Experiment supported target gene (TarBase v.8) |
| ENSG00000157593 | SLC35B2 | miR-29a/c-3p common target gene | Experiment supported target gene (TarBase v.8) |
| ENSG00000157601 | MX1 | miR-29c-3p target gene only | Experiment supported target gene (TarBase v.8) |
| ENSG00000157654 | PALM2-AKAP2 | miR-29a/c-3p common target gene | Experiment supported target gene (TarBase v.8) |
| ENSG00000157851 | DPYSL5 | miR-29a/c-3p common target gene | Experiment supported target gene (TarBase v.8) |
| ENSG00000157873 | TNFRSF14 | miR-29c-3p target gene only | Experiment supported target gene (TarBase v.8) |
| ENSG00000157916 | RER1 | miR-29c-3p target gene only | Experiment supported target gene (TarBase v.8) |
| ENSG00000157933 | SKI | miR-29a/c-3p common target gene | Experiment supported target gene (TarBase v.8) |
| ENSG00000157978 | LDLRAP1 | miR-29a/c-3p common target gene | Experiment supported target gene (TarBase v.8) |
| ENSG00000158022 | TRIM63 | miR-29a/c-3p common target gene | Theoretical prediction (microT-CDS algorithm) |
| ENSG00000158042 | MRPL17 | miR-29a/c-3p common target gene | Experiment supported target gene (TarBase v.8) |
| ENSG00000158050 | DUSP2 | miR-29a/c-3p common target gene | Experiment supported target gene (TarBase v.8) |
| ENSG00000158109 | TPRG1L | miR-29c-3p target gene only | Experiment supported target gene (TarBase v.8) |
| ENSG00000158125 | XDH | miR-29a/c-3p common target gene | Theoretical prediction (microT-CDS algorithm) |
| ENSG00000158169 | FANCC | miR-29a/c-3p common target gene | Experiment supported target gene (TarBase v.8) |
| ENSG00000158195 | WASF2 | miR-29a/c-3p common target gene | Experiment supported target gene (TarBase v.8) |
| ENSG00000158301 | GPRASP2 | miR-29a-3p target gene only | Experiment supported target gene (TarBase v.8) |
| ENSG00000158373 | HIST1H2BD | miR-29a-3p target gene only | Experiment supported target gene (TarBase v.8) |
| ENSG00000158445 | KCNB1 | miR-29a/c-3p common target gene | Experiment supported target gene (TarBase v.8) |
| ENSG00000158470 | B4GALT5 | miR-29a/c-3p common target gene | Experiment supported target gene (TarBase v.8) |
| ENSG00000158615 | PPP1R15B | miR-29a/c-3p common target gene | Experiment supported target gene (TarBase v.8) |
| ENSG00000158623 | COPG2 | miR-29c-3p target gene only | Experiment supported target gene (TarBase v.8) |
| ENSG00000158769 | F11R | miR-29a/c-3p common target gene | Experiment supported target gene (TarBase v.8) |
| ENSG00000158863 | FAM160B2 | miR-29a-3p target gene only | Experiment supported target gene (TarBase v.8) |
| ENSG00000158941 | CCAR2 | miR-29c-3p target gene only | Experiment supported target gene (TarBase v.8) |
| ENSG00000159111 | MRPL10 | miR-29c-3p target gene only | Experiment supported target gene (TarBase v.8) |
| ENSG00000159140 | SON | miR-29a/c-3p common target gene | Experiment supported target gene (TarBase v.8) |
| ENSG00000159167 | STC1 | miR-29a-3p target gene only | Experiment supported target gene (TarBase v.8) |
| ENSG00000159184 | HOXB13 | miR-29c-3p target gene only | Experiment supported target gene (TarBase v.8) |
| ENSG00000159199 | ATP5G1 | miR-29a/c-3p common target gene | Experiment supported target gene (TarBase v.8) |
| ENSG00000159214 | CCDC24 | miR-29c-3p target gene only | Experiment supported target gene (TarBase v.8) |
| ENSG00000159388 | BTG2 | miR-29a/c-3p common target gene | Experiment supported target gene (TarBase v.8) |
| ENSG00000159479 | MED8 | miR-29a/c-3p common target gene | Experiment supported target gene (TarBase v.8) |
| ENSG00000159579 | RSPRY1 | miR-29a/c-3p common target gene | Experiment supported target gene (TarBase v.8) |
| ENSG00000159792 | PSKH1 | miR-29a/c-3p common target gene | Experiment supported target gene (TarBase v.8) |
| ENSG00000159842 | ABR | miR-29a-3p target gene only | Experiment supported target gene (TarBase v.8) |
| ENSG00000159873 | CCDC117 | miR-29a/c-3p common target gene | Experiment supported target gene (TarBase v.8) |
| ENSG00000160007 | ARHGAP35 | miR-29a/c-3p common target gene | Experiment supported target gene (TarBase v.8) |
| ENSG00000160014 | CALM3 | miR-29a/c-3p common target gene | Experiment supported target gene (TarBase v.8) |
| ENSG00000160050 | CCDC28B | miR-29c-3p target gene only | Experiment supported target gene (TarBase v.8) |
| ENSG00000160094 | ZNF362 | miR-29a/c-3p common target gene | Theoretical prediction (microT-CDS algorithm) |
| ENSG00000160097 | FNDC5 | miR-29a/c-3p common target gene | Theoretical prediction (microT-CDS algorithm) |
| ENSG00000160124 | CCDC58 | miR-29c-3p target gene only | Experiment supported target gene (TarBase v.8) |
| ENSG00000160131 | VMA21 | miR-29a-3p target gene only | Experiment supported target gene (TarBase v.8) |
| ENSG00000160161 | CILP2 | miR-29a/c-3p common target gene | Theoretical prediction (microT-CDS algorithm) |
| ENSG00000160183 | TMPRSS3 | miR-29a/c-3p common target gene | Theoretical prediction (microT-CDS algorithm) |
| ENSG00000160190 | SLC37A1 | miR-29a/c-3p common target gene | Experiment supported target gene (TarBase v.8) |
| ENSG00000160193 | WDR4 | miR-29a/c-3p common target gene | Experiment supported target gene (TarBase v.8) |
| ENSG00000160216 | AGPAT3 | miR-29a-3p target gene only | Experiment supported target gene (TarBase v.8) |
| ENSG00000160305 | DIP2A | miR-29a-3p target gene only | Experiment supported target gene (TarBase v.8) |
| ENSG00000160310 | PRMT2 | miR-29a/c-3p common target gene | Experiment supported target gene (TarBase v.8) |
| ENSG00000160408 | ST6GALNAC6 | miR-29c-3p target gene only | Experiment supported target gene (TarBase v.8) |
| ENSG00000160446 | ZDHHC12 | miR-29c-3p target gene only | Experiment supported target gene (TarBase v.8) |
| ENSG00000160460 | SPTBN4 | miR-29a/c-3p common target gene | Experiment supported target gene (TarBase v.8) |
| ENSG00000160551 | TAOK1 | miR-29a/c-3p common target gene | Experiment supported target gene (TarBase v.8) |

| | | | |
|---|---|---|---|
| ENSG00000160588 | MPZL3 | miR-29a/c-3p common target gene | Experiment supported target gene (TarBase v.8) |
| ENSG00000160679 | CHTOP | miR-29a/c-3p common target gene | Experiment supported target gene (TarBase v.8) |
| ENSG00000160703 | NLRX1 | miR-29a/c-3p common target gene | Experiment supported target gene (TarBase v.8) |
| ENSG00000160714 | UBE2Q1 | miR-29a/c-3p common target gene | Theoretical prediction (microT-CDS algorithm) |
| ENSG00000160716 | CHRNB2 | miR-29a/c-3p common target gene | Experiment supported target gene (TarBase v.8) |
| ENSG00000160752 | FDPS | miR-29a-3p target gene only | Experiment supported target gene (TarBase v.8) |
| ENSG00000160753 | RUSC1 | miR-29a/c-3p common target gene | Experiment supported target gene (TarBase v.8) |
| ENSG00000160796 | NBEAL2 | miR-29a-3p target gene only | Experiment supported target gene (TarBase v.8) |
| ENSG00000160868 | CYP3A4 | miR-29c-3p target gene only | Experiment supported target gene (TarBase v.8) |
| ENSG00000160870 | CYP3A7 | miR-29c-3p target gene only | Experiment supported target gene (TarBase v.8) |
| ENSG00000160888 | IER2 | miR-29a-3p target gene only | Experiment supported target gene (TarBase v.8) |
| ENSG00000161057 | PSMC2 | miR-29c-3p target gene only | Experiment supported target gene (TarBase v.8) |
| ENSG00000161202 | DVL3 | miR-29a/c-3p common target gene | Experiment supported target gene (TarBase v.8) |
| ENSG00000161203 | AP2M1 | miR-29a-3p target gene only | Experiment supported target gene (TarBase v.8) |
| ENSG00000161298 | ZNF382 | miR-29c-3p target gene only | Theoretical prediction (microT-CDS algorithm) |
| ENSG00000161551 | ZNF577 | miR-29a/c-3p common target gene | Theoretical prediction (microT-CDS algorithm) |
| ENSG00000161638 | ITGA5 | miR-29c-3p target gene only | Experiment supported target gene (TarBase v.8) |
| ENSG00000161813 | LARP4 | miR-29a/c-3p common target gene | Theoretical prediction (microT-CDS algorithm) |
| ENSG00000161904 | LEMD2 | miR-29c-3p target gene only | Experiment supported target gene (TarBase v.8) |
| ENSG00000161955 | TNFSF13 | miR-29c-3p target gene only | Experiment supported target gene (TarBase v.8) |
| ENSG00000162063 | CCNF | miR-29a/c-3p common target gene | Experiment supported target gene (TarBase v.8) |
| ENSG00000162065 | TBC1D24 | miR-29a-3p target gene only | Experiment supported target gene (TarBase v.8) |
| ENSG00000162144 | CYB561A3 | miR-29a-3p target gene only | Experiment supported target gene (TarBase v.8) |
| ENSG00000162188 | GNG3 | miR-29c-3p target gene only | Experiment supported target gene (TarBase v.8) |
| ENSG00000162231 | NXF1 | miR-29a/c-3p common target gene | Experiment supported target gene (TarBase v.8) |
| ENSG00000162367 | TAL1 | miR-29a-3p target gene only | Experiment supported target gene (TarBase v.8) |
| ENSG00000162368 | CMPK1 | miR-29a/c-3p common target gene | Theoretical prediction (microT-CDS algorithm) |
| ENSG00000162390 | ACOT11 | miR-29a/c-3p common target gene | Experiment supported target gene (TarBase v.8) |
| ENSG00000162430 | SEPN1 | miR-29a-3p target gene only | Experiment supported target gene (TarBase v.8) |
| ENSG00000162433 | AK4 | miR-29a/c-3p common target gene | Theoretical prediction (microT-CDS algorithm) |
| ENSG00000162434 | JAK1 | miR-29a-3p target gene only | Experiment supported target gene (TarBase v.8) |
| ENSG00000162437 | RAVER2 | miR-29a-3p target gene only | Experiment supported target gene (TarBase v.8) |
| ENSG00000162441 | LZIC | miR-29a/c-3p common target gene | Experiment supported target gene (TarBase v.8) |
| ENSG00000162493 | PDPN | miR-29a/c-3p common target gene | Theoretical prediction (microT-CDS algorithm) |
| ENSG00000162522 | KIAA1522 | miR-29a/c-3p common target gene | Experiment supported target gene (TarBase v.8) |
| ENSG00000162542 | TMCO4 | miR-29c-3p target gene only | Experiment supported target gene (TarBase v.8) |
| ENSG00000162599 | NFIA | miR-29a/c-3p common target gene | Experiment supported target gene (TarBase v.8) |
| ENSG00000162613 | FUBP1 | miR-29a-3p target gene only | Experiment supported target gene (TarBase v.8) |
| ENSG00000162639 | HENMT1 | miR-29c-3p target gene only | Experiment supported target gene (TarBase v.8) |
| ENSG00000162642 | C1orf52 | miR-29a/c-3p common target gene | Experiment supported target gene (TarBase v.8) |
| ENSG00000162654 | GBP4 | miR-29c-3p target gene only | Experiment supported target gene (TarBase v.8) |
| ENSG00000162688 | AGL | miR-29a/c-3p common target gene | Experiment supported target gene (TarBase v.8) |
| ENSG00000162695 | SLC30A7 | miR-29a/c-3p common target gene | Theoretical prediction (microT-CDS algorithm) |
| ENSG00000162704 | ARPC5 | miR-29c-3p target gene only | Experiment supported target gene (TarBase v.8) |
| ENSG00000162711 | NLRP3 | miR-29c-3p target gene only | Experiment supported target gene (TarBase v.8) |
| ENSG00000162734 | PEA15 | miR-29a/c-3p common target gene | Experiment supported target gene (TarBase v.8) |
| ENSG00000162755 | KLHDC9 | miR-29c-3p target gene only | Experiment supported target gene (TarBase v.8) |
| ENSG00000162769 | FLVCR1 | miR-29c-3p target gene only | Experiment supported target gene (TarBase v.8) |
| ENSG00000162814 | SPATA17 | miR-29c-3p target gene only | Experiment supported target gene (TarBase v.8) |
| ENSG00000162849 | KIF26B | miR-29a/c-3p common target gene | Experiment supported target gene (TarBase v.8) |
| ENSG00000162851 | TFB2M | miR-29c-3p target gene only | Experiment supported target gene (TarBase v.8) |
| ENSG00000162892 | IL24 | miR-29c-3p target gene only | Experiment supported target gene (TarBase v.8) |
| ENSG00000162923 | WDR26 | miR-29a/c-3p common target gene | Experiment supported target gene (TarBase v.8) |
| ENSG00000162924 | REL | miR-29a/c-3p common target gene | Experiment supported target gene (TarBase v.8) |
| ENSG00000162989 | KCNJ3 | miR-29a/c-3p common target gene | Experiment supported target gene (TarBase v.8) |
| ENSG00000163006 | CCDC138 | miR-29a/c-3p common target gene | Experiment supported target gene (TarBase v.8) |
| ENSG00000163013 | FBXO41 | miR-29a/c-3p common target gene | Experiment supported target gene (TarBase v.8) |
| ENSG00000163017 | ACTG2 | miR-29a-3p target gene only | Experiment supported target gene (TarBase v.8) |
| ENSG00000163032 | VSNL1 | miR-29a/c-3p common target gene | Theoretical prediction (microT-CDS algorithm) |
| ENSG00000163041 | H3F3A | miR-29c-3p target gene only | Experiment supported target gene (TarBase v.8) |
| ENSG00000163053 | SLC16A14 | miR-29a/c-3p common target gene | Experiment supported target gene (TarBase v.8) |
| ENSG00000163069 | SGCB | miR-29a/c-3p common target gene | Theoretical prediction (microT-CDS algorithm) |

| | | | |
|---|---|---|---|
| ENSG00000163155 | LYSMD1 | miR-29a/c-3p common target gene | Experiment supported target gene (TarBase v.8) |
| ENSG00000163156 | SCNM1 | miR-29c-3p target gene only | Experiment supported target gene (TarBase v.8) |
| ENSG00000163209 | SPRR3 | miR-29c-3p target gene only | Experiment supported target gene (TarBase v.8) |
| ENSG00000163249 | CCNYL1 | miR-29a/c-3p common target gene | Experiment supported target gene (TarBase v.8) |
| ENSG00000163251 | FZD5 | miR-29a/c-3p common target gene | Experiment supported target gene (TarBase v.8) |
| ENSG00000163320 | CGGBP1 | miR-29a/c-3p common target gene | Experiment supported target gene (TarBase v.8) |
| ENSG00000163328 | GPR155 | miR-29a-3p target gene only | Experiment supported target gene (TarBase v.8) |
| ENSG00000163347 | CLDN1 | miR-29a/c-3p common target gene | Experiment supported target gene (TarBase v.8) |
| ENSG00000163349 | HIPK1 | miR-29a/c-3p common target gene | Experiment supported target gene (TarBase v.8) |
| ENSG00000163359 | COL6A3 | miR-29a/c-3p common target gene | Experiment supported target gene (TarBase v.8) |
| ENSG00000163376 | KBTBD8 | miR-29c-3p target gene only | Experiment supported target gene (TarBase v.8) |
| ENSG00000163428 | LRRC58 | miR-29a/c-3p common target gene | Experiment supported target gene (TarBase v.8) |
| ENSG00000163430 | FSTL1 | miR-29a/c-3p common target gene | Experiment supported target gene (TarBase v.8) |
| ENSG00000163444 | TMEM183A | miR-29a/c-3p common target gene | Experiment supported target gene (TarBase v.8) |
| ENSG00000163449 | TMEM169 | miR-29a/c-3p common target gene | Theoretical prediction (microT-CDS algorithm) |
| ENSG00000163466 | ARPC2 | miR-29a/c-3p common target gene | Experiment supported target gene (TarBase v.8) |
| ENSG00000163479 | SSR2 | miR-29c-3p target gene only | Experiment supported target gene (TarBase v.8) |
| ENSG00000163508 | EOMES | miR-29a/c-3p common target gene | Theoretical prediction (microT-CDS algorithm) |
| ENSG00000163513 | TGFBR2 | miR-29a-3p target gene only | Experiment supported target gene (TarBase v.8) |
| ENSG00000163527 | STT3B | miR-29a-3p target gene only | Experiment supported target gene (TarBase v.8) |
| ENSG00000163558 | PRKCI | miR-29a/c-3p common target gene | Experiment supported target gene (TarBase v.8) |
| ENSG00000163565 | IFI16 | miR-29a/c-3p common target gene | Experiment supported target gene (TarBase v.8) |
| ENSG00000163568 | AIM2 | miR-29c-3p target gene only | Experiment supported target gene (TarBase v.8) |
| ENSG00000163577 | EIF5A2 | miR-29a/c-3p common target gene | Experiment supported target gene (TarBase v.8) |
| ENSG00000163586 | FABP1 | miR-29a/c-3p common target gene | Theoretical prediction (microT-CDS algorithm) |
| ENSG00000163600 | ICOS | miR-29a/c-3p common target gene | Theoretical prediction (microT-CDS algorithm) |
| ENSG00000163625 | WDFY3 | miR-29a/c-3p common target gene | Experiment supported target gene (TarBase v.8) |
| ENSG00000163630 | SYNPR | miR-29a/c-3p common target gene | Experiment supported target gene (TarBase v.8) |
| ENSG00000163638 | ADAMTS9 | miR-29a/c-3p common target gene | Theoretical prediction (microT-CDS algorithm) |
| ENSG00000163655 | GMPS | miR-29c-3p target gene only | Experiment supported target gene (TarBase v.8) |
| ENSG00000163659 | TIPARP | miR-29a-3p target gene only | Experiment supported target gene (TarBase v.8) |
| ENSG00000163660 | CCNL1 | miR-29a/c-3p common target gene | Theoretical prediction (microT-CDS algorithm) |
| ENSG00000163661 | PTX3 | miR-29a/c-3p common target gene | Experiment supported target gene (TarBase v.8) |
| ENSG00000163704 | PRRT3 | miR-29c-3p target gene only | Experiment supported target gene (TarBase v.8) |
| ENSG00000163739 | CXCL1 | miR-29c-3p target gene only | Experiment supported target gene (TarBase v.8) |
| ENSG00000163762 | TM4SF18 | miR-29a/c-3p common target gene | Experiment supported target gene (TarBase v.8) |
| ENSG00000163788 | SNRK | miR-29a/c-3p common target gene | Experiment supported target gene (TarBase v.8) |
| ENSG00000163798 | SLC4A1AP | miR-29c-3p target gene only | Experiment supported target gene (TarBase v.8) |
| ENSG00000163820 | FYCO1 | miR-29c-3p target gene only | Experiment supported target gene (TarBase v.8) |
| ENSG00000163840 | DTX3L | miR-29c-3p target gene only | Experiment supported target gene (TarBase v.8) |
| ENSG00000163848 | ZNF148 | miR-29a/c-3p common target gene | Theoretical prediction (microT-CDS algorithm) |
| ENSG00000163864 | NMNAT3 | miR-29c-3p target gene only | Experiment supported target gene (TarBase v.8) |
| ENSG00000163877 | SNIP1 | miR-29a/c-3p common target gene | Experiment supported target gene (TarBase v.8) |
| ENSG00000163898 | LIPH | miR-29a-3p target gene only | Experiment supported target gene (TarBase v.8) |
| ENSG00000163902 | RPN1 | miR-29a-3p target gene only | Experiment supported target gene (TarBase v.8) |
| ENSG00000163909 | HEYL | miR-29c-3p target gene only | Experiment supported target gene (TarBase v.8) |
| ENSG00000163913 | IFT122 | miR-29c-3p target gene only | Theoretical prediction (microT-CDS algorithm) |
| ENSG00000163931 | TKT | miR-29a/c-3p common target gene | Experiment supported target gene (TarBase v.8) |
| ENSG00000163946 | FAM208A | miR-29a/c-3p common target gene | Experiment supported target gene (TarBase v.8) |
| ENSG00000163950 | SLBP | miR-29a-3p target gene only | Experiment supported target gene (TarBase v.8) |
| ENSG00000163960 | UBXN7 | miR-29a/c-3p common target gene | Theoretical prediction (microT-CDS algorithm) |
| ENSG00000163961 | RNF168 | miR-29a/c-3p common target gene | Experiment supported target gene (TarBase v.8) |
| ENSG00000163995 | ABLIM2 | miR-29c-3p target gene only | Experiment supported target gene (TarBase v.8) |
| ENSG00000164023 | SGMS2 | miR-29a/c-3p common target gene | Theoretical prediction (microT-CDS algorithm) |
| ENSG00000164032 | H2AFZ | miR-29c-3p target gene only | Experiment supported target gene (TarBase v.8) |
| ENSG00000164056 | SPRY1 | miR-29a/c-3p common target gene | Experiment supported target gene (TarBase v.8) |
| ENSG00000164066 | INTU | miR-29a-3p target gene only | Experiment supported target gene (TarBase v.8) |
| ENSG00000164068 | RNF123 | miR-29a/c-3p common target gene | Experiment supported target gene (TarBase v.8) |
| ENSG00000164096 | C4orf3 | miR-29c-3p target gene only | Experiment supported target gene (TarBase v.8) |
| ENSG00000164104 | HMGB2 | miR-29c-3p target gene only | Experiment supported target gene (TarBase v.8) |
| ENSG00000164111 | ANXA5 | miR-29a/c-3p common target gene | Experiment supported target gene (TarBase v.8) |
| ENSG00000164123 | C4orf45 | miR-29a/c-3p common target gene | Theoretical prediction (microT-CDS algorithm) |

| | | | |
|---|---|---|---|
| ENSG00000164151 | ICE1 | miR-29a-3p target gene only | Experiment supported target gene (TarBase v.8) |
| ENSG00000164163 | ABCE1 | miR-29a/c-3p common target gene | Theoretical prediction (microT-CDS algorithm) |
| ENSG00000164164 | OTUD4 | miR-29a/c-3p common target gene | Experiment supported target gene (TarBase v.8) |
| ENSG00000164171 | ITGA2 | miR-29a-3p target gene only | Experiment supported target gene (TarBase v.8) |
| ENSG00000164211 | STARD4 | miR-29c-3p target gene only | Experiment supported target gene (TarBase v.8) |
| ENSG00000164253 | WDR41 | miR-29a/c-3p common target gene | Experiment supported target gene (TarBase v.8) |
| ENSG00000164270 | HTR4 | miR-29c-3p target gene only | Experiment supported target gene (TarBase v.8) |
| ENSG00000164284 | GRPEL2 | miR-29a/c-3p common target gene | Theoretical prediction (microT-CDS algorithm) |
| ENSG00000164305 | CASP3 | miR-29c-3p target gene only | Experiment supported target gene (TarBase v.8) |
| ENSG00000164307 | ERAP1 | miR-29a/c-3p common target gene | Experiment supported target gene (TarBase v.8) |
| ENSG00000164309 | CMYA5 | miR-29a/c-3p common target gene | Theoretical prediction (microT-CDS algorithm) |
| ENSG00000164327 | RICTOR | miR-29a/c-3p common target gene | Experiment supported target gene (TarBase v.8) |
| ENSG00000164366 | CCDC127 | miR-29a/c-3p common target gene | Experiment supported target gene (TarBase v.8) |
| ENSG00000164400 | CSF2 | miR-29c-3p target gene only | Experiment supported target gene (TarBase v.8) |
| ENSG00000164488 | DACT2 | miR-29c-3p target gene only | Experiment supported target gene (TarBase v.8) |
| ENSG00000164542 | KIAA0895 | miR-29a/c-3p common target gene | Experiment supported target gene (TarBase v.8) |
| ENSG00000164574 | GALNT10 | miR-29a/c-3p common target gene | Theoretical prediction (microT-CDS algorithm) |
| ENSG00000164576 | SAP30L | miR-29a/c-3p common target gene | Theoretical prediction (microT-CDS algorithm) |
| ENSG00000164587 | RPS14 | miR-29c-3p target gene only | Experiment supported target gene (TarBase v.8) |
| ENSG00000164600 | NEUROD6 | miR-29c-3p target gene only | Experiment supported target gene (TarBase v.8) |
| ENSG00000164604 | GPR85 | miR-29a/c-3p common target gene | Theoretical prediction (microT-CDS algorithm) |
| ENSG00000164619 | BMPER | miR-29a/c-3p common target gene | Experiment supported target gene (TarBase v.8) |
| ENSG00000164626 | KCNK5 | miR-29c-3p target gene only | Experiment supported target gene (TarBase v.8) |
| ENSG00000164663 | USP49 | miR-29a/c-3p common target gene | Theoretical prediction (microT-CDS algorithm) |
| ENSG00000164675 | IQUB | miR-29a/c-3p common target gene | Theoretical prediction (microT-CDS algorithm) |
| ENSG00000164684 | ZNF704 | miR-29a/c-3p common target gene | Theoretical prediction (microT-CDS algorithm) |
| ENSG00000164692 | COL1A2 | miR-29a/c-3p common target gene | Experiment supported target gene (TarBase v.8) |
| ENSG00000164749 | HNF4G | miR-29a/c-3p common target gene | Theoretical prediction (microT-CDS algorithm) |
| ENSG00000164761 | TNFRSF11B | miR-29c-3p target gene only | Experiment supported target gene (TarBase v.8) |
| ENSG00000164828 | SUN1 | miR-29a/c-3p common target gene | Experiment supported target gene (TarBase v.8) |
| ENSG00000164830 | OXR1 | miR-29a/c-3p common target gene | Theoretical prediction (microT-CDS algorithm) |
| ENSG00000164889 | SLC4A2 | miR-29a-3p target gene only | Experiment supported target gene (TarBase v.8) |
| ENSG00000164916 | FOXK1 | miR-29a-3p target gene only | Experiment supported target gene (TarBase v.8) |
| ENSG00000164938 | TP53INP1 | miR-29a/c-3p common target gene | Experiment supported target gene (TarBase v.8) |
| ENSG00000164946 | FREM1 | miR-29a/c-3p common target gene | Theoretical prediction (microT-CDS algorithm) |
| ENSG00000164951 | PDP1 | miR-29a/c-3p common target gene | Experiment supported target gene (TarBase v.8) |
| ENSG00000164983 | TMEM65 | miR-29a/c-3p common target gene | Experiment supported target gene (TarBase v.8) |
| ENSG00000165028 | NIPSNAP3B | miR-29a/c-3p common target gene | Theoretical prediction (microT-CDS algorithm) |
| ENSG00000165066 | NKX6-3 | miR-29a/c-3p common target gene | Theoretical prediction (microT-CDS algorithm) |
| ENSG00000165072 | MAMDC2 | miR-29a-3p target gene only | Experiment supported target gene (TarBase v.8) |
| ENSG00000165124 | SVEP1 | miR-29a/c-3p common target gene | Theoretical prediction (microT-CDS algorithm) |
| ENSG00000165195 | PIGA | miR-29a/c-3p common target gene | Experiment supported target gene (TarBase v.8) |
| ENSG00000165282 | PIGO | miR-29a-3p target gene only | Experiment supported target gene (TarBase v.8) |
| ENSG00000165288 | BRWD3 | miR-29a/c-3p common target gene | Experiment supported target gene (TarBase v.8) |
| ENSG00000165309 | ARMC3 | miR-29c-3p target gene only | Experiment supported target gene (TarBase v.8) |
| ENSG00000165312 | OTUD1 | miR-29a-3p target gene only | Experiment supported target gene (TarBase v.8) |
| ENSG00000165323 | FAT3 | miR-29a/c-3p common target gene | Experiment supported target gene (TarBase v.8) |
| ENSG00000165389 | SPTSSA | miR-29c-3p target gene only | Experiment supported target gene (TarBase v.8) |
| ENSG00000165410 | CFL2 | miR-29a/c-3p common target gene | Theoretical prediction (microT-CDS algorithm) |
| ENSG00000165416 | SUGT1 | miR-29a-3p target gene only | Experiment supported target gene (TarBase v.8) |
| ENSG00000165443 | PHYHIPL | miR-29a/c-3p common target gene | Experiment supported target gene (TarBase v.8) |
| ENSG00000165449 | SLC16A9 | miR-29c-3p target gene only | Experiment supported target gene (TarBase v.8) |
| ENSG00000165475 | CRYL1 | miR-29c-3p target gene only | Experiment supported target gene (TarBase v.8) |
| ENSG00000165476 | REEP3 | miR-29c-3p target gene only | Experiment supported target gene (TarBase v.8) |
| ENSG00000165495 | PKNOX2 | miR-29a/c-3p common target gene | Theoretical prediction (microT-CDS algorithm) |
| ENSG00000165512 | ZNF22 | miR-29a-3p target gene only | Experiment supported target gene (TarBase v.8) |
| ENSG00000165521 | EML5 | miR-29a/c-3p common target gene | Theoretical prediction (microT-CDS algorithm) |
| ENSG00000165527 | ARF6 | miR-29a/c-3p common target gene | Experiment supported target gene (TarBase v.8) |
| ENSG00000165629 | ATP5C1 | miR-29a/c-3p common target gene | Experiment supported target gene (TarBase v.8) |
| ENSG00000165671 | NSD1 | miR-29a/c-3p common target gene | Experiment supported target gene (TarBase v.8) |
| ENSG00000165684 | SNAPC4 | miR-29c-3p target gene only | Experiment supported target gene (TarBase v.8) |
| ENSG00000165685 | TMEM52B | miR-29a-3p target gene only | Experiment supported target gene (TarBase v.8) |

| | | | |
|---|---|---|---|
| ENSG00000165716 | FAM69B | miR-29a-3p target gene only | Experiment supported target gene (TarBase v.8) |
| ENSG00000165724 | ZMYND19 | miR-29c-3p target gene only | Experiment supported target gene (TarBase v.8) |
| ENSG00000165732 | DDX21 | miR-29a/c-3p common target gene | Experiment supported target gene (TarBase v.8) |
| ENSG00000165806 | CASP7 | miR-29a/c-3p common target gene | Experiment supported target gene (TarBase v.8) |
| ENSG00000165821 | SALL2 | miR-29c-3p target gene only | Experiment supported target gene (TarBase v.8) |
| ENSG00000165891 | E2F7 | miR-29a/c-3p common target gene | Experiment supported target gene (TarBase v.8) |
| ENSG00000165915 | SLC39A13 | miR-29a/c-3p common target gene | Theoretical prediction (microT-CDS algorithm) |
| ENSG00000165949 | IFI27 | miR-29c-3p target gene only | Experiment supported target gene (TarBase v.8) |
| ENSG00000165959 | CLMN | miR-29a/c-3p common target gene | Experiment supported target gene (TarBase v.8) |
| ENSG00000165972 | CCDC38 | miR-29a/c-3p common target gene | Theoretical prediction (microT-CDS algorithm) |
| ENSG00000165983 | PTER | miR-29c-3p target gene only | Experiment supported target gene (TarBase v.8) |
| ENSG00000165985 | C1QL3 | miR-29a/c-3p common target gene | Experiment supported target gene (TarBase v.8) |
| ENSG00000165997 | ARL5B | miR-29a/c-3p common target gene | Experiment supported target gene (TarBase v.8) |
| ENSG00000166006 | KCNC2 | miR-29a/c-3p common target gene | Theoretical prediction (microT-CDS algorithm) |
| ENSG00000166035 | LIPC | miR-29c-3p target gene only | Experiment supported target gene (TarBase v.8) |
| ENSG00000166106 | ADAMTS15 | miR-29a-3p target gene only | Experiment supported target gene (TarBase v.8) |
| ENSG00000166123 | GPT2 | miR-29c-3p target gene only | Experiment supported target gene (TarBase v.8) |
| ENSG00000166145 | SPINT1 | miR-29a/c-3p common target gene | Experiment supported target gene (TarBase v.8) |
| ENSG00000166147 | FBN1 | miR-29a/c-3p common target gene | Experiment supported target gene (TarBase v.8) |
| ENSG00000166165 | CKB | miR-29a/c-3p common target gene | Experiment supported target gene (TarBase v.8) |
| ENSG00000166225 | FRS2 | miR-29a/c-3p common target gene | Experiment supported target gene (TarBase v.8) |
| ENSG00000166272 | WBP1L | miR-29a/c-3p common target gene | Experiment supported target gene (TarBase v.8) |
| ENSG00000166326 | TRIM44 | miR-29a/c-3p common target gene | Experiment supported target gene (TarBase v.8) |
| ENSG00000166341 | DCHS1 | miR-29a-3p target gene only | Experiment supported target gene (TarBase v.8) |
| ENSG00000166396 | SERPINB7 | miR-29a-3p target gene only | Experiment supported target gene (TarBase v.8) |
| ENSG00000166398 | KIAA0355 | miR-29a/c-3p common target gene | Experiment supported target gene (TarBase v.8) |
| ENSG00000166402 | TUB | miR-29a/c-3p common target gene | Experiment supported target gene (TarBase v.8) |
| ENSG00000166405 | RIC3 | miR-29a/c-3p common target gene | Theoretical prediction (microT-CDS algorithm) |
| ENSG00000166439 | RNF169 | miR-29a-3p target gene only | Experiment supported target gene (TarBase v.8) |
| ENSG00000166452 | AKIP1 | miR-29c-3p target gene only | Experiment supported target gene (TarBase v.8) |
| ENSG00000166483 | WEE1 | miR-29a/c-3p common target gene | Experiment supported target gene (TarBase v.8) |
| ENSG00000166509 | CLEC3A | miR-29a/c-3p common target gene | Theoretical prediction (microT-CDS algorithm) |
| ENSG00000166578 | IQCD | miR-29c-3p target gene only | Experiment supported target gene (TarBase v.8) |
| ENSG00000166664 | CHRFAM7A | miR-29a/c-3p common target gene | Theoretical prediction (microT-CDS algorithm) |
| ENSG00000166741 | NNMT | miR-29a/c-3p common target gene | Experiment supported target gene (TarBase v.8) |
| ENSG00000166747 | AP1G1 | miR-29a/c-3p common target gene | Experiment supported target gene (TarBase v.8) |
| ENSG00000166780 | C16orf45 | miR-29c-3p target gene only | Experiment supported target gene (TarBase v.8) |
| ENSG00000166797 | FAM96A | miR-29c-3p target gene only | Experiment supported target gene (TarBase v.8) |
| ENSG00000166816 | LDHD | miR-29c-3p target gene only | Experiment supported target gene (TarBase v.8) |
| ENSG00000166833 | NAV2 | miR-29a/c-3p common target gene | Theoretical prediction (microT-CDS algorithm) |
| ENSG00000166847 | DCTN5 | miR-29a/c-3p common target gene | Experiment supported target gene (TarBase v.8) |
| ENSG00000166848 | TERF2IP | miR-29a/c-3p common target gene | Experiment supported target gene (TarBase v.8) |
| ENSG00000166851 | PLK1 | miR-29a/c-3p common target gene | Experiment supported target gene (TarBase v.8) |
| ENSG00000166902 | MRPL16 | miR-29c-3p target gene only | Experiment supported target gene (TarBase v.8) |
| ENSG00000166925 | TSC22D4 | miR-29a-3p target gene only | Experiment supported target gene (TarBase v.8) |
| ENSG00000166986 | MARS | miR-29a/c-3p common target gene | Theoretical prediction (microT-CDS algorithm) |
| ENSG00000166997 | CNPY4 | miR-29c-3p target gene only | Experiment supported target gene (TarBase v.8) |
| ENSG00000167103 | PIP5KL1 | miR-29a/c-3p common target gene | Theoretical prediction (microT-CDS algorithm) |
| ENSG00000167118 | URM1 | miR-29a/c-3p common target gene | Theoretical prediction (microT-CDS algorithm) |
| ENSG00000167130 | DOLPP1 | miR-29c-3p target gene only | Theoretical prediction (microT-CDS algorithm) |
| ENSG00000167178 | ISLR2 | miR-29a/c-3p common target gene | Theoretical prediction (microT-CDS algorithm) |
| ENSG00000167191 | GPRC5B | miR-29c-3p target gene only | Experiment supported target gene (TarBase v.8) |
| ENSG00000167220 | HDHD2 | miR-29c-3p target gene only | Experiment supported target gene (TarBase v.8) |
| ENSG00000167257 | RNF214 | miR-29a/c-3p common target gene | Theoretical prediction (microT-CDS algorithm) |
| ENSG00000167281 | RBFOX3 | miR-29a/c-3p common target gene | Theoretical prediction (microT-CDS algorithm) |
| ENSG00000167306 | MYO5B | miR-29c-3p target gene only | Experiment supported target gene (TarBase v.8) |
| ENSG00000167325 | RRM1 | miR-29a/c-3p common target gene | Experiment supported target gene (TarBase v.8) |
| ENSG00000167378 | IRGQ | miR-29a/c-3p common target gene | Experiment supported target gene (TarBase v.8) |
| ENSG00000167522 | ANKRD11 | miR-29a/c-3p common target gene | Experiment supported target gene (TarBase v.8) |
| ENSG00000167536 | DHRS13 | miR-29c-3p target gene only | Experiment supported target gene (TarBase v.8) |
| ENSG00000167566 | NCKAP5L | miR-29a/c-3p common target gene | Theoretical prediction (microT-CDS algorithm) |
| ENSG00000167580 | AQP2 | miR-29a/c-3p common target gene | Theoretical prediction (microT-CDS algorithm) |
| ENSG00000167601 | AXL | miR-29a/c-3p common target gene | Experiment supported target gene (TarBase v.8) |
| ENSG00000167615 | LENG8 | miR-29a/c-3p common target gene | Experiment supported target gene (TarBase v.8) |
| ENSG00000167619 | TMEM145 | miR-29c-3p target gene only | Experiment supported target gene (TarBase v.8) |
| ENSG00000167656 | LY6D | miR-29c-3p target gene only | Experiment supported target gene (TarBase v.8) |
| ENSG00000167658 | EEF2 | miR-29c-3p target gene only | Experiment supported target gene (TarBase v.8) |
| ENSG00000167664 | TMIGD2 | miR-29c-3p target gene only | Experiment supported target gene (TarBase v.8) |
| ENSG00000167670 | CHAF1A | miR-29c-3p target gene only | Experiment supported target gene (TarBase v.8) |
| ENSG00000167702 | KIFC2 | miR-29a/c-3p common target gene | Theoretical prediction (microT-CDS algorithm) |
| ENSG00000167703 | SLC43A2 | miR-29a/c-3p common target gene | Experiment supported target gene (TarBase v.8) |
| ENSG00000167767 | KRT80 | miR-29a/c-3p common target gene | Experiment supported target gene (TarBase v.8) |
| ENSG00000167768 | KRT1 | miR-29c-3p target gene only | Experiment supported target gene (TarBase v.8) |
| ENSG00000167772 | ANGPTL4 | miR-29c-3p target gene only | Experiment supported target gene (TarBase v.8) |
| ENSG00000167778 | SPRYD3 | miR-29a-3p target gene only | Experiment supported target gene (TarBase v.8) |
| ENSG00000167861 | HID1 | miR-29a-3p target gene only | Experiment supported target gene (TarBase v.8) |
| ENSG00000167874 | TMEM88 | miR-29a-3p target gene only | Experiment supported target gene (TarBase v.8) |
| ENSG00000167977 | KCTD5 | miR-29a/c-3p common target gene | Experiment supported target gene (TarBase v.8) |
| ENSG00000167986 | DDB1 | miR-29a-3p target gene only | Experiment supported target gene (TarBase v.8) |
| ENSG00000167987 | VPS37C | miR-29a/c-3p common target gene | Experiment supported target gene (TarBase v.8) |
| ENSG00000167992 | VWCE | miR-29c-3p target gene only | Experiment supported target gene (TarBase v.8) |
| ENSG00000168010 | ATG16L2 | miR-29a/c-3p common target gene | Theoretical prediction (microT-CDS algorithm) |
| ENSG00000168036 | CTNNB1 | miR-29a/c-3p common target gene | Experiment supported target gene (TarBase v.8) |
| ENSG00000168062 | BATF2 | miR-29c-3p target gene only | Experiment supported target gene (TarBase v.8) |
| ENSG00000168101 | NUDT16L1 | miR-29a/c-3p common target gene | Experiment supported target gene (TarBase v.8) |
| ENSG00000168116 | KIAA1586 | miR-29a/c-3p common target gene | Experiment supported target gene (TarBase v.8) |
| ENSG00000168137 | SETD5 | miR-29c-3p target gene only | Experiment supported target gene (TarBase v.8) |
| ENSG00000168143 | FAM83B | miR-29a-3p target gene only | Experiment supported target gene (TarBase v.8) |
| ENSG00000168172 | HOOK3 | miR-29a/c-3p common target gene | Experiment supported target gene (TarBase v.8) |
| ENSG00000168175 | MAPK1IP1L | miR-29c-3p target gene only | Experiment supported target gene (TarBase v.8) |
| ENSG00000168246 | UBTD2 | miR-29a/c-3p common target gene | Experiment supported target gene (TarBase v.8) |
| ENSG00000168256 | NKIRAS2 | miR-29a/c-3p common target gene | Experiment supported target gene (TarBase v.8) |
| ENSG00000168291 | PDHB | miR-29c-3p target gene only | Experiment supported target gene (TarBase v.8) |
| ENSG00000168297 | PXK | miR-29c-3p target gene only | Experiment supported target gene (TarBase v.8) |
| ENSG00000168306 | ACOX2 | miR-29c-3p target gene only | Experiment supported target gene (TarBase v.8) |
| ENSG00000168314 | MOBP | miR-29a/c-3p common target gene | Theoretical prediction (microT-CDS algorithm) |
| ENSG00000168385 | 2-Sep | miR-29c-3p target gene only | Experiment supported target gene (TarBase v.8) |
| ENSG00000168394 | TAP1 | miR-29c-3p target gene only | Experiment supported target gene (TarBase v.8) |
| ENSG00000168398 | BDKRB2 | miR-29a/c-3p common target gene | Theoretical prediction (microT-CDS algorithm) |
| ENSG00000168411 | RFWD3 | miR-29c-3p target gene only | Experiment supported target gene (TarBase v.8) |
| ENSG00000168488 | ATXN2L | miR-29a-3p target gene only | Experiment supported target gene (TarBase v.8) |
| ENSG00000168495 | POLR3D | miR-29c-3p target gene only | Experiment supported target gene (TarBase v.8) |
| ENSG00000168517 | HEXIM2 | miR-29c-3p target gene only | Experiment supported target gene (TarBase v.8) |
| ENSG00000168539 | CHRM1 | miR-29c-3p target gene only | Experiment supported target gene (TarBase v.8) |
| ENSG00000168542 | COL3A1 | miR-29a/c-3p common target gene | Experiment supported target gene (TarBase v.8) |
| ENSG00000168575 | SLC20A2 | miR-29c-3p target gene only | Experiment supported target gene (TarBase v.8) |
| ENSG00000168615 | ADAM9 | miR-29a-3p target gene only | Experiment supported target gene (TarBase v.8) |
| ENSG00000168675 | LDLRAD4 | miR-29a/c-3p common target gene | Experiment supported target gene (TarBase v.8) |
| ENSG00000168679 | SLC16A4 | miR-29a-3p target gene only | Experiment supported target gene (TarBase v.8) |
| ENSG00000168710 | AHCYL1 | miR-29a/c-3p common target gene | Experiment supported target gene (TarBase v.8) |
| ENSG00000168769 | TET2 | miR-29a/c-3p common target gene | Experiment supported target gene (TarBase v.8) |
| ENSG00000168795 | ZBTB5 | miR-29a/c-3p common target gene | Experiment supported target gene (TarBase v.8) |
| ENSG00000168807 | SNTB2 | miR-29c-3p target gene only | Theoretical prediction (microT-CDS algorithm) |
| ENSG00000168813 | ZNF507 | miR-29a/c-3p common target gene | Experiment supported target gene (TarBase v.8) |
| ENSG00000168887 | C2orf68 | miR-29a/c-3p common target gene | Experiment supported target gene (TarBase v.8) |
| ENSG00000168913 | ENHO | miR-29a/c-3p common target gene | Theoretical prediction (microT-CDS algorithm) |
| ENSG00000168916 | ZNF608 | miR-29c-3p target gene only | Experiment supported target gene (TarBase v.8) |
| ENSG00000168918 | INPP5D | miR-29c-3p target gene only | Experiment supported target gene (TarBase v.8) |
| ENSG00000168938 | PPIC | miR-29a/c-3p common target gene | Experiment supported target gene (TarBase v.8) |
| ENSG00000168952 | STXBP6 | miR-29c-3p target gene only | Experiment supported target gene (TarBase v.8) |
| ENSG00000168994 | PXDC1 | miR-29c-3p target gene only | Experiment supported target gene (TarBase v.8) |
| ENSG00000169018 | FEM1B | miR-29a/c-3p common target gene | Experiment supported target gene (TarBase v.8) |
| ENSG00000169031 | COL4A3 | miR-29a/c-3p common target gene | Theoretical prediction (microT-CDS algorithm) |
| ENSG00000169083 | AR | miR-29a/c-3p common target gene | Theoretical prediction (microT-CDS algorithm) |
| ENSG00000169084 | DHRSX | miR-29c-3p target gene only | Experiment supported target gene (TarBase v.8) |
| ENSG00000169093 | ASMTL | miR-29a-3p target gene only | Experiment supported target gene (TarBase v.8) |
| ENSG00000169118 | CSNK1G1 | miR-29a/c-3p common target gene | Experiment supported target gene (TarBase v.8) |
| ENSG00000169155 | ZBTB43 | miR-29a/c-3p common target gene | Theoretical prediction (microT-CDS algorithm) |
| ENSG00000169174 | PCSK9 | miR-29c-3p target gene only | Experiment supported target gene (TarBase v.8) |
| ENSG00000169193 | CCDC126 | miR-29c-3p target gene only | Experiment supported target gene (TarBase v.8) |
| ENSG00000169248 | CXCL11 | miR-29c-3p target gene only | Experiment supported target gene (TarBase v.8) |
| ENSG00000169330 | KIAA1024 | miR-29a/c-3p common target gene | Theoretical prediction (microT-CDS algorithm) |
| ENSG00000169429 | CXCL8 | miR-29a-3p target gene only | Experiment supported target gene (TarBase v.8) |
| ENSG00000169436 | COL22A1 | miR-29a/c-3p common target gene | Theoretical prediction (microT-CDS algorithm) |
| ENSG00000169439 | SDC2 | miR-29a/c-3p common target gene | Experiment supported target gene (TarBase v.8) |
| ENSG00000169474 | SPRR1A | miR-29c-3p target gene only | Experiment supported target gene (TarBase v.8) |
| ENSG00000169490 | TM2D2 | miR-29a/c-3p common target gene | Experiment supported target gene (TarBase v.8) |
| ENSG00000169507 | SLC38A11 | miR-29c-3p target gene only | Experiment supported target gene (TarBase v.8) |
| ENSG00000169567 | HINT1 | miR-29a-3p target gene only | Experiment supported target gene (TarBase v.8) |
| ENSG00000169570 | DTWD2 | miR-29a/c-3p common target gene | Experiment supported target gene (TarBase v.8) |
| ENSG00000169599 | NFU1 | miR-29c-3p target gene only | Experiment supported target gene (TarBase v.8) |
| ENSG00000169641 | LUZP1 | miR-29a/c-3p common target gene | Experiment supported target gene (TarBase v.8) |
| ENSG00000169679 | BUB1 | miR-29a/c-3p common target gene | Experiment supported target gene (TarBase v.8) |
| ENSG00000169682 | SPNS1 | miR-29a/c-3p common target gene | Theoretical prediction (microT-CDS algorithm) |
| ENSG00000169710 | FASN | miR-29a/c-3p common target gene | Experiment supported target gene (TarBase v.8) |
| ENSG00000169714 | CNBP | miR-29c-3p target gene only | Experiment supported target gene (TarBase v.8) |
| ENSG00000169756 | LIMS1 | miR-29a/c-3p common target gene | Experiment supported target gene (TarBase v.8) |
| ENSG00000169760 | NLGN1 | miR-29a-3p target gene only | Experiment supported target gene (TarBase v.8) |
| ENSG00000169789 | PRY | miR-29a/c-3p common target gene | Theoretical prediction (microT-CDS algorithm) |
| ENSG00000169807 | PRY2 | miR-29a/c-3p common target gene | Theoretical prediction (microT-CDS algorithm) |
| ENSG00000169813 | HNRNPF | miR-29a/c-3p common target gene | Experiment supported target gene (TarBase v.8) |
| ENSG00000169826 | CSGALNACT2 | miR-29a/c-3p common target gene | Theoretical prediction (microT-CDS algorithm) |
| ENSG00000169855 | ROBO1 | miR-29a/c-3p common target gene | Experiment supported target gene (TarBase v.8) |
| ENSG00000169860 | P2RY1 | miR-29c-3p target gene only | Theoretical prediction (microT-CDS algorithm) |
| ENSG00000169891 | REPS2 | miR-29a/c-3p common target gene | Theoretical prediction (microT-CDS algorithm) |
| ENSG00000169925 | BRD3 | miR-29a/c-3p common target gene | Experiment supported target gene (TarBase v.8) |
| ENSG00000169964 | TMEM42 | miR-29c-3p target gene only | Experiment supported target gene (TarBase v.8) |
| ENSG00000169991 | IFFO2 | miR-29a/c-3p common target gene | Theoretical prediction (microT-CDS algorithm) |
| ENSG00000170027 | YWHAG | miR-29a/c-3p common target gene | Experiment supported target gene (TarBase v.8) |
| ENSG00000170153 | RNF150 | miR-29a/c-3p common target gene | Experiment supported target gene (TarBase v.8) |
| ENSG00000170265 | ZNF282 | miR-29a/c-3p common target gene | Experiment supported target gene (TarBase v.8) |
| ENSG00000170291 | ELP5 | miR-29c-3p target gene only | Theoretical prediction (microT-CDS algorithm) |
| ENSG00000170340 | B3GNT2 | miR-29a-3p target gene only | Experiment supported target gene (TarBase v.8) |
| ENSG00000170385 | SLC30A1 | miR-29a/c-3p common target gene | Experiment supported target gene (TarBase v.8) |
| ENSG00000170448 | NFXL1 | miR-29a/c-3p common target gene | Experiment supported target gene (TarBase v.8) |
| ENSG00000170456 | DENND5B | miR-29a/c-3p common target gene | Experiment supported target gene (TarBase v.8) |
| ENSG00000170468 | C14orf169 | miR-29c-3p target gene only | Experiment supported target gene (TarBase v.8) |
| ENSG00000170522 | ELOVL6 | miR-29a-3p target gene only | Experiment supported target gene (TarBase v.8) |
| ENSG00000170540 | ARL6IP1 | miR-29c-3p target gene only | Experiment supported target gene (TarBase v.8) |
| ENSG00000170542 | SERPINB9 | miR-29a/c-3p common target gene | Experiment supported target gene (TarBase v.8) |
| ENSG00000170606 | HSPA4 | miR-29a/c-3p common target gene | Experiment supported target gene (TarBase v.8) |
| ENSG00000170633 | RNF34 | miR-29a/c-3p common target gene | Experiment supported target gene (TarBase v.8) |
| ENSG00000170759 | KIF5B | miR-29a/c-3p common target gene | Experiment supported target gene (TarBase v.8) |
| ENSG00000170775 | GPR37 | miR-29a/c-3p common target gene | Experiment supported target gene (TarBase v.8) |
| ENSG00000170779 | CDCA4 | miR-29a/c-3p common target gene | Experiment supported target gene (TarBase v.8) |
| ENSG00000170802 | FOXN2 | miR-29c-3p target gene only | Experiment supported target gene (TarBase v.8) |
| ENSG00000170836 | PPM1D | miR-29a/c-3p common target gene | Experiment supported target gene (TarBase v.8) |
| ENSG00000170848 | PSG6 | miR-29c-3p target gene only | Experiment supported target gene (TarBase v.8) |
| ENSG00000170855 | TRIAP1 | miR-29c-3p target gene only | Experiment supported target gene (TarBase v.8) |
| ENSG00000170881 | RNF139 | miR-29c-3p target gene only | Experiment supported target gene (TarBase v.8) |
| ENSG00000170899 | GSTA4 | miR-29a/c-3p common target gene | Experiment supported target gene (TarBase v.8) |
| ENSG00000170917 | NUDT6 | miR-29c-3p target gene only | Experiment supported target gene (TarBase v.8) |
| ENSG00000170949 | ZNF160 | miR-29a/c-3p common target gene | Experiment supported target gene (TarBase v.8) |
| ENSG00000170961 | HAS2 | miR-29a/c-3p common target gene | Experiment supported target gene (TarBase v.8) |
| ENSG00000170962 | PDGFD | miR-29a-3p target gene only | Theoretical prediction (microT-CDS algorithm) |
| ENSG00000171044 | XKR6 | miR-29a/c-3p common target gene | Theoretical prediction (microT-CDS algorithm) |
| ENSG00000171135 | JAGN1 | miR-29a/c-3p common target gene | Experiment supported target gene (TarBase v.8) |
| ENSG00000171195 | MUC7 | miR-29a/c-3p common target gene | Theoretical prediction (microT-CDS algorithm) |
| ENSG00000171246 | NPTX1 | miR-29a-3p target gene only | Experiment supported target gene (TarBase v.8) |
| ENSG00000171346 | KRT15 | miR-29c-3p target gene only | Experiment supported target gene (TarBase v.8) |
| ENSG00000171365 | CLCN5 | miR-29a/c-3p common target gene | Theoretical prediction (microT-CDS algorithm) |
| ENSG00000171408 | PDE7B | miR-29a/c-3p common target gene | Experiment supported target gene (TarBase v.8) |
| ENSG00000171428 | NAT1 | miR-29a/c-3p common target gene | Theoretical prediction (microT-CDS algorithm) |
| ENSG00000171444 | MCC | miR-29a-3p target gene only | Theoretical prediction (microT-CDS algorithm) |
| ENSG00000171456 | ASXL1 | miR-29c-3p target gene only | Experiment supported target gene (TarBase v.8) |
| ENSG00000171467 | ZNF318 | miR-29a/c-3p common target gene | Experiment supported target gene (TarBase v.8) |
| ENSG00000171497 | PPID | miR-29a/c-3p common target gene | Experiment supported target gene (TarBase v.8) |
| ENSG00000171503 | ETFDH | miR-29c-3p target gene only | Experiment supported target gene (TarBase v.8) |
| ENSG00000171522 | PTGER4 | miR-29a-3p target gene only | Experiment supported target gene (TarBase v.8) |
| ENSG00000171533 | MAP6 | miR-29a/c-3p common target gene | Theoretical prediction (microT-CDS algorithm) |
| ENSG00000171557 | FGG | miR-29a/c-3p common target gene | Experiment supported target gene (TarBase v.8) |
| ENSG00000171560 | FGA | miR-29c-3p target gene only | Experiment supported target gene (TarBase v.8) |
| ENSG00000171564 | FGB | miR-29c-3p target gene only | Experiment supported target gene (TarBase v.8) |
| ENSG00000171612 | SLC25A33 | miR-29c-3p target gene only | Experiment supported target gene (TarBase v.8) |
| ENSG00000171680 | PLEKHG5 | miR-29c-3p target gene only | Experiment supported target gene (TarBase v.8) |
| ENSG00000171714 | ANO5 | miR-29a/c-3p common target gene | Experiment supported target gene (TarBase v.8) |
| ENSG00000171723 | GPHN | miR-29a/c-3p common target gene | Experiment supported target gene (TarBase v.8) |
| ENSG00000171791 | BCL2 | miR-29a/c-3p common target gene | Experiment supported target gene (TarBase v.8) |
| ENSG00000171848 | RRM2 | miR-29a-3p target gene only | Experiment supported target gene (TarBase v.8) |
| ENSG00000171855 | IFNB1 | miR-29c-3p target gene only | Experiment supported target gene (TarBase v.8) |
| ENSG00000171862 | PTEN | miR-29a/c-3p common target gene | Experiment supported target gene (TarBase v.8) |
| ENSG00000171988 | JMJD1C | miR-29a/c-3p common target gene | Experiment supported target gene (TarBase v.8) |
| ENSG00000172007 | RAB33B | miR-29a/c-3p common target gene | Experiment supported target gene (TarBase v.8) |
| ENSG00000172009 | THOP1 | miR-29a/c-3p common target gene | Experiment supported target gene (TarBase v.8) |
| ENSG00000172086 | KRCC1 | miR-29c-3p target gene only | Experiment supported target gene (TarBase v.8) |
| ENSG00000172115 | CYCS | miR-29a/c-3p common target gene | Experiment supported target gene (TarBase v.8) |
| ENSG00000172183 | ISG20 | miR-29a-3p target gene only | Experiment supported target gene (TarBase v.8) |
| ENSG00000172197 | MBOAT1 | miR-29c-3p target gene only | Experiment supported target gene (TarBase v.8) |
| ENSG00000172243 | CLEC7A | miR-29a/c-3p common target gene | Theoretical prediction (microT-CDS algorithm) |
| ENSG00000172260 | NEGR1 | miR-29a/c-3p common target gene | Experiment supported target gene (TarBase v.8) |
| ENSG00000172270 | BSG | miR-29a-3p target gene only | Experiment supported target gene (TarBase v.8) |
| ENSG00000172331 | BPGM | miR-29c-3p target gene only | Experiment supported target gene (TarBase v.8) |
| ENSG00000172380 | GNG12 | miR-29a/c-3p common target gene | Experiment supported target gene (TarBase v.8) |
| ENSG00000172404 | DNAJB7 | miR-29a/c-3p common target gene | Theoretical prediction (microT-CDS algorithm) |
| ENSG00000172432 | GTPBP2 | miR-29a-3p target gene only | Experiment supported target gene (TarBase v.8) |
| ENSG00000172478 | C2orf54 | miR-29c-3p target gene only | Experiment supported target gene (TarBase v.8) |
| ENSG00000172572 | PDE3A | miR-29a/c-3p common target gene | Experiment supported target gene (TarBase v.8) |
| ENSG00000172663 | TMEM134 | miR-29c-3p target gene only | Experiment supported target gene (TarBase v.8) |
| ENSG00000172667 | ZMAT3 | miR-29c-3p target gene only | Experiment supported target gene (TarBase v.8) |
| ENSG00000172716 | SLFN11 | miR-29a/c-3p common target gene | Experiment supported target gene (TarBase v.8) |
| ENSG00000172733 | PURG | miR-29a/c-3p common target gene | Experiment supported target gene (TarBase v.8) |
| ENSG00000172757 | CFL1 | miR-29a/c-3p common target gene | Theoretical prediction (microT-CDS algorithm) |
| ENSG00000172795 | DCP2 | miR-29a/c-3p common target gene | Theoretical prediction (microT-CDS algorithm) |
| ENSG00000172830 | SSH3 | miR-29c-3p target gene only | Experiment supported target gene (TarBase v.8) |
| ENSG00000172878 | METAP1D | miR-29c-3p target gene only | Experiment supported target gene (TarBase v.8) |
| ENSG00000172915 | NBEA | miR-29a/c-3p common target gene | Experiment supported target gene (TarBase v.8) |
| ENSG00000172927 | MYEOV | miR-29c-3p target gene only | Experiment supported target gene (TarBase v.8) |
| ENSG00000172936 | MYD88 | miR-29c-3p target gene only | Experiment supported target gene (TarBase v.8) |
| ENSG00000172954 | LCLAT1 | miR-29a/c-3p common target gene | Experiment supported target gene (TarBase v.8) |
| ENSG00000172986 | GXYLT2 | miR-29a/c-3p common target gene | Theoretical prediction (microT-CDS algorithm) |
| ENSG00000172995 | ARPP21 | miR-29c-3p target gene only | Experiment supported target gene (TarBase v.8) |
| ENSG00000173011 | TADA2B | miR-29a/c-3p common target gene | Theoretical prediction (microT-CDS algorithm) |
| ENSG00000173013 | CCDC96 | miR-29c-3p target gene only | Experiment supported target gene (TarBase v.8) |
| ENSG00000173039 | RELA | miR-29a/c-3p common target gene | Experiment supported target gene (TarBase v.8) |
| ENSG00000173064 | HECTD4 | miR-29a/c-3p common target gene | Experiment supported target gene (TarBase v.8) |
| ENSG00000173114 | LRRN3 | miR-29a-3p target gene only | Experiment supported target gene (TarBase v.8) |
| ENSG00000173166 | RAPH1 | miR-29a/c-3p common target gene | Experiment supported target gene (TarBase v.8) |
| ENSG00000173218 | VANGL1 | miR-29a/c-3p common target gene | Theoretical prediction (microT-CDS algorithm) |

| | | | |
|---|---|---|---|
| ENSG00000173237 | C11orf86 | miR-29c-3p target gene only | Experiment supported target gene (TarBase v.8) |
| ENSG00000173262 | SLC2A14 | miR-29c-3p target gene only | Experiment supported target gene (TarBase v.8) |
| ENSG00000173276 | ZBTB21 | miR-29a/c-3p common target gene | Experiment supported target gene (TarBase v.8) |
| ENSG00000173391 | OLR1 | miR-29a/c-3p common target gene | Experiment supported target gene (TarBase v.8) |
| ENSG00000173406 | DAB1 | miR-29a/c-3p common target gene | Theoretical prediction (microT-CDS algorithm) |
| ENSG00000173431 | RNASE8 | miR-29c-3p target gene only | Experiment supported target gene (TarBase v.8) |
| ENSG00000173451 | THAP2 | miR-29a/c-3p common target gene | Experiment supported target gene (TarBase v.8) |
| ENSG00000173456 | RNF26 | miR-29a-3p target gene only | Experiment supported target gene (TarBase v.8) |
| ENSG00000173473 | SMARCC1 | miR-29a/c-3p common target gene | Experiment supported target gene (TarBase v.8) |
| ENSG00000173482 | PTPRM | miR-29a/c-3p common target gene | Experiment supported target gene (TarBase v.8) |
| ENSG00000173531 | MST1 | miR-29a/c-3p common target gene | Theoretical prediction (microT-CDS algorithm) |
| ENSG00000173535 | TNFRSF10C | miR-29c-3p target gene only | Experiment supported target gene (TarBase v.8) |
| ENSG00000173546 | CSPG4 | miR-29a/c-3p common target gene | Theoretical prediction (microT-CDS algorithm) |
| ENSG00000173557 | C2orf70 | miR-29c-3p target gene only | Experiment supported target gene (TarBase v.8) |
| ENSG00000173621 | LRFN4 | miR-29c-3p target gene only | Experiment supported target gene (TarBase v.8) |
| ENSG00000173626 | TRAPPC3L | miR-29c-3p target gene only | Experiment supported target gene (TarBase v.8) |
| ENSG00000173638 | SLC19A1 | miR-29a/c-3p common target gene | Experiment supported target gene (TarBase v.8) |
| ENSG00000173726 | TOMM20 | miR-29a/c-3p common target gene | Experiment supported target gene (TarBase v.8) |
| ENSG00000173801 | JUP | miR-29a-3p target gene only | Experiment supported target gene (TarBase v.8) |
| ENSG00000173818 | ENDOV | miR-29a-3p target gene only | Experiment supported target gene (TarBase v.8) |
| ENSG00000173848 | NET1 | miR-29a-3p target gene only | Experiment supported target gene (TarBase v.8) |
| ENSG00000173894 | CBX2 | miR-29a/c-3p common target gene | Experiment supported target gene (TarBase v.8) |
| ENSG00000173950 | XXYLT1 | miR-29a/c-3p common target gene | Experiment supported target gene (TarBase v.8) |
| ENSG00000173960 | UBXN2A | miR-29a/c-3p common target gene | Experiment supported target gene (TarBase v.8) |
| ENSG00000174010 | KLHL15 | miR-29a/c-3p common target gene | Experiment supported target gene (TarBase v.8) |
| ENSG00000174013 | FBXO45 | miR-29a-3p target gene only | Experiment supported target gene (TarBase v.8) |
| ENSG00000174136 | RGMB | miR-29a-3p target gene only | Experiment supported target gene (TarBase v.8) |
| ENSG00000174151 | CYB561D1 | miR-29c-3p target gene only | Theoretical prediction (microT-CDS algorithm) |
| ENSG00000174197 | MGA | miR-29a/c-3p common target gene | Theoretical prediction (microT-CDS algorithm) |
| ENSG00000174282 | ZBTB4 | miR-29c-3p target gene only | Experiment supported target gene (TarBase v.8) |
| ENSG00000174306 | ZHX3 | miR-29a/c-3p common target gene | Experiment supported target gene (TarBase v.8) |
| ENSG00000174307 | PHLDA3 | miR-29c-3p target gene only | Experiment supported target gene (TarBase v.8) |
| ENSG00000174373 | RALGAPA1 | miR-29a/c-3p common target gene | Theoretical prediction (microT-CDS algorithm) |
| ENSG00000174405 | LIG4 | miR-29c-3p target gene only | Experiment supported target gene (TarBase v.8) |
| ENSG00000174437 | ATP2A2 | miR-29a/c-3p common target gene | Theoretical prediction (microT-CDS algorithm) |
| ENSG00000174442 | ZWILCH | miR-29c-3p target gene only | Experiment supported target gene (TarBase v.8) |
| ENSG00000174456 | C12orf76 | miR-29a/c-3p common target gene | Experiment supported target gene (TarBase v.8) |
| ENSG00000174482 | LINGO2 | miR-29a/c-3p common target gene | Theoretical prediction (microT-CDS algorithm) |
| ENSG00000174485 | DENND4A | miR-29a/c-3p common target gene | Experiment supported target gene (TarBase v.8) |
| ENSG00000174502 | SLC26A9 | miR-29c-3p target gene only | Experiment supported target gene (TarBase v.8) |
| ENSG00000174516 | PELI3 | miR-29c-3p target gene only | Experiment supported target gene (TarBase v.8) |
| ENSG00000174567 | GOLT1A | miR-29c-3p target gene only | Experiment supported target gene (TarBase v.8) |
| ENSG00000174611 | KY | miR-29a/c-3p common target gene | Theoretical prediction (microT-CDS algorithm) |
| ENSG00000174705 | SH3PXD2B | miR-29a/c-3p common target gene | Experiment supported target gene (TarBase v.8) |
| ENSG00000174749 | C4orf32 | miR-29a/c-3p common target gene | Theoretical prediction (microT-CDS algorithm) |
| ENSG00000174792 | C4orf26 | miR-29c-3p target gene only | Experiment supported target gene (TarBase v.8) |
| ENSG00000174804 | FZD4 | miR-29a/c-3p common target gene | Theoretical prediction (microT-CDS algorithm) |
| ENSG00000174807 | CD248 | miR-29c-3p target gene only | Experiment supported target gene (TarBase v.8) |
| ENSG00000174839 | DENND6A | miR-29c-3p target gene only | Experiment supported target gene (TarBase v.8) |
| ENSG00000174939 | ASPHD1 | miR-29a/c-3p common target gene | Experiment supported target gene (TarBase v.8) |
| ENSG00000174951 | FUT1 | miR-29c-3p target gene only | Experiment supported target gene (TarBase v.8) |
| ENSG00000175040 | CHST2 | miR-29a-3p target gene only | Experiment supported target gene (TarBase v.8) |
| ENSG00000175073 | VCPIP1 | miR-29a/c-3p common target gene | Experiment supported target gene (TarBase v.8) |
| ENSG00000175087 | PDIK1L | miR-29a/c-3p common target gene | Experiment supported target gene (TarBase v.8) |
| ENSG00000175130 | MARCKSL1 | miR-29c-3p target gene only | Experiment supported target gene (TarBase v.8) |
| ENSG00000175137 | SH3BP5L | miR-29a/c-3p common target gene | Experiment supported target gene (TarBase v.8) |
| ENSG00000175155 | YPEL2 | miR-29a/c-3p common target gene | Experiment supported target gene (TarBase v.8) |
| ENSG00000175161 | CADM2 | miR-29a/c-3p common target gene | Experiment supported target gene (TarBase v.8) |
| ENSG00000175170 | FAM182B | miR-29a/c-3p common target gene | Theoretical prediction (microT-CDS algorithm) |
| ENSG00000175198 | PCCA | miR-29c-3p target gene only | Experiment supported target gene (TarBase v.8) |
| ENSG00000175215 | CTDSP2 | miR-29c-3p target gene only | Experiment supported target gene (TarBase v.8) |
| ENSG00000175229 | GAL3ST3 | miR-29a/c-3p common target gene | Experiment supported target gene (TarBase v.8) |
| ENSG00000175344 | CHRNA7 | miR-29a/c-3p common target gene | Theoretical prediction (microT-CDS algorithm) |
| ENSG00000175387 | SMAD2 | miR-29a/c-3p common target gene | Experiment supported target gene (TarBase v.8) |
| ENSG00000175416 | CLTB | miR-29a-3p target gene only | Experiment supported target gene (TarBase v.8) |
| ENSG00000175445 | LPL | miR-29a/c-3p common target gene | Experiment supported target gene (TarBase v.8) |
| ENSG00000175505 | CLCF1 | miR-29c-3p target gene only | Experiment supported target gene (TarBase v.8) |
| ENSG00000175606 | TMEM70 | miR-29a/c-3p common target gene | Experiment supported target gene (TarBase v.8) |
| ENSG00000175646 | PRM1 | miR-29c-3p target gene only | Experiment supported target gene (TarBase v.8) |
| ENSG00000175727 | MLXIP | miR-29a/c-3p common target gene | Experiment supported target gene (TarBase v.8) |
| ENSG00000175866 | BAIAP2 | miR-29a/c-3p common target gene | Theoretical prediction (microT-CDS algorithm) |
| ENSG00000175868 | CALCB | miR-29a/c-3p common target gene | Experiment supported target gene (TarBase v.8) |
| ENSG00000175893 | ZDHHC21 | miR-29a/c-3p common target gene | Theoretical prediction (microT-CDS algorithm) |
| ENSG00000175920 | DOK7 | miR-29c-3p target gene only | Experiment supported target gene (TarBase v.8) |
| ENSG00000175928 | LRRN1 | miR-29a-3p target gene only | Experiment supported target gene (TarBase v.8) |
| ENSG00000176022 | B3GALT6 | miR-29c-3p target gene only | Experiment supported target gene (TarBase v.8) |
| ENSG00000176046 | NUPR1 | miR-29a-3p target gene only | Experiment supported target gene (TarBase v.8) |
| ENSG00000176092 | AIM1L | miR-29c-3p target gene only | Experiment supported target gene (TarBase v.8) |
| ENSG00000176095 | IP6K1 | miR-29a-3p target gene only | Experiment supported target gene (TarBase v.8) |
| ENSG00000176171 | BNIP3 | miR-29c-3p target gene only | Experiment supported target gene (TarBase v.8) |
| ENSG00000176209 | SMIM19 | miR-29c-3p target gene only | Experiment supported target gene (TarBase v.8) |
| ENSG00000176273 | SLC35G1 | miR-29c-3p target gene only | Experiment supported target gene (TarBase v.8) |
| ENSG00000176407 | KCMF1 | miR-29a/c-3p common target gene | Experiment supported target gene (TarBase v.8) |
| ENSG00000176444 | CLK2 | miR-29a/c-3p common target gene | Theoretical prediction (microT-CDS algorithm) |
| ENSG00000176490 | DIRAS1 | miR-29c-3p target gene only | Experiment supported target gene (TarBase v.8) |
| ENSG00000176597 | B3GNT5 | miR-29c-3p target gene only | Experiment supported target gene (TarBase v.8) |
| ENSG00000176619 | LMNB2 | miR-29c-3p target gene only | Experiment supported target gene (TarBase v.8) |
| ENSG00000176641 | RNF152 | miR-29a/c-3p common target gene | Experiment supported target gene (TarBase v.8) |
| ENSG00000176771 | NCKAP5 | miR-29a/c-3p common target gene | Experiment supported target gene (TarBase v.8) |
| ENSG00000176842 | IRX5 | miR-29c-3p target gene only | Experiment supported target gene (TarBase v.8) |
| ENSG00000176887 | SOX11 | miR-29a/c-3p common target gene | Experiment supported target gene (TarBase v.8) |
| ENSG00000176890 | TYMS | miR-29a/c-3p common target gene | Experiment supported target gene (TarBase v.8) |
| ENSG00000176928 | GCNT4 | miR-29a-3p target gene only | Experiment supported target gene (TarBase v.8) |
| ENSG00000176986 | SEC24C | miR-29c-3p target gene only | Experiment supported target gene (TarBase v.8) |
| ENSG00000177025 | C19orf18 | miR-29c-3p target gene only | Experiment supported target gene (TarBase v.8) |
| ENSG00000177125 | ZBTB34 | miR-29a/c-3p common target gene | Experiment supported target gene (TarBase v.8) |
| ENSG00000177169 | ULK1 | miR-29a/c-3p common target gene | Experiment supported target gene (TarBase v.8) |
| ENSG00000177200 | CHD9 | miR-29c-3p target gene only | Experiment supported target gene (TarBase v.8) |
| ENSG00000177335 | C8orf31 | miR-29a/c-3p common target gene | Theoretical prediction (microT-CDS algorithm) |
| ENSG00000177459 | ERICH5 | miR-29c-3p target gene only | Experiment supported target gene (TarBase v.8) |
| ENSG00000177469 | PTRF | miR-29a/c-3p common target gene | Experiment supported target gene (TarBase v.8) |
| ENSG00000177494 | ZBED2 | miR-29c-3p target gene only | Experiment supported target gene (TarBase v.8) |
| ENSG00000177542 | SLC25A22 | miR-29a/c-3p common target gene | Experiment supported target gene (TarBase v.8) |
| ENSG00000177565 | TBL1XR1 | miR-29a/c-3p common target gene | Theoretical prediction (microT-CDS algorithm) |
| ENSG00000177595 | PIDD1 | miR-29c-3p target gene only | Experiment supported target gene (TarBase v.8) |
| ENSG00000177606 | JUN | miR-29c-3p target gene only | Experiment supported target gene (TarBase v.8) |
| ENSG00000177721 | ANXA2R | miR-29c-3p target gene only | Experiment supported target gene (TarBase v.8) |
| ENSG00000177732 | SOX12 | miR-29a/c-3p common target gene | Experiment supported target gene (TarBase v.8) |
| ENSG00000177853 | ZNF518A | miR-29a/c-3p common target gene | Experiment supported target gene (TarBase v.8) |
| ENSG00000177888 | ZBTB41 | miR-29a/c-3p common target gene | Experiment supported target gene (TarBase v.8) |
| ENSG00000177971 | IMP3 | miR-29a/c-3p common target gene | Experiment supported target gene (TarBase v.8) |
| ENSG00000178053 | MLF1 | miR-29a/c-3p common target gene | Theoretical prediction (microT-CDS algorithm) |
| ENSG00000178105 | DDX10 | miR-29c-3p target gene only | Experiment supported target gene (TarBase v.8) |
| ENSG00000178163 | ZNF518B | miR-29a/c-3p common target gene | Experiment supported target gene (TarBase v.8) |
| ENSG00000178175 | ZNF366 | miR-29a/c-3p common target gene | Theoretical prediction (microT-CDS algorithm) |
| ENSG00000178177 | LCORL | miR-29a/c-3p common target gene | Experiment supported target gene (TarBase v.8) |
| ENSG00000178202 | KDELC2 | miR-29a-3p target gene only | Experiment supported target gene (TarBase v.8) |
| ENSG00000178252 | WDR6 | miR-29a-3p target gene only | Experiment supported target gene (TarBase v.8) |
| ENSG00000178385 | PLEKHM3 | miR-29a/c-3p common target gene | Experiment supported target gene (TarBase v.8) |
| ENSG00000178462 | TUBAL3 | miR-29c-3p target gene only | Experiment supported target gene (TarBase v.8) |
| ENSG00000178467 | P4HTM | miR-29c-3p target gene only | Experiment supported target gene (TarBase v.8) |
| ENSG00000178531 | CTXN1 | miR-29c-3p target gene only | Experiment supported target gene (TarBase v.8) |
| ENSG00000178585 | CTNNBIP1 | miR-29a/c-3p common target gene | Experiment supported target gene (TarBase v.8) |
| ENSG00000178719 | GRINA | miR-29a/c-3p common target gene | Experiment supported target gene (TarBase v.8) |
| ENSG00000178971 | CTC1 | miR-29a/c-3p common target gene | Theoretical prediction (microT-CDS algorithm) |
| ENSG00000179010 | MRFAP1 | miR-29a/c-3p common target gene | Experiment supported target gene (TarBase v.8) |
| ENSG00000179051 | RCC2 | miR-29a/c-3p common target gene | Experiment supported target gene (TarBase v.8) |
| ENSG00000179094 | PER1 | miR-29a/c-3p common target gene | Experiment supported target gene (TarBase v.8) |
| ENSG00000179119 | SPTY2D1 | miR-29a/c-3p common target gene | Experiment supported target gene (TarBase v.8) |
| ENSG00000179133 | C10orf67 | miR-29a/c-3p common target gene | Theoretical prediction (microT-CDS algorithm) |
| ENSG00000179361 | ARID3B | miR-29c-3p target gene only | Experiment supported target gene (TarBase v.8) |
| ENSG00000179454 | KLHL28 | miR-29a/c-3p common target gene | Experiment supported target gene (TarBase v.8) |
| ENSG00000179456 | ZBTB18 | miR-29c-3p target gene only | Experiment supported target gene (TarBase v.8) |
| ENSG00000179750 | APOBEC3B | miR-29a-3p target gene only | Experiment supported target gene (TarBase v.8) |
| ENSG00000179820 | MYADM | miR-29a-3p target gene only | Experiment supported target gene (TarBase v.8) |
| ENSG00000179841 | AKAP5 | miR-29a/c-3p common target gene | Experiment supported target gene (TarBase v.8) |
| ENSG00000179886 | TIGD5 | miR-29a/c-3p common target gene | Theoretical prediction (microT-CDS algorithm) |
| ENSG00000179913 | B3GNT3 | miR-29c-3p target gene only | Experiment supported target gene (TarBase v.8) |
| ENSG00000179965 | ZNF771 | miR-29a/c-3p common target gene | Experiment supported target gene (TarBase v.8) |
| ENSG00000180182 | MED14 | miR-29a/c-3p common target gene | Experiment supported target gene (TarBase v.8) |
| ENSG00000180228 | PRKRA | miR-29a/c-3p common target gene | Experiment supported target gene (TarBase v.8) |
| ENSG00000180304 | OAZ2 | miR-29a-3p target gene only | Experiment supported target gene (TarBase v.8) |
| ENSG00000180329 | CCDC43 | miR-29c-3p target gene only | Experiment supported target gene (TarBase v.8) |
| ENSG00000180354 | MTURN | miR-29a/c-3p common target gene | Experiment supported target gene (TarBase v.8) |
| ENSG00000180530 | NRIP1 | miR-29a/c-3p common target gene | Experiment supported target gene (TarBase v.8) |
| ENSG00000180596 | HIST1H2BC | miR-29a-3p target gene only | Experiment supported target gene (TarBase v.8) |
| ENSG00000180667 | YOD1 | miR-29a/c-3p common target gene | Experiment supported target gene (TarBase v.8) |
| ENSG00000180801 | ARSJ | miR-29a-3p target gene only | Experiment supported target gene (TarBase v.8) |
| ENSG00000180891 | CUEDC1 | miR-29a/c-3p common target gene | Experiment supported target gene (TarBase v.8) |
| ENSG00000180914 | OXTR | miR-29a/c-3p common target gene | Theoretical prediction (microT-CDS algorithm) |
| ENSG00000180921 | FAM83H | miR-29a/c-3p common target gene | Experiment supported target gene (TarBase v.8) |
| ENSG00000180957 | PITPNB | miR-29a/c-3p common target gene | Experiment supported target gene (TarBase v.8) |
| ENSG00000181026 | AEN | miR-29a/c-3p common target gene | Theoretical prediction (microT-CDS algorithm) |
| ENSG00000181045 | SLC26A11 | miR-29a/c-3p common target gene | Experiment supported target gene (TarBase v.8) |
| ENSG00000181090 | EHMT1 | miR-29a/c-3p common target gene | Theoretical prediction (microT-CDS algorithm) |
| ENSG00000181104 | F2R | miR-29a-3p target gene only | Experiment supported target gene (TarBase v.8) |
| ENSG00000181274 | FRAT2 | miR-29a/c-3p common target gene | Experiment supported target gene (TarBase v.8) |
| ENSG00000181467 | RAP2B | miR-29a-3p target gene only | Experiment supported target gene (TarBase v.8) |
| ENSG00000181555 | SETD2 | miR-29a-3p target gene only | Experiment supported target gene (TarBase v.8) |
| ENSG00000181690 | PLAG1 | miR-29a/c-3p common target gene | Experiment supported target gene (TarBase v.8) |
| ENSG00000181744 | C3orf58 | miR-29a/c-3p common target gene | Experiment supported target gene (TarBase v.8) |
| ENSG00000181788 | SIAH2 | miR-29c-3p target gene only | Experiment supported target gene (TarBase v.8) |
| ENSG00000181827 | RFX7 | miR-29a/c-3p common target gene | Theoretical prediction (microT-CDS algorithm) |
| ENSG00000181904 | C5orf24 | miR-29a/c-3p common target gene | Experiment supported target gene (TarBase v.8) |
| ENSG00000181915 | ADO | miR-29a/c-3p common target gene | Experiment supported target gene (TarBase v.8) |
| ENSG00000182010 | RTKN2 | miR-29a/c-3p common target gene | Theoretical prediction (microT-CDS algorithm) |
| ENSG00000182035 | ADIG | miR-29c-3p target gene only | Experiment supported target gene (TarBase v.8) |
| ENSG00000182095 | TNRC18 | miR-29a/c-3p common target gene | Experiment supported target gene (TarBase v.8) |
| ENSG00000182118 | FAM89A | miR-29c-3p target gene only | Experiment supported target gene (TarBase v.8) |
| ENSG00000182158 | CREB3L2 | miR-29a-3p target gene only | Experiment supported target gene (TarBase v.8) |
| ENSG00000182197 | EXT1 | miR-29a/c-3p common target gene | Theoretical prediction (microT-CDS algorithm) |
| ENSG00000182218 | HHIPL1 | miR-29a/c-3p common target gene | Theoretical prediction (microT-CDS algorithm) |
| ENSG00000182240 | BACE2 | miR-29a-3p target gene only | Experiment supported target gene (TarBase v.8) |
| ENSG00000182263 | FIGN | miR-29a/c-3p common target gene | Theoretical prediction (microT-CDS algorithm) |
| ENSG00000182504 | CEP97 | miR-29a/c-3p common target gene | Experiment supported target gene (TarBase v.8) |
| ENSG00000182557 | SPNS3 | miR-29c-3p target gene only | Experiment supported target gene (TarBase v.8) |
| ENSG00000182580 | EPHB3 | miR-29c-3p target gene only | Experiment supported target gene (TarBase v.8) |
| ENSG00000182636 | NDN | miR-29a/c-3p common target gene | Experiment supported target gene (TarBase v.8) |
| ENSG00000182700 | IGIP | miR-29c-3p target gene only | Experiment supported target gene (TarBase v.8) |
| ENSG00000182831 | C16orf72 | miR-29a/c-3p common target gene | Experiment supported target gene (TarBase v.8) |
| ENSG00000182890 | GLUD2 | miR-29a/c-3p common target gene | Experiment supported target gene (TarBase v.8) |
| ENSG00000182983 | ZNF662 | miR-29a/c-3p common target gene | Theoretical prediction (microT-CDS algorithm) |
| ENSG00000182985 | CADM1 | miR-29a-3p target gene only | Experiment supported target gene (TarBase v.8) |
| ENSG00000183018 | SPNS2 | miR-29c-3p target gene only | Experiment supported target gene (TarBase v.8) |
| ENSG00000183049 | CAMK1D | miR-29a/c-3p common target gene | Experiment supported target gene (TarBase v.8) |
| ENSG00000183054 | RGPD6 | miR-29a/c-3p common target gene | Theoretical prediction (microT-CDS algorithm) |
| ENSG00000183091 | NEB | miR-29c-3p target gene only | Experiment supported target gene (TarBase v.8) |
| ENSG00000183255 | PTTG1IP | miR-29a/c-3p common target gene | Experiment supported target gene (TarBase v.8) |
| ENSG00000183283 | DAZAP2 | miR-29c-3p target gene only | Experiment supported target gene (TarBase v.8) |
| ENSG00000183287 | CCBE1 | miR-29a/c-3p common target gene | Experiment supported target gene (TarBase v.8) |
| ENSG00000183337 | BCOR | miR-29a/c-3p common target gene | Experiment supported target gene (TarBase v.8) |
| ENSG00000183474 | GTF2H2C | miR-29c-3p target gene only | Experiment supported target gene (TarBase v.8) |
| ENSG00000183486 | MX2 | miR-29c-3p target gene only | Experiment supported target gene (TarBase v.8) |
| ENSG00000183496 | MEX3B | miR-29a/c-3p common target gene | Theoretical prediction (microT-CDS algorithm) |
| ENSG00000183530 | PRR14L | miR-29a/c-3p common target gene | Experiment supported target gene (TarBase v.8) |
| ENSG00000183655 | KLHL25 | miR-29c-3p target gene only | Experiment supported target gene (TarBase v.8) |
| ENSG00000183665 | TRMT12 | miR-29c-3p target gene only | Experiment supported target gene (TarBase v.8) |
| ENSG00000183688 | FAM101B | miR-29a/c-3p common target gene | Theoretical prediction (microT-CDS algorithm) |
| ENSG00000183690 | EFHC2 | miR-29c-3p target gene only | Experiment supported target gene (TarBase v.8) |
| ENSG00000183696 | UPP1 | miR-29c-3p target gene only | Experiment supported target gene (TarBase v.8) |
| ENSG00000183709 | IFNL2 | miR-29c-3p target gene only | Experiment supported target gene (TarBase v.8) |
| ENSG00000183741 | CBX6 | miR-29a/c-3p common target gene | Experiment supported target gene (TarBase v.8) |
| ENSG00000183742 | MACC1 | miR-29a/c-3p common target gene | Experiment supported target gene (TarBase v.8) |
| ENSG00000183783 | KCTD8 | miR-29a/c-3p common target gene | Experiment supported target gene (TarBase v.8) |
| ENSG00000183808 | RBM12B | miR-29c-3p target gene only | Experiment supported target gene (TarBase v.8) |
| ENSG00000183833 | MAATS1 | miR-29a/c-3p common target gene | Experiment supported target gene (TarBase v.8) |
| ENSG00000183853 | KIRREL | miR-29a/c-3p common target gene | Theoretical prediction (microT-CDS algorithm) |
| ENSG00000184007 | PTP4A2 | miR-29a-3p target gene only | Experiment supported target gene (TarBase v.8) |
| ENSG00000184012 | TMPRSS2 | miR-29c-3p target gene only | Experiment supported target gene (TarBase v.8) |
| ENSG00000184056 | VPS33B | miR-29a/c-3p common target gene | Theoretical prediction (microT-CDS algorithm) |
| ENSG00000184060 | ADAP2 | miR-29c-3p target gene only | Experiment supported target gene (TarBase v.8) |
| ENSG00000184113 | CLDN5 | miR-29a-3p target gene only | Experiment supported target gene (TarBase v.8) |
| ENSG00000184178 | SCFD2 | miR-29c-3p target gene only | Experiment supported target gene (TarBase v.8) |
| ENSG00000184226 | PCDH9 | miR-29a/c-3p common target gene | Experiment supported target gene (TarBase v.8) |
| ENSG00000184305 | CCSER1 | miR-29a/c-3p common target gene | Experiment supported target gene (TarBase v.8) |
| ENSG00000184349 | EFNA5 | miR-29a/c-3p common target gene | Experiment supported target gene (TarBase v.8) |
| ENSG00000184402 | SS18L1 | miR-29a/c-3p common target gene | Experiment supported target gene (TarBase v.8) |
| ENSG00000184428 | TOP1MT | miR-29c-3p target gene only | Experiment supported target gene (TarBase v.8) |
| ENSG00000184500 | PROS1 | miR-29a-3p target gene only | Experiment supported target gene (TarBase v.8) |
| ENSG00000184508 | HDDC3 | miR-29c-3p target gene only | Experiment supported target gene (TarBase v.8) |
| ENSG00000184575 | XPOT | miR-29a/c-3p common target gene | Experiment supported target gene (TarBase v.8) |
| ENSG00000184602 | SNN | miR-29a/c-3p common target gene | Experiment supported target gene (TarBase v.8) |
| ENSG00000184611 | KCNH7 | miR-29a-3p target gene only | Experiment supported target gene (TarBase v.8) |
| ENSG00000184675 | AMER1 | miR-29a/c-3p common target gene | Experiment supported target gene (TarBase v.8) |
| ENSG00000184677 | ZBTB40 | miR-29a/c-3p common target gene | Theoretical prediction (microT-CDS algorithm) |
| ENSG00000184678 | HIST2H2BE | miR-29a-3p target gene only | Experiment supported target gene (TarBase v.8) |
| ENSG00000184702 | 5-Sep | miR-29a/c-3p common target gene | Theoretical prediction (microT-CDS algorithm) |
| ENSG00000184840 | TMED9 | miR-29a/c-3p common target gene | Experiment supported target gene (TarBase v.8) |
| ENSG00000184845 | DRD1 | miR-29a/c-3p common target gene | Theoretical prediction (microT-CDS algorithm) |
| ENSG00000184863 | RBM33 | miR-29a/c-3p common target gene | Experiment supported target gene (TarBase v.8) |
| ENSG00000184905 | TCEAL2 | miR-29c-3p target gene only | Experiment supported target gene (TarBase v.8) |
| ENSG00000184916 | JAG2 | miR-29c-3p target gene only | Experiment supported target gene (TarBase v.8) |
| ENSG00000184979 | USP18 | miR-29c-3p target gene only | Experiment supported target gene (TarBase v.8) |
| ENSG00000184986 | TMEM121 | miR-29c-3p target gene only | Experiment supported target gene (TarBase v.8) |
| ENSG00000184992 | BRI3BP | miR-29a/c-3p common target gene | Experiment supported target gene (TarBase v.8) |
| ENSG00000185002 | RFX6 | miR-29a/c-3p common target gene | Experiment supported target gene (TarBase v.8) |
| ENSG00000185009 | AP3M1 | miR-29a-3p target gene only | Experiment supported target gene (TarBase v.8) |
| ENSG00000185090 | MANEAL | miR-29a/c-3p common target gene | Experiment supported target gene (TarBase v.8) |
| ENSG00000185129 | PURA | miR-29a/c-3p common target gene | Experiment supported target gene (TarBase v.8) |
| ENSG00000185156 | MFSD6L | miR-29c-3p target gene only | Experiment supported target gene (TarBase v.8) |
| ENSG00000185201 | IFITM2 | miR-29c-3p target gene only | Experiment supported target gene (TarBase v.8) |
| ENSG00000185278 | ZBTB37 | miR-29a/c-3p common target gene | Experiment supported target gene (TarBase v.8) |
| ENSG00000185298 | CCDC137 | miR-29c-3p target gene only | Experiment supported target gene (TarBase v.8) |
| ENSG00000185479 | KRT6B | miR-29c-3p target gene only | Experiment supported target gene (TarBase v.8) |
| ENSG00000185483 | ROR1 | miR-29a/c-3p common target gene | Theoretical prediction (microT-CDS algorithm) |
| ENSG00000185507 | IRF7 | miR-29c-3p target gene only | Experiment supported target gene (TarBase v.8) |
| ENSG00000185591 | SP1 | miR-29a/c-3p common target gene | Experiment supported target gene (TarBase v.8) |
| ENSG00000185619 | PCGF3 | miR-29a/c-3p common target gene | Experiment supported target gene (TarBase v.8) |

| | | | |
|---|---|---|---|
| ENSG00000185621 | LMLN | miR-29a-3p target gene only | Theoretical prediction (microT-CDS algorithm) |
| ENSG00000185650 | ZFP36L1 | miR-29a/c-3p common target gene | Experiment supported target gene (TarBase v.8) |
| ENSG00000185658 | BRWD1 | miR-29a/c-3p common target gene | Theoretical prediction (microT-CDS algorithm) |
| ENSG00000185670 | ZBTB3 | miR-29c-3p target gene only | Experiment supported target gene (TarBase v.8) |
| ENSG00000185730 | ZNF696 | miR-29a/c-3p common target gene | Experiment supported target gene (TarBase v.8) |
| ENSG00000185742 | C11orf87 | miR-29a/c-3p common target gene | Experiment supported target gene (TarBase v.8) |
| ENSG00000185745 | IFIT1 | miR-29a/c-3p common target gene | Experiment supported target gene (TarBase v.8) |
| ENSG00000185787 | MORF4L1 | miR-29a/c-3p common target gene | Experiment supported target gene (TarBase v.8) |
| ENSG00000185818 | NAT8L | miR-29c-3p target gene only | Experiment supported target gene (TarBase v.8) |
| ENSG00000185829 | ARL17A | miR-29a/c-3p common target gene | Theoretical prediction (microT-CDS algorithm) |
| ENSG00000185838 | GNB1L | miR-29c-3p target gene only | Experiment supported target gene (TarBase v.8) |
| ENSG00000185885 | IFITM1 | miR-29c-3p target gene only | Experiment supported target gene (TarBase v.8) |
| ENSG00000185896 | LAMP1 | miR-29a/c-3p common target gene | Experiment supported target gene (TarBase v.8) |
| ENSG00000185900 | POMK | miR-29a/c-3p common target gene | Experiment supported target gene (TarBase v.8) |
| ENSG00000185947 | ZNF267 | miR-29c-3p target gene only | Experiment supported target gene (TarBase v.8) |
| ENSG00000186026 | ZNF284 | miR-29a-3p target gene only | Experiment supported target gene (TarBase v.8) |
| ENSG00000186073 | C15orf41 | miR-29a-3p target gene only | Experiment supported target gene (TarBase v.8) |
| ENSG00000186141 | POLR3C | miR-29a-3p target gene only | Experiment supported target gene (TarBase v.8) |
| ENSG00000186174 | BCL9L | miR-29a/c-3p common target gene | Experiment supported target gene (TarBase v.8) |
| ENSG00000186188 | FFAR4 | miR-29c-3p target gene only | Experiment supported target gene (TarBase v.8) |
| ENSG00000186204 | CYP4F12 | miR-29c-3p target gene only | Experiment supported target gene (TarBase v.8) |
| ENSG00000186310 | NAP1L3 | miR-29a/c-3p common target gene | Experiment supported target gene (TarBase v.8) |
| ENSG00000186318 | BACE1 | miR-29a/c-3p common target gene | Experiment supported target gene (TarBase v.8) |
| ENSG00000186329 | TMEM212 | miR-29a/c-3p common target gene | Theoretical prediction (microT-CDS algorithm) |
| ENSG00000186416 | NKRF | miR-29a/c-3p common target gene | Theoretical prediction (microT-CDS algorithm) |
| ENSG00000186432 | KPNA4 | miR-29a/c-3p common target gene | Experiment supported target gene (TarBase v.8) |
| ENSG00000186448 | ZNF197 | miR-29c-3p target gene only | Theoretical prediction (microT-CDS algorithm) |
| ENSG00000186469 | GNG2 | miR-29a/c-3p common target gene | Experiment supported target gene (TarBase v.8) |
| ENSG00000186480 | INSIG1 | miR-29a/c-3p common target gene | Experiment supported target gene (TarBase v.8) |
| ENSG00000186496 | ZNF396 | miR-29a/c-3p common target gene | Theoretical prediction (microT-CDS algorithm) |
| ENSG00000186526 | CYP4F8 | miR-29c-3p target gene only | Experiment supported target gene (TarBase v.8) |
| ENSG00000186529 | CYP4F3 | miR-29a/c-3p common target gene | Theoretical prediction (microT-CDS algorithm) |
| ENSG00000186660 | ZFP91 | miR-29a/c-3p common target gene | Experiment supported target gene (TarBase v.8) |
| ENSG00000186687 | LYRM7 | miR-29a/c-3p common target gene | Experiment supported target gene (TarBase v.8) |
| ENSG00000186767 | SPIN4 | miR-29a/c-3p common target gene | Theoretical prediction (microT-CDS algorithm) |
| ENSG00000186814 | ZSCAN30 | miR-29a/c-3p common target gene | Theoretical prediction (microT-CDS algorithm) |
| ENSG00000187231 | SESTD1 | miR-29a/c-3p common target gene | Experiment supported target gene (TarBase v.8) |
| ENSG00000187239 | FNBP1 | miR-29a/c-3p common target gene | Experiment supported target gene (TarBase v.8) |
| ENSG00000187243 | MAGED4B | miR-29c-3p target gene only | Experiment supported target gene (TarBase v.8) |
| ENSG00000187288 | CIDEC | miR-29a/c-3p common target gene | Experiment supported target gene (TarBase v.8) |
| ENSG00000187498 | COL4A1 | miR-29a/c-3p common target gene | Experiment supported target gene (TarBase v.8) |
| ENSG00000187510 | PLEKHG7 | miR-29a/c-3p common target gene | Theoretical prediction (microT-CDS algorithm) |
| ENSG00000187605 | TET3 | miR-29a/c-3p common target gene | Experiment supported target gene (TarBase v.8) |
| ENSG00000187626 | ZKSCAN4 | miR-29a/c-3p common target gene | Experiment supported target gene (TarBase v.8) |
| ENSG00000187678 | SPRY4 | miR-29a/c-3p common target gene | Experiment supported target gene (TarBase v.8) |
| ENSG00000187699 | C2orf88 | miR-29a/c-3p common target gene | Theoretical prediction (microT-CDS algorithm) |
| ENSG00000187736 | NHEJ1 | miR-29c-3p target gene only | Experiment supported target gene (TarBase v.8) |
| ENSG00000188001 | TPRG1 | miR-29c-3p target gene only | Experiment supported target gene (TarBase v.8) |
| ENSG00000188010 | MORN2 | miR-29c-3p target gene only | Experiment supported target gene (TarBase v.8) |
| ENSG00000188042 | ARL4C | miR-29c-3p target gene only | Experiment supported target gene (TarBase v.8) |
| ENSG00000188070 | C11orf95 | miR-29c-3p target gene only | Experiment supported target gene (TarBase v.8) |
| ENSG00000188153 | COL4A5 | miR-29a/c-3p common target gene | Experiment supported target gene (TarBase v.8) |
| ENSG00000188176 | SMTNL2 | miR-29a/c-3p common target gene | Theoretical prediction (microT-CDS algorithm) |
| ENSG00000188313 | PLSCR1 | miR-29c-3p target gene only | Experiment supported target gene (TarBase v.8) |
| ENSG00000188419 | CHM | miR-29a/c-3p common target gene | Experiment supported target gene (TarBase v.8) |
| ENSG00000188517 | COL25A1 | miR-29a/c-3p common target gene | Theoretical prediction (microT-CDS algorithm) |
| ENSG00000188529 | SRSF10 | miR-29a/c-3p common target gene | Experiment supported target gene (TarBase v.8) |
| ENSG00000188549 | C15orf52 | miR-29c-3p target gene only | Experiment supported target gene (TarBase v.8) |
| ENSG00000188613 | NANOS1 | miR-29a/c-3p common target gene | Experiment supported target gene (TarBase v.8) |
| ENSG00000188636 | LDOC1L | miR-29a/c-3p common target gene | Experiment supported target gene (TarBase v.8) |
| ENSG00000188643 | S100A16 | miR-29c-3p target gene only | Experiment supported target gene (TarBase v.8) |
| ENSG00000188647 | PTAR1 | miR-29a/c-3p common target gene | Theoretical prediction (microT-CDS algorithm) |

| | | | |
|---|---|---|---|
| ENSG00000188672 | RHCE | miR-29a/c-3p common target gene | Theoretical prediction (microT-CDS algorithm) |
| ENSG00000188738 | FSIP2 | miR-29a/c-3p common target gene | Theoretical prediction (microT-CDS algorithm) |
| ENSG00000188807 | TMEM201 | miR-29a-3p target gene only | Experiment supported target gene (TarBase v.8) |
| ENSG00000188895 | MSL1 | miR-29a/c-3p common target gene | Theoretical prediction (microT-CDS algorithm) |
| ENSG00000188910 | GJB3 | miR-29c-3p target gene only | Experiment supported target gene (TarBase v.8) |
| ENSG00000188994 | ZNF292 | miR-29a/c-3p common target gene | Experiment supported target gene (TarBase v.8) |
| ENSG00000189060 | H1F0 | miR-29c-3p target gene only | Experiment supported target gene (TarBase v.8) |
| ENSG00000189067 | LITAF | miR-29c-3p target gene only | Experiment supported target gene (TarBase v.8) |
| ENSG00000189079 | ARID2 | miR-29a/c-3p common target gene | Experiment supported target gene (TarBase v.8) |
| ENSG00000189143 | CLDN4 | miR-29c-3p target gene only | Experiment supported target gene (TarBase v.8) |
| ENSG00000189159 | HN1 | miR-29a/c-3p common target gene | Experiment supported target gene (TarBase v.8) |
| ENSG00000189184 | PCDH18 | miR-29c-3p target gene only | Experiment supported target gene (TarBase v.8) |
| ENSG00000189221 | MAOA | miR-29a-3p target gene only | Experiment supported target gene (TarBase v.8) |
| ENSG00000189241 | TSPYL1 | miR-29c-3p target gene only | Experiment supported target gene (TarBase v.8) |
| ENSG00000189280 | GJB5 | miR-29c-3p target gene only | Experiment supported target gene (TarBase v.8) |
| ENSG00000189298 | ZKSCAN3 | miR-29a/c-3p common target gene | Theoretical prediction (microT-CDS algorithm) |
| ENSG00000189377 | CXCL17 | miR-29c-3p target gene only | Experiment supported target gene (TarBase v.8) |
| ENSG00000196072 | BLOC1S2 | miR-29a/c-3p common target gene | Experiment supported target gene (TarBase v.8) |
| ENSG00000196083 | IL1RAP | miR-29a/c-3p common target gene | Theoretical prediction (microT-CDS algorithm) |
| ENSG00000196116 | TDRD7 | miR-29c-3p target gene only | Experiment supported target gene (TarBase v.8) |
| ENSG00000196139 | AKR1C3 | miR-29a/c-3p common target gene | Experiment supported target gene (TarBase v.8) |
| ENSG00000196182 | STK40 | miR-29c-3p target gene only | Experiment supported target gene (TarBase v.8) |
| ENSG00000196230 | TUBB | miR-29a-3p target gene only | Experiment supported target gene (TarBase v.8) |
| ENSG00000196345 | ZKSCAN7 | miR-29a/c-3p common target gene | Theoretical prediction (microT-CDS algorithm) |
| ENSG00000196368 | NUDT11 | miR-29a/c-3p common target gene | Theoretical prediction (microT-CDS algorithm) |
| ENSG00000196396 | PTPN1 | miR-29a/c-3p common target gene | Theoretical prediction (microT-CDS algorithm) |
| ENSG00000196405 | EVL | miR-29a/c-3p common target gene | Experiment supported target gene (TarBase v.8) |
| ENSG00000196408 | NOXO1 | miR-29c-3p target gene only | Experiment supported target gene (TarBase v.8) |
| ENSG00000196422 | PPP1R26 | miR-29c-3p target gene only | Experiment supported target gene (TarBase v.8) |
| ENSG00000196428 | TSC22D2 | miR-29a/c-3p common target gene | Experiment supported target gene (TarBase v.8) |
| ENSG00000196449 | YRDC | miR-29a/c-3p common target gene | Theoretical prediction (microT-CDS algorithm) |
| ENSG00000196470 | SIAH1 | miR-29a/c-3p common target gene | Experiment supported target gene (TarBase v.8) |
| ENSG00000196504 | PRPF40A | miR-29a/c-3p common target gene | Experiment supported target gene (TarBase v.8) |
| ENSG00000196511 | TPK1 | miR-29a/c-3p common target gene | Experiment supported target gene (TarBase v.8) |
| ENSG00000196526 | AFAP1 | miR-29a/c-3p common target gene | Experiment supported target gene (TarBase v.8) |
| ENSG00000196549 | MME | miR-29a-3p target gene only | Experiment supported target gene (TarBase v.8) |
| ENSG00000196562 | SULF2 | miR-29a-3p target gene only | Experiment supported target gene (TarBase v.8) |
| ENSG00000196569 | LAMA2 | miR-29a/c-3p common target gene | Experiment supported target gene (TarBase v.8) |
| ENSG00000196628 | TCF4 | miR-29a/c-3p common target gene | Experiment supported target gene (TarBase v.8) |
| ENSG00000196642 | RABL6 | miR-29a/c-3p common target gene | Experiment supported target gene (TarBase v.8) |
| ENSG00000196653 | ZNF502 | miR-29c-3p target gene only | Experiment supported target gene (TarBase v.8) |
| ENSG00000196659 | TTC30B | miR-29a/c-3p common target gene | Experiment supported target gene (TarBase v.8) |
| ENSG00000196684 | HSH2D | miR-29c-3p target gene only | Experiment supported target gene (TarBase v.8) |
| ENSG00000196700 | ZNF512B | miR-29a/c-3p common target gene | Experiment supported target gene (TarBase v.8) |
| ENSG00000196704 | AMZ2 | miR-29c-3p target gene only | Experiment supported target gene (TarBase v.8) |
| ENSG00000196705 | ZNF431 | miR-29a/c-3p common target gene | Experiment supported target gene (TarBase v.8) |
| ENSG00000196712 | NF1 | miR-29a/c-3p common target gene | Theoretical prediction (microT-CDS algorithm) |
| ENSG00000196739 | COL27A1 | miR-29a/c-3p common target gene | Theoretical prediction (microT-CDS algorithm) |
| ENSG00000196743 | GM2A | miR-29a/c-3p common target gene | Theoretical prediction (microT-CDS algorithm) |
| ENSG00000196754 | S100A2 | miR-29c-3p target gene only | Experiment supported target gene (TarBase v.8) |
| ENSG00000196757 | ZNF700 | miR-29c-3p target gene only | Experiment supported target gene (TarBase v.8) |
| ENSG00000196782 | MAML3 | miR-29a-3p target gene only | Experiment supported target gene (TarBase v.8) |
| ENSG00000196793 | ZNF239 | miR-29c-3p target gene only | Experiment supported target gene (TarBase v.8) |
| ENSG00000196865 | NHLRC2 | miR-29a/c-3p common target gene | Experiment supported target gene (TarBase v.8) |
| ENSG00000196932 | TMEM26 | miR-29a-3p target gene only | Theoretical prediction (microT-CDS algorithm) |
| ENSG00000196937 | FAM3C | miR-29a/c-3p common target gene | Theoretical prediction (microT-CDS algorithm) |
| ENSG00000196950 | SLC39A10 | miR-29a/c-3p common target gene | Experiment supported target gene (TarBase v.8) |
| ENSG00000196968 | FUT11 | miR-29a/c-3p common target gene | Experiment supported target gene (TarBase v.8) |
| ENSG00000197008 | ZNF138 | miR-29c-3p target gene only | Experiment supported target gene (TarBase v.8) |
| ENSG00000197037 | ZSCAN25 | miR-29a/c-3p common target gene | Experiment supported target gene (TarBase v.8) |
| ENSG00000197045 | GMFB | miR-29a/c-3p common target gene | Experiment supported target gene (TarBase v.8) |
| ENSG00000197061 | HIST1H4C | miR-29a/c-3p common target gene | Experiment supported target gene (TarBase v.8) |

| | | | |
|---|---|---|---|
| ENSG00000197063 | MAFG | miR-29a/c-3p common target gene | Experiment supported target gene (TarBase v.8) |
| ENSG00000197102 | DYNC1H1 | miR-29a-3p target gene only | Experiment supported target gene (TarBase v.8) |
| ENSG00000197110 | IFNL3 | miR-29c-3p target gene only | Experiment supported target gene (TarBase v.8) |
| ENSG00000197119 | SLC25A29 | miR-29a-3p target gene only | Experiment supported target gene (TarBase v.8) |
| ENSG00000197124 | ZNF682 | miR-29a/c-3p common target gene | Experiment supported target gene (TarBase v.8) |
| ENSG00000197134 | ZNF257 | miR-29c-3p target gene only | Experiment supported target gene (TarBase v.8) |
| ENSG00000197153 | HIST1H3J | miR-29c-3p target gene only | Experiment supported target gene (TarBase v.8) |
| ENSG00000197157 | SND1 | miR-29a/c-3p common target gene | Experiment supported target gene (TarBase v.8) |
| ENSG00000197183 | NOL4L | miR-29a/c-3p common target gene | Experiment supported target gene (TarBase v.8) |
| ENSG00000197208 | SLC22A4 | miR-29a-3p target gene only | Experiment supported target gene (TarBase v.8) |
| ENSG00000197296 | FITM2 | miR-29a/c-3p common target gene | Experiment supported target gene (TarBase v.8) |
| ENSG00000197312 | DDI2 | miR-29a/c-3p common target gene | Experiment supported target gene (TarBase v.8) |
| ENSG00000197321 | SVIL | miR-29a/c-3p common target gene | Theoretical prediction (microT-CDS algorithm) |
| ENSG00000197355 | UAP1L1 | miR-29a-3p target gene only | Experiment supported target gene (TarBase v.8) |
| ENSG00000197386 | HTT | miR-29a/c-3p common target gene | Experiment supported target gene (TarBase v.8) |
| ENSG00000197415 | VEPH1 | miR-29c-3p target gene only | Experiment supported target gene (TarBase v.8) |
| ENSG00000197444 | OGDHL | miR-29c-3p target gene only | Experiment supported target gene (TarBase v.8) |
| ENSG00000197461 | PDGFA | miR-29a/c-3p common target gene | Experiment supported target gene (TarBase v.8) |
| ENSG00000197479 | PCDHB11 | miR-29a/c-3p common target gene | Experiment supported target gene (TarBase v.8) |
| ENSG00000197565 | COL4A6 | miR-29a/c-3p common target gene | Theoretical prediction (microT-CDS algorithm) |
| ENSG00000197579 | TOPORS | miR-29a/c-3p common target gene | Experiment supported target gene (TarBase v.8) |
| ENSG00000197622 | CDC42SE1 | miR-29a/c-3p common target gene | Experiment supported target gene (TarBase v.8) |
| ENSG00000197694 | SPTAN1 | miR-29a/c-3p common target gene | Experiment supported target gene (TarBase v.8) |
| ENSG00000197696 | NMB | miR-29c-3p target gene only | Experiment supported target gene (TarBase v.8) |
| ENSG00000197860 | SGTB | miR-29a/c-3p common target gene | Experiment supported target gene (TarBase v.8) |
| ENSG00000197872 | FAM49A | miR-29c-3p target gene only | Experiment supported target gene (TarBase v.8) |
| ENSG00000197894 | ADH5 | miR-29a-3p target gene only | Experiment supported target gene (TarBase v.8) |
| ENSG00000197948 | FCHSD1 | miR-29a/c-3p common target gene | Theoretical prediction (microT-CDS algorithm) |
| ENSG00000197959 | DNM3 | miR-29a/c-3p common target gene | Experiment supported target gene (TarBase v.8) |
| ENSG00000197969 | VPS13A | miR-29a/c-3p common target gene | Experiment supported target gene (TarBase v.8) |
| ENSG00000198010 | DLGAP2 | miR-29a/c-3p common target gene | Experiment supported target gene (TarBase v.8) |
| ENSG00000198018 | ENTPD7 | miR-29a/c-3p common target gene | Experiment supported target gene (TarBase v.8) |
| ENSG00000198034 | RPS4X | miR-29a/c-3p common target gene | Experiment supported target gene (TarBase v.8) |
| ENSG00000198042 | MAK16 | miR-29c-3p target gene only | Experiment supported target gene (TarBase v.8) |
| ENSG00000198055 | GRK6 | miR-29c-3p target gene only | Experiment supported target gene (TarBase v.8) |
| ENSG00000198074 | AKR1B10 | miR-29c-3p target gene only | Experiment supported target gene (TarBase v.8) |
| ENSG00000198113 | TOR4A | miR-29a-3p target gene only | Experiment supported target gene (TarBase v.8) |
| ENSG00000198121 | LPAR1 | miR-29a/c-3p common target gene | Experiment supported target gene (TarBase v.8) |
| ENSG00000198142 | SOWAHC | miR-29a/c-3p common target gene | Experiment supported target gene (TarBase v.8) |
| ENSG00000198146 | ZNF770 | miR-29a/c-3p common target gene | Experiment supported target gene (TarBase v.8) |
| ENSG00000198185 | ZNF334 | miR-29c-3p target gene only | Experiment supported target gene (TarBase v.8) |
| ENSG00000198252 | STYX | miR-29a/c-3p common target gene | Experiment supported target gene (TarBase v.8) |
| ENSG00000198265 | HELZ | miR-29a/c-3p common target gene | Experiment supported target gene (TarBase v.8) |
| ENSG00000198363 | ASPH | miR-29c-3p target gene only | Experiment supported target gene (TarBase v.8) |
| ENSG00000198417 | MT1F | miR-29c-3p target gene only | Experiment supported target gene (TarBase v.8) |
| ENSG00000198431 | TXNRD1 | miR-29a/c-3p common target gene | Experiment supported target gene (TarBase v.8) |
| ENSG00000198464 | ZNF480 | miR-29c-3p target gene only | Experiment supported target gene (TarBase v.8) |
| ENSG00000198513 | ATL1 | miR-29a-3p target gene only | Experiment supported target gene (TarBase v.8) |
| ENSG00000198538 | ZNF28 | miR-29a/c-3p common target gene | Theoretical prediction (microT-CDS algorithm) |
| ENSG00000198554 | WDHD1 | miR-29a/c-3p common target gene | Experiment supported target gene (TarBase v.8) |
| ENSG00000198561 | CTNND1 | miR-29a/c-3p common target gene | Experiment supported target gene (TarBase v.8) |
| ENSG00000198576 | ARC | miR-29c-3p target gene only | Experiment supported target gene (TarBase v.8) |
| ENSG00000198626 | RYR2 | miR-29a/c-3p common target gene | Experiment supported target gene (TarBase v.8) |
| ENSG00000198690 | FAN1 | miR-29a/c-3p common target gene | Experiment supported target gene (TarBase v.8) |
| ENSG00000198700 | IPO9 | miR-29a-3p target gene only | Experiment supported target gene (TarBase v.8) |
| ENSG00000198720 | ANKRD13B | miR-29a/c-3p common target gene | Experiment supported target gene (TarBase v.8) |
| ENSG00000198727 | MT-CYB | miR-29a-3p target gene only | Experiment supported target gene (TarBase v.8) |
| ENSG00000198729 | PPP1R14C | miR-29c-3p target gene only | Experiment supported target gene (TarBase v.8) |
| ENSG00000198740 | ZNF652 | miR-29a/c-3p common target gene | Experiment supported target gene (TarBase v.8) |
| ENSG00000198742 | SMURF1 | miR-29c-3p target gene only | Experiment supported target gene (TarBase v.8) |
| ENSG00000198743 | SLC5A3 | miR-29a-3p target gene only | Experiment supported target gene (TarBase v.8) |
| ENSG00000198746 | GPATCH3 | miR-29c-3p target gene only | Experiment supported target gene (TarBase v.8) |

| | | | |
|---|---|---|---|
| ENSG00000198794 | SCAMP5 | miR-29a/c-3p common target gene | Experiment supported target gene (TarBase v.8) |
| ENSG00000198804 | MT-CO1 | miR-29a-3p target gene only | Experiment supported target gene (TarBase v.8) |
| ENSG00000198824 | CHAMP1 | miR-29a/c-3p common target gene | Experiment supported target gene (TarBase v.8) |
| ENSG00000198825 | INPP5F | miR-29c-3p target gene only | Experiment supported target gene (TarBase v.8) |
| ENSG00000198856 | OSTC | miR-29a/c-3p common target gene | Experiment supported target gene (TarBase v.8) |
| ENSG00000198858 | R3HDM4 | miR-29a/c-3p common target gene | Experiment supported target gene (TarBase v.8) |
| ENSG00000198862 | LTN1 | miR-29c-3p target gene only | Experiment supported target gene (TarBase v.8) |
| ENSG00000198863 | RUNDC1 | miR-29a/c-3p common target gene | Experiment supported target gene (TarBase v.8) |
| ENSG00000198876 | DCAF12 | miR-29a/c-3p common target gene | Experiment supported target gene (TarBase v.8) |
| ENSG00000198881 | ASB12 | miR-29a/c-3p common target gene | Theoretical prediction (microT-CDS algorithm) |
| ENSG00000198889 | DCAF12L1 | miR-29a/c-3p common target gene | Theoretical prediction (microT-CDS algorithm) |
| ENSG00000198890 | PRMT6 | miR-29a/c-3p common target gene | Experiment supported target gene (TarBase v.8) |
| ENSG00000198911 | SREBF2 | miR-29a-3p target gene only | Experiment supported target gene (TarBase v.8) |
| ENSG00000198925 | ATG9A | miR-29a/c-3p common target gene | Experiment supported target gene (TarBase v.8) |
| ENSG00000198932 | GPRASP1 | miR-29a/c-3p common target gene | Experiment supported target gene (TarBase v.8) |
| ENSG00000198937 | CCDC167 | miR-29c-3p target gene only | Experiment supported target gene (TarBase v.8) |
| ENSG00000198959 | TGM2 | miR-29a-3p target gene only | Experiment supported target gene (TarBase v.8) |
| ENSG00000203668 | CHML | miR-29c-3p target gene only | Experiment supported target gene (TarBase v.8) |
| ENSG00000203727 | SAMD5 | miR-29c-3p target gene only | Experiment supported target gene (TarBase v.8) |
| ENSG00000203780 | FANK1 | miR-29c-3p target gene only | Experiment supported target gene (TarBase v.8) |
| ENSG00000203814 | HIST2H2BF | miR-29a/c-3p common target gene | Experiment supported target gene (TarBase v.8) |
| ENSG00000203943 | SAMD13 | miR-29a/c-3p common target gene | Theoretical prediction (microT-CDS algorithm) |
| ENSG00000204001 | LCN8 | miR-29c-3p target gene only | Experiment supported target gene (TarBase v.8) |
| ENSG00000204019 | CT83 | miR-29c-3p target gene only | Experiment supported target gene (TarBase v.8) |
| ENSG00000204084 | INPP5B | miR-29a/c-3p common target gene | Theoretical prediction (microT-CDS algorithm) |
| ENSG00000204086 | RPA4 | miR-29c-3p target gene only | Experiment supported target gene (TarBase v.8) |
| ENSG00000204103 | MAFB | miR-29a/c-3p common target gene | Experiment supported target gene (TarBase v.8) |
| ENSG00000204118 | NAP1L6 | miR-29a/c-3p common target gene | Theoretical prediction (microT-CDS algorithm) |
| ENSG00000204120 | GIGYF2 | miR-29a/c-3p common target gene | Experiment supported target gene (TarBase v.8) |
| ENSG00000204262 | COL5A2 | miR-29a/c-3p common target gene | Experiment supported target gene (TarBase v.8) |
| ENSG00000204291 | COL15A1 | miR-29a/c-3p common target gene | Experiment supported target gene (TarBase v.8) |
| ENSG00000204308 | RNF5 | miR-29c-3p target gene only | Experiment supported target gene (TarBase v.8) |
| ENSG00000204314 | PRRT1 | miR-29c-3p target gene only | Experiment supported target gene (TarBase v.8) |
| ENSG00000204344 | STK19 | miR-29c-3p target gene only | Experiment supported target gene (TarBase v.8) |
| ENSG00000204406 | MBD5 | miR-29a/c-3p common target gene | Theoretical prediction (microT-CDS algorithm) |
| ENSG00000204435 | CSNK2B | miR-29c-3p target gene only | Experiment supported target gene (TarBase v.8) |
| ENSG00000204463 | BAG6 | miR-29a/c-3p common target gene | Experiment supported target gene (TarBase v.8) |
| ENSG00000204498 | NFKBIL1 | miR-29a/c-3p common target gene | Experiment supported target gene (TarBase v.8) |
| ENSG00000204516 | MICB | miR-29a/c-3p common target gene | Experiment supported target gene (TarBase v.8) |
| ENSG00000204520 | MICA | miR-29c-3p target gene only | Experiment supported target gene (TarBase v.8) |
| ENSG00000204560 | DHX16 | miR-29a/c-3p common target gene | Experiment supported target gene (TarBase v.8) |
| ENSG00000204569 | PPP1R10 | miR-29c-3p target gene only | Experiment supported target gene (TarBase v.8) |
| ENSG00000204576 | PRR3 | miR-29a/c-3p common target gene | Theoretical prediction (microT-CDS algorithm) |
| ENSG00000204618 | RNF39 | miR-29a/c-3p common target gene | Experiment supported target gene (TarBase v.8) |
| ENSG00000204655 | MOG | miR-29a/c-3p common target gene | Theoretical prediction (microT-CDS algorithm) |
| ENSG00000204815 | TTC25 | miR-29c-3p target gene only | Experiment supported target gene (TarBase v.8) |
| ENSG00000204843 | DCTN1 | miR-29c-3p target gene only | Experiment supported target gene (TarBase v.8) |
| ENSG00000204856 | FAM216A | miR-29c-3p target gene only | Experiment supported target gene (TarBase v.8) |
| ENSG00000204920 | ZNF155 | miR-29a/c-3p common target gene | Experiment supported target gene (TarBase v.8) |
| ENSG00000204946 | ZNF783 | miR-29a/c-3p common target gene | Theoretical prediction (microT-CDS algorithm) |
| ENSG00000204961 | PCDHA9 | miR-29a/c-3p common target gene | Experiment supported target gene (TarBase v.8) |
| ENSG00000204962 | PCDHA8 | miR-29a/c-3p common target gene | Theoretical prediction (microT-CDS algorithm) |
| ENSG00000204963 | PCDHA7 | miR-29a/c-3p common target gene | Theoretical prediction (microT-CDS algorithm) |
| ENSG00000204965 | PCDHA5 | miR-29a/c-3p common target gene | Theoretical prediction (microT-CDS algorithm) |
| ENSG00000204967 | PCDHA4 | miR-29a/c-3p common target gene | Theoretical prediction (microT-CDS algorithm) |
| ENSG00000204969 | PCDHA2 | miR-29a/c-3p common target gene | Theoretical prediction (microT-CDS algorithm) |
| ENSG00000204970 | PCDHA1 | miR-29a/c-3p common target gene | Theoretical prediction (microT-CDS algorithm) |
| ENSG00000205060 | SLC35B4 | miR-29c-3p target gene only | Experiment supported target gene (TarBase v.8) |
| ENSG00000205089 | CCNI2 | miR-29c-3p target gene only | Experiment supported target gene (TarBase v.8) |
| ENSG00000205189 | ZBTB10 | miR-29a/c-3p common target gene | Experiment supported target gene (TarBase v.8) |
| ENSG00000205212 | CCDC144NL | miR-29a/c-3p common target gene | Theoretical prediction (microT-CDS algorithm) |
| ENSG00000205581 | HMGN1 | miR-29a-3p target gene only | Theoretical prediction (microT-CDS algorithm) |

| | | | |
|---|---|---|---|
| ENSG00000205643 | CDPF1 | miR-29a/c-3p common target gene | Experiment supported target gene (TarBase v.8) |
| ENSG00000205730 | ITPRIPL2 | miR-29a/c-3p common target gene | Experiment supported target gene (TarBase v.8) |
| ENSG00000205791 | LOH12CR2 | miR-29a-3p target gene only | Theoretical prediction (microT-CDS algorithm) |
| ENSG00000206150 | RNASE13 | miR-29a/c-3p common target gene | Theoretical prediction (microT-CDS algorithm) |
| ENSG00000206190 | ATP10A | miR-29a/c-3p common target gene | Theoretical prediction (microT-CDS algorithm) |
| ENSG00000206418 | RAB12 | miR-29a/c-3p common target gene | Experiment supported target gene (TarBase v.8) |
| ENSG00000206557 | TRIM71 | miR-29a-3p target gene only | Experiment supported target gene (TarBase v.8) |
| ENSG00000206560 | ANKRD28 | miR-29a/c-3p common target gene | Experiment supported target gene (TarBase v.8) |
| ENSG00000206579 | XKR4 | miR-29a/c-3p common target gene | Theoretical prediction (microT-CDS algorithm) |
| ENSG00000211448 | DIO2 | miR-29a/c-3p common target gene | Experiment supported target gene (TarBase v.8) |
| ENSG00000211455 | STK38L | miR-29a/c-3p common target gene | Experiment supported target gene (TarBase v.8) |
| ENSG00000212127 | TAS2R14 | miR-29c-3p target gene only | Experiment supported target gene (TarBase v.8) |
| ENSG00000213024 | NUP62 | miR-29a/c-3p common target gene | Experiment supported target gene (TarBase v.8) |
| ENSG00000213047 | DENND1B | miR-29a/c-3p common target gene | Theoretical prediction (microT-CDS algorithm) |
| ENSG00000213064 | SFT2D2 | miR-29c-3p target gene only | Experiment supported target gene (TarBase v.8) |
| ENSG00000213190 | MLLT11 | miR-29c-3p target gene only | Experiment supported target gene (TarBase v.8) |
| ENSG00000213281 | NRAS | miR-29a/c-3p common target gene | Experiment supported target gene (TarBase v.8) |
| ENSG00000213380 | COG8 | miR-29a/c-3p common target gene | Experiment supported target gene (TarBase v.8) |
| ENSG00000213585 | VDAC1 | miR-29a/c-3p common target gene | Experiment supported target gene (TarBase v.8) |
| ENSG00000213694 | S1PR3 | miR-29c-3p target gene only | Experiment supported target gene (TarBase v.8) |
| ENSG00000213699 | SLC35F6 | miR-29a-3p target gene only | Experiment supported target gene (TarBase v.8) |
| ENSG00000213782 | DDX47 | miR-29a-3p target gene only | Experiment supported target gene (TarBase v.8) |
| ENSG00000213923 | CSNK1E | miR-29a/c-3p common target gene | Experiment supported target gene (TarBase v.8) |
| ENSG00000213928 | IRF9 | miR-29c-3p target gene only | Experiment supported target gene (TarBase v.8) |
| ENSG00000214063 | TSPAN4 | miR-29c-3p target gene only | Experiment supported target gene (TarBase v.8) |
| ENSG00000214097 | SMCO1 | miR-29a/c-3p common target gene | Theoretical prediction (microT-CDS algorithm) |
| ENSG00000214107 | MAGEB1 | miR-29a-3p target gene only | Experiment supported target gene (TarBase v.8) |
| ENSG00000214216 | IQCJ | miR-29a/c-3p common target gene | Theoretical prediction (microT-CDS algorithm) |
| ENSG00000214357 | NEURL1B | miR-29c-3p target gene only | Experiment supported target gene (TarBase v.8) |
| ENSG00000214575 | CPEB1 | miR-29c-3p target gene only | Experiment supported target gene (TarBase v.8) |
| ENSG00000214595 | EML6 | miR-29a/c-3p common target gene | Theoretical prediction (microT-CDS algorithm) |
| ENSG00000214756 | METTL12 | miR-29c-3p target gene only | Experiment supported target gene (TarBase v.8) |
| ENSG00000214814 | FER1L6 | miR-29a-3p target gene only | Experiment supported target gene (TarBase v.8) |
| ENSG00000214860 | EVPLL | miR-29c-3p target gene only | Experiment supported target gene (TarBase v.8) |
| ENSG00000215012 | C22orf29 | miR-29a/c-3p common target gene | Experiment supported target gene (TarBase v.8) |
| ENSG00000215193 | PEX26 | miR-29a/c-3p common target gene | Theoretical prediction (microT-CDS algorithm) |
| ENSG00000215301 | DDX3X | miR-29a/c-3p common target gene | Experiment supported target gene (TarBase v.8) |
| ENSG00000215305 | VPS16 | miR-29a/c-3p common target gene | Experiment supported target gene (TarBase v.8) |
| ENSG00000215717 | TMEM167B | miR-29a-3p target gene only | Experiment supported target gene (TarBase v.8) |
| ENSG00000215788 | TNFRSF25 | miR-29c-3p target gene only | Experiment supported target gene (TarBase v.8) |
| ENSG00000216490 | IFI30 | miR-29a/c-3p common target gene | Experiment supported target gene (TarBase v.8) |
| ENSG00000217930 | PAM16 | miR-29c-3p target gene only | Experiment supported target gene (TarBase v.8) |
| ENSG00000218336 | TENM3 | miR-29a/c-3p common target gene | Theoretical prediction (microT-CDS algorithm) |
| ENSG00000219073 | CELA3B | miR-29c-3p target gene only | Experiment supported target gene (TarBase v.8) |
| ENSG00000221886 | ZBED8 | miR-29c-3p target gene only | Experiment supported target gene (TarBase v.8) |
| ENSG00000221926 | TRIM16 | miR-29a/c-3p common target gene | Theoretical prediction (microT-CDS algorithm) |
| ENSG00000221978 | CCNL2 | miR-29a/c-3p common target gene | Theoretical prediction (microT-CDS algorithm) |
| ENSG00000222047 | C10orf55 | miR-29a/c-3p common target gene | Theoretical prediction (microT-CDS algorithm) |
| ENSG00000223572 | CKMT1A | miR-29c-3p target gene only | Experiment supported target gene (TarBase v.8) |
| ENSG00000225614 | ZNF469 | miR-29a/c-3p common target gene | Experiment supported target gene (TarBase v.8) |
| ENSG00000225830 | ERCC6 | miR-29a/c-3p common target gene | Experiment supported target gene (TarBase v.8) |
| ENSG00000227345 | PARG | miR-29a/c-3p common target gene | Experiment supported target gene (TarBase v.8) |
| ENSG00000229415 | SFTA3 | miR-29a/c-3p common target gene | Theoretical prediction (microT-CDS algorithm) |
| ENSG00000229809 | ZNF688 | miR-29c-3p target gene only | Experiment supported target gene (TarBase v.8) |
| ENSG00000231274 | SBK3 | miR-29a/c-3p common target gene | Experiment supported target gene (TarBase v.8) |
| ENSG00000234602 | MCIDAS | miR-29a/c-3p common target gene | Theoretical prediction (microT-CDS algorithm) |
| ENSG00000236279 | CLEC2L | miR-29a/c-3p common target gene | Theoretical prediction (microT-CDS algorithm) |
| ENSG00000239389 | PCDHA13 | miR-29a/c-3p common target gene | Theoretical prediction (microT-CDS algorithm) |
| ENSG00000239590 | OR1J4 | miR-29a-3p target gene only | Experiment supported target gene (TarBase v.8) |
| ENSG00000239697 | TNFSF12 | miR-29c-3p target gene only | Experiment supported target gene (TarBase v.8) |
| ENSG00000239900 | ADSL | miR-29a/c-3p common target gene | Experiment supported target gene (TarBase v.8) |
| ENSG00000240583 | AQP1 | miR-29a/c-3p common target gene | Experiment supported target gene (TarBase v.8) |
| ENSG00000240682 | ISY1 | miR-29a/c-3p common target gene | Experiment supported target gene (TarBase v.8) |
| ENSG00000240764 | PCDHGC5 | miR-29a/c-3p common target gene | Experiment supported target gene (TarBase v.8) |
| ENSG00000240849 | TMEM189 | miR-29a-3p target gene only | Theoretical prediction (microT-CDS algorithm) |
| ENSG00000241233 | KRTAP5-8 | miR-29c-3p target gene only | Experiment supported target gene (TarBase v.8) |
| ENSG00000241468 | ATP5J2 | miR-29a/c-3p common target gene | Experiment supported target gene (TarBase v.8) |
| ENSG00000241563 | CORT | miR-29a/c-3p common target gene | Theoretical prediction (microT-CDS algorithm) |
| ENSG00000241685 | ARPC1A | miR-29c-3p target gene only | Experiment supported target gene (TarBase v.8) |
| ENSG00000241697 | TMEFF1 | miR-29a/c-3p common target gene | Experiment supported target gene (TarBase v.8) |
| ENSG00000241878 | PISD | miR-29c-3p target gene only | Experiment supported target gene (TarBase v.8) |
| ENSG00000241973 | PI4KA | miR-29a/c-3p common target gene | Experiment supported target gene (TarBase v.8) |
| ENSG00000242265 | PEG10 | miR-29a-3p target gene only | Experiment supported target gene (TarBase v.8) |
| ENSG00000242372 | EIF6 | miR-29a/c-3p common target gene | Experiment supported target gene (TarBase v.8) |
| ENSG00000242732 | RGAG4 | miR-29a/c-3p common target gene | Theoretical prediction (microT-CDS algorithm) |
| ENSG00000243232 | PCDHAC2 | miR-29a/c-3p common target gene | Theoretical prediction (microT-CDS algorithm) |
| ENSG00000243251 | PGBD3 | miR-29c-3p target gene only | Experiment supported target gene (TarBase v.8) |
| ENSG00000243317 | C7orf73 | miR-29a/c-3p common target gene | Experiment supported target gene (TarBase v.8) |
| ENSG00000243364 | EFNA4 | miR-29c-3p target gene only | Experiment supported target gene (TarBase v.8) |
| ENSG00000243444 | PALM2 | miR-29a/c-3p common target gene | Theoretical prediction (microT-CDS algorithm) |
| ENSG00000243646 | IL10RB | miR-29a/c-3p common target gene | Experiment supported target gene (TarBase v.8) |
| ENSG00000243709 | LEFTY1 | miR-29c-3p target gene only | Experiment supported target gene (TarBase v.8) |
| ENSG00000243725 | TTC4 | miR-29c-3p target gene only | Experiment supported target gene (TarBase v.8) |
| ENSG00000244038 | DDOST | miR-29a/c-3p common target gene | Experiment supported target gene (TarBase v.8) |
| ENSG00000244187 | TMEM141 | miR-29c-3p target gene only | Experiment supported target gene (TarBase v.8) |
| ENSG00000244462 | RBM12 | miR-29a/c-3p common target gene | Experiment supported target gene (TarBase v.8) |
| ENSG00000244482 | LILRA6 | miR-29a/c-3p common target gene | Theoretical prediction (microT-CDS algorithm) |
| ENSG00000244617 | ASPRV1 | miR-29c-3p target gene only | Experiment supported target gene (TarBase v.8) |
| ENSG00000248383 | PCDHAC1 | miR-29a/c-3p common target gene | Theoretical prediction (microT-CDS algorithm) |
| ENSG00000248712 | CCDC153 | miR-29c-3p target gene only | Experiment supported target gene (TarBase v.8) |
| ENSG00000249158 | PCDHA11 | miR-29a/c-3p common target gene | Theoretical prediction (microT-CDS algorithm) |
| ENSG00000249459 | ZNF286B | miR-29a/c-3p common target gene | Theoretical prediction (microT-CDS algorithm) |
| ENSG00000250120 | PCDHA10 | miR-29a/c-3p common target gene | Theoretical prediction (microT-CDS algorithm) |
| ENSG00000251503 | APITD1-CORT | miR-29a/c-3p common target gene | Theoretical prediction (microT-CDS algorithm) |
| ENSG00000251664 | PCDHA12 | miR-29a/c-3p common target gene | Theoretical prediction (microT-CDS algorithm) |
| ENSG00000253159 | PCDHGA12 | miR-29c-3p target gene only | Experiment supported target gene (TarBase v.8) |
| ENSG00000253293 | HOXA10 | miR-29a/c-3p common target gene | Experiment supported target gene (TarBase v.8) |
| ENSG00000253719 | ATXN7L3B | miR-29a/c-3p common target gene | Experiment supported target gene (TarBase v.8) |
| ENSG00000253729 | PRKDC | miR-29a-3p target gene only | Experiment supported target gene (TarBase v.8) |
| ENSG00000254093 | PINX1 | miR-29a/c-3p common target gene | Theoretical prediction (microT-CDS algorithm) |
| ENSG00000254505 | CHMP4A | miR-29c-3p target gene only | Experiment supported target gene (TarBase v.8) |
| ENSG00000255408 | PCDHA3 | miR-29a/c-3p common target gene | Theoretical prediction (microT-CDS algorithm) |
| ENSG00000255690 | TRIL | miR-29a/c-3p common target gene | Experiment supported target gene (TarBase v.8) |
| ENSG00000256043 | CTSO | miR-29a/c-3p common target gene | Theoretical prediction (microT-CDS algorithm) |
| ENSG00000256061 | DYX1C1 | miR-29c-3p target gene only | Experiment supported target gene (TarBase v.8) |
| ENSG00000256683 | ZNF350 | miR-29c-3p target gene only | Experiment supported target gene (TarBase v.8) |
| ENSG00000256870 | SLC5A8 | miR-29a/c-3p common target gene | Theoretical prediction (microT-CDS algorithm) |
| ENSG00000257594 | GALNT4 | miR-29c-3p target gene only | Theoretical prediction (microT-CDS algorithm) |
| ENSG00000258289 | CHURC1 | miR-29a-3p target gene only | Experiment supported target gene (TarBase v.8) |
| ENSG00000258873 | DUXA | miR-29a-3p target gene only | Experiment supported target gene (TarBase v.8) |
| ENSG00000258890 | CEP95 | miR-29a/c-3p common target gene | Theoretical prediction (microT-CDS algorithm) |
| ENSG00000258941 | RP11-407N17.3 | miR-29c-3p target gene only | Experiment supported target gene (TarBase v.8) |
| ENSG00000259075 | POC1B-GALNT4 | miR-29c-3p target gene only | Theoretical prediction (microT-CDS algorithm) |
| ENSG00000259956 | RBM15B | miR-29a-3p target gene only | Experiment supported target gene (TarBase v.8) |
| ENSG00000260092 | RP11-77K12.7 | miR-29a/c-3p common target gene | Theoretical prediction (microT-CDS algorithm) |
| ENSG00000260272 | RP11-20I23.1 | miR-29a-3p target gene only | Experiment supported target gene (TarBase v.8) |
| ENSG00000260903 | XKR7 | miR-29a/c-3p common target gene | Experiment supported target gene (TarBase v.8) |
| ENSG00000260916 | CCPG1 | miR-29a/c-3p common target gene | Experiment supported target gene (TarBase v.8) |
| ENSG00000261115 | TMEM178B | miR-29a/c-3p common target gene | Experiment supported target gene (TarBase v.8) |
| ENSG00000261236 | BOP1 | miR-29c-3p target gene only | Experiment supported target gene (TarBase v.8) |
| ENSG00000261272 | MUC22 | miR-29a/c-3p common target gene | Theoretical prediction (microT-CDS algorithm) |
| ENSG00000261609 | GAN | miR-29a/c-3p common target gene | Experiment supported target gene (TarBase v.8) |
| ENSG00000265241 | RBM8A | miR-29c-3p target gene only | Experiment supported target gene (TarBase v.8) |
| ENSG00000265763 | ZNF488 | miR-29c-3p target gene only | Experiment supported target gene (TarBase v.8) |
| ENSG00000265972 | TXNIP | miR-29a-3p target gene only | Experiment supported target gene (TarBase v.8) |
| ENSG00000268104 | SLC6A14 | miR-29a/c-3p common target gene | Theoretical prediction (microT-CDS algorithm) |
| ENSG00000268182 | SMIM17 | miR-29a/c-3p common target gene | Theoretical prediction (microT-CDS algorithm) |
| ENSG00000269307 | CTD-2278I10.6 | miR-29a/c-3p common target gene | Theoretical prediction (microT-CDS algorithm) |
| ENSG00000271503 | CCL5 | miR-29c-3p target gene only | Experiment supported target gene (TarBase v.8) |
| ENSG00000271601 | LIX1L | miR-29a-3p target gene only | Experiment supported target gene (TarBase v.8) |
| ENSG00000272325 | NUDT3 | miR-29a/c-3p common target gene | Experiment supported target gene (TarBase v.8) |
| ENSG00000272442 | RP11-444E17.6 | miR-29a/c-3p common target gene | Theoretical prediction (microT-CDS algorithm) |
| ENSG00000272886 | DCP1A | miR-29a-3p target gene only | Experiment supported target gene (TarBase v.8) |
| ENSG00000273045 | C2ORF15 | miR-29c-3p target gene only | Experiment supported target gene (TarBase v.8) |
| ENSG00000273079 | GRIN2B | miR-29a/c-3p common target gene | Theoretical prediction (microT-CDS algorithm) |
| ENSG00000273703 | HIST1H2BM | miR-29a-3p target gene only | Experiment supported target gene (TarBase v.8) |
| ENSG00000273802 | HIST1H2BG | miR-29a/c-3p common target gene | Experiment supported target gene (TarBase v.8) |
| ENSG00000274211 | SOCS7 | miR-29a/c-3p common target gene | Experiment supported target gene (TarBase v.8) |
| ENSG00000274349 | ZNF658 | miR-29a/c-3p common target gene | Theoretical prediction (microT-CDS algorithm) |
| ENSG00000274641 | HIST1H2BO | miR-29a-3p target gene only | Experiment supported target gene (TarBase v.8) |
| ENSG00000275004 | ZNF280B | miR-29a-3p target gene only | Experiment supported target gene (TarBase v.8) |
| ENSG00000275183 | LENG9 | miR-29a/c-3p common target gene | Theoretical prediction (microT-CDS algorithm) |
| ENSG00000275302 | CCL4 | miR-29c-3p target gene only | Experiment supported target gene (TarBase v.8) |
| ENSG00000275713 | HIST1H2BH | miR-29a-3p target gene only | Experiment supported target gene (TarBase v.8) |
| ENSG00000275714 | HIST1H3A | miR-29c-3p target gene only | Experiment supported target gene (TarBase v.8) |
| ENSG00000275793 | RIMBP3 | miR-29c-3p target gene only | Experiment supported target gene (TarBase v.8) |
| ENSG00000276043 | UHRF1 | miR-29c-3p target gene only | Experiment supported target gene (TarBase v.8) |
| ENSG00000276085 | CCL3L3 | miR-29c-3p target gene only | Experiment supported target gene (TarBase v.8) |
| ENSG00000276966 | HIST1H4E | miR-29a/c-3p common target gene | Experiment supported target gene (TarBase v.8) |
| ENSG00000277224 | HIST1H2BF | miR-29a/c-3p common target gene | Experiment supported target gene (TarBase v.8) |
| ENSG00000277443 | MARCKS | miR-29a/c-3p common target gene | Experiment supported target gene (TarBase v.8) |
| ENSG00000277632 | CCL3 | miR-29c-3p target gene only | Experiment supported target gene (TarBase v.8) |
| ENSG00000278540 | ACACA | miR-29a/c-3p common target gene | Experiment supported target gene (TarBase v.8) |
| ENSG00000278637 | HIST1H4A | miR-29a/c-3p common target gene | Experiment supported target gene (TarBase v.8) |
| ENSG00000278828 | HIST1H3H | miR-29c-3p target gene only | Experiment supported target gene (TarBase v.8) |
| ENSG00000280055 | TMEM75 | miR-29a/c-3p common target gene | Theoretical prediction (microT-CDS algorithm) |
| ENSG00000283154 | IQCJ-SCHIP1 | miR-29a/c-3p common target gene | Experiment supported target gene (TarBase v.8) |

**Table S3. Differntially expressed miR-29a/c-3p target genes in female and male P0-HUVECs from PE and NT pregnancies.***

| miR-29a/c-3p target genes | | | Male vs. Female in NT P0-HUVECs | | PE vs. NT in Female P0-HUVECs | | PE vs. NT in Male P0-HUVECs | |
|---|---|---|---|---|---|---|---|---|
| GeneID | gene | Human miR-29a/c-3p target | log2[fold change)] | FDR-P value | log2[fold change] | FDR-P value | log2[fold change] | FDR-P value |
| ENSG00000198576 | ARC | miR-29c-3p target gene only | -5.72 | 0.000 | -5.94 | 0.003 | 5.24 | 0.001 |
| ENSG00000160161 | CILP2 | miR-29a/c-3p common target gene | -5.32 | 0.000 | -4.14 | 0.042 | 4.65 | 0.005 |
| ENSG00000104722 | NEFM | miR-29a/c-3p common target gene | -4.31 | 0.001 | -4.32 | 0.040 | 3.59 | 0.016 |
| ENSG00000273079 | GRIN2B | miR-29a/c-3p common target gene | -4.13 | 0.001 | -4.01 | 0.017 | 3.57 | 0.013 |
| ENSG00000175344 | CHRNA7 | miR-29a/c-3p common target gene | -3.52 | 0.001 | -3.76 | 0.010 | 3.33 | 0.017 |
| ENSG00000099812 | MISP | miR-29c-3p target gene only | -5.32 | 0.006 | | NS | 5.00 | 0.029 |
| ENSG00000158050 | DUSP2 | miR-29a/c-3p common target gene | -4.19 | 0.002 | | NS | 3.29 | 0.013 |
| ENSG00000100292 | HMOX1 | miR-29c-3p target gene only | -3.66 | 0.001 | | NS | 3.13 | 0.004 |
| ENSG00000085741 | WNT11 | miR-29c-3p target gene only | -3.56 | 0.009 | | NS | 4.66 | 0.016 |
| ENSG00000104267 | CA2 | miR-29a/c-3p common target gene | -3.27 | 0.006 | | NS | 3.50 | 0.002 |
| ENSG00000129757 | CDKN1C | miR-29c-3p target gene only | -2.89 | 0.003 | | NS | 3.95 | 0.001 |
| ENSG00000108821 | COL1A1 | miR-29a/c-3p common target gene | -2.78 | 0.003 | | NS | 3.13 | 0.000 |
| ENSG00000182580 | EPHB3 | miR-29c-3p target gene only | -2.62 | 0.002 | | NS | 2.90 | 0.031 |
| ENSG00000035499 | DEPDC1B | miR-29c-3p target gene only | -2.19 | 0.040 | | NS | 3.35 | 0.045 |
| ENSG00000136014 | USP44 | miR-29a/c-3p common target gene | -2.00 | 0.014 | | NS | 2.64 | 0.020 |
| ENSG00000163376 | KBTBD8 | miR-29c-3p target gene only | -1.86 | 0.009 | | NS | 2.13 | 0.047 |
| ENSG00000086061 | DNAJA1 | miR-29a-3p target gene only | -1.19 | 0.039 | | NS | 1.51 | 0.046 |
| ENSG00000203814 | HIST2H2BF | miR-29a/c-3p common target gene | | NS | | NS | 1.47 | 0.049 |
| ENSG00000141582 | CBX4 | miR-29a/c-3p common target gene | | NS | | NS | 1.71 | 0.047 |
| ENSG00000166165 | CKB | miR-29a/c-3p common target gene | | NS | | NS | 2.49 | 0.002 |
| ENSG00000076716 | GPC4 | miR-29a/c-3p common target gene | | NS | | NS | 2.52 | 0.037 |
| ENSG00000069493 | CLEC2D | miR-29a/c-3p common target gene | | NS | | NS | 2.74 | 0.021 |
| ENSG00000049540 | ELN | miR-29a/c-3p common target gene | | NS | | NS | 2.81 | 0.001 |
| ENSG00000104415 | WISP1 | miR-29a/c-3p common target gene | | NS | | NS | 2.96 | 0.011 |
| ENSG00000170775 | GPR37 | miR-29a/c-3p common target gene | | NS | | NS | -3.13 | 0.013 |
| ENSG00000189221 | MAOA | miR-29a-3p target gene only | | NS | | NS | -2.57 | 0.031 |
| ENSG00000170917 | NUDT6 | miR-29c-3p target gene only | | NS | | NS | -1.89 | 0.048 |
| ENSG00000134986 | NREP | miR-29a/c-3p common target gene | | NS | | NS | -1.86 | 0.021 |
| ENSG00000061273 | HDAC7 | miR-29a-3p target gene only | 0.87 | 0.049 | 1.48 | 0.016 | | NS |
| ENSG00000163638 | ADAMTS9 | miR-29a/c-3p common target gene | 1.50 | 0.031 | 2.17 | 0.019 | | NS |
| ENSG00000101265 | RASSF2 | miR-29c-3p target gene only | 1.75 | 0.028 | 2.18 | 0.015 | | NS |
| ENSG00000130508 | PXDN | miR-29a/c-3p common target gene | 2.33 | 0.004 | 2.68 | 0.015 | | NS |
| ENSG00000187498 | COL4A1 | miR-29a/c-3p common target gene | 2.37 | 0.001 | 2.88 | 0.014 | | NS |
| ENSG00000167601 | AXL | miR-29a/c-3p common target gene | 2.67 | 0.005 | 2.48 | 0.011 | | NS |
| ENSG00000134871 | COL4A2 | miR-29a/c-3p common target gene | 2.70 | 0.001 | 3.43 | 0.004 | | NS |
| ENSG00000184113 | CLDN5 | miR-29a-3p target gene only | 2.85 | 0.014 | 3.51 | 0.001 | | NS |
| ENSG00000109079 | TNFAIP1 | miR-29c-3p target gene only | | NS | 1.20 | 0.048 | | NS |
| ENSG00000140548 | ZNF710 | miR-29a-3p target gene only | | NS | 1.31 | 0.045 | | NS |
| ENSG00000171680 | PLEKHG5 | miR-29c-3p target gene only | | NS | 1.39 | 0.024 | | NS |
| ENSG00000069399 | BCL3 | miR-29c-3p target gene only | | NS | 1.40 | 0.025 | | NS |
| ENSG00000148154 | UGCG | miR-29a-3p target gene only | | NS | 1.42 | 0.021 | | NS |
| ENSG00000163430 | FSTL1 | miR-29a/c-3p common target gene | | NS | 1.46 | 0.032 | | NS |
| ENSG00000149115 | TNKS1BP1 | miR-29a/c-3p common target gene | | NS | 1.47 | 0.003 | | NS |
| ENSG00000101255 | TRIB3 | miR-29c-3p target gene only | | NS | 1.50 | 0.028 | | NS |
| ENSG00000110446 | SLC15A3 | miR-29c-3p target gene only | | NS | 1.50 | 0.035 | | NS |
| ENSG00000145901 | TNIP1 | miR-29a-3p target gene only | | NS | 1.52 | 0.035 | | NS |
| ENSG00000186480 | INSIG1 | miR-29a/c-3p common target gene | | NS | 1.52 | 0.039 | | NS |
| ENSG00000125089 | SH3TC1 | miR-29a-3p target gene only | | NS | 1.53 | 0.010 | | NS |
| ENSG00000072310 | SREBF1 | miR-29a/c-3p common target gene | | NS | 1.59 | 0.006 | | NS |
| ENSG00000070669 | ASNS | miR-29a-3p target gene only | | NS | 1.65 | 0.016 | | NS |
| ENSG00000169710 | FASN | miR-29a/c-3p common target gene | | NS | 1.72 | 0.024 | | NS |
| ENSG00000277443 | MARCKS | miR-29a/c-3p common target gene | | NS | 1.73 | 0.017 | | NS |
| ENSG00000172183 | ISG20 | miR-29a-3p target gene only | | NS | 1.73 | 0.002 | | NS |
| ENSG00000135069 | PSAT1 | miR-29a-3p target gene only | | NS | 1.74 | 0.048 | | NS |
| ENSG00000216490 | IFI30 | miR-29a/c-3p common target gene | | NS | 1.91 | 0.030 | | NS |
| ENSG00000103044 | HAS3 | miR-29a/c-3p common target gene | | NS | 1.92 | 0.020 | | NS |
| ENSG00000137571 | SLCO5A1 | miR-29a/c-3p common target gene | | NS | 1.95 | 0.042 | | NS |
| ENSG00000198121 | LPAR1 | miR-29a/c-3p common target gene | | NS | 2.10 | 0.021 | | NS |
| ENSG00000118503 | TNFAIP3 | miR-29a/c-3p common target gene | | NS | 2.15 | 0.003 | | NS |
| ENSG00000128274 | A4GALT | miR-29c-3p target gene only | | NS | 2.21 | 0.001 | | NS |
| ENSG00000162711 | NLRP3 | miR-29c-3p target gene only | | NS | 2.25 | 0.014 | | NS |
| ENSG00000141682 | PMAIP1 | miR-29c-3p target gene only | | NS | 2.35 | 0.000 | | NS |
| ENSG00000168398 | BDKRB2 | miR-29a/c-3p common target gene | | NS | 2.38 | 0.032 | | NS |
| ENSG00000167772 | ANGPTL4 | miR-29c-3p target gene only | | NS | 2.52 | 0.007 | | NS |
| ENSG00000169248 | CXCL11 | miR-29c-3p target gene only | | NS | 3.08 | 0.036 | | NS |
| ENSG00000058085 | LAMC2 | miR-29a/c-3p common target gene | | NS | 3.08 | 0.002 | | NS |
| ENSG00000277632 | CCL3 | miR-29c-3p target gene only | | NS | 3.08 | 0.028 | | NS |
| ENSG00000003989 | SLC7A2 | miR-29a-3p target gene only | | NS | 3.32 | 0.007 | | NS |
| ENSG00000078081 | LAMP3 | miR-29c-3p target gene only | | NS | 3.41 | 0.023 | | NS |
| ENSG00000117525 | F3 | miR-29c-3p target gene only | | NS | 3.45 | 0.001 | | NS |
| ENSG00000128342 | LIF | miR-29a/c-3p common target gene | | NS | 3.82 | 0.000 | | NS |
| ENSG00000123610 | TNFAIP6 | miR-29a-3p target gene only | | NS | 3.90 | 0.001 | | NS |
| ENSG00000276085 | CCL3L3 | miR-29c-3p target gene only | | NS | 4.37 | 0.006 | | NS |
| ENSG00000123689 | G0S2 | miR-29c-3p target gene only | | NS | 4.38 | 0.014 | | NS |
| ENSG00000164400 | CSF2 | miR-29c-3p target gene only | | NS | 4.40 | 0.006 | | NS |
| ENSG00000006210 | CX3CL1 | miR-29a/c-3p common target gene | | NS | 4.90 | 0.000 | | NS |
| ENSG00000172478 | C2orf54 | miR-29c-3p target gene only | -5.97 | 0.002 | -7.88 | 0.009 | | NS |
| ENSG00000123405 | NFE2 | miR-29c-3p target gene only | -4.68 | 0.003 | -4.76 | 0.040 | | NS |
| ENSG00000176490 | DIRAS1 | miR-29c-3p target gene only | -4.14 | 0.009 | -7.01 | 0.012 | | NS |
| ENSG00000171533 | MAP6 | miR-29a/c-3p common target gene | -4.03 | 0.001 | -4.05 | 0.032 | | NS |
| ENSG00000157542 | KCNJ6 | miR-29a/c-3p common target gene | -3.60 | 0.002 | -3.98 | 0.036 | | NS |
| ENSG00000153253 | SCN3A | miR-29a-3p target gene only | -3.58 | 0.038 | -6.01 | 0.042 | | NS |
| ENSG00000119938 | PPP1R3C | miR-29c-3p target gene only | -3.38 | 0.000 | -4.06 | 0.003 | | NS |
| ENSG00000198010 | DLGAP2 | miR-29a/c-3p common target gene | -3.32 | 0.013 | -4.85 | 0.014 | | NS |
| ENSG00000204103 | MAFB | miR-29a/c-3p common target gene | -3.10 | 0.003 | -3.12 | 0.044 | | NS |
| ENSG00000158445 | KCNB1 | miR-29a/c-3p common target gene | -2.84 | 0.006 | -3.10 | 0.044 | | NS |

| | | | | | | | |
|---|---|---|---|---|---|---|---|
| ENSG00000183091 | NEB | miR-29c-3p target gene only | -2.68 | 0.000 | -2.93 | 0.005 | NS |
| ENSG00000101057 | MYBL2 | miR-29a/c-3p common target gene | -2.48 | 0.014 | -3.33 | 0.020 | NS |
| ENSG00000184226 | PCDH9 | miR-29a/c-3p common target gene | -1.84 | 0.024 | -3.32 | 0.002 | NS |
| ENSG00000117461 | PIK3R3 | miR-29a-3p target gene only | -1.27 | 0.023 | -1.63 | 0.049 | NS |
| ENSG00000118432 | CNR1 | miR-29a-3p target gene only | -1.13 | 0.045 | -2.62 | 0.001 | NS |
| ENSG00000186204 | CYP4F12 | miR-29c-3p target gene only | | NS | -6.89 | 0.041 | NS |
| ENSG00000198626 | RYR2 | miR-29a/c-3p common target gene | | NS | -6.67 | 0.042 | NS |
| ENSG00000110203 | FOLR3 | miR-29c-3p target gene only | | NS | -6.59 | 0.019 | NS |
| ENSG00000084628 | NKAIN1 | miR-29a-3p target gene only | | NS | -6.52 | 0.039 | NS |
| ENSG00000173626 | TRAPPC3L | miR-29c-3p target gene only | | NS | -6.12 | 0.026 | NS |
| ENSG00000261115 | TMEM178B | miR-29a/c-3p common target gene | | NS | -5.16 | 0.041 | NS |
| ENSG00000114771 | AADAC | miR-29c-3p target gene only | | NS | -4.93 | 0.002 | NS |
| ENSG00000112562 | SMOC2 | miR-29c-3p target gene only | | NS | -4.86 | 0.022 | NS |
| ENSG00000073737 | DHRS9 | miR-29c-3p target gene only | | NS | -4.70 | 0.021 | NS |
| ENSG00000198074 | AKR1B10 | miR-29c-3p target gene only | | NS | -4.70 | 0.019 | NS |
| ENSG00000100505 | TRIM9 | miR-29a/c-3p common target gene | | NS | -4.42 | 0.010 | NS |
| ENSG00000214814 | FER1L6 | miR-29a-3p target gene only | | NS | -4.28 | 0.014 | NS |
| ENSG00000136235 | GPNMB | miR-29a-3p target gene only | | NS | -4.23 | 0.000 | NS |
| ENSG00000135547 | HEY2 | miR-29c-3p target gene only | | NS | -4.22 | 0.041 | NS |
| ENSG00000231274 | SBK3 | miR-29a/c-3p common target gene | | NS | -4.15 | 0.036 | NS |
| ENSG00000151150 | ANK3 | miR-29a/c-3p common target gene | | NS | -4.05 | 0.001 | NS |
| ENSG00000117152 | RGS4 | miR-29a/c-3p common target gene | | NS | -3.81 | 0.001 | NS |
| ENSG00000196569 | LAMA2 | miR-29a/c-3p common target gene | | NS | -3.73 | 0.007 | NS |
| ENSG00000085563 | ABCB1 | miR-29a-3p target gene only | | NS | -3.67 | 0.010 | NS |
| ENSG00000134321 | RSAD2 | miR-29c-3p target gene only | | NS | -3.53 | 0.035 | NS |
| ENSG00000102385 | DRP2 | miR-29a/c-3p common target gene | | NS | -3.51 | 0.048 | NS |
| ENSG00000172927 | MYEOV | miR-29c-3p target gene only | | NS | -3.33 | 0.041 | NS |
| ENSG00000154310 | TNIK | miR-29a/c-3p common target gene | | NS | -3.04 | 0.015 | NS |
| ENSG00000108984 | MAP2K6 | miR-29a/c-3p common target gene | | NS | -2.84 | 0.019 | NS |
| ENSG00000060718 | COL11A1 | miR-29a/c-3p common target gene | | NS | -2.77 | 0.049 | NS |
| ENSG00000164309 | CMYA5 | miR-29a/c-3p common target gene | | NS | -2.70 | 0.032 | NS |
| ENSG00000110876 | SELPLG | miR-29a-3p target gene only | | NS | -2.69 | 0.007 | NS |
| ENSG00000103942 | HOMER2 | miR-29c-3p target gene only | | NS | -2.58 | 0.045 | NS |
| ENSG00000087494 | PTHLH | miR-29c-3p target gene only | | NS | -2.47 | 0.017 | NS |
| ENSG00000139174 | PRICKLE1 | miR-29a-3p target gene only | | NS | -2.37 | 0.007 | NS |
| ENSG00000148483 | TMEM236 | miR-29a/c-3p common target gene | | NS | -2.31 | 0.047 | NS |
| ENSG00000064012 | CASP8 | miR-29a/c-3p common target gene | | NS | -2.19 | 0.003 | NS |
| ENSG00000004799 | PDK4 | miR-29a-3p target gene only | | NS | -2.11 | 0.026 | NS |
| ENSG00000082497 | SERTAD4 | miR-29a/c-3p common target gene | | NS | -2.00 | 0.037 | NS |
| ENSG00000168675 | LDLRAD4 | miR-29a/c-3p common target gene | | NS | -1.97 | 0.009 | NS |
| ENSG00000168916 | ZNF608 | miR-29c-3p target gene only | | NS | -1.94 | 0.008 | NS |
| ENSG00000079102 | RUNX1T1 | miR-29a/c-3p common target gene | | NS | -1.84 | 0.021 | NS |
| ENSG00000186310 | NAP1L3 | miR-29a/c-3p common target gene | | NS | -1.77 | 0.042 | NS |
| ENSG00000124588 | NQO2 | miR-29c-3p target gene only | | NS | -1.64 | 0.019 | NS |
| ENSG00000145990 | GFOD1 | miR-29a/c-3p common target gene | | NS | -1.58 | 0.028 | NS |
| ENSG00000162599 | NFIA | miR-29a/c-3p common target gene | | NS | -1.49 | 0.030 | NS |
| ENSG00000170836 | PPM1D | miR-29a/c-3p common target gene | | NS | -1.39 | 0.036 | NS |
| ENSG00000206557 | TRIM71 | miR-29a-3p target gene only | -6.44 | 0.011 | | NS | NS |
| ENSG00000205212 | CCDC144NL | miR-29a/c-3p common target gene | -6.06 | 0.011 | | NS | NS |
| ENSG00000135116 | HRK | miR-29a/c-3p common target gene | -5.62 | 0.013 | | NS | NS |
| ENSG00000168539 | CHRM1 | miR-29c-3p target gene only | -5.61 | 0.014 | | NS | NS |
| ENSG00000169174 | PCSK9 | miR-29c-3p target gene only | -5.25 | 0.039 | | NS | NS |
| ENSG00000157851 | DPYSL5 | miR-29a/c-3p common target gene | -4.98 | 0.025 | | NS | NS |
| ENSG00000110777 | POU2AF1 | miR-29c-3p target gene only | -4.92 | 0.032 | | NS | NS |
| ENSG00000115112 | TFCP2L1 | miR-29c-3p target gene only | -4.90 | 0.023 | | NS | NS |
| ENSG00000204655 | MOG | miR-29a/c-3p common target gene | -4.87 | 0.041 | | NS | NS |
| ENSG00000255690 | TRIL | miR-29a/c-3p common target gene | -4.84 | 0.037 | | NS | NS |
| ENSG00000175868 | CALCB | miR-29a/c-3p common target gene | -4.84 | 0.046 | | NS | NS |
| ENSG00000088002 | SULT2B1 | miR-29c-3p target gene only | -4.80 | 0.026 | | NS | NS |
| ENSG00000187288 | CIDEC | miR-29a/c-3p common target gene | -4.40 | 0.033 | | NS | NS |
| ENSG00000126218 | F10 | miR-29a-3p target gene only | -4.39 | 0.037 | | NS | NS |
| ENSG00000185818 | NAT8L | miR-29c-3p target gene only | -4.24 | 0.007 | | NS | NS |
| ENSG00000139219 | COL2A1 | miR-29a/c-3p common target gene | -4.21 | 0.007 | | NS | NS |
| ENSG00000167281 | RBFOX3 | miR-29a/c-3p common target gene | -4.12 | 0.008 | | NS | NS |
| ENSG00000006047 | YBX2 | miR-29c-3p target gene only | -4.12 | 0.020 | | NS | NS |
| ENSG00000196684 | HSH2D | miR-29c-3p target gene only | -3.94 | 0.012 | | NS | NS |
| ENSG00000164270 | HTR4 | miR-29c-3p target gene only | -3.91 | 0.030 | | NS | NS |
| ENSG00000186329 | TMEM212 | miR-29a/c-3p common target gene | -3.82 | 0.015 | | NS | NS |
| ENSG00000160460 | SPTBN4 | miR-29a/c-3p common target gene | -3.70 | 0.024 | | NS | NS |
| ENSG00000175920 | DOK7 | miR-29c-3p target gene only | -3.67 | 0.028 | | NS | NS |
| ENSG00000155980 | KIF5A | miR-29a/c-3p common target gene | -3.66 | 0.014 | | NS | NS |
| ENSG00000177459 | ERICH5 | miR-29c-3p target gene only | -3.65 | 0.029 | | NS | NS |
| ENSG00000118402 | ELOVL4 | miR-29a/c-3p common target gene | -3.51 | 0.028 | | NS | NS |
| ENSG00000197153 | HIST1H3J | miR-29c-3p target gene only | -3.45 | 0.015 | | NS | NS |
| ENSG00000163017 | ACTG2 | miR-29a-3p target gene only | -3.37 | 0.017 | | NS | NS |
| ENSG00000135423 | GLS2 | miR-29c-3p target gene only | -3.33 | 0.036 | | NS | NS |
| ENSG00000166664 | CHRFAM7A | miR-29a/c-3p common target gene | -3.30 | 0.008 | | NS | NS |
| ENSG00000092850 | TEKT2 | miR-29c-3p target gene only | -3.21 | 0.011 | | NS | NS |
| ENSG00000150656 | CNDP1 | miR-29a/c-3p common target gene | -3.11 | 0.035 | | NS | NS |
| ENSG00000273703 | HIST1H2BM | miR-29a-3p target gene only | -2.95 | 0.024 | | NS | NS |
| ENSG00000274641 | HIST1H2BO | miR-29a-3p target gene only | -2.93 | 0.008 | | NS | NS |
| ENSG00000240583 | AQP1 | miR-29a/c-3p common target gene | -2.86 | 0.000 | | NS | NS |
| ENSG00000091831 | ESR1 | miR-29a-3p target gene only | -2.75 | 0.002 | | NS | NS |
| ENSG00000138769 | CDKL2 | miR-29a/c-3p common target gene | -2.72 | 0.020 | | NS | NS |
| ENSG00000189143 | CLDN4 | miR-29c-3p target gene only | -2.65 | 0.041 | | NS | NS |
| ENSG00000171848 | RRM2 | miR-29a-3p target gene only | -2.65 | 0.005 | | NS | NS |
| ENSG00000143320 | CRABP2 | miR-29a/c-3p common target gene | -2.54 | 0.008 | | NS | NS |
| ENSG00000123612 | ACVR1C | miR-29c-3p target gene only | -2.44 | 0.017 | | NS | NS |
| ENSG00000149201 | CCDC81 | miR-29a-3p target gene only | -2.33 | 0.011 | | NS | NS |
| ENSG00000016391 | CHDH | miR-29a/c-3p common target gene | -2.22 | 0.047 | | NS | NS |
| ENSG00000117016 | RIMS3 | miR-29c-3p target gene only | -2.18 | 0.028 | | NS | NS |

| | | | | | | |
|---|---|---|---|---|---|---|
| ENSG00000127324 | TSPAN8 | miR-29a/c-3p common target gene | -2.17 | 0.044 | NS | NS |
| ENSG00000103056 | SMPD3 | miR-29a/c-3p common target gene | -2.15 | 0.039 | NS | NS |
| ENSG00000116741 | RGS2 | miR-29c-3p target gene only | -2.03 | 0.030 | NS | NS |
| ENSG00000132510 | KDM6B | miR-29a/c-3p common target gene | -1.95 | 0.031 | NS | NS |
| ENSG00000044574 | HSPA5 | miR-29a-3p target gene only | -1.90 | 0.010 | NS | NS |
| ENSG00000117472 | TSPAN1 | miR-29a/c-3p common target gene | -1.87 | 0.029 | NS | NS |
| ENSG00000170385 | SLC30A1 | miR-29a/c-3p common target gene | -1.79 | 0.003 | NS | NS |
| ENSG00000175866 | BAIAP2 | miR-29a/c-3p common target gene | -1.78 | 0.030 | NS | NS |
| ENSG00000130766 | SESN2 | miR-29c-3p target gene only | -1.70 | 0.018 | NS | NS |
| ENSG00000106462 | EZH2 | miR-29a/c-3p common target gene | -1.68 | 0.015 | NS | NS |
| ENSG00000162231 | NXF1 | miR-29a/c-3p common target gene | -1.61 | 0.011 | NS | NS |
| ENSG00000169155 | ZBTB43 | miR-29a/c-3p common target gene | -1.61 | 0.015 | NS | NS |
| ENSG00000108448 | TRIM16L | miR-29a/c-3p common target gene | -1.61 | 0.048 | NS | NS |
| ENSG00000131351 | HAUS8 | miR-29a/c-3p common target gene | -1.60 | 0.024 | NS | NS |
| ENSG00000101544 | ADNP2 | miR-29a/c-3p common target gene | -1.59 | 0.018 | NS | NS |
| ENSG00000166106 | ADAMTS15 | miR-29a-3p target gene only | -1.57 | 0.048 | NS | NS |
| ENSG00000198932 | GPRASP1 | miR-29a/c-3p common target gene | -1.53 | 0.024 | NS | NS |
| ENSG00000213047 | DENND1B | miR-29a/c-3p common target gene | -1.53 | 0.011 | NS | NS |
| ENSG00000069667 | RORA | miR-29a/c-3p common target gene | -1.47 | 0.036 | NS | NS |
| ENSG00000152495 | CAMK4 | miR-29a/c-3p common target gene | -1.47 | 0.048 | NS | NS |
| ENSG00000171988 | JMJD1C | miR-29a/c-3p common target gene | -1.41 | 0.006 | NS | NS |
| ENSG00000196428 | TSC22D2 | miR-29a/c-3p common target gene | -1.41 | 0.031 | NS | NS |
| ENSG00000047346 | FAM214A | miR-29a/c-3p common target gene | -1.41 | 0.042 | NS | NS |
| ENSG00000178053 | MLF1 | miR-29a/c-3p common target gene | -1.40 | 0.047 | NS | NS |
| ENSG00000154429 | CCSAP | miR-29a/c-3p common target gene | -1.38 | 0.047 | NS | NS |
| ENSG00000096717 | SIRT1 | miR-29a/c-3p common target gene | -1.38 | 0.018 | NS | NS |
| ENSG00000138759 | FRAS1 | miR-29a/c-3p common target gene | -1.31 | 0.028 | NS | NS |
| ENSG00000204569 | PPP1R10 | miR-29c-3p target gene only | -1.26 | 0.045 | NS | NS |
| ENSG00000151623 | NR3C2 | miR-29a/c-3p common target gene | -1.26 | 0.037 | NS | NS |
| ENSG00000111011 | RSRC2 | miR-29c-3p target gene only | -1.25 | 0.013 | NS | NS |
| ENSG00000092969 | TGFB2 | miR-29a/c-3p common target gene | -1.24 | 0.033 | NS | NS |
| ENSG00000112972 | HMGCS1 | miR-29a/c-3p common target gene | -1.19 | 0.036 | NS | NS |
| ENSG00000064309 | CDON | miR-29a-3p target gene only | -1.18 | 0.035 | NS | NS |
| ENSG00000145780 | FEM1C | miR-29a/c-3p common target gene | -1.17 | 0.030 | NS | NS |
| ENSG00000117000 | RLF | miR-29a/c-3p common target gene | -1.04 | 0.039 | NS | NS |
| ENSG00000068878 | PSME4 | miR-29a/c-3p common target gene | -1.01 | 0.049 | NS | NS |
| ENSG00000215301 | DDX3X | miR-29a/c-3p common target gene | -0.86 | 0.034 | NS | NS |
| ENSG00000077238 | IL4R | miR-29a/c-3p common target gene | 1.09 | 0.050 | NS | NS |
| ENSG00000151474 | FRMD4A | miR-29a/c-3p common target gene | 1.63 | 0.030 | NS | NS |
| ENSG00000067048 | DDX3Y | miR-29a/c-3p common target gene | 10.86 | 0.000 | NS | NS |

* NS: Positively detected but not statistically different in comparison.

**Table S4. PE-dysregulated miR-29a/c-3p target diseases and biological functions in female and male P0-HUVECs.***

| PE-dysregulated miR-29a/c-3p target diseases and biological functions | PE vs. NT in Female P0-HUVECs FDR P-value | PE vs. NT in Male P0-HUVECs FDR P-value |
|---|---|---|
| Seizures | 1.67E-04 | 1.91E-03 |
| Cell viability | 2.00E-05 | 3.73E-03 |
| Disorder of blood pressure | 1.39E-03 | 8.83E-03 |
| Angiogenesis | 6.64E-08 | 9.04E-03 |
| Cell movement | 4.32E-10 | 1.02E-02 |
| Concentration of Ca2+ | 1.42E-03 | 1.45E-02 |
| Vasculogenesis | 1.46E-06 | 1.58E-02 |
| Obesity | 8.73E-05 | 1.59E-02 |
| Atherosclerosis | 2.70E-04 | 2.39E-02 |
| Cellular infiltration | 1.88E-08 | NS |
| Recruitment of mononuclear leukocytes | 3.57E-06 | NS |
| Inflammatory response | 1.71E-05 | NS |
| Recruitment of monocytes | 1.80E-05 | NS |
| Invasion of cells | 2.52E-05 | NS |
| Migration of endothelial cells | 2.54E-05 | NS |
| Recruitment of phagocytes | 2.68E-05 | NS |
| Recruitment of lymphatic system cells | 5.04E-05 | NS |
| Recruitment of leukocytes | 5.44E-05 | NS |
| Recruitment of myeloid cells | 5.49E-05 | NS |
| Concentration of lipid | 7.33E-05 | NS |
| Recruitment of T lymphocytes | 9.04E-05 | NS |
| Concentration of fatty acid | 9.12E-05 | NS |
| Transmigration of mononuclear leukocytes | 1.03E-04 | NS |
| Accumulation of lipid | 1.35E-04 | NS |
| Cell death of endothelial cells | 1.47E-04 | NS |
| Concentration of triacylglycerol | 1.67E-04 | NS |
| Conversion of leukocytes | 1.80E-04 | NS |
| Seizure disorder | 2.86E-04 | NS |
| Conversion of thymocytes | 2.98E-04 | NS |
| Cell death of antigen presenting cells | 3.07E-04 | NS |
| Recruitment of Th1 cells | 3.31E-04 | NS |
| Apoptosis of antigen presenting cells | 4.39E-04 | NS |
| Apoptosis of endothelial cells | 4.43E-04 | NS |
| Chemotaxis | 4.43E-04 | NS |
| Adhesion of immune cells | 5.35E-04 | NS |
| Perinatal death | 6.84E-04 | NS |
| Disruption of microvascular endothelial cells | 7.03E-04 | NS |
| Atherosclerotic lesion | 1.44E-03 | NS |
| Malignant hypertension | NS | 7.28E-03 |
| Accumulation of extracellular matrix | NS | 1.02E-02 |
| Cardiogenesis of embryo | NS | 1.02E-02 |
| Chronic inflammatory disorder | NS | 1.02E-02 |
| Delay in initiation of conversion of inflammatory macrophages | NS | 1.02E-02 |
| Delay in initiation of conversion of M2 macrophages | NS | 1.02E-02 |
| Development of artery | NS | 1.02E-02 |
| Formation of vascular lesion | NS | 1.02E-02 |
| Oxidative stress injury of aorta | NS | 1.02E-02 |
| Growth of vessel | NS | 1.04E-02 |
| Conversion of heme | NS | 1.29E-02 |
| Entry into S phase of endothelial cell lines | NS | 1.29E-02 |
| Function of blood vessel | NS | 1.29E-02 |
| Hypertension | NS | 1.29E-02 |
| Inhibition of lymphokine activated killer cells | NS | 1.29E-02 |
| Outgrowth of vascular endothelial cells | NS | 1.56E-02 |
| Thickness of vessel tunica externa | NS | 1.56E-02 |
| Vasodilation of abdominal aorta | NS | 1.56E-02 |
| Pediatric obesity | NS | 1.58E-02 |
| Response of adaptive T-regulatory cells | NS | 1.83E-02 |
| Vasodilation of pulmonary artery | NS | 1.83E-02 |
| Concentration of 5-hydroxytryptamine | NS | 1.95E-02 |
| Abnormal function of cardiovascular system | NS | 2.04E-02 |

| | | |
|---|---|---|
| Adhesion of endothelial progenitor cells | NS | 2.04E-02 |
| Concentration of reactive oxygen species | NS | 2.04E-02 |
| Maturation of adipocytes | NS | 2.04E-02 |
| Remodeling of blood vessel | NS | 2.13E-02 |
| Cytotoxic reaction of macrophages | NS | 2.21E-02 |
| Growth of blood vessel | NS | 2.21E-02 |
| Disorganization of blood vessel | NS | 2.39E-02 |
| Stroke | NS | 2.39E-02 |

*NS: Not statistically Significant

**Table S5. MiR-29a/c-3p target diseases and biological functions enriched in PE-upregulated genes in female and male P0-HUVECs.**

| PE-upregulated miR-29a/c-3p target diseases and biological functions | PE vs. NT in Female P0-HUVECs FDR P-value | PE vs. NT in Male P0-HUVECs FDR P-value |
|---|---|---|
| * Angiogenesis | 3.73E-06 | 1.13E-02 |
| * Vasculogenesis | 7.78E-06 | 1.52E-02 |
| * Atherosclerosis | NS | 2.14E-02 |
| * Cell viability | 8.40E-04 | 9.91E-03 |
| * Concentration of Ca2+ | NS | 1.29E-02 |
| * Disorder of blood pressure | NS | 1.52E-02 |
| * Obesity | 1.36E-04 | NS |
| * Seizures | NS | 8.49E-04 |
| * Accumulation of lipid | 4.36E-05 | NS |
| * Cellular infiltration | 9.39E-06 | NS |
| * Invasion of cells | 9.39E-06 | NS |
| * Migration of endothelial cells | 9.52E-06 | NS |
| * Chemotaxis | 4.13E-05 | NS |
| * Inflammatory response | 6.43E-06 | NS |
| * Adhesion of immune cells | 4.58E-05 | NS |
| * Recruitment of mononuclear leukocytes | 1.33E-06 | NS |
| * Apoptosis of antigen presenting cells | 7.01E-04 | NS |
| * Apoptosis of endothelial cells | 3.66E-05 | NS |
| * Disruption of microvascular endothelial cells | 3.00E-04 | NS |
| * Perinatal death | 2.46E-03 | 2.81E-02 |
| * Accumulation of extracellular matrix | NS | 1.29E-02 |
| * Adhesion of endothelial progenitor cells | NS | 2.04E-02 |
| * Chronic inflammatory disorder | 2.80E-03 | 1.29E-02 |
| * Concentration of reactive oxygen species | NS | 2.24E-02 |
| * Remodeling of blood vessel | NS | 2.33E-02 |
| * Vasodilation of pulmonary artery | NS | 1.52E-02 |
| * Hypertension | NS | 2.57E-02 |
| * Stroke | NS | 1.29E-02 |
| * Permeability of blood vessel | 2.00E-03 | NS |
| * Abnormal function of cardiovascular system | NS | 2.18E-02 |
| * Atherosclerotic lesion | 2.34E-04 | NS |
| * Cardiogenesis of embryo | NS | 1.29E-02 |
| * Cell death of antigen presenting cells | 2.50E-04 | NS |
| * Concentration of fatty acid | 9.35E-04 | NS |
| * Concentration of lipid | 4.88E-03 | NS |
| * Concentration of triacylglycerol | 1.42E-03 | NS |
| * Conversion of heme | NS | 1.52E-02 |
| * Conversion of leukocytes | 3.09E-05 | NS |
| * Conversion of thymocytes | 1.36E-04 | NS |
| * Cytotoxic reaction of macrophages | NS | 2.04E-02 |
| * Delay in initiation of conversion of inflammatory macrophages | NS | 1.29E-02 |
| * Delay in initiation of conversion of M2 macrophages | NS | 1.29E-02 |
| * Development of artery | NS | 1.40E-02 |
| * Disorganization of blood vessel | NS | 2.57E-02 |
| * Entry into S phase of endothelial cell lines | NS | 1.52E-02 |
| * Formation of vascular lesion | NS | 1.29E-02 |
| * Function of blood vessel | NS | 1.52E-02 |
| * Growth of blood vessel | NS | 2.00E-02 |
| * Growth of vessel | NS | 1.29E-02 |
| * Inhibition of lymphokine activated killer cells | NS | 1.52E-02 |
| * Malignant hypertension | NS | 1.28E-02 |
| * Maturation of adipocytes | NS | 2.04E-02 |
| * Outgrowth of vascular endothelial cells | NS | 1.84E-02 |
| * Oxidative stress injury of aorta | NS | 1.29E-02 |
| * Recruitment of leukocytes | 3.50E-05 | NS |
| * Recruitment of monocytes | 2.35E-06 | NS |
| * Recruitment of myeloid cells | 6.90E-05 | NS |
| * Recruitment of phagocytes | 4.62E-05 | NS |
| * Recruitment of T lymphocytes | 7.36E-05 | NS |
| * Recruitment of Th1 cells | 2.56E-03 | NS |
| * Response of adaptive T-regulatory cells | NS | 2.04E-02 |
| * Thickness of vessel tunica externa | NS | 1.84E-02 |
| * Transmigration of mononuclear leukocytes | 9.35E-04 | NS |
| * Vasodilation of abdominal aorta | NS | 2.04E-02 |
| Epilepsy | NS | 2.34E-03 |
| Organismal death | 1.84E-07 | 5.00E-03 |
| Proliferation of smooth muscle cells | 1.91E-03 | 6.34E-03 |
| AMPA mediated synaptic current | NS | 6.34E-03 |
| Chronic respiratory disorder | NS | 6.34E-03 |
| Entry into S phase of carcinoma cell lines | NS | 6.34E-03 |
| Entry into S phase of lung cancer cell lines | NS | 6.34E-03 |
| Homeostasis of ion | NS | 6.34E-03 |
| Maturation of osteoblasts | NS | 6.34E-03 |
| Opioid-related disorder | NS | 7.10E-03 |
| Dyskinesia | NS | 9.16E-03 |
| Migration of smooth muscle cells | NS | 9.16E-03 |
| Multiple cancers | NS | 9.16E-03 |
| Peripheral vascular disease | NS | 9.16E-03 |

| | | |
|---|---|---|
| Landau Kleffner syndrome | NS | 9.91E-03 |
| Cell viability of tumor cell lines | NS | 1.28E-02 |
| Head and neck tumor | NS | 1.28E-02 |
| Inflammation of gastrointestinal tract | NS | 1.28E-02 |
| Breast or gynecological cancer | 9.78E-06 | 1.29E-02 |
| Pelvic cancer | 9.31E-05 | 1.29E-02 |
| Cancer of secretory structure | 4.34E-04 | 1.29E-02 |
| Diabetic complication | 1.49E-03 | 1.29E-02 |
| Breast or ovarian cancer | 2.15E-03 | 1.29E-02 |
| Breast or pancreatic cancer | 2.38E-03 | 1.29E-02 |
| Intraabdominal organ tumor | 3.94E-03 | 1.29E-02 |
| Epileptic seizure | NS | 1.29E-02 |
| Vascular calcification | NS | 1.29E-02 |
| Vasospasm | NS | 1.29E-02 |
| Familial psychiatric disease | NS | 1.29E-02 |
| Assembly of intercellular junctions | NS | 1.29E-02 |
| Formation of plasma membrane | NS | 1.29E-02 |
| Digestive system cancer | NS | 1.29E-02 |
| Adenoma | NS | 1.29E-02 |
| Aggregation of intestinal cell lines | NS | 1.29E-02 |
| Alcoholism | NS | 1.29E-02 |
| Arrest in cell cycle progression of blood cells | NS | 1.29E-02 |
| Arrest in cell cycle progression of blood platelets | NS | 1.29E-02 |
| Autosomal dominant mental retardation type 6 | NS | 1.29E-02 |
| Autosomal dominant osteogenesis imperfecta type I | NS | 1.29E-02 |
| Autosomal dominant supravalvular aortic stenosis | NS | 1.29E-02 |
| Autosomal recessive osteopetrosis type 3 | NS | 1.29E-02 |
| Binge eating disorder | NS | 1.29E-02 |
| Brain damage | NS | 1.29E-02 |
| Breakdown of heme | NS | 1.29E-02 |
| Cell movement of sperm | NS | 1.29E-02 |
| Chronic obstructive pulmonary disease | NS | 1.29E-02 |
| Combined osteogenesis imperfecta and Ehlers-Danlos syndrome type I | NS | 1.29E-02 |
| Congenital malformation of genital organs | NS | 1.29E-02 |
| Degenerative brain disorder | NS | 1.29E-02 |
| Delay in initiation of turnover of epidermal cells | NS | 1.29E-02 |
| Deposition of vascular smooth muscle cells | NS | 1.29E-02 |
| Disorder of basal ganglia | NS | 1.29E-02 |
| Doubling time of cervical cancer cell lines | NS | 1.29E-02 |
| Early infantile epileptic encephalopathy type 27 | NS | 1.29E-02 |
| Edema of brain | NS | 1.29E-02 |
| Entry into S phase | NS | 1.29E-02 |
| Entry into S phase of mesothelioma cell lines | NS | 1.29E-02 |
| Essential tremor | NS | 1.29E-02 |
| Excitatory postsynaptic potential | NS | 1.29E-02 |
| Experimental glomerular disease | NS | 1.29E-02 |
| Familial gigantism | NS | 1.29E-02 |
| Fibrosis of red pulp | NS | 1.29E-02 |
| Formation of aneurysm | NS | 1.29E-02 |
| Fragility of humerus | NS | 1.29E-02 |
| Functional hyposplenism | NS | 1.29E-02 |
| Generalized epilepsy | NS | 1.29E-02 |
| Generation of bilirubin | NS | 1.29E-02 |
| Growth failure or short stature | NS | 1.29E-02 |
| Growth of Ebola virus | NS | 1.29E-02 |
| Head and neck carcinoma | NS | 1.29E-02 |
| Heme oxygenase-1 deficiency | NS | 1.29E-02 |
| Huntington Disease | NS | 1.29E-02 |
| Hypertension of lung | NS | 1.29E-02 |
| Hypospadia | NS | 1.29E-02 |
| IMAGE syndrome | NS | 1.29E-02 |
| Injury of gastric mucosa | NS | 1.29E-02 |
| Jewett stage D prostatic carcinoma | NS | 1.29E-02 |
| Lesioning of renal glomerulus | NS | 1.29E-02 |
| Maturation of bone marrow-derived immature dendritic cells | NS | 1.29E-02 |
| Maturation of myoblasts | NS | 1.29E-02 |
| Maturation of satellite cells | NS | 1.29E-02 |
| Memory consolidation | NS | 1.29E-02 |
| Migration of Paneth cells | NS | 1.29E-02 |
| Migration of stomach cancer cell | NS | 1.29E-02 |
| Migration of vascular smooth muscle cells | NS | 1.29E-02 |
| Mild osteogenesis imperfecta type I | NS | 1.29E-02 |
| Nasodigitoacoustic syndrome | NS | 1.29E-02 |
| Neurovascular coupling | NS | 1.29E-02 |
| Osteogenesis imperfect type III/IV | NS | 1.29E-02 |
| Osteogenesis imperfecta type 2 thin-bone variant | NS | 1.29E-02 |
| Osteogenesis imperfecta type IIC | NS | 1.29E-02 |
| Oxidative stress response of melanoma cell lines | NS | 1.29E-02 |
| Progressive encephalopathy | NS | 1.29E-02 |
| Psychomotor agitation | NS | 1.29E-02 |
| Quantity of lung tissue | NS | 1.29E-02 |
| Reorganization of smooth muscle | NS | 1.29E-02 |

| | | |
|---|---|---|
| Replication of Japanese encephalitis virus | NS | 1.29E-02 |
| Respiratory insufficiency | NS | 1.29E-02 |
| Survival of brain tissue | NS | 1.29E-02 |
| Survival of smooth muscle cell lines | NS | 1.29E-02 |
| Thickness of ascending aorta | NS | 1.29E-02 |
| Tinnitus | NS | 1.29E-02 |
| Traumatic brain injury | NS | 1.29E-02 |
| Vasospasm of basilar artery | NS | 1.29E-02 |
| Secretion of molecule | NS | 1.32E-02 |
| Incidence of tumor | 1.81E-03 | 1.39E-02 |
| Abdominal cancer | 4.40E-03 | 1.39E-02 |
| Microtubule dynamics | 4.40E-03 | 1.39E-02 |
| Gastroenteritis | NS | 1.39E-02 |
| Liver tumor | NS | 1.39E-02 |
| Response of aorta | NS | 1.39E-02 |
| Rheumatic Disease | NS | 1.39E-02 |
| Associative learning | NS | 1.42E-02 |
| Non-malignant disorder | 6.17E-06 | 1.52E-02 |
| Cell movement of monocytes | 2.36E-04 | 1.52E-02 |
| Neuromuscular disease | 9.63E-04 | 1.52E-02 |
| Non-traumatic arthropathy | 1.33E-03 | 1.52E-02 |
| Ion homeostasis of cells | 4.28E-03 | 1.52E-02 |
| Hepato-pancreato-biliary cancer | NS | 1.52E-02 |
| Breast or gastric cancer | NS | 1.52E-02 |
| Morphogenesis of outflow tract | NS | 1.52E-02 |
| Abnormal morphology of cerebral cortex | NS | 1.52E-02 |
| Acidification of phagolysosome | NS | 1.52E-02 |
| Amyotrophic lateral sclerosis | NS | 1.52E-02 |
| Apoptosis of gastric mucous cells | NS | 1.52E-02 |
| Autosomal dominant Beckwith-Wiedemann syndrome | NS | 1.52E-02 |
| Autosomal dominant cutis laxa type 1 | NS | 1.52E-02 |
| Autosomal dominant osteogenesis imperfecta type III | NS | 1.52E-02 |
| Beat of sperm | NS | 1.52E-02 |
| Branching of axon branches | NS | 1.52E-02 |
| Cell transformation | NS | 1.52E-02 |
| Cell viability of chronic myelogenous leukemia cells | NS | 1.52E-02 |
| Classical phenylketonuria | NS | 1.52E-02 |
| Cleavage of heme | NS | 1.52E-02 |
| Congenital cortical hyperostosis | NS | 1.52E-02 |
| Cytotoxicity of nervous tissue cell lines | NS | 1.52E-02 |
| Damage of brain tissue | NS | 1.52E-02 |
| Dendritic growth/branching | NS | 1.52E-02 |
| Developmental delay and intractable seizure | NS | 1.52E-02 |
| Dominant perinatal lethal osteogenesis imperfecta | NS | 1.52E-02 |
| Early onset osteoarthritis | NS | 1.52E-02 |
| Ehlers-Danlos syndrome type VIIB | NS | 1.52E-02 |
| Endocrine carcinoma | NS | 1.52E-02 |
| Extensibility of carotid artery | NS | 1.52E-02 |
| Extension of axon branches | NS | 1.52E-02 |
| Familial aortic disorder | NS | 1.52E-02 |
| Fusion of Golgi membrane | NS | 1.52E-02 |
| Grade 3 glioma | NS | 1.52E-02 |
| Growth Failure | NS | 1.52E-02 |
| Growth of rhabdomyosarcoma | NS | 1.52E-02 |
| Healing of gastric mucosa | NS | 1.52E-02 |
| Homeostasis of bilirubin | NS | 1.52E-02 |
| Homeostasis of lymphatic fluid | NS | 1.52E-02 |
| Hypercalciuria | NS | 1.52E-02 |
| Hyperexcitation of hippocampal neurons | NS | 1.52E-02 |
| Inflammation of airway | NS | 1.52E-02 |
| Lennox-Gastaut syndrome | NS | 1.52E-02 |
| Long-term potentiation of dorsal striatum | NS | 1.52E-02 |
| Morphology of bone | NS | 1.52E-02 |
| Multiple epiphyseal dysplasia 1 | NS | 1.52E-02 |
| Muscularization of descending aorta | NS | 1.52E-02 |
| Nicotine dependence | NS | 1.52E-02 |
| Nicotine-mediated receptor current | NS | 1.52E-02 |
| Obsessive-compulsive disorder | NS | 1.52E-02 |
| Omphalocele | NS | 1.52E-02 |
| Orientation of smooth muscle cells | NS | 1.52E-02 |
| Osteogenesis imperfecta type IVB | NS | 1.52E-02 |
| Ovarian adenocarcinoma | NS | 1.52E-02 |
| Oxidation of heme | NS | 1.52E-02 |
| Oxidative stress response of macrophage cancer cell lines | NS | 1.52E-02 |
| Quantity of elastin in blood vessel | NS | 1.52E-02 |
| Serous adenocarcinoma in ovary | NS | 1.52E-02 |
| Short-term potentiation of hippocampus | NS | 1.52E-02 |
| Simpson-Golabi-Behmel syndrome type 1 | NS | 1.52E-02 |
| Spatial memory consolidation | NS | 1.52E-02 |
| Survival of thymic epithelial cells | NS | 1.52E-02 |
| Thickness of fibrous cap | NS | 1.52E-02 |
| Unilateral ureteral obstruction nephropathy | NS | 1.52E-02 |

| | | |
|---|---|---|
| Accumulation of tumor cell lines | NS | 1.59E-02 |
| Non-affective psychosis | NS | 1.67E-02 |
| Spatial learning | NS | 1.70E-02 |
| Thickness of connective tissue | NS | 1.70E-02 |
| Migration of carcinoma cell lines | 1.89E-05 | 1.72E-02 |
| Lymphatic system tumor | 3.66E-05 | 1.72E-02 |
| Malignant genitourinary solid tumor | 4.74E-05 | 1.72E-02 |
| Genital tract cancer | 5.34E-05 | 1.74E-02 |
| Syndromic encephalopathy | NS | 1.76E-02 |
| Edema | NS | 1.77E-02 |
| Damage of kidney | NS | 1.77E-02 |
| Serous adenocarcinoma | NS | 1.77E-02 |
| Premature aging | NS | 1.80E-02 |
| Attention deficit hyperactivity disorder | NS | 1.80E-02 |
| Adenocarcinoma | NS | 1.81E-02 |
| Proliferation of fibroblast cell lines | NS | 1.81E-02 |
| Genital tumor | 2.97E-05 | 1.84E-02 |
| Lymphoma | 3.09E-05 | 1.84E-02 |
| Partial seizure | NS | 1.84E-02 |
| Abnormal morphology of ureteric bud tip | NS | 1.84E-02 |
| Abnormal morphology of vestibular dark cells | NS | 1.84E-02 |
| Abnormality of endolymph | NS | 1.84E-02 |
| Abnormality of vestibular endolymph | NS | 1.84E-02 |
| Clustering of neurofilaments | NS | 1.84E-02 |
| Cognitive impairment | NS | 1.84E-02 |
| Contraction of arteriole | NS | 1.84E-02 |
| Delay in reassembly of Golgi apparatus | NS | 1.84E-02 |
| Delay in transmigration of smooth muscle cells | NS | 1.84E-02 |
| Disarray of myofibrils | NS | 1.84E-02 |
| Dissociation of neurofilaments | NS | 1.84E-02 |
| Ehlers-Danlos syndrome type VIIA | NS | 1.84E-02 |
| Entry into S phase of lung cell lines | NS | 1.84E-02 |
| Fragility of femur | NS | 1.84E-02 |
| Frontal lobe dementia | NS | 1.84E-02 |
| Inflammation of vascular smooth muscle cells | NS | 1.84E-02 |
| Long-term potentiation of prefrontal cortex | NS | 1.84E-02 |
| Morphology of RPE cells | NS | 1.84E-02 |
| Open-angle glaucoma | NS | 1.84E-02 |
| Operant self-administration of ethanol | NS | 1.84E-02 |
| Recovery of heart ventricle | NS | 1.84E-02 |
| Secretion of taurocholic acid | NS | 1.84E-02 |
| Semantic dementia | NS | 1.84E-02 |
| Symptomatic stage leptomeningeal metastasis | NS | 1.84E-02 |
| Symptomatic stage metastatic breast cancer | NS | 1.84E-02 |
| Tumorigenesis of epithelial neoplasm | NS | 1.84E-02 |
| Differentiation of connective tissue cells | 4.62E-05 | 1.85E-02 |
| Morphogenesis of dendritic spines | NS | 1.85E-02 |
| Coordination | NS | 1.85E-02 |
| Familial epilepsy | NS | 1.85E-02 |
| Dementia | NS | 1.85E-02 |
| Familial vascular disease | NS | 1.85E-02 |
| Subarachnoid hemorrhage | NS | 1.98E-02 |
| Anogenital cancer | 3.09E-05 | 2.00E-02 |
| Breast or ovarian carcinoma | NS | 2.00E-02 |
| Quantity of synapse | NS | 2.00E-02 |
| Thyroid carcinoma | NS | 2.00E-02 |
| Familial encephalopathy | NS | 2.00E-02 |
| Abnormal morphology of brain | NS | 2.00E-02 |
| Liver cancer | NS | 2.00E-02 |
| Social anxiety disorder | NS | 2.00E-02 |
| Frequency of tumor | NS | 2.03E-02 |
| Paralysis | NS | 2.03E-02 |
| Mature T-cell neoplasm | 7.10E-05 | 2.04E-02 |
| Progressive motor neuropathy | 2.94E-03 | 2.04E-02 |
| Vasodilation of artery | NS | 2.04E-02 |
| Streptozotocin-induced diabetic nephropathy | NS | 2.04E-02 |
| Abnormal morphology of disorganized barrel cortex | NS | 2.04E-02 |
| Accumulation of vascular smooth muscle cells | NS | 2.04E-02 |
| Activation of peripheral B lymphocytes | NS | 2.04E-02 |
| Acute kidney injury | NS | 2.04E-02 |
| Carcinoma | NS | 2.04E-02 |
| Cell death of hippocampal neurons | NS | 2.04E-02 |
| Concentration of oleoylethanolamide | NS | 2.04E-02 |
| Deformation of mice | NS | 2.04E-02 |
| Development of neurons | NS | 2.04E-02 |
| Development of peripherin inclusion | NS | 2.04E-02 |
| Epithelial-mesenchymal transition of AT2 cells | NS | 2.04E-02 |
| Familial neurological disorder | NS | 2.04E-02 |
| Maturation of fibroblast cell lines | NS | 2.04E-02 |
| Memory reconsolidation | NS | 2.04E-02 |
| Mingling of cells | NS | 2.04E-02 |
| Myhre syndrome | NS | 2.04E-02 |

| | | |
|---|---|---|
| Nephrotoxic acute renal failure | NS | 2.04E-02 |
| Opening of pore | NS | 2.04E-02 |
| Primary periodic paralysis | NS | 2.04E-02 |
| Progression of gastric carcinoma | NS | 2.04E-02 |
| Sensory processing | NS | 2.04E-02 |
| Swelling of perikaryon | NS | 2.04E-02 |
| Ulcerative colitis | NS | 2.04E-02 |
| Wiedemann-Rautenstrauch-like progeroid syndrome | NS | 2.04E-02 |
| Extrapancreatic malignant tumor | NS | 2.05E-02 |
| T-cell non-Hodgkin lymphoma | 7.54E-05 | 2.06E-02 |
| Rheumatoid arthritis | 1.10E-03 | 2.10E-02 |
| Conjunctivitis | NS | 2.11E-02 |
| Maturation of cells | NS | 2.13E-02 |
| Inflammation of joint | 2.67E-06 | 2.13E-02 |
| Morphology of nervous system | NS | 2.13E-02 |
| Fear | NS | 2.15E-02 |
| Congenital anomaly of skin | NS | 2.16E-02 |
| Delirium | NS | 2.22E-02 |
| Immune mediated inflammatory disease | 2.25E-08 | 2.24E-02 |
| Cancer of cells | 8.15E-06 | 2.24E-02 |
| Non-Hodgkin lymphoma | 1.63E-05 | 2.24E-02 |
| Pregnancy in diabetics | NS | 2.24E-02 |
| Congenital malformation of genitourinary system | NS | 2.24E-02 |
| Abdominal aortic aneurysm | NS | 2.24E-02 |
| Abnormal pruning of axons | NS | 2.24E-02 |
| Accumulation of macrophages | NS | 2.24E-02 |
| Adhesion of chondrocytes | NS | 2.24E-02 |
| Cell death of chronic myelogenous leukemia cells | NS | 2.24E-02 |
| Chronic phase rheumatoid arthritis | NS | 2.24E-02 |
| Development of thrombus | NS | 2.24E-02 |
| Developmental process of synapse | NS | 2.24E-02 |
| Formation of neurofilaments | NS | 2.24E-02 |
| Function of endothelial progenitor cells | NS | 2.24E-02 |
| Function of vestibular organ | NS | 2.24E-02 |
| Homeostasis of inorganic cation | NS | 2.24E-02 |
| Leakage of plasma | NS | 2.24E-02 |
| Learning | NS | 2.24E-02 |
| Length of primary neurites | NS | 2.24E-02 |
| Mass of right ventricle | NS | 2.24E-02 |
| Metastasis of cutaneous melanoma | NS | 2.24E-02 |
| Mobility of embryonic cell lines | NS | 2.24E-02 |
| Replication of Flaviviridae | NS | 2.24E-02 |
| Spontaneous fracture | NS | 2.24E-02 |
| Stenosis of pulmonary artery | NS | 2.24E-02 |
| Thickness of aorta wall | NS | 2.24E-02 |
| Transport of carbon dioxide | NS | 2.24E-02 |
| Post-traumatic stress disorder | NS | 2.25E-02 |
| Startle response | NS | 2.25E-02 |
| Hypotension | NS | 2.32E-02 |
| Formation of cellular inclusion bodies | NS | 2.32E-02 |
| Morphogenesis of cardiovascular system | NS | 2.33E-02 |
| Abnormal aortic valve physiology | NS | 2.33E-02 |
| Bipolar disorder | NS | 2.33E-02 |
| Schizophrenia | NS | 2.33E-02 |
| Long-term potentiation | NS | 2.34E-02 |
| Hepatobiliary carcinoma | NS | 2.34E-02 |
| Breast cancer | NS | 2.36E-02 |
| Quantity of metal | NS | 2.36E-02 |
| Neoplasia of cells | 3.05E-05 | 2.40E-02 |
| Quantity of neurons | NS | 2.41E-02 |
| T-cell malignant neoplasm | 6.30E-04 | 2.41E-02 |
| Prepulse inhibition | NS | 2.43E-02 |
| Stenosis of aorta | NS | 2.43E-02 |
| Inflammation of the large intestine | NS | 2.44E-02 |
| Benign lesion | 1.63E-05 | 2.44E-02 |
| Cigarette smoking-related carcinoma | NS | 2.44E-02 |
| Concentration of palmitoylethanolamide | NS | 2.44E-02 |
| Dissociation of microtubules | NS | 2.44E-02 |
| Hyperkalemic periodic paralysis | NS | 2.44E-02 |
| Idiopathic fetal growth restriction | NS | 2.44E-02 |
| Metastatic recurrent melanoma | NS | 2.44E-02 |
| Migration of osteosarcoma cells | NS | 2.44E-02 |
| Paramyotonia congenita | NS | 2.44E-02 |
| Pharmacologically induced seizure | NS | 2.44E-02 |
| Quantity of neurofilaments | NS | 2.44E-02 |
| Russell-Silver syndrome | NS | 2.44E-02 |
| Synaptogenesis of hippocampal neurons | NS | 2.44E-02 |
| Vomiting | NS | 2.44E-02 |
| Development of genital tumor | 2.96E-03 | 2.45E-02 |
| Postoperative pain | NS | 2.45E-02 |
| Development of palate | NS | 2.47E-02 |
| Female genital tract cancer | 6.68E-04 | 2.50E-02 |

| | | |
|---|---|---|
| Mature lymphocytic neoplasm | 1.17E-04 | 2.52E-02 |
| Early-onset neurological disorder | NS | 2.52E-02 |
| Formation of dendrites | NS | 2.52E-02 |
| Neuronal cell death | NS | 2.52E-02 |
| Quantity of neurites | NS | 2.52E-02 |
| Sickle cell anemia | NS | 2.55E-02 |
| Renal impairment | NS | 2.56E-02 |
| Abnormality of left ventricle | NS | 2.57E-02 |
| Accumulation of carcinoma cell lines | NS | 2.57E-02 |
| Accumulation of neurofilaments | NS | 2.57E-02 |
| Apoptosis of medium spiny neurons | NS | 2.57E-02 |
| Branching morphogenesis of ureter | NS | 2.57E-02 |
| Bronchopulmonary dysplasia | NS | 2.57E-02 |
| Delay in amyotrophic lateral sclerosis | NS | 2.57E-02 |
| Disorder of stature | NS | 2.57E-02 |
| Entry into S phase of epithelial cell lines | NS | 2.57E-02 |
| Familial partial lipodystrophy type 2 | NS | 2.57E-02 |
| Formation of paraxial mesoderm | NS | 2.57E-02 |
| Function of extraembryonic tissue | NS | 2.57E-02 |
| Function of thymus gland | NS | 2.57E-02 |
| Grade 3 malignant glioma | NS | 2.57E-02 |
| Hypertrophy | NS | 2.57E-02 |
| Hypokalemic periodic paralysis | NS | 2.57E-02 |
| Morphogenesis of notochord | NS | 2.57E-02 |
| Morphology of epithelial cells | NS | 2.57E-02 |
| Neuritogenesis | NS | 2.57E-02 |
| Overgrowth syndrome | NS | 2.57E-02 |
| Pervasive developmental disorder not otherwise specified | NS | 2.57E-02 |
| Plasticity of neurons | NS | 2.57E-02 |
| Progressive muscle atrophy | NS | 2.57E-02 |
| Quantity of heme | NS | 2.57E-02 |
| Sickle cell trait | NS | 2.57E-02 |
| Stabilization of atherosclerotic lesion | NS | 2.57E-02 |
| Survival of Saccharomyces cerevisiae | NS | 2.57E-02 |
| Tonicity of blood vessel | NS | 2.57E-02 |
| Abnormal morphology of axons | NS | 2.58E-02 |
| Secretion of steroid | NS | 2.61E-02 |
| Depressive disorder | NS | 2.63E-02 |
| Development of gastrointestinal tract | NS | 2.63E-02 |
| Diabetes mellitus | NS | 2.64E-02 |
| Asthma | NS | 2.65E-02 |
| Lymphocytic cancer | 7.81E-05 | 2.69E-02 |
| Malignant lymphocytic neoplasm | 7.81E-05 | 2.69E-02 |
| Cell movement of mononuclear leukocytes | 1.42E-04 | 2.69E-02 |
| Cell death of smooth muscle cells | NS | 2.75E-02 |
| Schizoaffective disorder | NS | 2.75E-02 |
| Tumorigenesis of reproductive tract | 4.99E-05 | 2.77E-02 |
| Liver carcinoma | NS | 2.77E-02 |
| Activation of dopaminergic neurons | NS | 2.77E-02 |
| Cell death of multilineage progenitor cells | NS | 2.77E-02 |
| Inflammation of aorta | NS | 2.77E-02 |
| Lymphangioleiomyomatosis | NS | 2.77E-02 |
| Metastasis of cervical cancer cell lines | NS | 2.77E-02 |
| Mineralization of fibroblast cell lines | NS | 2.77E-02 |
| Mobility of fibroblast cell lines | NS | 2.77E-02 |
| Primary lateral sclerosis | NS | 2.77E-02 |
| Stimulation of CD8+ T lymphocyte | NS | 2.77E-02 |
| WHO grade II glioma | NS | 2.77E-02 |
| WHO grade IV glioma | NS | 2.77E-02 |
| Apoptosis | 1.33E-06 | 2.78E-02 |
| Asthenozoospermia | NS | 2.78E-02 |
| Congenital heart disease | NS | 2.78E-02 |
| Formation of cellular protrusions | NS | 2.82E-02 |
| Cell viability of stomach cancer cell lines | NS | 2.87E-02 |
| Hematological or lymphatic system tumor | 1.35E-04 | 2.88E-02 |
| Laminopathy | NS | 2.89E-02 |
| Migration of squamous cell carcinoma cell lines | 2.76E-04 | 2.91E-02 |
| Lymphoreticular neoplasm | 2.94E-04 | 2.91E-02 |
| Homeostasis of H+ | NS | 2.91E-02 |
| Development of epithelial tissue | NS | 2.91E-02 |
| Abnormal morphology of optic tract | NS | 2.91E-02 |
| Arrest in proliferation of tumor cell lines | NS | 2.91E-02 |
| Carotid atherosclerosis | NS | 2.91E-02 |
| Cell rounding of epithelial cell lines | NS | 2.91E-02 |
| Chronic pain | NS | 2.91E-02 |
| Differentiation of adipocytes | NS | 2.91E-02 |
| Familial idiopathic basal ganglia calcification | NS | 2.91E-02 |
| Formation of neural crest | NS | 2.91E-02 |
| Function of vestibular system | NS | 2.91E-02 |
| Generalized convulsive epilepsy | NS | 2.91E-02 |
| Hyperalgesia of paw | NS | 2.91E-02 |
| Length of anogenital area | NS | 2.91E-02 |

| | | |
|---|---|---|
| Loss of brush border | NS | 2.91E-02 |
| Mild skin manifestation | NS | 2.91E-02 |
| Morphology of neurites | NS | 2.91E-02 |
| Pigment dispersion syndrome | NS | 2.91E-02 |
| Reperfusion injury of liver | NS | 2.91E-02 |
| Severe osteoarthritis | NS | 2.91E-02 |
| Tyrosine nitration of protein | NS | 2.91E-02 |
| WHO grade III glioma | NS | 2.91E-02 |
| Inflammation of organ | 6.43E-06 | 2.94E-02 |
| Cell viability of leukemia cell lines | NS | 2.95E-02 |
| Maturation of dendritic cells | NS | 2.95E-02 |
| Maturation of nervous tissue | NS | 2.95E-02 |
| Memory | NS | 2.95E-02 |
| Proliferation of vascular cells | NS | 2.96E-02 |
| Emotional behavior | NS | 2.97E-02 |
| Formation of solid tumor | 1.33E-03 | 3.02E-02 |
| Pelvic carcinoma | 1.38E-03 | 3.07E-02 |
| Degeneration of blood vessel | NS | 3.07E-02 |
| Abnormal morphology of Paneth cells | NS | 3.07E-02 |
| Accumulation of lung cancer cell lines | NS | 3.07E-02 |
| Arrest in cell cycle progression of hematopoietic cells | NS | 3.07E-02 |
| Community acquired pneumonia | NS | 3.07E-02 |
| Development of cutaneous melanoma | NS | 3.07E-02 |
| Development of myeloblasts | NS | 3.07E-02 |
| Drug seeking behavior | NS | 3.07E-02 |
| Dynamics of actin cytoskeleton | NS | 3.07E-02 |
| Familial generalized epilepsy | NS | 3.07E-02 |
| Fibrosis of dermis | NS | 3.07E-02 |
| Ischemic acute renal failure | NS | 3.07E-02 |
| Neoplasia of melanoma cell lines | NS | 3.07E-02 |
| Pustulosis palmaris et plantaris | NS | 3.07E-02 |
| Transformation of mammary epithelial cells | NS | 3.07E-02 |
| Fibrosis of liver | NS | 3.09E-02 |
| Transport of molecule | NS | 3.18E-02 |
| Long-term potentiation of cerebral cortex | NS | 3.23E-02 |
| Quantity of cells | 8.35E-04 | 3.27E-02 |
| Abnormal morphology of aorta wall | NS | 3.27E-02 |
| Binding of progesterone | NS | 3.27E-02 |
| Organ Degeneration | NS | 3.27E-02 |
| Migration of cells | 3.26E-09 | NS |
| Cell movement of tumor cell lines | 4.43E-08 | NS |
| Survival of organism | 5.22E-07 | NS |
| Migration of tumor cell lines | 1.20E-06 | NS |
| Leukocyte migration | 1.33E-06 | NS |
| Fibrosis | 1.33E-06 | NS |
| Quantity of macrophages | 1.33E-06 | NS |
| Chemotaxis of tumor cell lines | 1.33E-06 | NS |
| Cell movement of carcinoma cell lines | 1.33E-06 | NS |
| Inflammation of absolute anatomical region | 3.73E-06 | NS |
| Growth of tumor | 6.17E-06 | NS |
| Demyelination of spinal cord | 7.78E-06 | NS |
| Necrosis | 8.15E-06 | NS |
| Psoriasis | 8.15E-06 | NS |
| Cell movement of leukocytes | 8.57E-06 | NS |
| Necrosis of epithelial tissue | 8.57E-06 | NS |
| Modification of leukocytes | 9.18E-06 | NS |
| Cell death of carcinoma cell lines | 9.39E-06 | NS |
| Systemic autoimmune syndrome | 9.39E-06 | NS |
| Biosynthesis of amide | 9.78E-06 | NS |
| Lupus erythematosus | 1.53E-05 | NS |
| Apoptosis of carcinoma cell lines | 1.63E-05 | NS |
| Morphology of body cavity | 1.63E-05 | NS |
| Invasion of carcinoma cell lines | 2.21E-05 | NS |
| Invasion of tumor cell lines | 2.21E-05 | NS |
| Inflammation of body cavity | 2.71E-05 | NS |
| Cell movement of granulocytes | 3.01E-05 | NS |
| Cell movement of myeloid cells | 3.05E-05 | NS |
| Cellular homeostasis | 3.09E-05 | NS |
| Cell movement of phagocytes | 3.28E-05 | NS |
| Attraction of leukocytes | 3.50E-05 | NS |
| Osteoclastogenesis | 3.50E-05 | NS |
| Abnormal morphology of epithelial tissue | 3.50E-05 | NS |
| Hematologic cancer of cells | 3.56E-05 | NS |
| Activation of cells | 4.04E-05 | NS |
| Pelvic tumor | 4.13E-05 | NS |
| Binding of endothelial cells | 4.67E-05 | NS |
| Protein kinase cascade | 4.67E-05 | NS |
| Differentiation of osteoclasts | 4.67E-05 | NS |
| Invasion of lung cancer cell lines | 5.04E-05 | NS |
| Female genital neoplasm | 5.04E-05 | NS |
| Quantity of connective tissue | 5.31E-05 | NS |
| Systemic lupus erythematosus | 5.34E-05 | NS |

| | | |
|---|---|---|
| Apoptosis of bone marrow-derived dendritic cells | 5.79E-05 | NS |
| Migration of myeloid cells | 6.31E-05 | NS |
| Glioblastoma | 6.76E-05 | NS |
| Dermatitis | 7.75E-05 | NS |
| Adhesion of mononuclear leukocytes | 8.48E-05 | NS |
| Neoplasia of leukocytes | 9.06E-05 | NS |
| Cellular infiltration by leukocytes | 9.31E-05 | NS |
| Cell movement of lymphoma cell lines | 9.32E-05 | NS |
| Metastasis of cells | 9.44E-05 | NS |
| Function of antigen presenting cells | 1.00E-04 | NS |
| Synthesis of hyaluronic acid | 1.02E-04 | NS |
| Lymphocyte migration | 1.07E-04 | NS |
| Abnormal morphology of body cavity | 1.23E-04 | NS |
| Migration of endothelial cell lines | 1.35E-04 | NS |
| Cell movement of antigen presenting cells | 1.36E-04 | NS |
| Cell death of lung cancer cell lines | 1.36E-04 | NS |
| Familial porencephaly | 1.36E-04 | NS |
| Structural integrity of Reichert's membrane | 1.36E-04 | NS |
| Invasive tumor | 1.42E-04 | NS |
| Synthesis of fatty acid | 1.43E-04 | NS |
| Quantity of myeloid cells | 1.55E-04 | NS |
| Peripheral T-cell lymphoma | 1.55E-04 | NS |
| Function of phagocytes | 1.86E-04 | NS |
| Interaction of tumor cell lines | 1.95E-04 | NS |
| Clearance of cells | 2.05E-04 | NS |
| Benign solid tumor | 2.09E-04 | NS |
| Synthesis of lipid | 2.14E-04 | NS |
| Metastasis of tumor cell lines | 2.21E-04 | NS |
| Viral entry by HIV | 2.23E-04 | NS |
| Lipolysis of adipose tissue | 2.26E-04 | NS |
| Function of leukocytes | 2.26E-04 | NS |
| Cell movement of dendritic cells | 2.30E-04 | NS |
| Cell death of tumor cell lines | 2.36E-04 | NS |
| Cell proliferation of tumor cell lines | 2.37E-04 | NS |
| Hematologic cancer | 2.37E-04 | NS |
| Lipolysis of cells | 2.43E-04 | NS |
| Apoptosis of tumor cell lines | 2.47E-04 | NS |
| Cell movement of eosinophils | 2.50E-04 | NS |
| Cell movement of neutrophils | 2.53E-04 | NS |
| Nephritis | 2.66E-04 | NS |
| Neoplasia of carcinoma cell lines | 2.66E-04 | NS |
| Cerebral hemorrhage | 2.76E-04 | NS |
| Sepsis | 2.88E-04 | NS |
| Structural integrity of basement membrane | 3.00E-04 | NS |
| Killing of adenocarcinoma cell lines | 3.00E-04 | NS |
| Recruitment of M1 macrophages | 3.00E-04 | NS |
| Modification of neutrophils | 3.00E-04 | NS |
| Senescence of liver cell lines | 3.00E-04 | NS |
| Binding of monocytes | 3.24E-04 | NS |
| Apoptosis of lung cancer cell lines | 3.29E-04 | NS |
| Neoplasia of tumor cell lines | 3.29E-04 | NS |
| Morphology of digestive system | 3.46E-04 | NS |
| Development of bone marrow cells | 3.58E-04 | NS |
| Cell movement of T lymphocytes | 3.89E-04 | NS |
| Quantity of adipose tissue | 3.93E-04 | NS |
| Quantity of leukocytes | 3.99E-04 | NS |
| Tumorigenesis of lymphocytes | 3.99E-04 | NS |
| Damage of intestine | 4.00E-04 | NS |
| Transmigration of cells | 4.07E-04 | NS |
| Development of abdomen | 4.16E-04 | NS |
| Binding of professional phagocytic cells | 4.34E-04 | NS |
| Apoptosis of dendritic cells | 4.34E-04 | NS |
| Necrosis of vascular endothelial cells | 4.34E-04 | NS |
| Conversion of T lymphocytes | 4.34E-04 | NS |
| Osteoclastogenesis of bone marrow cells | 4.34E-04 | NS |
| Quantity of glycosylceramide | 4.34E-04 | NS |
| Anoikis of carcinoma cell lines | 4.78E-04 | NS |
| Single suture craniosynostosis | 4.78E-04 | NS |
| Loss of epithelial cells | 4.78E-04 | NS |
| Susceptibility to intracerebral hemorrhage | 5.07E-04 | NS |
| Production of myristic acid | 5.07E-04 | NS |
| Quantity of colfosceril palmitate | 5.07E-04 | NS |
| Retention of carbohydrate | 5.07E-04 | NS |
| Celiac disease | 5.11E-04 | NS |
| Chemotaxis of phagocytes | 5.38E-04 | NS |
| Colitis-associated cancer | 5.71E-04 | NS |
| Glucose tolerance | 5.72E-04 | NS |
| Metastasis of lung cancer cell lines | 5.75E-04 | NS |
| Mobilization of Ca2+ | 6.22E-04 | NS |
| Attraction of antigen presenting cells | 6.22E-04 | NS |
| Microangiopathy | 6.27E-04 | NS |
| Response of lymphatic system cells | 6.77E-04 | NS |

| | | |
|---|---|---|
| Cell death of macrophages | 6.83E-04 | NS |
| Cell movement of lung cancer cell lines | 6.83E-04 | NS |
| Metastasis of carcinoma cell lines | 6.83E-04 | NS |
| Abnormal morphology of abdomen | 6.83E-04 | NS |
| Myeloid or lymphoid neoplasm | 6.83E-04 | NS |
| Allergy | 6.89E-04 | NS |
| Expression of RNA | 7.00E-04 | NS |
| Binding of myeloid cells | 7.00E-04 | NS |
| Expansion of cells | 7.30E-04 | NS |
| Accumulation of carbohydrate | 7.30E-04 | NS |
| Phagocytosis by macrophages | 7.30E-04 | NS |
| Binding of tumor cell lines | 7.30E-04 | NS |
| Abnormality of peritoneum | 7.30E-04 | NS |
| Angioimmunoblastic T-cell lymphoma | 7.30E-04 | NS |
| Biosynthesis of stearic acid | 7.30E-04 | NS |
| Apoptosis of pneumocytes | 7.33E-04 | NS |
| Quantity of pulmonary alveolus | 7.33E-04 | NS |
| Uterine serous carcinoma | 7.33E-04 | NS |
| Genital carcinoma | 7.34E-04 | NS |
| Metabolism of acylglycerol | 7.86E-04 | NS |
| Cell death of phagocytes | 8.02E-04 | NS |
| Adhesion of endothelial cells | 8.18E-04 | NS |
| Transdifferentiation | 8.20E-04 | NS |
| Cell death of epithelial cells | 8.22E-04 | NS |
| Adhesion of phagocytes | 8.40E-04 | NS |
| Morphology of vessel | 8.64E-04 | NS |
| Neoplasia of blood cells | 8.64E-04 | NS |
| Vascular lesion | 8.70E-04 | NS |
| Benign oral disorder | 8.70E-04 | NS |
| Response of mononuclear leukocytes | 8.85E-04 | NS |
| Cell proliferation of carcinoma cell lines | 8.85E-04 | NS |
| Lipolysis of adipocytes | 8.85E-04 | NS |
| Viral life cycle | 9.35E-04 | NS |
| Cellular infiltration of dendritic cells | 9.36E-04 | NS |
| Immune response of leukocytes | 9.36E-04 | NS |
| Morphology of lung cells | 9.36E-04 | NS |
| Metabolism of membrane lipid derivative | 9.42E-04 | NS |
| Lymphopenia of bone marrow | 9.42E-04 | NS |
| Activation of enzyme | 9.42E-04 | NS |
| Aggregation of carbohydrate | 9.42E-04 | NS |
| Differentiation of dendritic precursor cells | 9.42E-04 | NS |
| Migration of plasmacytoid precursor dendritic cells | 9.42E-04 | NS |
| Bleeding | 9.43E-04 | NS |
| Immune response of phagocytes | 9.43E-04 | NS |
| Chemotaxis of eosinophils | 9.66E-04 | NS |
| Quantity of IL-1b in blood | 9.66E-04 | NS |
| Chemoattraction | 9.76E-04 | NS |
| Chemotaxis of neutrophils | 1.05E-03 | NS |
| Embryonal tumor | 1.05E-03 | NS |
| Endotoxin shock response | 1.07E-03 | NS |
| Accumulation of cells | 1.08E-03 | NS |
| Quantity of stem cells | 1.08E-03 | NS |
| Function of macrophages | 1.10E-03 | NS |
| Progressive neurological disorder | 1.10E-03 | NS |
| Chemotaxis of antigen presenting cells | 1.11E-03 | NS |
| Quantity of carbohydrate | 1.11E-03 | NS |
| Cell proliferation of breast cancer cell lines | 1.11E-03 | NS |
| Polyarticular juvenile rheumatoid arthritis | 1.11E-03 | NS |
| Neovascularization | 1.14E-03 | NS |
| Hematopoiesis of mononuclear leukocytes | 1.14E-03 | NS |
| Extravasation of cells | 1.14E-03 | NS |
| Migration of granulocytes | 1.16E-03 | NS |
| Fatty acid metabolism | 1.18E-03 | NS |
| Lung injury | 1.18E-03 | NS |
| Proliferation of tumor cells | 1.18E-03 | NS |
| Concentration of colfosceril palmitate | 1.18E-03 | NS |
| Migration of liver cell lines | 1.18E-03 | NS |
| Cell death of leukemia cell lines | 1.20E-03 | NS |
| Viral Infection | 1.26E-03 | NS |
| Attraction of mononuclear leukocytes | 1.26E-03 | NS |
| Formation of extracellular matrix | 1.26E-03 | NS |
| Migration of monocytes | 1.29E-03 | NS |
| Accumulation of lipid droplets | 1.33E-03 | NS |
| Damage of digestive system | 1.33E-03 | NS |
| Morphology of adipocytes | 1.33E-03 | NS |
| Morphology of white adipose tissue | 1.34E-03 | NS |
| Chemotaxis of monocytes | 1.38E-03 | NS |
| Chronic periodontal disease | 1.38E-03 | NS |
| Apoptosis of muscle cells | 1.38E-03 | NS |
| Proliferation of cancer cells | 1.40E-03 | NS |
| Adhesion of myeloid cells | 1.42E-03 | NS |
| Polyarthritis | 1.42E-03 | NS |

| | | |
|---|---|---|
| Immune response of myeloid cells | 1.44E-03 | NS |
| Cellular infiltration by monocytes | 1.45E-03 | NS |
| Leakage of blood-brain barrier | 1.45E-03 | NS |
| Glucose metabolism disorder | 1.48E-03 | NS |
| Apoptosis of lymphoma cell lines | 1.56E-03 | NS |
| I-kappaB kinase/NF-kappaB cascade | 1.61E-03 | NS |
| Abnormal morphology of adipose tissue | 1.68E-03 | NS |
| Chemotaxis of mononuclear leukocytes | 1.70E-03 | NS |
| Cell movement of monocyte-derived dendritic cells | 1.70E-03 | NS |
| Metabolism of triacylglycerol | 1.72E-03 | NS |
| Breast or colorectal cancer | 1.76E-03 | NS |
| Cytolysis of endothelial cells | 1.78E-03 | NS |
| Abnormal morphology of white adipocytes | 1.78E-03 | NS |
| Stimulation of osteoclast precursor cells | 1.78E-03 | NS |
| Abnormal metabolism | 1.82E-03 | NS |
| Quantity of Ca2+ | 1.82E-03 | NS |
| Synthesis of acylglycerol | 1.82E-03 | NS |
| Healing of lesion | 1.83E-03 | NS |
| Transmigration of monocytes | 1.85E-03 | NS |
| Progressive malignant solid tumor | 1.88E-03 | NS |
| NK cell proliferation | 1.91E-03 | NS |
| Adhesion of monocytes | 1.92E-03 | NS |
| Chemotaxis of lymphoma cell lines | 1.92E-03 | NS |
| Migration of lymphoma cell lines | 1.92E-03 | NS |
| Synthesis of polysaccharide | 1.92E-03 | NS |
| Synthesis of nitric oxide | 1.93E-03 | NS |
| Permeability of vascular system | 1.94E-03 | NS |
| Genitourinary carcinoma | 1.94E-03 | NS |
| Apoptosis of vascular endothelial cells | 2.00E-03 | NS |
| Proteinuria | 2.00E-03 | NS |
| Abdominal lesion | 2.00E-03 | NS |
| Apoptosis of phagocytes | 2.00E-03 | NS |
| Activation of blood cells | 2.00E-03 | NS |
| Differentiation of epithelial tissue | 2.00E-03 | NS |
| Apoptosis of monocyte-derived dendritic cells | 2.00E-03 | NS |
| Differentiation of proerythroblasts | 2.00E-03 | NS |
| Abnormal morphology of lymphoid organ | 2.00E-03 | NS |
| Advanced stage amyotrophic lateral sclerosis | 2.00E-03 | NS |
| Chemotaxis of leukocyte cell lines | 2.00E-03 | NS |
| Colorectal tumor | 2.00E-03 | NS |
| Dermatomyositis | 2.00E-03 | NS |
| Glomerulonephritis | 2.00E-03 | NS |
| Hematocrit | 2.00E-03 | NS |
| Intimal hyperplasia | 2.00E-03 | NS |
| Quantity of ganglioside GM1 | 2.00E-03 | NS |
| Synthesis of palmitic acid | 2.00E-03 | NS |
| Accumulation of myeloid cells | 2.02E-03 | NS |
| Female genital tract serous carcinoma | 2.04E-03 | NS |
| Pelvic serous carcinoma | 2.04E-03 | NS |
| Recruitment of neutrophils | 2.07E-03 | NS |
| Lymphoid hyperplasia | 2.07E-03 | NS |
| Transmigration of T lymphocytes | 2.07E-03 | NS |
| Remodeling of bone | 2.15E-03 | NS |
| Generation of tumor | 2.15E-03 | NS |
| Inflammation of peritoneum | 2.15E-03 | NS |
| Shape change of myeloid cells | 2.15E-03 | NS |
| Uterine serous papillary cancer | 2.21E-03 | NS |
| Transcription | 2.24E-03 | NS |
| Juvenile dermatomyositis | 2.24E-03 | NS |
| Foam cells | 2.24E-03 | NS |
| Recruitment of macrophages | 2.25E-03 | NS |
| Accumulation of triacylglycerol | 2.25E-03 | NS |
| Apoptosis of leukemia cell lines | 2.26E-03 | NS |
| Anoikis of lung cancer cell lines | 2.26E-03 | NS |
| Colitis-associated colon cancer | 2.26E-03 | NS |
| Loss of intestinal epithelial cells | 2.26E-03 | NS |
| Mobilization of bone marrow cells | 2.26E-03 | NS |
| Thickening of glomerular basement membrane | 2.26E-03 | NS |
| Autoimmune bullous skin disease | 2.26E-03 | NS |
| Progression of tumor | 2.28E-03 | NS |
| Nonhematologic malignant neoplasm | 2.29E-03 | NS |
| Attraction of phagocytes | 2.29E-03 | NS |
| Development of urinary tract | 2.32E-03 | NS |
| Migration of lung cancer cell lines | 2.33E-03 | NS |
| Accumulation of blood cells | 2.34E-03 | NS |
| Proliferation of muscle cells | 2.34E-03 | NS |
| Interaction of prostate cancer cell lines | 2.37E-03 | NS |
| Adhesion of lymphocytes | 2.44E-03 | NS |
| Morphology of blood vessel | 2.45E-03 | NS |
| Flux of Ca2+ | 2.46E-03 | NS |
| Diabetic nephropathy | 2.46E-03 | NS |
| Skin cancer | 2.52E-03 | NS |

| | | |
|---|---|---|
| Quantity of mononuclear leukocytes | 2.53E-03 | NS |
| Autosomal dominant Emery-Dreifuss muscular dystrophy | 2.55E-03 | NS |
| Migration of phagocytes | 2.55E-03 | NS |
| Viral entry | 2.61E-03 | NS |
| Response of myeloid leukocytes | 2.61E-03 | NS |
| Vascular leak syndrome | 2.64E-03 | NS |
| Reorganization of cytoskeleton | 2.66E-03 | NS |
| Metastatic solid tumor | 2.74E-03 | NS |
| Chemoattraction of leukocytes | 2.74E-03 | NS |
| Response of lymphocytes | 2.80E-03 | NS |
| Colony formation | 2.86E-03 | NS |
| Abnormal morphology of spleen | 2.89E-03 | NS |
| Mass of fat pad | 2.89E-03 | NS |
| Infection of vascular endothelial cells | 2.91E-03 | NS |
| Activation of alveolar macrophages | 2.91E-03 | NS |
| Concentration of eicosanoid | 2.93E-03 | NS |
| Mass of epigonadal fat pad | 2.94E-03 | NS |
| Apoptosis of epithelial cells | 3.14E-03 | NS |
| Killing of cells | 3.14E-03 | NS |
| Skin carcinoma | 3.15E-03 | NS |
| Quantity of sphingolipid | 3.15E-03 | NS |
| Cell death of bone marrow-derived macrophages | 3.15E-03 | NS |
| High grade astrocytoma | 3.22E-03 | NS |
| Hypoplasia of optic nerve | 3.26E-03 | NS |
| Bleeding of tissue | 3.26E-03 | NS |
| Arthritis of ankle joint | 3.26E-03 | NS |
| Proliferation of osteoclast precursor cells | 3.26E-03 | NS |
| Release of mitochondrial DNA | 3.26E-03 | NS |
| Runting | 3.26E-03 | NS |
| Urination disorder | 3.27E-03 | NS |
| Colorectal cancer | 3.34E-03 | NS |
| Ductal carcinoma | 3.34E-03 | NS |
| Metabolism of reactive oxygen species | 3.43E-03 | NS |
| Metastasis | 3.44E-03 | NS |
| Hypersensitive reaction | 3.44E-03 | NS |
| Proliferation of endothelial cells | 3.48E-03 | NS |
| Abnormal bone density | 3.49E-03 | NS |
| Inflammation of central nervous system | 3.54E-03 | NS |
| Cell movement of macrophages | 3.61E-03 | NS |
| Growth of lung | 3.73E-03 | NS |
| Damage of epithelial tissue | 3.76E-03 | NS |
| Apoptosis of leukocytes | 3.82E-03 | NS |
| Adhesion of tumor cell lines | 3.86E-03 | NS |
| Extrapulmonary squamous cell carcinoma | 3.90E-03 | NS |
| Abortion | 3.90E-03 | NS |
| Angiogenesis of tumor | 3.94E-03 | NS |
| Hematopoiesis of phagocytes | 3.94E-03 | NS |
| Viral entry by HIV-1 | 3.99E-03 | NS |
| Concentration of cholesterol | 3.99E-03 | NS |
| Degranulation | 3.99E-03 | NS |
| Cutaneous T-cell lymphoma | 3.99E-03 | NS |
| Delay in apoptosis of phagocytes | 3.99E-03 | NS |
| Immune response of splenocytes | 3.99E-03 | NS |
| Oxidation of NADPH | 3.99E-03 | NS |
| Pyroptosis of bone marrow-derived macrophages | 3.99E-03 | NS |
| Quantity of type II pneumocytes | 3.99E-03 | NS |
| Recruitment of CD8+ T lymphocyte | 3.99E-03 | NS |
| Development of genitourinary system | 3.99E-03 | NS |
| Quantity of neuroglia | 3.99E-03 | NS |
| Abnormal morphology of epithelial cells | 4.03E-03 | NS |
| Morphology of myeloid cells | 4.07E-03 | NS |
| Enlargement of spleen | 4.07E-03 | NS |
| Activation of leukocytes | 4.09E-03 | NS |
| Dupuytren contracture | 4.10E-03 | NS |
| Pemphigus foliaceus | 4.10E-03 | NS |
| Vascularization of absolute anatomical region | 4.27E-03 | NS |
| Invasive cancer | 4.27E-03 | NS |
| Development of body trunk | 4.28E-03 | NS |
| Genitourinary adenocarcinoma | 4.28E-03 | NS |
| Mass of genitourinary system | 4.28E-03 | NS |
| Adhesion of PBMCs | 4.31E-03 | NS |
| Transdifferentiation of cells | 4.31E-03 | NS |
| Accumulation of cholesterol | 4.31E-03 | NS |
| Active stage psoriasis | 4.31E-03 | NS |
| Differentiation of eosinophils | 4.31E-03 | NS |
| Mobilization of neutrophils | 4.31E-03 | NS |
| Lipolysis | 4.32E-03 | NS |
| Ductal breast carcinoma | 4.40E-03 | NS |
| Loss of hair | 4.40E-03 | NS |
| Organization of cytoskeleton | 4.40E-03 | NS |
| Extracranial solid tumor | 4.52E-03 | NS |
| Development of benign tumor | 4.57E-03 | NS |

| | | |
|---|---|---|
| Cellular infiltration by mononuclear leukocytes | 4.66E-03 | NS |
| Atopic dermatitis | 4.66E-03 | NS |
| Transdifferentiation of epithelial cells | 4.67E-03 | NS |
| Arrest in proliferation of leukemia cell lines | 4.67E-03 | NS |
| Development of peripheral blood leukocytes | 4.67E-03 | NS |
| Migration of fibroblast-like synoviocytes | 4.67E-03 | NS |
| Quantity of cerebroside | 4.67E-03 | NS |
| Stimulation of stem cells | 4.67E-03 | NS |
| Visual function | 4.67E-03 | NS |
| Ovarian tumor | 4.69E-03 | NS |
| Vascular tumor | 4.75E-03 | NS |
| Binding of gonadal cell lines | 4.76E-03 | NS |
| Concentration of choline-phospholipid | 4.76E-03 | NS |
| Dysgenesis | 4.82E-03 | NS |
| Necrosis of liver | 4.83E-03 | NS |
| Cell movement of cancer cells | 4.83E-03 | NS |
| Cellular infiltration by eosinophils | 4.88E-03 | NS |
| Development of head | 4.97E-03 | NS |
| Abnormal morphology of blood vessel | 4.98E-03 | NS |
| Quantity of cytokine | 5.04E-03 | NS |
| Attraction of monocytes | 5.06E-03 | NS |
| Quantity of inflammatory leukocytes | 5.06E-03 | NS |
| Primary ovarian cancer | 5.17E-03 | NS |
| Immune response of cells | 5.25E-03 | NS |
| Binding of DNA | 5.25E-03 | NS |
| Quantity of TNF in blood | 5.29E-03 | NS |
| Abnormal morphology of macrophages | 5.29E-03 | NS |
| Urological disorder of kidney | 5.29E-03 | NS |
| Synthesis of carbohydrate | 5.33E-03 | NS |
| Killing of bacteria | 5.45E-03 | NS |
| Pelvic adenocarcinoma | 5.48E-03 | NS |
| Hematoma | 5.48E-03 | NS |
| Quantity of alveolar macrophages | 5.48E-03 | NS |
| Stimulation of hematopoietic progenitor cells | 5.48E-03 | NS |
| Infection by Flaviviridae | 5.49E-03 | NS |
| Quantity of protein in blood | 5.52E-03 | NS |
| Colon tumor | 5.52E-03 | NS |
| Growth of renal glomerulus | 5.52E-03 | NS |
| Proliferation of neuroblasts | 5.52E-03 | NS |
| Synthesis of triacylglycerol | 5.52E-03 | NS |
| Activation of lymphocytes | 5.54E-03 | NS |
| Apoptosis of leukocyte cell lines | 5.61E-03 | NS |
| Cellular infiltration by granulocytes | 5.87E-03 | NS |
| Cell movement of connective tissue cells | 5.87E-03 | NS |
| Blindness | 5.87E-03 | NS |
| Apoptosis of endometrial cancer cell lines | 5.87E-03 | NS |

*: Also enriched in analysis with all PE-dysregulated miR-29a/c-3p target genes; NS: Not statistically Significant.

**Table S6. MiR-29a/c-3p target diseases and biological functions enriched in PE-downregulated genes in female and male P0-HUVECs.**

| PE-downregulated miR-29a/c-3p target diseases and biological functions | PE vs. NT in Female P0-HUVECs FDR P-value | PE vs. NT in Male P0-HUVECs FDR P-value |
|---|---|---|
| *Seizures | 1.90E-04 | NS |
| *Concentration of Ca2+ | 1.57E-04 | NS |
| *Angiogenesis | 1.32E-02 | NS |
| *Cell viability | 1.75E-02 | NS |
| *Concentration of lipid | 1.36E-02 | NS |
| *Cell movement | 5.04E-03 | 3.90E-02 |
| *Pediatric obesity | NS | 2.95E-02 |
| *Concentration of 5-hydroxytryptamine | NS | 3.14E-02 |
| *Seizure disorder | 2.09E-04 | NS |
| Branching of cells | 8.15E-03 | NS |
| Formation of cellular protrusions | 1.10E-02 | NS |
| Cellular infiltration by granulocytes | 1.57E-02 | NS |
| Migration of cells | 3.78E-03 | NS |
| Migration of monocytes | NS | 3.80E-02 |
| Oscillation of Ca2+ | 7.71E-03 | NS |
| Transport of Ca2+ | 1.40E-02 | NS |
| Hypotension | 7.25E-04 | NS |
| Schizophrenia | 3.02E-06 | 2.87E-03 |
| Oxidation of 2-phenethylamine | NS | 2.87E-03 |
| Oxidation of 5-hydroxytryptamine | NS | 2.87E-03 |
| Oxidation of epinephrine | NS | 2.87E-03 |
| Oxidation of norepinephrine | NS | 2.87E-03 |
| Accumulation of 5-hydroxytryptamine | NS | 2.87E-03 |
| Braak stage I-II presymptomatic Alzheimer's disease | NS | 2.87E-03 |
| Brunner syndrome | NS | 2.87E-03 |
| Cheese reaction | NS | 2.87E-03 |
| Concentration of dopamine | NS | 2.87E-03 |
| Deamination of dopamine | NS | 2.87E-03 |
| Deamination of norepinephrine | NS | 2.87E-03 |
| Efficacy of mirtazapine | NS | 2.87E-03 |
| Expansion of thalamocortical axons | NS | 2.87E-03 |
| Fusion of sensory cortex | NS | 2.87E-03 |
| Metabolism of 2-phenethylamine | NS | 2.87E-03 |
| Metabolism of dopamine | NS | 2.87E-03 |
| Non-endogenous depressive disorder | NS | 2.87E-03 |
| Organization of axon terminals | NS | 2.87E-03 |
| Quantity of apoptotic neuroepithelial cells | NS | 2.87E-03 |
| Quantity of thalamocortical axons | NS | 2.87E-03 |
| Segregation of sensory projections | NS | 2.87E-03 |
| Susceptibility to antisocial behavior | NS | 2.87E-03 |
| Braak stage VI symptomatic Alzheimer's disease | NS | 3.81E-03 |
| Cell survival of cortical astrocytes | NS | 3.81E-03 |
| Patterning of thalamocortical axons | NS | 3.81E-03 |
| Oxidation of dopamine | NS | 4.56E-03 |
| Clearance of dopamine | NS | 4.56E-03 |
| Olfactory-discrimination memory | NS | 4.56E-03 |
| Arrest in spermatogenesis of spermatocytes | NS | 4.59E-03 |
| Atrophia bulborum hereditaria | NS | 4.59E-03 |
| Cell death of neuroblastoma cell lines | NS | 4.59E-03 |
| Inattentiveness | NS | 4.59E-03 |
| Neurotic depressive disorder | NS | 4.59E-03 |
| Regulation of neurotransmitter | NS | 4.59E-03 |
| Size of infarct | NS | 4.59E-03 |
| Abnormal morphology of substantia nigra | NS | 5.78E-03 |
| Catabolism of dopamine | NS | 5.78E-03 |
| Exogenous obesity | NS | 5.78E-03 |
| Morphology of somatosensory cortex | NS | 5.78E-03 |
| Familial psychiatric disease | NS | 5.91E-03 |
| Stress response of mice | NS | 6.29E-03 |
| Dysfunction of mice | NS | 6.91E-03 |
| Inhibition of cyclic AMP | NS | 7.34E-03 |
| Social transmission of food preference | NS | 7.34E-03 |
| Emotional behavior | NS | 8.91E-03 |
| Metabolism of 5-hydroxytryptamine | NS | 9.13E-03 |
| Catalepsy | NS | 1.01E-02 |
| Quantity of prostate cancer cell lines | NS | 1.01E-02 |
| Emotional lability | NS | 1.06E-02 |

| | | |
|---|---|---|
| Movement Disorders | 5.47E-03 | 1.06E-02 |
| Syndromic behavioral deficit | NS | 1.18E-02 |
| Systolic pressure of left ventricle | NS | 1.18E-02 |
| Oppositional defiant disorder | NS | 1.28E-02 |
| Methamphetamine dependence | NS | 1.50E-02 |
| Apoptosis of spermatocytes | NS | 1.50E-02 |
| Uptake of dopamine | NS | 1.50E-02 |
| Narcolepsy | NS | 1.51E-02 |
| Synthesis of dopamine | NS | 1.51E-02 |
| Early-onset high myopia | NS | 1.54E-02 |
| Learning | 1.57E-02 | 1.69E-02 |
| Familial encephalopathy | NS | 1.80E-02 |
| Neurodegeneration of dopaminergic neurons | NS | 1.80E-02 |
| Swimming behavior | NS | 1.80E-02 |
| Gliosis of cerebral cortex | NS | 1.83E-02 |
| Morphology of brain | 5.04E-03 | 2.05E-02 |
| Autophagy of epithelial cell lines | NS | 2.05E-02 |
| Autophagy of kidney cell lines | NS | 2.05E-02 |
| Function of cardiomyocytes | NS | 2.05E-02 |
| Cell death of epithelial cells | 6.10E-03 | 2.08E-02 |
| Quantity of epinephrine | NS | 2.13E-02 |
| Panic disorder | NS | 2.20E-02 |
| Gout | NS | 2.21E-02 |
| Inhibitory postsynaptic current | NS | 2.45E-02 |
| Locomotor activity | NS | 2.45E-02 |
| Low-grade lymphoma | NS | 2.45E-02 |
| Obsessive-compulsive disorder | NS | 2.65E-02 |
| Ventricular fibrillation | NS | 2.70E-02 |
| M2 polarization | NS | 2.79E-02 |
| Tourette syndrome | NS | 2.80E-02 |
| Huntington Disease | NS | 2.84E-02 |
| Quantity of sperm | NS | 2.93E-02 |
| Activation of astrocytes | NS | 2.95E-02 |
| Attention deficit hyperactivity disorder | NS | 2.95E-02 |
| Autophagy of embryonic cell lines | NS | 2.95E-02 |
| Concentration of norepinephrine | NS | 2.95E-02 |
| Growth of cholangiocarcinoma | NS | 2.95E-02 |
| Aggressive behavior | NS | 3.07E-02 |
| Endoplasmic reticulum stress response of cells | NS | 3.15E-02 |
| Formation of reactive oxygen species | NS | 3.15E-02 |
| Production of hydrogen peroxide | NS | 3.15E-02 |
| Ventricular tachycardia | NS | 3.15E-02 |
| Grooming | NS | 3.15E-02 |
| Volume of infarct | NS | 3.15E-02 |
| Response of heart | NS | 3.62E-02 |
| Apoptosis of cortical neurons | NS | 3.72E-02 |
| Body mass index | NS | 3.77E-02 |
| Contextual conditioning | NS | 3.77E-02 |
| B-cell non-Hodgkin lymphoma | NS | 3.90E-02 |
| Concentration of glutathione | NS | 4.07E-02 |
| Cognitive impairment | 6.13E-05 | 4.16E-02 |
| Benign solid tumor | NS | 4.16E-02 |
| Alcoholism | NS | 4.35E-02 |
| Activation of Adenylate cyclase | NS | 4.42E-02 |
| Metabolism of cholesterol | NS | 4.42E-02 |
| Apoptosis of neuroblastoma cell lines | NS | 4.43E-02 |
| Function of neurons | NS | 4.54E-02 |
| Lymphoreticular neoplasm | 2.09E-04 | 4.64E-02 |
| Hyperactive behavior | NS | 4.80E-02 |
| Cell movement of sperm | NS | 4.85E-02 |
| Sporadic amyotrophic lateral sclerosis | NS | 5.14E-02 |
| Atrial fibrillation | NS | 5.49E-02 |
| Mature B cell malignant tumor | NS | 5.54E-02 |
| Invasion of prostate cancer cell lines | NS | 5.75E-02 |
| Hypoactivity of mice | NS | 5.78E-02 |
| Eating Disorders | NS | 6.00E-02 |
| Cancer of head | 1.57E-02 | 6.04E-02 |
| Migraines | NS | 6.04E-02 |
| Concentration of ATP | NS | 6.09E-02 |
| Apoptosis of cardiomyocytes | NS | 6.76E-02 |
| Hemangioma | NS | 7.01E-02 |
| Spatial learning | NS | 7.01E-02 |

| | | |
|---|---|---|
| Major depression | 1.21E-02 | 7.54E-02 |
| Coordination | NS | 7.54E-02 |
| Signal transduction | NS | 7.54E-02 |
| Skin cancer | 3.84E-07 | NS |
| Cutaneous melanoma | 1.76E-06 | NS |
| Urinary tract cancer | 1.76E-06 | NS |
| Melanoma | 1.02E-05 | NS |
| Basal cell carcinoma | 2.68E-05 | NS |
| Skin carcinoma | 2.74E-05 | NS |
| Genitourinary tumor | 6.24E-05 | NS |
| Bladder cancer | 8.89E-05 | NS |
| Progressive neurological disorder | 1.90E-04 | NS |
| Action potential of cells | 2.09E-04 | NS |
| Ductal carcinoma | 2.09E-04 | NS |
| Prostatic carcinoma | 2.24E-04 | NS |
| Prostatic adenocarcinoma | 2.60E-04 | NS |
| Organization of cells | 2.76E-04 | NS |
| Bladder carcinoma | 2.76E-04 | NS |
| Malignant genitourinary solid tumor | 2.96E-04 | NS |
| Malignant neoplasm of retroperitoneum | 2.96E-04 | NS |
| Pelvic tumor | 2.96E-04 | NS |
| Prostate cancer | 2.96E-04 | NS |
| Genital tumor | 3.31E-04 | NS |
| Extracranial solid tumor | 3.45E-04 | NS |
| HER2 negative hormone receptor negative breast cancer | 3.59E-04 | NS |
| Genitourinary adenocarcinoma | 3.74E-04 | NS |
| Development of neural cells | 3.84E-04 | NS |
| Respiratory system tumor | 4.66E-04 | NS |
| Cancer | 5.06E-04 | NS |
| Neurotransmission | 5.08E-04 | NS |
| Abdominal neoplasm | 5.08E-04 | NS |
| Cancer of secretory structure | 5.08E-04 | NS |
| Extrapancreatic malignant tumor | 5.92E-04 | NS |
| Gastrointestinal adenocarcinoma | 6.33E-04 | NS |
| Anogenital cancer | 7.17E-04 | NS |
| Development of neurons | 7.83E-04 | NS |
| Excitatory postsynaptic potential | 7.83E-04 | NS |
| Non-small cell lung carcinoma | 7.83E-04 | NS |
| Malignant neoplasm of respiratory system | 8.30E-04 | NS |
| Neoplasia of blood cells | 8.49E-04 | NS |
| Postoperative pain | 8.84E-04 | NS |
| Cell transformation | 8.91E-04 | NS |
| Acute leukemia | 8.91E-04 | NS |
| Nonhematologic malignant neoplasm | 8.91E-04 | NS |
| Lung tumor | 9.56E-04 | NS |
| Liquid tumor | 9.69E-04 | NS |
| Intraabdominal organ tumor | 1.03E-03 | NS |
| Arrest in differentiation of cancer cells | 1.04E-03 | NS |
| Delay in fragmentation of DNA | 1.04E-03 | NS |
| Desensitization of hippocampal neurons | 1.04E-03 | NS |
| Extraadrenal retroperitoneal tumor | 1.04E-03 | NS |
| Pelvic cancer | 1.05E-03 | NS |
| Genital tract cancer | 1.06E-03 | NS |
| Incidence of tumor | 1.06E-03 | NS |
| Uterine tumor | 1.23E-03 | NS |
| Adenocarcinoma | 1.23E-03 | NS |
| Autism spectrum disorder or intellectual disability | 1.31E-03 | NS |
| Epilepsy or neurodevelopmental disorder | 1.31E-03 | NS |
| Long-term recognition memory | 1.31E-03 | NS |
| Hematologic cancer | 1.33E-03 | NS |
| Genitourinary carcinoma | 1.39E-03 | NS |
| Myeloid or lymphoid neoplasm | 1.54E-03 | NS |
| Lung cancer | 1.62E-03 | NS |
| Lung carcinoma | 1.73E-03 | NS |
| Tauopathy | 1.73E-03 | NS |
| Secretion of molecule | 1.84E-03 | NS |
| Progressive encephalopathy | 1.84E-03 | NS |
| Familial generalized epilepsy | 1.90E-03 | NS |
| Mature lymphocytic neoplasm | 1.90E-03 | NS |
| Bone marrow lesion | 2.02E-03 | NS |
| Quantity of cells | 2.03E-03 | NS |
| Sézary syndrome | 2.21E-03 | NS |

| | | |
|---|---|---|
| Development of genital tumor | 2.23E-03 | NS |
| Clear-cell adenocarcinoma | 2.32E-03 | NS |
| COVID-19 | 2.32E-03 | NS |
| Differentiation of myeloid leukocytes | 2.32E-03 | NS |
| Digestive organ tumor | 2.32E-03 | NS |
| Ductal breast carcinoma | 2.32E-03 | NS |
| Epilepsy | 2.32E-03 | NS |
| Genital carcinoma | 2.32E-03 | NS |
| Leukemia | 2.32E-03 | NS |
| Lymphoproliferative disorder of the skin | 2.32E-03 | NS |
| Tubulation of epithelial cells | 2.32E-03 | NS |
| Tumorigenesis of tissue | 2.32E-03 | NS |
| Autosomal dominant epilepsy | 2.40E-03 | NS |
| Gastrointestinal tract cancer | 2.48E-03 | NS |
| Development of adenocarcinoma | 2.64E-03 | NS |
| Frequency of tumor | 2.69E-03 | NS |
| Granulopoiesis | 2.69E-03 | NS |
| Pancreaticobiliary cancer | 2.69E-03 | NS |
| Renal lesion | 2.69E-03 | NS |
| Hepato-pancreato-biliary cancer | 2.71E-03 | NS |
| Autosomal dominant early infantile epileptic encephalopathy | 2.71E-03 | NS |
| Breast or gynecological cancer | 2.71E-03 | NS |
| Large intestine adenocarcinoma | 2.71E-03 | NS |
| Pelvic carcinoma | 2.71E-03 | NS |
| Neoplasia of cells | 2.77E-03 | NS |
| Bone marrow cancer | 2.77E-03 | NS |
| Dentin disease | 2.77E-03 | NS |
| Progressive motor neuropathy | 2.85E-03 | NS |
| Hepatobiliary neoplasm | 2.95E-03 | NS |
| Blue round small cell tumor | 2.95E-03 | NS |
| Musculoskeletal pain | 2.95E-03 | NS |
| Non-melanoma solid tumor | 2.95E-03 | NS |
| Pelvic adenocarcinoma | 2.95E-03 | NS |
| Recognition memory | 2.95E-03 | NS |
| Tumorigenesis of reproductive tract | 2.95E-03 | NS |
| Female genital neoplasm | 2.96E-03 | NS |
| Gastroesophageal adenocarcinoma | 2.96E-03 | NS |
| Action potential of neurons | 3.14E-03 | NS |
| Familial epilepsy | 3.14E-03 | NS |
| Development of carcinoma | 3.22E-03 | NS |
| Pancreaticobiliary carcinoma | 3.27E-03 | NS |
| Transport of cation | 3.35E-03 | NS |
| Synthesis of 2-arachidonoylglycerol | 3.35E-03 | NS |
| Colorectal adenocarcinoma | 3.41E-03 | NS |
| Organismal death | 3.45E-03 | NS |
| Invasive tumor | 3.45E-03 | NS |
| Non-Hodgkin lymphoma | 3.48E-03 | NS |
| Mature T-cell or NK-cell neoplasm | 3.52E-03 | NS |
| Renal clear cell adenocarcinoma | 3.56E-03 | NS |
| Autism or intellectual disability | 3.74E-03 | NS |
| Transport of ion | 3.99E-03 | NS |
| Cell spreading of embryonic cells | 4.11E-03 | NS |
| Volume of trabecular bone | 4.11E-03 | NS |
| Gastric adenocarcinoma | 4.27E-03 | NS |
| Myeloid neoplasm | 4.28E-03 | NS |
| Skin squamous cell carcinoma | 4.30E-03 | NS |
| Gastro-esophageal carcinoma | 4.32E-03 | NS |
| Malignant neoplasm of large intestine | 4.32E-03 | NS |
| Hyperesthesia | 4.33E-03 | NS |
| Amyloidosis | 4.49E-03 | NS |
| Transport of metal | 4.56E-03 | NS |
| Advanced malignant tumor | 4.70E-03 | NS |
| AMPA mediated synaptic current | 4.87E-03 | NS |
| Endometrial adenocarcinoma | 4.89E-03 | NS |
| Transport of metal ion | 5.04E-03 | NS |
| Breast or colorectal cancer | 5.21E-03 | NS |
| Undifferentiated sarcoma | 5.22E-03 | NS |
| Organization of cytoskeleton | 5.40E-03 | NS |
| Malignant myeloid neoplasm | 5.47E-03 | NS |
| Drug seeking behavior | 5.64E-03 | NS |
| Isometric tension | 5.64E-03 | NS |
| Uterine cancer | 5.73E-03 | NS |

| | | |
|---|---|---|
| Carcinoma | 5.87E-03 | NS |
| Lymphoma | 5.98E-03 | NS |
| Neuritogenesis | 6.31E-03 | NS |
| Metastasis | 6.35E-03 | NS |
| Organization of neurons | 6.45E-03 | NS |
| Renal cancer | 6.46E-03 | NS |
| Progressive neuromuscular disease | 6.49E-03 | NS |
| Development of mesenchyme | 6.64E-03 | NS |
| G2/M phase transition | 6.64E-03 | NS |
| Parkinson's disease | 6.64E-03 | NS |
| Abnormality of endometrium | 6.92E-03 | NS |
| Renal cell carcinoma | 7.09E-03 | NS |
| Breast or gastric cancer | 7.10E-03 | NS |
| Cell death of kidney cells | 7.29E-03 | NS |
| Idiopathic generalized epilepsy | 7.29E-03 | NS |
| Length of muscle cells | 7.29E-03 | NS |
| Mammary tumor | 7.29E-03 | NS |
| Quantity of neutrophils | 7.29E-03 | NS |
| Rheumatic disease of joint | 7.29E-03 | NS |
| Small-cell carcinoma | 7.29E-03 | NS |
| Survival of CD8+ T lymphocyte | 7.29E-03 | NS |
| Uterine carcinoma | 7.29E-03 | NS |
| Mental retardation | 7.40E-03 | NS |
| Place preference | 7.42E-03 | NS |
| Endometrioid carcinoma | 7.42E-03 | NS |
| Endometrioid endometrial adenocarcinoma | 7.42E-03 | NS |
| Female genital tract cancer | 7.42E-03 | NS |
| Morphology of trabecular bone | 7.44E-03 | NS |
| Non-colon gastrointestinal cancer | 7.52E-03 | NS |
| Abdominal carcinoma | 7.63E-03 | NS |
| Apoptosis of embryonic cell lines | 7.63E-03 | NS |
| Cell death of embryonic cell lines | 7.63E-03 | NS |
| Colon carcinoma | 7.63E-03 | NS |
| Pancreatobiliary tumor | 7.63E-03 | NS |
| Pancreatic cancer | 7.67E-03 | NS |
| Female genital tract adenocarcinoma | 7.68E-03 | NS |
| Colorectal cancer | 7.71E-03 | NS |
| Quantity of granulomonocytic cells | 7.71E-03 | NS |
| Abdominal cancer | 7.77E-03 | NS |
| Neurodegeneration of brain | 7.77E-03 | NS |
| Digestive system cancer | 7.78E-03 | NS |
| Endometrial cancer | 7.78E-03 | NS |
| Liver tumor | 8.01E-03 | NS |
| Abdominal adenocarcinoma | 8.13E-03 | NS |
| Alzheimer disease | 8.13E-03 | NS |
| Morphology of osteoclasts | 8.13E-03 | NS |
| Formation of solid tumor | 8.15E-03 | NS |
| Female genital carcinoma | 8.15E-03 | NS |
| T-cell non-Hodgkin lymphoma | 8.15E-03 | NS |
| Neuromuscular disease with neuropathy | 8.45E-03 | NS |
| Long term synaptic depression | 8.64E-03 | NS |
| Breast or ovarian cancer | 8.64E-03 | NS |
| Pervasive developmental disorder | 8.64E-03 | NS |
| Thickness of cortical bone | 8.64E-03 | NS |
| Breast cancer | 8.72E-03 | NS |
| Depressive disorder | 8.85E-03 | NS |
| Uterine serous papillary cancer | 8.86E-03 | NS |
| Hematologic cancer of cells | 8.88E-03 | NS |
| Pancreatic carcinoma | 8.96E-03 | NS |
| Microtubule dynamics | 9.01E-03 | NS |
| Enlargement of cells | 9.08E-03 | NS |
| Neoplasia of leukocytes | 9.08E-03 | NS |
| Lymphocytic cancer | 9.25E-03 | NS |
| Malignant lymphocytic neoplasm | 9.25E-03 | NS |
| Development of digestive organ tumor | 9.32E-03 | NS |
| Cellular infiltration by leukocytes | 9.50E-03 | NS |
| Generation of tumor | 9.90E-03 | NS |
| Breast or pancreatic cancer | 1.01E-02 | NS |
| Transport of molecule | 1.01E-02 | NS |
| Organization of muscle cells | 1.03E-02 | NS |
| Sprouting | 1.06E-02 | NS |
| Female genital tract serous carcinoma | 1.07E-02 | NS |

| | | |
|---|---|---|
| Laryngeal squamous cell carcinoma | 1.07E-02 | NS |
| Pelvic serous carcinoma | 1.07E-02 | NS |
| Familial neurological disorder | 1.07E-02 | NS |
| Apoptosis of thymocytes | 1.10E-02 | NS |
| Colon adenocarcinoma | 1.10E-02 | NS |
| Development of muscle tissue | 1.10E-02 | NS |
| Pancreatic adenocarcinoma | 1.10E-02 | NS |
| Cell death of kidney cell lines | 1.10E-02 | NS |
| Breast carcinoma | 1.10E-02 | NS |
| Formation of muscle | 1.10E-02 | NS |
| Gastric carcinoma | 1.10E-02 | NS |
| Hyperalgesia | 1.10E-02 | NS |
| Cognition | 1.10E-02 | NS |
| Brain cancer | 1.11E-02 | NS |
| Morphology of nervous system | 1.11E-02 | NS |
| Esophageal carcinoma | 1.11E-02 | NS |
| Opioid-related disorder | 1.11E-02 | NS |
| Size of bone | 1.11E-02 | NS |
| Quantity of osteoclasts | 1.11E-02 | NS |
| Acute myeloid leukemia | 1.12E-02 | NS |
| Advanced extracranial solid tumor | 1.12E-02 | NS |
| Drug dependence | 1.12E-02 | NS |
| Arrest in G2 phase of fibroblasts | 1.13E-02 | NS |
| Cloning | 1.13E-02 | NS |
| Quantity of megakaryocyte/erythrocyte lineage-restricted progenitor cells | 1.13E-02 | NS |
| Memory | 1.17E-02 | NS |
| Breast or ovarian carcinoma | 1.21E-02 | NS |
| Cell death of epithelial cell lines | 1.21E-02 | NS |
| Autosomal dominant deafness | 1.28E-02 | NS |
| Small cell lung carcinoma | 1.29E-02 | NS |
| Neuroendocrine tumor | 1.30E-02 | NS |
| Hepatobiliary carcinoma | 1.30E-02 | NS |
| Childhood epilepsy | 1.30E-02 | NS |
| Abnormal bone density | 1.32E-02 | NS |
| Lung adenocarcinoma | 1.33E-02 | NS |
| Neurogenesis of brain | 1.33E-02 | NS |
| Early-onset neurological disorder | 1.35E-02 | NS |
| Neuronal cell death | 1.35E-02 | NS |
| Upper gastrointestinal carcinoma | 1.35E-02 | NS |
| Advanced malignant solid tumor | 1.36E-02 | NS |
| Pain of muscle | 1.40E-02 | NS |
| Morphology of muscle | 1.40E-02 | NS |
| Visceral metastasis | 1.40E-02 | NS |
| Familial mental retardation | 1.40E-02 | NS |
| Pedal edema | 1.41E-02 | NS |
| Mechanical allodynia behavior | 1.41E-02 | NS |
| Mood Disorders | 1.41E-02 | NS |
| Advanced malignant gastrointestinal neoplasm | 1.45E-02 | NS |
| Differentiation of connective tissue cells | 1.46E-02 | NS |
| Squamous-cell carcinoma | 1.46E-02 | NS |
| Abnormal morphology of brain | 1.48E-02 | NS |
| Urological disorder of kidney | 1.48E-02 | NS |
| Neuroprotection | 1.53E-02 | NS |
| Necrosis | 1.53E-02 | NS |
| Apoptosis of kidney cell lines | 1.54E-02 | NS |
| Development of striated muscle | 1.54E-02 | NS |
| Pancreatic ductal adenocarcinoma | 1.56E-02 | NS |
| Proliferation of endothelioma cell lines | 1.57E-02 | NS |
| Transport of corticosterone | 1.57E-02 | NS |
| Transport of hydrocortisone | 1.57E-02 | NS |
| Synaptic transmission | 1.57E-02 | NS |
| Outgrowth of neuroblastoma cell lines | 1.57E-02 | NS |
| Long-term potentiation | 1.57E-02 | NS |
| Maturation of cells | 1.57E-02 | NS |
| Cellular infiltration by phagocytes | 1.57E-02 | NS |
| Accumulation of cells | 1.57E-02 | NS |
| Extraintestinal functional disorder | 1.57E-02 | NS |
| Abnormal morphology of nervous system | 1.57E-02 | NS |
| Abnormality of cerebral cortex | 1.57E-02 | NS |
| Adenocarcinoma in right upper lobe of lung | 1.57E-02 | NS |
| Aggregation of squamous carcinoma cells | 1.57E-02 | NS |
| Aggressive behavior toward mice | 1.57E-02 | NS |

| | | |
|---|---|---|
| Amyloidosis cutis dyschromia | 1.57E-02 | NS |
| Apoptosis of bronchiolar cell | 1.57E-02 | NS |
| Apoptosis of mesangial cells | 1.57E-02 | NS |
| Arrest in differentiation of macrophages | 1.57E-02 | NS |
| Arrest in differentiation of myelomonocytic cells | 1.57E-02 | NS |
| Arrest in differentiation of neuroblastoma cells | 1.57E-02 | NS |
| Arrest in function of multipotential hemopoietic progenitor cells | 1.57E-02 | NS |
| Arrest in G1/S phase transition of embryonic stem cells | 1.57E-02 | NS |
| Arrest in G2 phase of colorectal cancer cell lines | 1.57E-02 | NS |
| Arrest in maturation of megakaryocytes | 1.57E-02 | NS |
| Arrest in migration of facial branchiomotor neurons | 1.57E-02 | NS |
| Arthrogryposis multiplex congenita-6 | 1.57E-02 | NS |
| Autoimmune lymphoproliferative syndrome type 2B | 1.57E-02 | NS |
| Autosomal dominant deafness type 37 | 1.57E-02 | NS |
| Autosomal dominant deafness type 68 | 1.57E-02 | NS |
| Autosomal dominant encephalopathy | 1.57E-02 | NS |
| Autosomal dominant mental retardation type 6 | 1.57E-02 | NS |
| Autosomal recessive mental retardation type 37 | 1.57E-02 | NS |
| Autosomal recessive mental retardation type 54 | 1.57E-02 | NS |
| Axonogenesis | 1.57E-02 | NS |
| Benign pelvic disease | 1.57E-02 | NS |
| Benign uterine disease | 1.57E-02 | NS |
| Biliary tract carcinoma | 1.57E-02 | NS |
| Biogenesis of lateral plasma membrane | 1.57E-02 | NS |
| Bipolar disorder | 1.57E-02 | NS |
| Blood pressure of corpus cavernosum penis | 1.57E-02 | NS |
| Brachydactyly type E2 | 1.57E-02 | NS |
| Brain astrocytoma | 1.57E-02 | NS |
| Brain malformation and urinary tract defect | 1.57E-02 | NS |
| Cecum adenocarcinoma | 1.57E-02 | NS |
| Cell death of myelomonocytic cells | 1.57E-02 | NS |
| Cell death of progenitor cells | 1.57E-02 | NS |
| Cell rolling of blood platelets | 1.57E-02 | NS |
| Cell rolling of pre-B lymphocytes | 1.57E-02 | NS |
| Cell spreading of mesenchymal cells | 1.57E-02 | NS |
| Chemokinesis of embryonic cell lines | 1.57E-02 | NS |
| Chemokinesis of epithelial cell lines | 1.57E-02 | NS |
| Chromosome 1p32-p31 deletion syndrome | 1.57E-02 | NS |
| Chronic colitis | 1.57E-02 | NS |
| Clearance of doxorubicin | 1.57E-02 | NS |
| Clearance of SN-38 | 1.57E-02 | NS |
| Climbing ability | 1.57E-02 | NS |
| Colchicine resistance | 1.57E-02 | NS |
| Colony formation of osteoprogenitor cells | 1.57E-02 | NS |
| Congenital muscular dystrophy due to partial merosin deficiency | 1.57E-02 | NS |
| Cranial chondrodystrophy | 1.57E-02 | NS |
| Cytokinesis of B cell lymphoma cells | 1.57E-02 | NS |
| Delay in acidification of gonadal cell lines | 1.57E-02 | NS |
| Delay in initiation of development of oligodendrocytes | 1.57E-02 | NS |
| Delay in initiation of turnover of epidermal cells | 1.57E-02 | NS |
| Delay in production of endocrine cells | 1.57E-02 | NS |
| Delay in tubulation of epithelial cells | 1.57E-02 | NS |
| Dentin dysplasia with extreme microdontia and misshapen teeth type I | 1.57E-02 | NS |
| Depolarization-induced suppression of excitation of hippocampal neurons | 1.57E-02 | NS |
| Differentiation of periosteal cells | 1.57E-02 | NS |
| Discomfort | 1.57E-02 | NS |
| Dissociation of embryonic stem cells | 1.57E-02 | NS |
| Drug resistance of lung cell lines | 1.57E-02 | NS |
| Duane retraction syndrome 3 with deafness | 1.57E-02 | NS |
| Dyskinesia of axial skeleton | 1.57E-02 | NS |
| Early infantile epileptic encephalopathy type 26 | 1.57E-02 | NS |
| Early infantile epileptic encephalopathy type 27 | 1.57E-02 | NS |
| Early infantile epileptic encephalopathy type 62 | 1.57E-02 | NS |
| Efficacy of etoposide | 1.57E-02 | NS |
| Efficacy of vincristine | 1.57E-02 | NS |
| Efflux of cyclosporin A | 1.57E-02 | NS |
| Efflux of digoxin | 1.57E-02 | NS |
| Efflux of fexofenadine | 1.57E-02 | NS |
| Efflux of vincristine | 1.57E-02 | NS |
| Elimination of osteocytes | 1.57E-02 | NS |
| Entry into S phase of mesothelioma cell lines | 1.57E-02 | NS |
| Epileptic seizure | 1.57E-02 | NS |

| | | |
|---|---|---|
| Excitotoxic lesion | 1.57E-02 | NS |
| Extension of parallel fiber | 1.57E-02 | NS |
| Extrapancreatic neuroendocrine tumor | 1.57E-02 | NS |
| Familial focal epilepsy with variable foci 4 | 1.57E-02 | NS |
| Familial multicentric carpotarsal osteolysis syndrome | 1.57E-02 | NS |
| Fear | 1.57E-02 | NS |
| Flipping of glucosylceramide | 1.57E-02 | NS |
| Formation of sling cells | 1.57E-02 | NS |
| Gait disturbance | 1.57E-02 | NS |
| Gliomatosis cerebri | 1.57E-02 | NS |
| Glucocorticoid-resistant inflammatory bowel disease | 1.57E-02 | NS |
| Grade 4 astrocytoma | 1.57E-02 | NS |
| Grade 4 high grade glioma | 1.57E-02 | NS |
| Hepatobiliary system cancer | 1.57E-02 | NS |
| High grade astrocytoma | 1.57E-02 | NS |
| Homeostasis of peripheral T lymphocyte | 1.57E-02 | NS |
| Inflammatory bowel disease type 13 | 1.57E-02 | NS |
| Intellectual developmental disorder with gastrointestinal difficulties and high pain threshold | 1.57E-02 | NS |
| Isometric tension of myofiber | 1.57E-02 | NS |
| Keppen-Lubinsky syndrome | 1.57E-02 | NS |
| Killing of pancreatic cancer cell lines | 1.57E-02 | NS |
| Lack of blood platelets | 1.57E-02 | NS |
| Length of apical membrane | 1.57E-02 | NS |
| Length of basal membrane | 1.57E-02 | NS |
| Length of smooth muscle cells | 1.57E-02 | NS |
| Lesioning of renal glomerulus | 1.57E-02 | NS |
| Lipogenesis by breast cell lines | 1.57E-02 | NS |
| Locally advanced nasopharyngeal squamous cell carcinoma | 1.57E-02 | NS |
| Loss of atrioventricular canal | 1.57E-02 | NS |
| Loss of bone marrow-derived dendritic cells | 1.57E-02 | NS |
| Loss of lateral plasma membrane | 1.57E-02 | NS |
| Lung squamous cell carcinoma | 1.57E-02 | NS |
| Lysis of germ cell tumor cell lines | 1.57E-02 | NS |
| Marshall/Stickler syndrome | 1.57E-02 | NS |
| Mature T-cell neoplasm | 1.57E-02 | NS |
| Merosin-negative congenital muscular dystrophy | 1.57E-02 | NS |
| Metabolism of alpha-tocopherol | 1.57E-02 | NS |
| Metastatic solid tumor | 1.57E-02 | NS |
| Migration of sling cells | 1.57E-02 | NS |
| Morphology of bone | 1.57E-02 | NS |
| Morphology of brain cells | 1.57E-02 | NS |
| Morphology of cerebral cortex | 1.57E-02 | NS |
| Mucoid diarrhea | 1.57E-02 | NS |
| Muscle spasticity of hindlimb | 1.57E-02 | NS |
| Muscle tumor | 1.57E-02 | NS |
| Nausea | 1.57E-02 | NS |
| Necroptosis of T lymphocytes | 1.57E-02 | NS |
| Nemaline myopathy type 2 | 1.57E-02 | NS |
| Neurogenesis of subventricular zone | 1.57E-02 | NS |
| Organization of synapse | 1.57E-02 | NS |
| Pancreatobiliary adenocarcinoma | 1.57E-02 | NS |
| Penetration of dexamethasone | 1.57E-02 | NS |
| Permeability transition of breast cancer cell lines | 1.57E-02 | NS |
| Physical stamina | 1.57E-02 | NS |
| Polyploidization of breast cancer cell lines | 1.57E-02 | NS |
| Polyploidization of carcinoma cell lines | 1.57E-02 | NS |
| Polyploidization of lung cancer cell lines | 1.57E-02 | NS |
| Presence of corpus callosum | 1.57E-02 | NS |
| Progressive early-onset ataxia | 1.57E-02 | NS |
| Proliferation of astrocytoma cells | 1.57E-02 | NS |
| Quantity of apoptotic intestinal cell lines | 1.57E-02 | NS |
| Quantity of bone | 1.57E-02 | NS |
| Quantity of femur | 1.57E-02 | NS |
| Quantity of hematopoietic progenitor cells | 1.57E-02 | NS |
| Quantity of intramyocellular lipid store | 1.57E-02 | NS |
| Quantity of myocardium | 1.57E-02 | NS |
| Rectosigmoid adenocarcinoma | 1.57E-02 | NS |
| Recurrent adult acute myeloid leukemia | 1.57E-02 | NS |
| Reinitiation of mitosis of breast cancer cell lines | 1.57E-02 | NS |
| Reinitiation of mitosis of carcinoma cell lines | 1.57E-02 | NS |
| Reinitiation of mitosis of lung cancer cell lines | 1.57E-02 | NS |
| Release of C6-NBD-glucosylceramide | 1.57E-02 | NS |

| | | |
|---|---|---|
| Release of C6-NBD-PE | 1.57E-02 | NS |
| Release of N-(1-hexanoyl)-D-erythro-glucosylsphingosine | 1.57E-02 | NS |
| Secretion of catecholamine | 1.57E-02 | NS |
| Single suture craniosynostosis | 1.57E-02 | NS |
| Stage III-IVB sinonasal squamous cell carcinoma | 1.57E-02 | NS |
| Stickler syndrome type 2 | 1.57E-02 | NS |
| Suppression of GABA | 1.57E-02 | NS |
| Survival of brain tissue | 1.57E-02 | NS |
| Survival of oocytes | 1.57E-02 | NS |
| Survival of tubular cells | 1.57E-02 | NS |
| Syringomyelia | 1.57E-02 | NS |
| Thrombocytosis | 1.57E-02 | NS |
| Tonic-clonic seizure | 1.57E-02 | NS |
| Toxicity of nevirapine | 1.57E-02 | NS |
| Translocation of etoposide | 1.57E-02 | NS |
| Translocation of mitoxantrone | 1.57E-02 | NS |
| Translocation of topotecan | 1.57E-02 | NS |
| Translocation of vinblastine | 1.57E-02 | NS |
| Translocation of vincristine | 1.57E-02 | NS |
| Transport of amiodarone | 1.57E-02 | NS |
| Transport of platelet activating factor | 1.57E-02 | NS |
| Tricuspid valve stenosis | 1.57E-02 | NS |
| Tumorigenesis of lymphocytes | 1.57E-02 | NS |
| Upper aerodigestive tract carcinoma | 1.57E-02 | NS |
| Uptake of dexamethasone | 1.57E-02 | NS |
| Uptake of hydrocortisone | 1.57E-02 | NS |
| Vasodilation of saphenous artery | 1.57E-02 | NS |
| Ventricular arrhythmias due to cardiac ryanodine receptor calcium release deficiency syndro | 1.57E-02 | NS |
| Head and neck tumor | 1.58E-02 | NS |
| Connective or soft tissue tumor | 1.58E-02 | NS |
| Neuromuscular disease | 1.58E-02 | NS |
| Memory consolidation | 1.65E-02 | NS |
| Upper aero-digestive squamous cell carcinoma | 1.65E-02 | NS |
| Brain lesion | 1.66E-02 | NS |
| Apoptosis of muscle cells | 1.66E-02 | NS |
| Major affective disorder | 1.67E-02 | NS |
| Long term synaptic depression of neurons | 1.69E-02 | NS |
| Upper airway cancer | 1.69E-02 | NS |
| Degradation of DNA | 1.71E-02 | NS |
| Cellular infiltration by myeloid cells | 1.72E-02 | NS |
| Cell death of keratinocytes | 1.72E-02 | NS |
| Neurogenesis of dentate gyrus | 1.72E-02 | NS |
| Signaling of lymphocytes | 1.72E-02 | NS |
| Degeneration of cells | 1.74E-02 | NS |
| Complex neurodevelopmental disorder | 1.80E-02 | NS |
| Morphology of trabecula | 1.81E-02 | NS |
| Multiple Sclerosis | 1.81E-02 | NS |
| Inflammation of the large intestine | 1.87E-02 | NS |

*: Also significantly enriched in IPA analysis with all PE-dysregulated miR-29a/c-3p target genes; NS: Not statistically Significant.

**Table S7. PE dysregulated miR-29a/c-3p target gene network in female and male P0-HUVECs**

| Gene network | PE vs. NT Female P0-HUVECs | | | PE vs. NT Male P0-HUVECs | | |
|---|---|---|---|---|---|---|
| | P-value | Differentially Expressed miR-29a/c-3p Target Genes | # of Genes | P-value | Differentially Expressed miR-29a/c-3p Target Genes | # of Genes |
| TNF-Regulated Genes | 1.35E-14 | A4GALT, AKR1B10, ANGPTL4, ANK3, ARC, BCL3, BDKRB2, CASP8, CCL3, CCL3L3, CHRNA7, CLDN5, CSF2, CX3CL1, CXCL11, F3, FASN, G0S2, HAS3, INSIG1, LAMC2, LAMP3, LIF, MAP2K6, NEFM, NLRP3, PMAIP1, PPP1R3C, PTHLH, RGS4, SELPLG, SLC15A3, SLC7A2, SREBF1, TNFAIP3, TNFAIP6, TNIP1, UGCG | 38 | 3.36E-04 | ARC, CA2, CCN4, CHRNA7, CLEC2D, COL1A1, DUSP2, HMOX1, NEFM | 9 |
| TGFB1-Regulated Genes | 1.81E-13 | ANGPTL4, ARC, ASNS, AXL, BCL3, BDKRB2, CASP8, CCL3, CCL3L3, CHRNA7, COL11A1, COL4A1, COL4A2, CSF2, CX3CL1, F3, FASN, GRIN2B, HAS3, IFI30, INSIG1, LAMC2, LIF, LPAR1, MYBL2, NLRP3, PPP1R3C, PSAT1, PTHLH, RSAD2, SELPLG, SMOC2, TNFAIP3, TNFAIP6, TRIB3, TRIM9 | 36 | 7.83E-07 | ARC, CCN4, CDKN1C, CHRNA7, CLEC2D, COL1A1, DNAJA1, ELN, GRIN2B, HMOX1, MAOA, WNT11 | 12 |
| IFNG-Regulated Genes | 2.39E-10 | ABCB1, ADAMTS9, ANGPTL4, ARC, ASNS, BCL3, CASP8, CCL3, CCL3L3, CNR1, CSF2, CX3CL1, CXCL11, F3, FASN, IFI30, ISG20, LAMC2, LAMP3, LIF, NEFM, NLRP3, PMAIP1, PTHLH, RSAD2, SLC15A3, SREBF1, TNFAIP6 | 28 | 1.65E-03 | ARC, CLEC2D, COL1A1, ELN, H2BC18, HMOX1, NEFM | 7 |
| IL1B-Regulated Genes | 4.06E-08 | A4GALT, ANGPTL4, ARC, BCL3, CCL3, CCL3L3, CSF2, CX3CL1, CXCL11, F3, G0S2, ISG20, LAMC2, LIF, MAP2K6, PTHLH, RSAD2, SREBF1, TNFAIP3, TNFAIP6, UGCG | 21 | 4.10E-02 | ARC, COL1A1, ELN, HMOX1 | 4 |
| NFkB-Regulated Genes | 1.06E-11 | ABCB1, ADAMTS9, BCL3, CASP8, CCL3, CCL3L3, CSF2, CX3CL1, CXCL11, F3, G0S2, LIF, PMAIP1, PPM1D, RSAD2, SELPLG, TNFAIP3, TNFAIP6, TNIP1, TRIB3 | 20 | - | - | - |
| MYC-Regulated Genes | 2.63E-04 | ANGPTL4, ASNS, AXL, CASP8, COL4A1, COL4A2, CXCL11, F3, FASN, FSTL1, ISG20, PMAIP1, PSAT1, PTHLH, RSAD2, SREBF1 | 16 | 3.25E-03 | CBX4, CCN4, COL1A1, DUSP2, EPHB3, HMOX1 | 6 |
| AGT-Regulated Genes | 2.71E-05 | ANK3, BDKRB2, CASP8, CCL3L3, CNR1, COL4A1, COL4A2, F3, FASN, GFOD1, LIF, MAFB, PPP1R3C, PTHLH, UGCG | 15 | 6.55E-04 | CCN4, CDKN1C, COL1A1, ELN, HMOX1, NREP | 6 |
| IL33-Regulated Genes | 5.62E-07 | AXL, BCL3, CCL3, CCL3L3, CSF2, CX3CL1, FER1L6, G0S2, NLRP3, PSAT1, SELPLG, TNFAIP3 | 12 | - | - | - |
| FOXO1-Regulated Genes | 1.36E-05 | ANGPTL4, CASP8, CLDN5, CNR1, COL4A1, COL4A2, FASN, PDK4, PMAIP1, SREBF1, TRIB3 | 11 | 2.24E-03 | CA2, CDKN1C, COL1A1, HMOX1 | 4 |
| MAPK1-Regulated Genes | 1.60E-06 | ANGPTL4, ARC, CCL3L3, CXCL11, GFOD1, IFI30, ISG20, LAMP3, LIF, MYBL2, PSAT1 | 11 | 9.93E-03 | ARC, DUSP2, HMOX1 | 3 |
| PDGF BB-Regulated Genes | 6.67E-07 | BCL3, BDKRB2, CSF2, F3, FASN, HOMER2, LIF, MYBL2, SREBF1, TNFAIP3, TRIB3 | 11 | 7.78E-03 | CBX4, COL1A1, HMOX1 | 3 |
| F2-Regulated Genes | 8.76E-07 | ADAMTS9, ANGPTL4, BCL3, COL4A1, CX3CL1, DHRS9, F3, MYBL2, PMAIP1, TNFAIP3 | 10 | 4.43E-02 | COL1A1, HMOX1 | 2 |
| IL1-Regulated Genes | 6.07E-06 | CCL3, CSF2, CXCL11, F3, LIF, MAP2K6, SREBF1, TNFAIP3, TNFAIP6, UGCG | 10 | - | - | - |
| CSF2-Regulated Genes | 1.86E-04 | BCL3, CCL3, CCL3L3, CSF2, F3, LAMP3, NFE2, NLRP3, SREBF1, TNFAIP3 | 10 | 2.42E-02 | DUSP2, ELN, HMOX1 | 3 |

### Supplemental Figures with Figure Legends

**Figure S1. Overexpression efficiency of miR29a-3p(+) (A) and miR29c-3p(+) (B) on miR29a/c-3p expression in HUVECs.** Sub-confluent HUVECs were transfected with miR-29a-3p(+), miR-29c-3p(+), Negative control (Neg.Ctrl), or vehicle Control (Veh). The miRNA levels were determined using miRNA RT-qPCR. Data normalized to housekeeping genes are expressed as medians ±SEM fold of Veh at each corresponding time point. * Differ from negative control at corresponding time. ($P < 0.05$; Mann-Whitney Rank Sum Test; n=4)

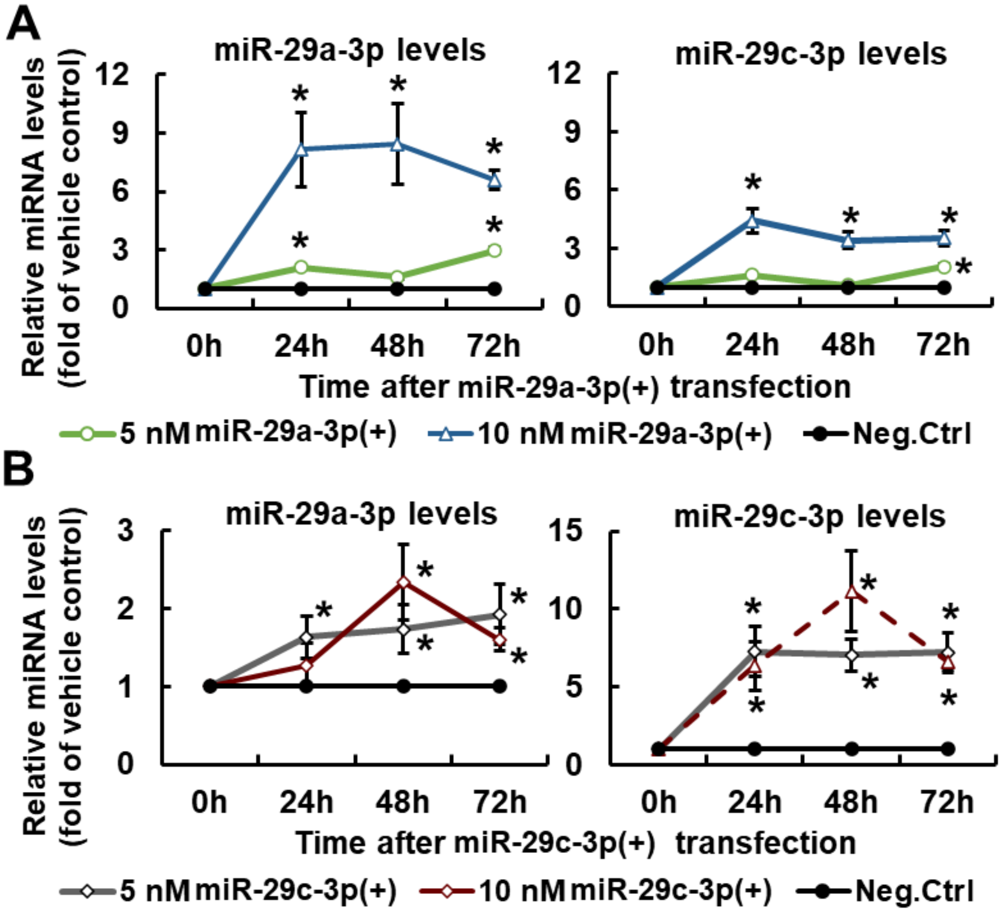

**Figure S2. Key canonical pathways enriched in preeclampsia upregulated (A) and downregulated (B) miR-29a/c-3p target genes in female and male P0-HUVECs.** Significant enrichments were determined using IPA software (P < 0.05, Fisher's exact test followed with BH-FDR multiple test correction). *: Canonical pathways also significantly enriched in overall preeclampsia-dysregulated miR-29a/c-3p target genes in Figure 2D.

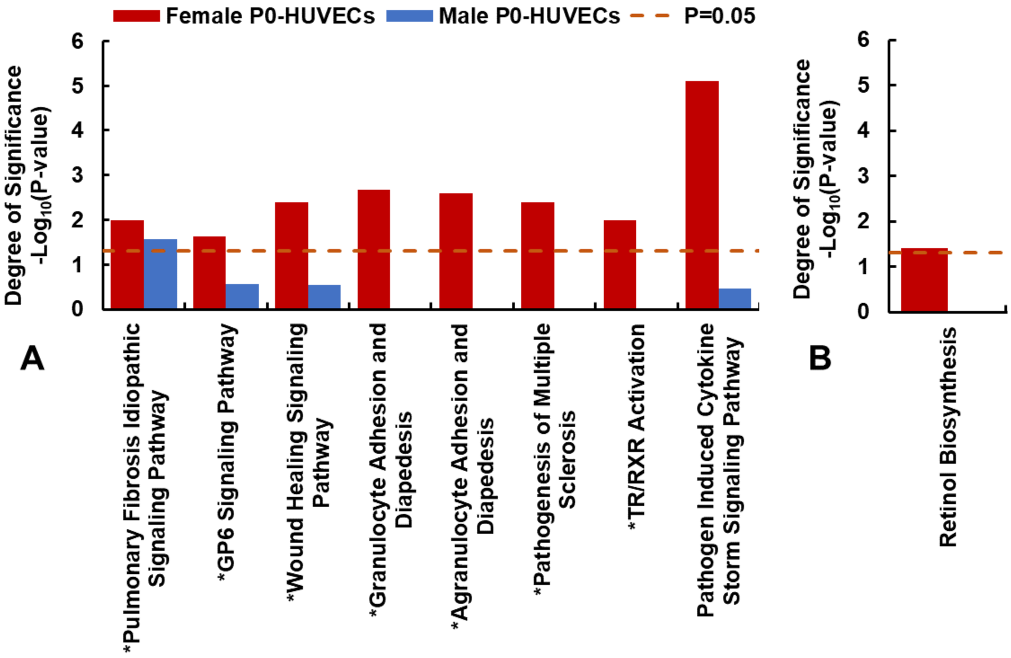

**Figure S3. Key diseases and biological functions enriched in preeclampsia upregulated (A) and downregulated (B) miR-29a/c-3p target genes in female and male P0-HUVECs.** Significant enrichments were determined using IPA software (P < 0.05, Fisher's exact test followed with BH-FDR multiple test correction). *: Diseases and biological functions also significantly enriched in overall preeclampsia-dysregulated miR-29a/c-3p target genes in figure 2E.

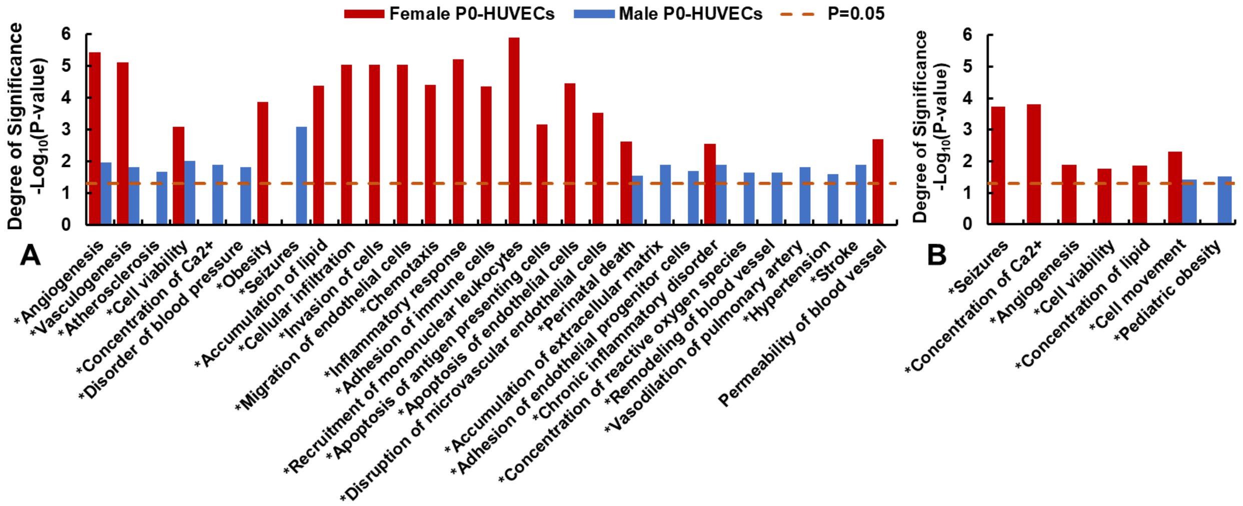

**Figure S4. MiR29-a/c-3p do not alter endothelial monolayer integrity in response to TNFα in female and male P1-HUVECs from NT and PE.** Cells were transfected with miR-29a/c-3p(+) or miR29-a/c-3p(i), and then cultured until confluence (~30h). After 6-8h of serum starvation, confluent cells were treated with ECMb (serum-free control) or TNFα (10ng/ml) for 25h. Electrical resistance at 4000Hz was constantly recorded. Data are expressed as medians ± SEM fold of Vehicle control treated with ECMb at corresponding time (n=5-10 cell preparations/sex/group).

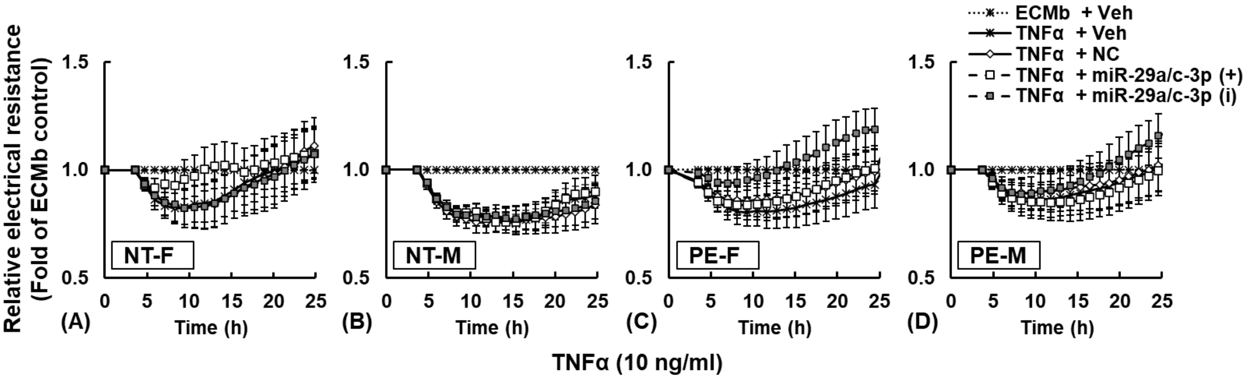
